# Genomic diversity of *Helicobacter pylori* populations from different regions of the human stomach

**DOI:** 10.1101/2022.01.18.476812

**Authors:** D.J. Wilkinson, B. Dickins, K. Robinson, J. Winter

## Abstract

Individuals infected with *Helicobacter pylori* harbour unique and diverse populations of quasispecies, but diversity between and within different regions of the human stomach and the process of bacterial adaptation to each location are not yet well understood.

We applied whole-genome deep sequencing to characterise the within- and between- stomach region genetic diversity *of H. pylori* populations from paired antrum and corpus biopsies of 15 patients, along with single biopsies from one region of 3 patients, by scanning allelic diversity. We combined population deep sequencing with more conventional sequencing of multiple *H. pylori* single colony isolates from individual biopsies to generate a unique dataset. Single colony isolates were used to validate the scanning allelic diversity pipelines.

We detected extensive population allelic diversity within the different regions of each patient’s stomach. Diversity was most commonly found within non-coding, hypothetical, outer membrane, restriction modification system, virulence, lipopolysaccharide biosynthesis, efflux systems and chemotaxis-associated genes. Antrum and corpus populations from the same patient grouped together phylogenetically, indicating that most patients were initially infected with a single strain, which then diversified. Single colonies from the antrum and corpus of the same patients grouped into distinct clades, suggesting mechanisms for within-location adaptation across multiple *H. pylori* isolates from different patients. Recombination was observed both within and between different regions of the same stomach.

The comparisons made available by combined sequencing and analysis of isolates and populations enabled comprehensive analysis of the genetic changes associated with *H. pylori* diversification and stomach region adaptation.

## Introduction

*Helicobacter pylori* typically first colonises people in early childhood and then persists as a chronic, lifelong infection^1^. This usually results in gastritis, which is most often asymptomatic in nature^2^. The chronic inflammatory nature of the infection, and the high mutation and recombination rate of this bacterium, are thought to contribute to a diverse bacterial population (quasispecies) within the infected gastric mucosa. Colonising bacteria in the antrum and corpus regions of the human stomach will be exposed to different levels of acidity, inflammatory factors, access to sheltering glands and mucus. This variation in environmental conditions within stomachs may drive bacterial diversification over time. Particularly in high-prevalence areas, humans are colonised by multiple strains^3^, further increasing this diversity. The high level of genetic diversity of *H. pylori* has implications for the design of successful eradication regimens and vaccines. However, the extent and characteristics of *H. pylori* diversity within infected individuals are not yet fully understood.

Genome sequences of *H. pylori* isolates across the world have revealed a global population structure and diversity^4–8^. Comparative genomics approaches have been taken, usually with single colony isolates from each patient, to investigate global population diversity in *H. pylori*. Other studies have characterised *H. pylori* strains from specific geographical regions^9, 10^. These global and geographical genetic analyses have facilitated the identification of genotypes associated with different disease types and severity. For example, a recent genome wide association study^11^ identified SNPs and genes in *H. pylori* genomes that could be used to assess gastric cancer risk in infected people. A similar study identified six genes that were associated with peptic ulcer disease and gastric adenocarcinoma, by comparing multi-ethnic populations at high to low risk of these diseases^12^. Therefore, despite there being a very high level of diversity in *H. pylori* genomes, analysis at global and local geographical levels has provided important insights into bacterial virulence and disease progression.

Studies of bacterial populations that rely on sequencing single colony isolates can only capture a subset of the genetic diversity, but bacterial diversity at population level can be more comprehensively assessed using deep sequencing methodologies. Within-patient genetic diversity of *Burkholderia dolosa* from chronically infected individuals was previously investigated using a population deep sequencing approach^13^, where the reads mapped onto a reference genome identified the most variable genes and regions, and captured a snapshot of within-patient genetic diversity of the whole population.

For this study, we adopted a similar approach^13^ and adapted it to *H. pylori* populations from gastric biopsies taken from the antrum and corpus of infected patients. Single colonies were also isolated from a subset of these biopsies for conventional whole-genome sequencing. Here we present a high-resolution analysis of *H. pylori* genetic structure and population diversity using these techniques in *H. pylori* populations for the first time.

## Materials and Methods

A summary of the methods used in this study is presented in Fig. 1.

**Fig. 1.**
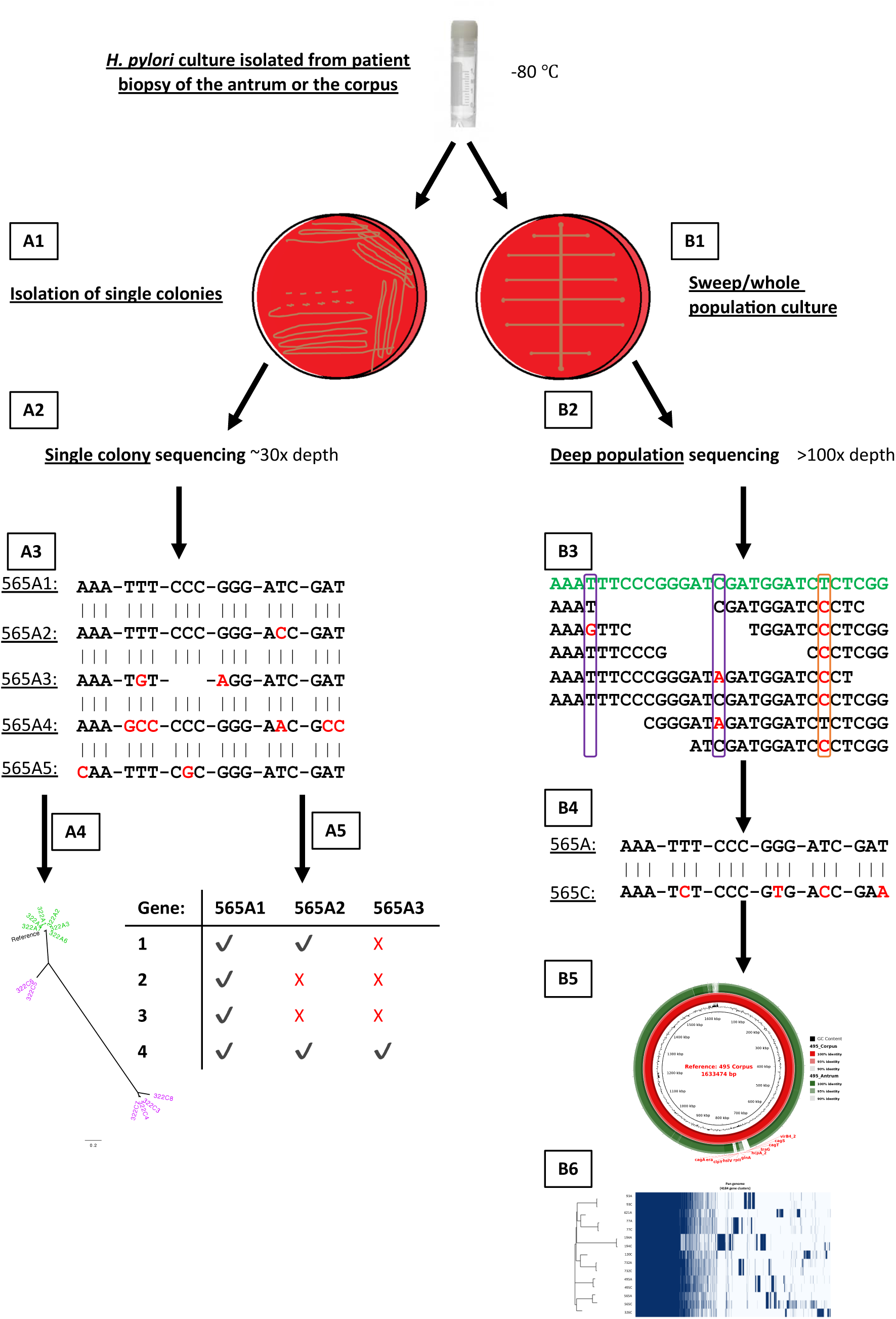
Overview of study design. *H. pylori* isolates from corpus and antrum biopsies were subjected to population deep sequencing (right; B1-6) and streaked out to single colonies which were individually sequenced (left; A1-5). Data from the two approaches were used to analyse *H. pylori* diversity within and between stomach regions and individuals. A1 – isolation of single colony isolates; A2 – single colony isolates sequenced to ∼30x depth; A3 – single colony isolate genomes aligned to investigate between strain diversity with red nucleotide bases representing SNPs; A4 – single colony isolate genomes used to create phylogenetic trees; A5 – gene presence and absence analysis. B1 – culture of *pylori* populations from each biopsy; B2 – population deep sequencing to >100x depth; B3 – within population diversity investigated by mapping reads to the consensus genome (green sequence) with minor (purple boxes) and common (orange box) allelic diversity highlighted; B4 – between stomach region diversity analysed by aligning the consensus genome assemblies from each regions’ population; B5 –BLASTN identity between stomach region populations investigated using the consensus assembled population genomes, highlighting areas of high diversity using BRIG^20^; B6 – pan-genome analysis.

### Bacterial culture

*H. pylori* was grown on blood agar base 2 with 5% horse blood (Oxoid, Basingstoke, UK) for 24–48 h at 37°C under humid, microaerobic conditions. Cultures were stored in iso-sensitest medium containing 15% (v/v) glycerol (Sigma Aldrich, UK) at -80°C.

### Clinical samples

Gastric biopsies were donated by 18 *H. pylori*-infected patients (median age 63.5 years, range 40–79 years; 38.9% male) attending the Queen’s Medical Centre, Nottingham, UK, for routine upper GI tract endoscopy to investigate dyspeptic symptoms. Written informed consent was obtained, and the study was approved by the Nottingham Research Ethics Committee 2 (08/H0408/195). Patients were excluded if taking antibiotics, proton pump inhibitors or >150 mg/day aspirin, 2 weeks preceding the endoscopy. Patients with variable Sydney scores and/or *H. pylori vacA, cagA* and *cagE* genotypes between their antrum and corpus were prioritised for inclusion in this study, which primarily aimed to test the potential power of a combined approach of deep population and single colony sequencing for *H. pylori* isolates. Biopsies were swept across blood base #2 agar plates containing 5% (v/v) horse blood and incubated for 2-3 days at 37°C under microaerophilic conditions (10% CO_2_, 5% O_2_, 85% N_2_). *H. pylori* growth from each biopsy sweep was picked and pooled into isosensitest broth containing 15% (v/v) glycerol for long term storage at -80°C. For single colony isolation, pooled cultures were streaked out to obtain multiple single colony isolates. These were picked at random and passaged one to two times to increase bacterial numbers for long term storage.

### DNA extraction and whole-genome sequencing

Genomic DNA from population sweeps and single colony isolates was extracted using the QIAGEN QIAmp DNA Mini Kit following manufacturer’s instructions. DNA quality was determined using the NanoDrop2000 spectrophotometer using strict absorbance ratio cut-off values of A260/A280 1.8 – 1.9 and A260/A230 1.9 - 2.2. Genomic DNA was quantified using the dsDNA high sensitivity assay on a Qubit v4.0 fluorometer and diluted to 0.3 ng/μl for creation of Nextera XT paired-end libraries. Sequencing used Illumina V3 chemistry cartridges spiked with 1% PhiX DNA run at 2 x 250 cycles on the MiSeq platform.

The raw sequencing data generated in this study were deposited on the NCBI website with the accession code PRJNA787419.

### Sequencing read curation, contamination detection, whole-genome assembly and annotation

All sequencing reads were trimmed for Illumina sequencing adapters and adapter read-through using Trimmomatic^14^ (version 0.38). Reads were further trimmed with Sickle^15^ (version 1.33) to Phred 30 with a minimum read length of 50bp. Trimmed reads were inspected by FastQC^16^ 0.11.7 for confirmation of expected trimming. Curated reads were passed through Kraken^17^ 1.0 and the MiniKraken database (RefSeq complete bacterial, archaeal and viral genomes captured on 18/10/2017) to detect contaminated samples, defined as <92% of curated sequencing reads mapped to *H. pylori* and/or <95% of the reads mapped to the *Helicobacter* genus. Two deep-sequenced samples (308A and 326A) contained contaminant reads from an unknown species. Curated reads were used to construct *de novo* assemblies using the SPAdes^18^ 3.11.1 assembler in careful mode creating consensus whole-genome assemblies for the deep-sequenced clinical sweep samples and contigs <500 bp were removed. Sequencing depth/coverage was determined using mosdepth 0.2.3^19^. Contaminant reads were assembled into separate contigs and excluded from all further analyses with the exception of BLAST ring image generator (BRIG; version 0.95) analysis^20^ (Fig. 2). Consensus population sweeps and single colony assemblies were annotated using Prokka^21^ (version 1.13) guided by the reference *H. pylori* strain 26695 (NC_000915.1) with an e-value threshold of 0.001 to account for gene diversity.

**Fig. 2.**
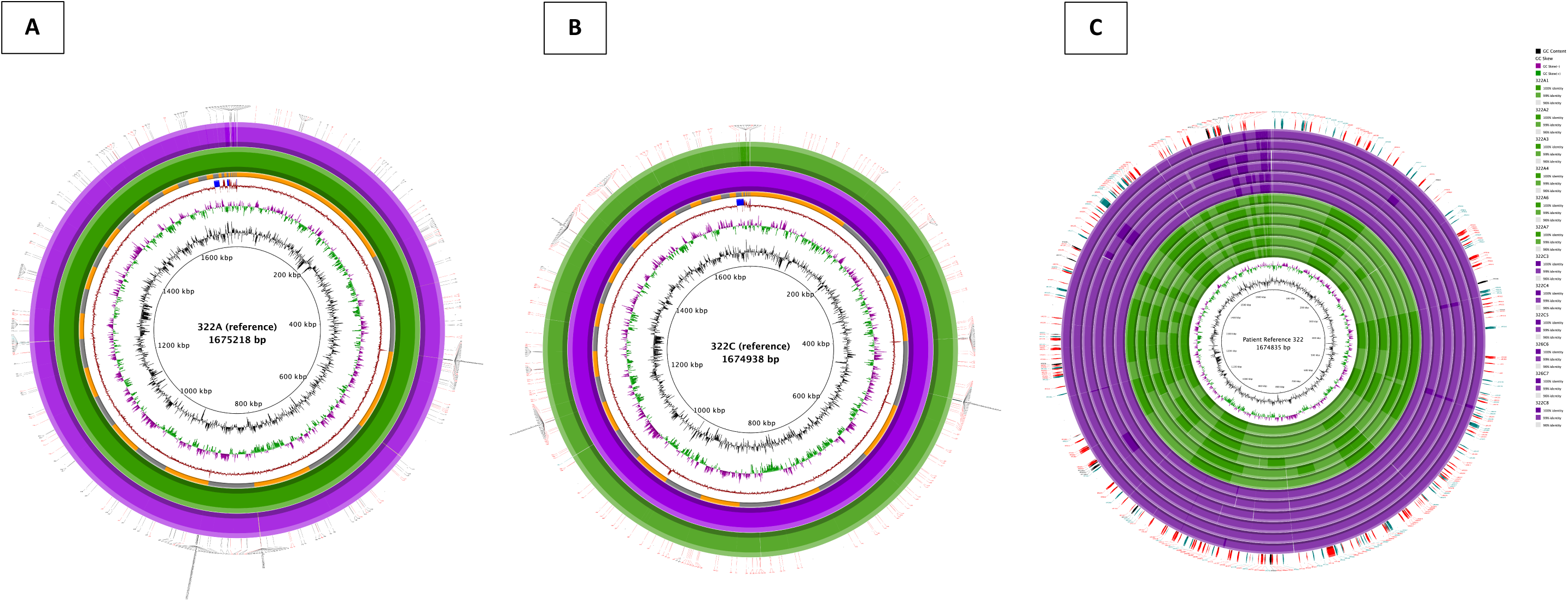
Representative whole-genome alignment for *H. pylori* strains isolated from patient 322. Whole-genome alignments of the consensus genomes generated by deep population sequencing of antrum and corpus *H. pylori* populations, using antrum (A) or corpus (B) consensus genome as the reference. Figure C depicts the alignment of colony isolate assembled genomes against the patient reference. Colour intensity of each ring indicates percentage identity between antrum and corpus consensus genomes. Positions of contigs within the assembled reference genome are shown as a ring alternating in colour between grey and orange in figs A-B to show contig boundaries. Peripheral ticks indicate positions of SNPs between antrum and corpus genomes identified by whole-genome alignment (black), read mapping (teal), or both methods (red) in figures A-B. Alignments for the other patients are shown in Suppl. Fig. 2-15.

### Within-region diversity of deep-sequenced bacterial sweeps

A read mapping and polymorphic detection pipeline was developed similar to that of Lieberman *et al.* (2014)^13^. An overview of the pipeline developed for this study is depicted in Table 1.

**Table 1.**
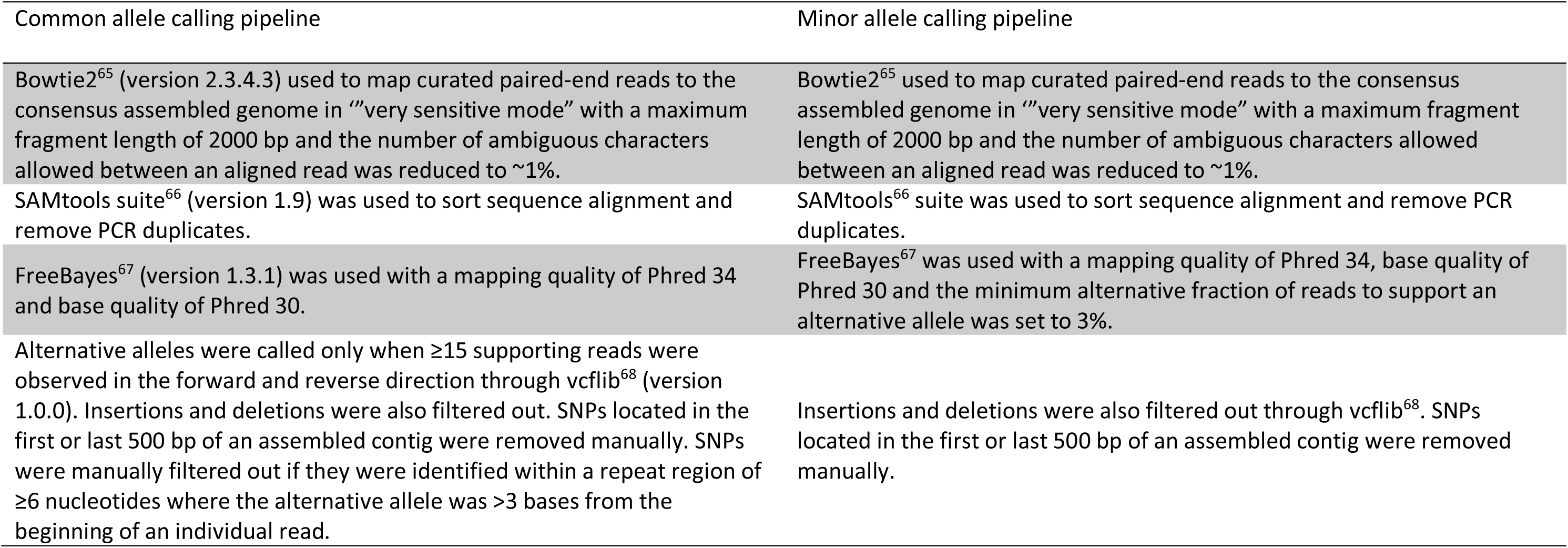
Common and minor allele calling pipelines.

Validation of the minor allele calling pipeline was achieved by mapping the curated single colony sequencing reads of the antrum and the corpus to the corresponding clinical sweep deep-sequenced consensus assembled reference genomes from each region (n=19) using Snippy^22^ 4.4.0 with an alternative allele support >90%, coverage of ≥6, minimum mapping quality of Phred 34 and minimum base quality of Phred 30. Data from 565C were excluded due to the potential presence of different *H. pylori* strains, 565C1 and 565C6, which had an unusually high number of contigs (565C1 – 1,501; 565C6 – 1,295), low N50 statistics (565C1 – 3,665; 565C6 – 3,977) and high genome lengths (565C1 – ∼2.62 Mbp; 565C6 – ∼2.58 Mbp).

Within-region diversity was further investigated using the single colony isolates from each patient and analysed through the BRIG^20^ (Fig. 2). Briefly, the single colony isolate genomes from both the antrum and corpus of each patient were aligned^23^ to the patient reference consensus genome which was assembled as previously described using all deep population sequencing reads from the antrum and corpus datasets and run through BRIG with a lower identity threshold of 96%. Single nucleotide polymorphisms between the aligned genomes were identified and compared to the allelic genes identified by the population minor allele calling pipeline. The locations of variant genes were then highlighted for reference.

### Between-region diversity

BRIG was also used to investigate diversity between stomach regions. Paired antrum and corpus consensus assembled genomes from the deep population sequencing were used, and BLASTN identity was set to 95% to account for more potential diversity in this dataset. The antrum and corpus consensus genomes for each patient were used as the reference sequence, with the alternative region used as the query sequence in order to fully investigate between-region genomic diversity.

Between-region diversity was further investigated using a consensus and read mapping alignment approach depicted in Suppl. Fig. 1.

High confidence alignment SNPs identified by consensus genome alignment and read mapping (Fig. 2) were passed through a custom build of SnpEff^24^ 4.3 to determine whether alignment identified SNPs were synonymous or nonsynonymous.

Core genome phylogenetic trees were constructed from patients with paired antrum and corpus single colony isolates through Snippy^22^ 4.4.0 and sites of recombination were detected and removed via Gubbins 2.3.1. Polymorphic sites between the aligned genomes were extracted to create a SNP alignment file using the SNP-sites tool^25^ 2.4.1. The SNP alignment-based phylogeny was constructed using FastTree^26^ 2.1.1 which infers approximately-maximum-likelihood trees.

Pan-genome analysis of all single colony isolates was undertaken by Panaroo^27^ version 1.2.2 using default settings (Fig. 6).

### Heatmaps and Venn diagrams

Heatmaps were created using ggplot2 and RColorBrewer through R statistical software^28^ version 3.5.1. “HP” gene reference numbers from the reference strain 26695 (NC_000915.1) were used for nomenclature followed by the gene product information as provided by the genome annotation pipeline described above. Where there were no matches to “HP” numbers, the gene abbreviation (if applicable) was used followed by the gene product information.

Venn diagrams were created using the R statistical software 3.5.1 VennDiagram package 1.6.20^29^.

## Results

### Genome alignments revealed larger scale differences between antrum and corpus populations

Using deep sequencing, hotspots of genetic diversity were observed between *H. pylori* populations in the antrum and corpus regions by aligning the population consensus genomes (Fig. 2A-B; Suppl. Fig. 2-15). However, some hotspots were located near assembly contig breaks with fewer SNPs supported by both alignment and read mapping methods (red SNPs; Fig. 2A-B) suggesting these could be assembly-associated artefacts. Noticeable gaps or stretches with <95% BLASTN identity were identified between the antrum and corpus aligned consensus assemblies from 12/16 patients.

Single colony isolate genomes from the antrum and corpus of patients with paired samples were aligned and visualised using high BLASTN identity between 96% and 100% (Fig. 2C; Suppl. Fig. 6C, 8C, 10C, 12C, 13C, 14C and 15C) to identify small scales differences between the aligned genomes. Defined stretches of BLASTN identity were evident between single colony isolates from the antrum and corpus of each patient. This “visual fingerprint” could differentiate the isolates taken from the antrum and corpus, suggesting stomach region-specific adaptation and between-region differences.

More nonsynonymous than synonymous mutations were detected between paired antrum and corpus populations suggesting that selection pressures were acting independently within and between these regions (Suppl. Fig. 16).

### Deep sequencing and read mapping identified the most variable genes within *H. pylori* populations

Within-region diversity was firstly detected using very stringent criteria to detect “common” allelic variation (Fig. 3A; Suppl. Fig. 17). Stringency thresholds were then lowered for some parameters (particularly the number of alternative reads mapping in the forward and reverse direction) to detect “minor” allele diversity (Fig. 3B; Suppl. Fig. 18). “Minor” alleles were present at lower frequencies in the population, while “common” alleles were present at higher frequencies and represent polymorphisms that may be approaching fixation in the population or represent genes/bases under more recent selective pressures^13^.

**Fig. 3.**
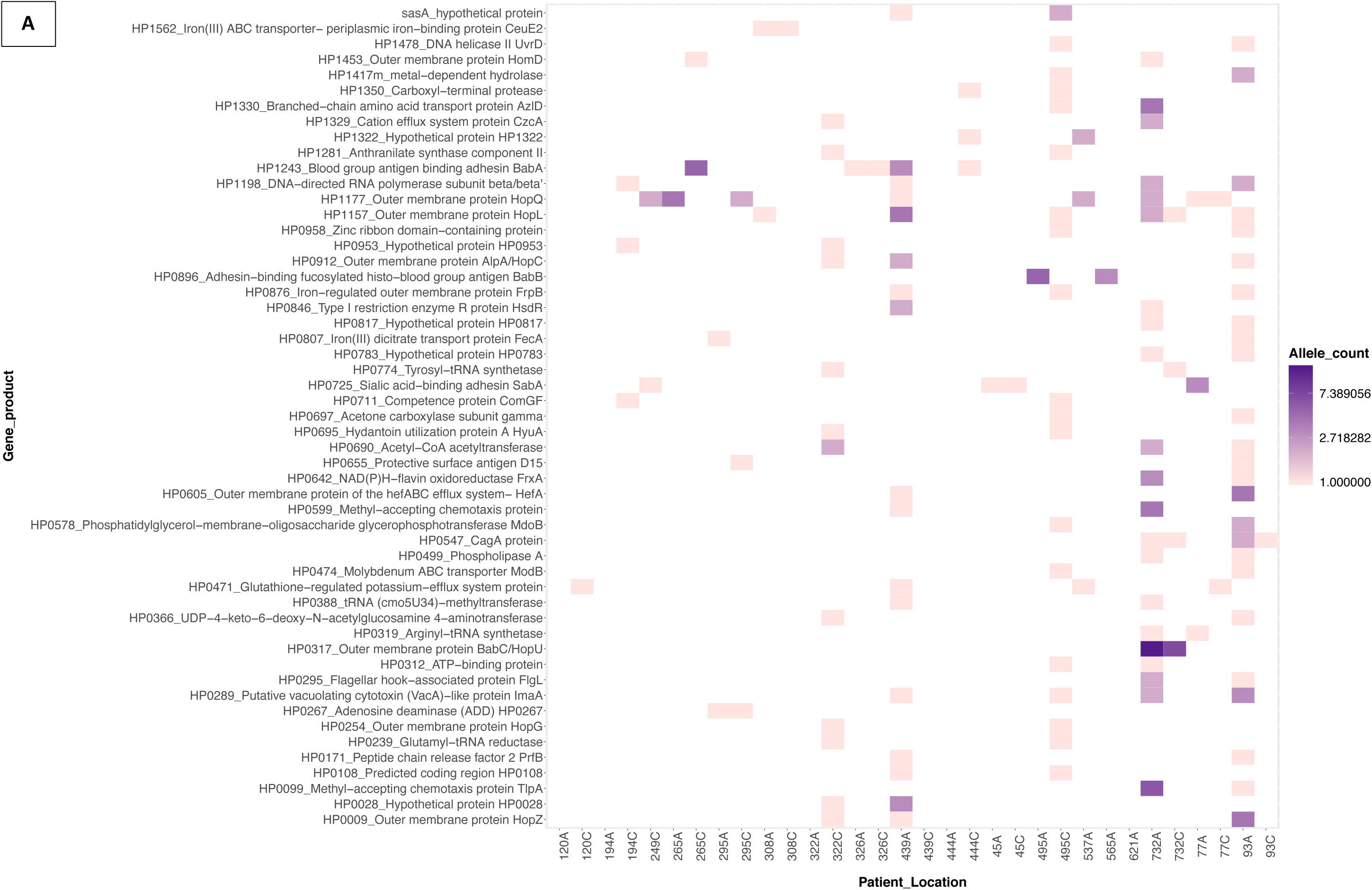
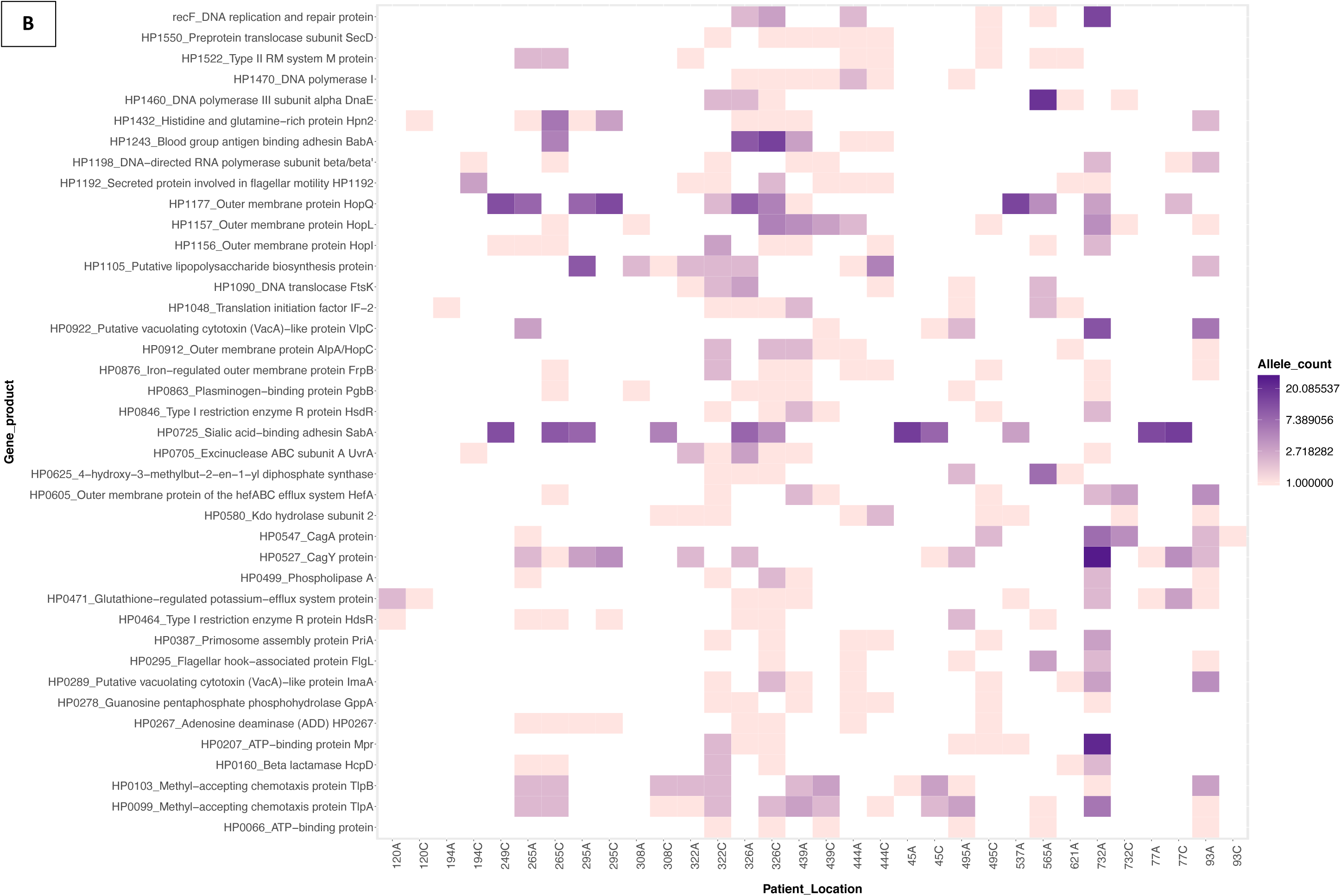
Heat maps showing common (A) and minor (B) allelic variation across genes/gene products for each antrum- and corpus-derived *H. pylori* population, derived from deep sequencing of the population from each biopsy and read-mapping back to the consensus genome to identify variant bases at each locus. Higher colour intensity indicates a larger number of variant bases within that gene/gene product. Only variable genes represented by two or more patient samples were included in panel A. Due to the high number of allelic genes in the minor allele variation, panel B was filtered to highlight allelic variant genes/gene products shared between six or more different patient samples. In both panels sample 565C was excluded due to the extreme variation observed. For an undocketed heatmap, including sample 565C, see Suppl. Fig. 17-18.

A total of 7,920 common allelic variants were detected within 1011 genes from 33 deep-sequenced samples. Excluding hypothetical proteins, the highest common allelic variation (Fig. 3A; Suppl. Fig. 17) was observed within outer membrane protein (OMP) associated genes (*babA/B/C, frpB, hopA/B/C/D/E/G/H/I/L/N/Q/U/Z, hofA/B/C/D/E/F/G/H, homA/C/D, hefA/D/G, horB/C/D/E/F/G/I/J/K, lptB/D* and *sabA*) with polymorphisms in these genes detected in *H. pylori* populations isolated from 22/33 of the patient biopsies. Polymorphisms in virulence-related genes including the *vacA* paralogue HP0289 (n=5), *babA* (n=5), *cagA* (n=4) and *sabA* (n=4) were also identified in the common allele dataset. Other notable genes with high allelic diversity within the common allele dataset were those encoding the glutathione-regulated potassium-efflux system protein HP0471 (n=5), DNA-directed RNA polymerase subunit beta/beta (HP1198; n=5) and acetyl-CoA acetyltransferase (HP0690; n=5).

There were 16,492 minor allelic variants within 1,738 genes from 33 deep-sequenced samples. As with the common allelic variant analysis, OMP genes (in addition to *hopF/H/K*, *hefB/C*, *horA/H/I/L* and *lptA*) (Fig 3B; Suppl. Fig. 18) had the highest number of variants in the minor allele dataset (n=27). Methyl-accepting chemotaxis genes (*tlpA*/*B*/*C* and HP0599; n=18), restriction modification genes (hsdM/R, hdsM/R, Hpy8I, MboIIR, HP0479, HP0592, HP1351, HP1367, HP1368, HP1371, HP1471, HP1472, HP1499, HP1517, HP1521, HP1522; n=19), *vacA* like genes (*imaA*, *vlpC*, HP0610; n=14), *cag*PAI genes (cagΔ/β/γ/A/C/D/E/F/H/M/N/S/W/X/Y/Z; n=18), salic acid-binding adhesin gene *sabA* (n=11), lipopolysaccharide biosynthesis genes (HP0208; HP0208; n=13) and the glutathione-regulated potassium-efflux system (HP0471; n=11) were also highly variable (Fig. 3B; Suppl. Fig. 18).

### Combining population deep sequencing, read mapping and genome alignment allows more powerful analysis of *H. pylori* population structure and diversity

Figure 4 brings together the key findings from the whole-genome alignments (Fig. 2A-B; Suppl. Fig. 2-15) and read mapping approach (Fig. 3B) and highlights the number of differences detected by each approach. Some genes, for example *cagY* (HP0527) in patient 265, were identified as having differences between the antrum and corpus regions by genome alignment but also had polymorphic diversity *within* one or both region(s) detected by read mapping. Other genes were different between the regions (by genome alignment) but showed no within niche diversity at either region e.g. *dnaE* (HP1460) in patient 120.

**Fig. 4.**
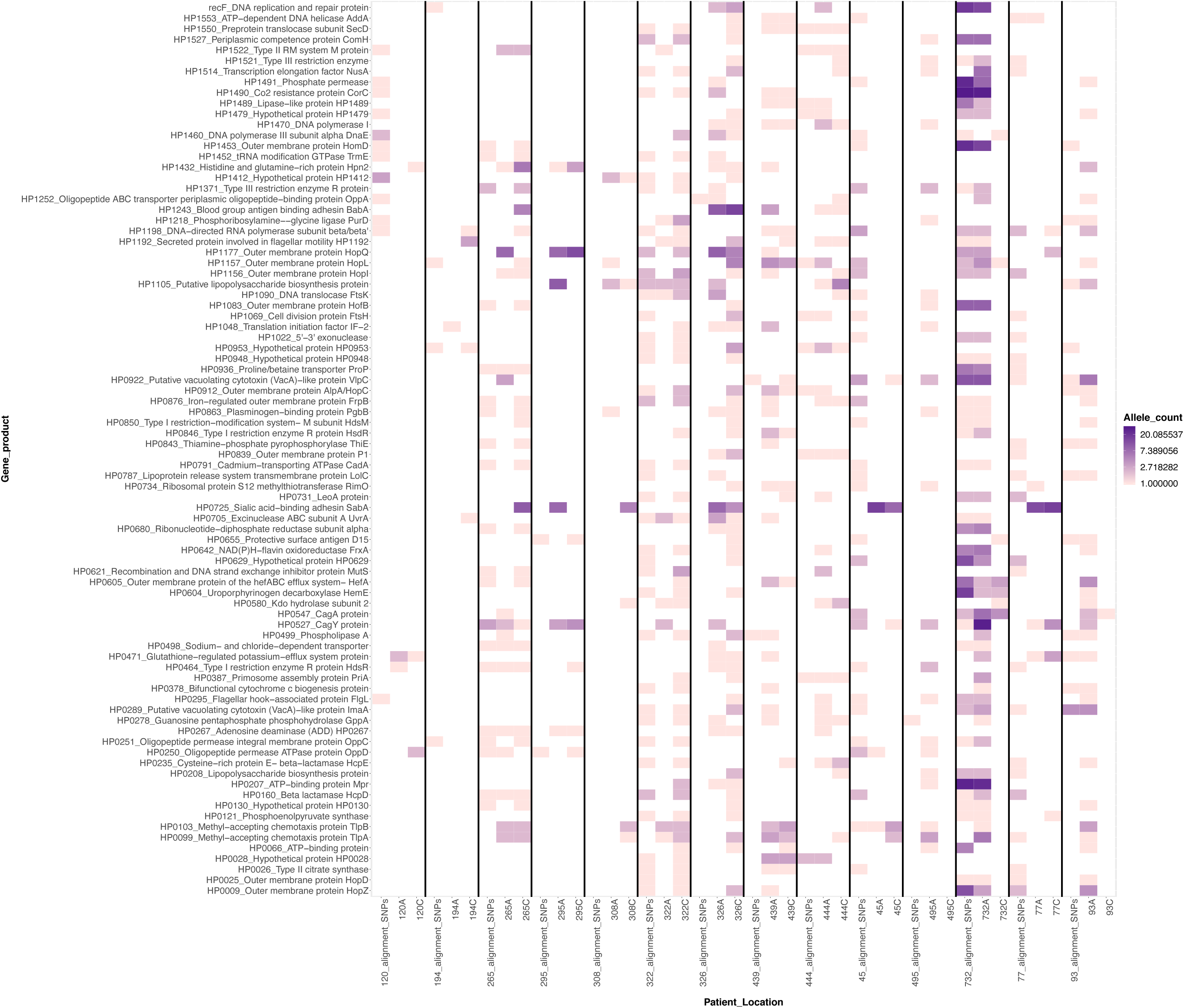
Heat map showing the most variable genes and populations identified by a combination of analytical approaches (within and between bacterial populations from the antrum and corpus regions of patient stomachs). Each column contains data from one patient. Within each column, three datasets are shown. From left to right these are: between stomach region variation from whole-genome alignment antrum versus corpus; within antrum minor allele variation; within corpus minor allele variation. Darker colour intensity indicates a larger number of variant bases within that gene/gene product. Gene products/genes that were shown to be diverse six or more times by any methodology across the patient datasets were included in this figure to reduce figure size and to highlight genes/gene products with higher observed genetic diversity. Hypothetical proteins and intergenic nucleotide diversity were removed from this dataset to improve visualisation. Only patients with paired antrum and corpus data were included. Patient 565 was excluded due to the extreme variation observed. A undocketed figure including patient 565 can be found in Suppl. Fig. 19. This Figure combines the information presented in Fig. 3B plus Suppl. Fig. 1 (detection of minor allelic variants within each stomach region by read-mapping of deep sequencing reads back to the consensus genome) and Fig. 2A-B (detection of validated ‘red’ SNPs by whole-genome alignment between stomach regions) plus Suppl. Fig. 2-15.

### Single colony sequencing added value by allowing phylogenetic and pangenome analyses

Phylogenetic trees were constructed for the single colony isolates from patient biopsies (Fig. 5). While the antrum and corpus isolates from some patients separated out into different clades, this was not always the case (e.g. Fig. 5B, 5F). There was evidence of migration of *H. pylori* between antrum and corpus in 5/9 patients with one or more isolate(s) clustering in the opposite region’s isolate cluster. Pangenome analysis (Fig. 6) confirmed that the isolates from each patient clustered together, indicating that most patients had originally been infected with a single strain that subsequently diversified. The exception was patient 565 in whom two distinct *H. pylori* clades were evident.

**Fig. 5.**
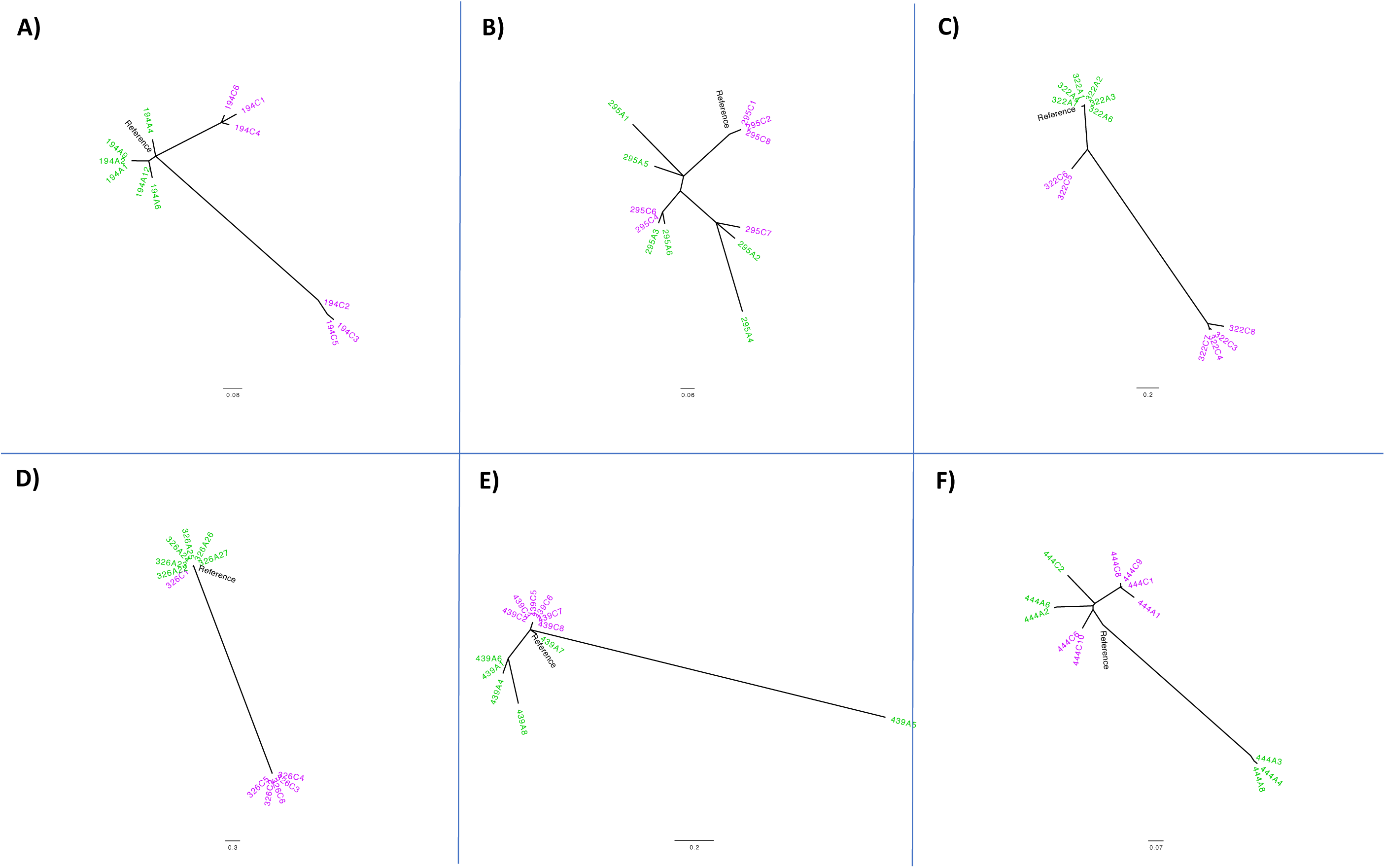
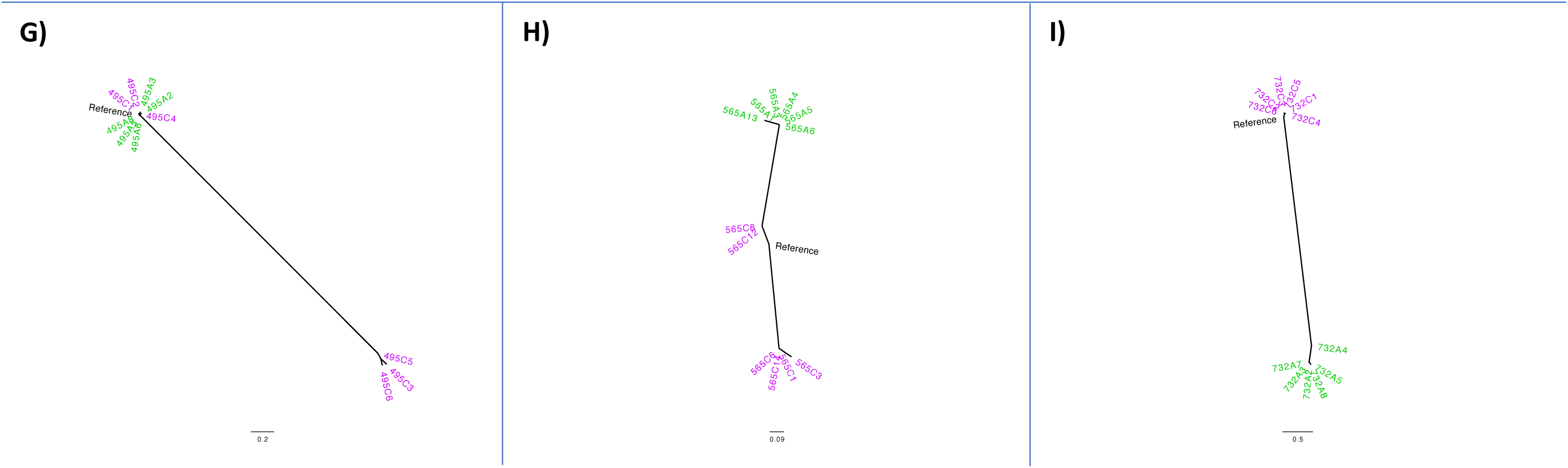
Phylogenetic trees depicting single colony isolates from patients: A) 194, B) 295, C) 322, D) 326, E) 439, F) 444, G) 495, H) 565 and I) 732. Antrum-derived strains are shown in green and corpus-derived strains in pink/purple. The patient reference consensus genome is shown in black. Phylogenies were rooted to the midpoint in decreasing order and drawn in a radial format.

### Population deep sequencing with read mapping is much more sensitive for detecting polymorphisms than conventional single colony sequencing, but combining both methods is best

Single colony isolates were used to validate the deep sequencing minor allele calling pipeline. Using a read mapping approach and exact base matching we determined an average of 68.69% (95% CL: 59.13% – 78.25%) alleles were detected only by the population deep sequencing, 8.13% (95% CL: 4.83% – 11.43%) by the single colony isolate analysis and 23.18% (95% CL: 14.93% – 31.43%) were concordant between methodologies (Suppl. Fig. 20).

## Discussion

By employing a combination of deep and single colony sequencing approaches in *H. pylori* for the first time, we detected extensive diversity within and between bacterial populations from the antrum and corpus regions of patient stomachs, including in virulence and colonisation associated genes. This combined approach generated a richer dataset than either individual approach.

Of the studies that have investigated genetic diversity of *H. pylori* within individual patients^30–38^, only three^30, 37, 38^ used a whole-genome sequencing approach. Others used PCR amplification and sequencing to focus on specific genes and loci. Didelot *et al*. (2013)^30^ sequenced single isolates from the antrum and corpus of 45 individuals and found evidence of micro-evolution of the bacterial population within each individual’s stomach.

In this study we used deep sequencing to detect much higher levels of genetic diversity in *H. pylori* populations than has previously been reported, even when very stringent parameters were applied (Fig. 3A). When the parameters were relaxed to detect minor allelic variants (Fig. 3B), a much larger number of polymorphic sites were detected. This study presents a comprehensive snapshot of *H. pylori* genetic diversity at a single point in time.

Some of the most frequently identified common allelic variant genes were related to virulence and colonisation, e.g. OMPs, *vacA* paralogue HP0289, *babA*, and *cagA*. In comparison, many of the minor allelic variants were in OMP genes which make up approximately 4% of the *H. pylori* coding genome^39^ and are highly diverse and polymorphic^39–41^. Most studies to date have studied polymorphic variation in *H. pylori* OMPs between patients, geographical regions, sequential isolates from animal models or familial isolated strains to reach these conclusions^30, 40, 42, 43^. The comparative genetics approaches of such studies have contributed to the identification and understanding of OMP polymorphic diversity, but polymorphic diversity has yet to be shown within populations taken from the same time point, despite the identification of OMP phase variation and gene conversion^42–44^. This is perhaps hampered by the difficulty and increased workload in isolating single colonies from population sweeps, the lack of paired biopsy samples from the same patient and the increased sequencing costs such investigations incur.

*cagA* gene diversity between individuals and geographical regions^45, 46^ is well characterised. To the best of our knowledge, *within*-patient genetic diversity of *cagA* at a single point in time has not yet been shown. In this study, we evidence both within- and between-stomach region genetic diversity of *cagA* (Fig. 3A-B; Fig. 4; Suppl. Fig. 17-19). Within- and between-region diversity in *cagA* sequences could result in variations in virulence activity within the stomach, thus influencing gastritis patterns and disease development. Within-region polymorphic diversity of *vacA* paralogues HP0289, HP0610 and HP0922 was also observed. These genes are thought to play roles in host colonisation and collagen degradation. Diversity here could therefore impact on colonisation, persistence, and disease outcomes.

Several important genes and groups of genes were identified within both the common and minor allele pipelines (Fig. 3A-B). These included the DNA-directed RNA polymerase beta subunit (HP1198; *rpoBC*). Certain mutations within the *rpoB* gene of *H. pylori* have been shown to increase resistance to rifamycins^47, 48^. We also observed diversity in the clarithromycin resistance-associated gene *kefB* (HP0471)^49, 50^. These observations might help to explain eradication therapy failure within patients whereby minority resistant strains persist within the population. Methyl-accepting chemotaxis associated genes (HP0082; HP0103; HP0099; HP0599) were also identified (including consensus genome alignment-based SNPs in patient 45). These genes are important for *H. pylori* colonisation and survival, as *tlpA* (HP0099) senses arginine, bicarbonate, and acidic pH^51, 52^, and *tlpB* (HP0103) is essential for chemotaxis away from acidic pH and towards more favourable conditions^53^. Such allelic diversity could affect the sensitivity of chemotactic responses and enable colonisation of different areas of the stomach by these strains.

In addition, restriction-modification system genes were identified. Furuta *et al.* (2015)^54^, observed similar genetic diversity among restriction-modification associated genes using a comparative genetics approach with single colony isolates obtained from five families. Other studies have also observed restriction-modification system diversity between strains taken from different patients^10, 55, 56^. Bacterial restriction-modification systems confer protection against invading foreign DNA^57^ but do not pose a barrier to homologous recombination^58^. Restriction-modification systems can also influence gene expression^59, 60^ and may have roles in adhesion and virulence^61, 62^. Diversity in restriction-modification genes could therefore play a role in niche adaptation and persistence.

Lipopolysaccharide biosynthesis associated genes were also highly polymorphic in the common and minor allele pipelines. 18 samples had within-region minor allelic diversity amongst genes of the putative outer membrane biogenesis complex components^63^. While we have focused on differences and diversity, our dataset could equally be used to identify useful regions of conservation that might be exploited in vaccine development.

Our combined approach revealed that some genes identified with *between*-region diversity by genome alignment also had polymorphic diversity *within* one or both regions of the stomach. For example, the iron regulated OMP gene of sample 265C was polymorphic within this population but was also identified by whole-genome alignment as variable between the consensus genome sequences of the antrum and corpus derived samples from patient 265. This observation was not uncommon across the dataset.

The phylogenetic relationships between single colony isolates obtained from the antrum and corpus in this study agree with Ailloud *et al.* (2019)^38^. There were distinct clades of *H. pylori* between the antrum and corpus in some patients, and there was evidence of migration between the two sites in others. Ailloud *et al.* (2019)^38^ showed that migration of *H. pylori* strains from the antrum to the corpus was relatively infrequent whereas migration between the corpus and fundus is a more common event, perhaps due to the more significant environmental differences between the antrum and corpus regions. Our phylogenetic analysis also indicated that where an antrum cluster was present, corpus strains were more frequently observed within antrum clades than vice versa. This suggests that while migrations between the antrum and corpus are infrequent, the corpus isolates are more likely to migrate to the antrum than vice versa. Again, this could be due to the differences between the antrum and corpus environments where antrum isolates are less fit or poorly adapted to colonise the harsher oxyntic epithelium whereas the corpus strains are able to colonise the more neutral antrum glands.

An alternative explanation for the non-clustering of antrum and corpus strains in patients 295 and 444 could be due to the biopsy sampling location and method. Fung *et al.* (2019)^64^, showed how founder strains initially colonise glands which spread to adjacent glands in the immediate vicinity. This creates islands of closely related *H. pylori* strains, and where island boundaries occur the inhabitants may then compete for space. At these boundaries there are glands containing a mixture of strains, or adjacent glands containing different strains side by side. Transition zones between the antrum and corpus typically contain mixed populations. The size of a biopsy (how many strain islands it spans) and its proximity to the antrum-corpus transition zone may influence the *H. pylori* diversity observed. This could potentially disrupt the genetic clustering and phylogenetic topology of the antrum and corpus. Therefore, standardised sampling locations within the antrum and corpus would be beneficial for this type of analysis. However, this may not be feasible in practice because biopsies are often taken from areas likely to harbour *H. pylori* infection, such as adjacent to visually diseased epithelium.

The pan genome analyses of the single colony isolates and the consensus assembled deep-sequenced populations, both showed that the strains from each patient had a unique pattern of gene presence and absences, distinct from the patterns observed in strains from other patients. The exceptionally diverse *H. pylori* samples from patient 565 all clustered together, but there were substantial gene content differences between single colony isolates from this patient. The corpus isolates from this patient were much more diverse than the antrum isolates, not just allelically but also in their accessory genes. Consequently, patient 565 may have been infected by more than one *H. pylori* strain that exchanged DNA through homologous recombination and/or natural transformation over time and now share a similar genetic signature (Fig 6).

By combining conventional sequencing of single colony isolates from antrum and corpus with population deep sequencing of the same samples, from multiple patients, we were able to comprehensively characterise *H. pylori* population diversity. The dual approach allowed for comparative analysis to determine how well the data generated from each approach agreed with each other. This confirmed that the deep sequencing allelic calling pipeline was able to detect 91.87% of the SNPs, with only 8.13% of SNPs identified from the single colony pipeline alone. The population deep sequencing minor allele calling methodology captured a more comprehensive snapshot of population genetic diversity compared to the single colony approach. However, the minor allele calling pipeline is still likely to be an underestimate of the true diversity present, due to the stringent quality control parameters that were applied, and single colony isolate sequencing from the same populations added value because it enabled additional phylogenetic and recombination analyses that would not have been possible using the deep sequencing data alone.

Combining the two complementary approaches of deep population sequencing and single colony isolate sequencing generated more information than either individual strategy. But this combined approach was intensive and was only applied to a relatively small number of UK-based patients in this study. This study did not have sufficient sample size to identify any significant associations between genomic traits in the *H. pylori* populations and the presence of ulcers, intestinal metaplasia, or severe inflammation in the patients’ stomachs. This study also relied on a culture-based approach, so may not provide a full picture of the genetic diversity of strains present in the stomach due to the selection pressures applied by culture on agar plates prior to sequencing. Although genetic changes over time can be inferred from the data, this study was not longitudinal, and samples were taken only once from each patient.

In conclusion, we have shown that single colony analysis alone can identify fixed differences in the genomes of *H. pylori* between stomach regions, but population deep sequencing reveals that underlying variation is still present and the population as a whole retains a high degree of potential plasticity. This may help explain why *H. pylori* can persist in a chronic infection. Identifying loci with minimal variation might usefully inform future vaccine design, while loci in which high numbers of minor allelic variants are concentrated indicate which genes are critical for niche adaptation and persistence.

## Acknowledgements

We thank Dr Kaisa Thorell for providing genome annotation expertise, technical and research staff at Nottingham Trent University and at the University of Nottingham for helping to drive this research forward. Mrs Joanne Rhead and Ms Melanie Lingaya isolated *H. pylori* from human gastric biopsies. We would also like to thank Professor John Atherton and Professor Alan McNally for advising on this project. This study was supported by the Vice-Chancellor’s Scholarship and the School of Science and Technology at Nottingham Trent University as well as the National Institute for Health Research (NIHR), through the Biomedical Research Centre at Nottingham University Hospitals NHS Trust and the University of Nottingham. The views expressed are those of the authors and not necessarily those of the NHS, the NIHR or the Department of Health.

Author names in bold designate shared co-first authorship

