## Supplemental Figures for "Genomic diversity of *Helicobacter pylori* populations from different regions of the human stomach"

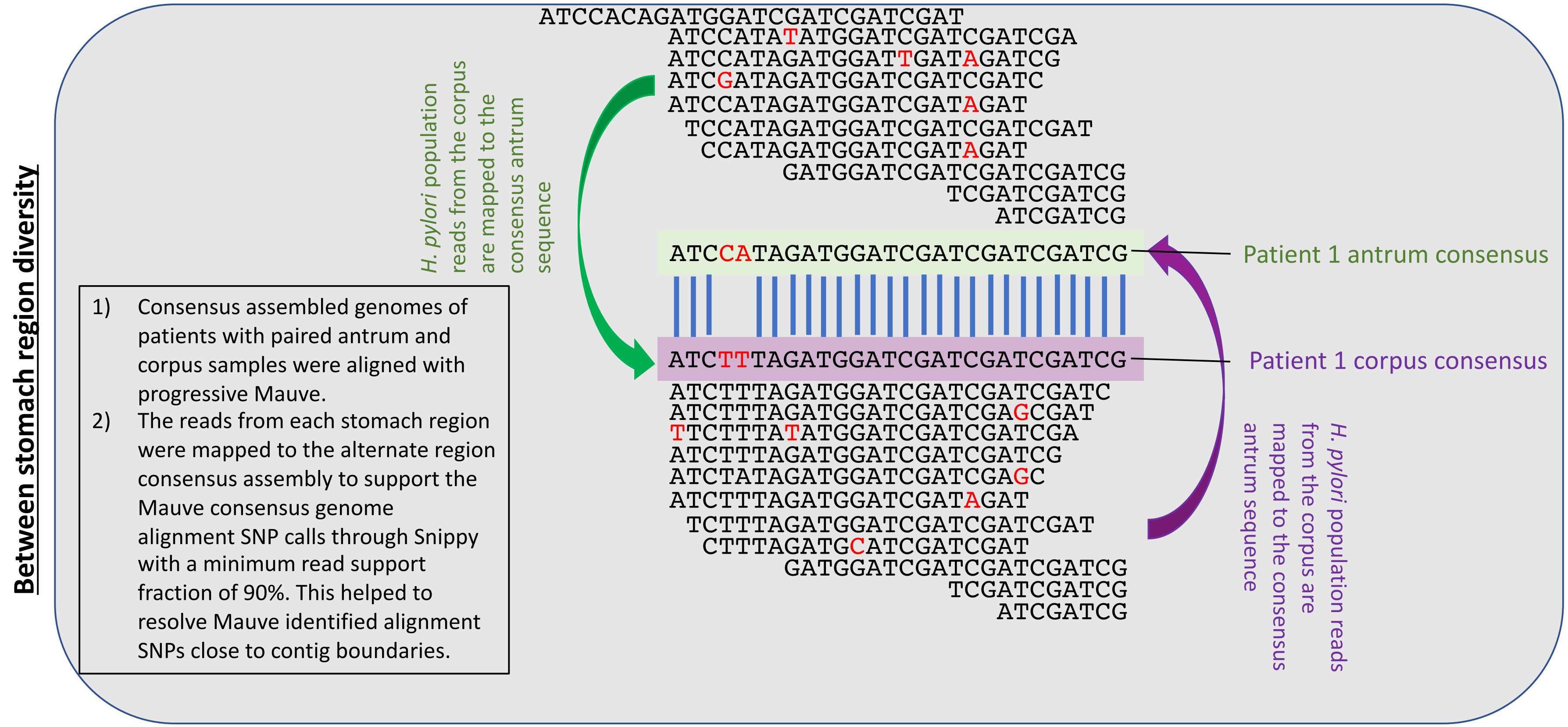


**Suppl. Fig. 1**. Between region diversity investigated using a consensus and read mapping alignment approach. This approach was taken to identify between region SNPs of high quality by supporting alignment SNPs with alternative stomach region read mapping to the opposite location consensus genome.

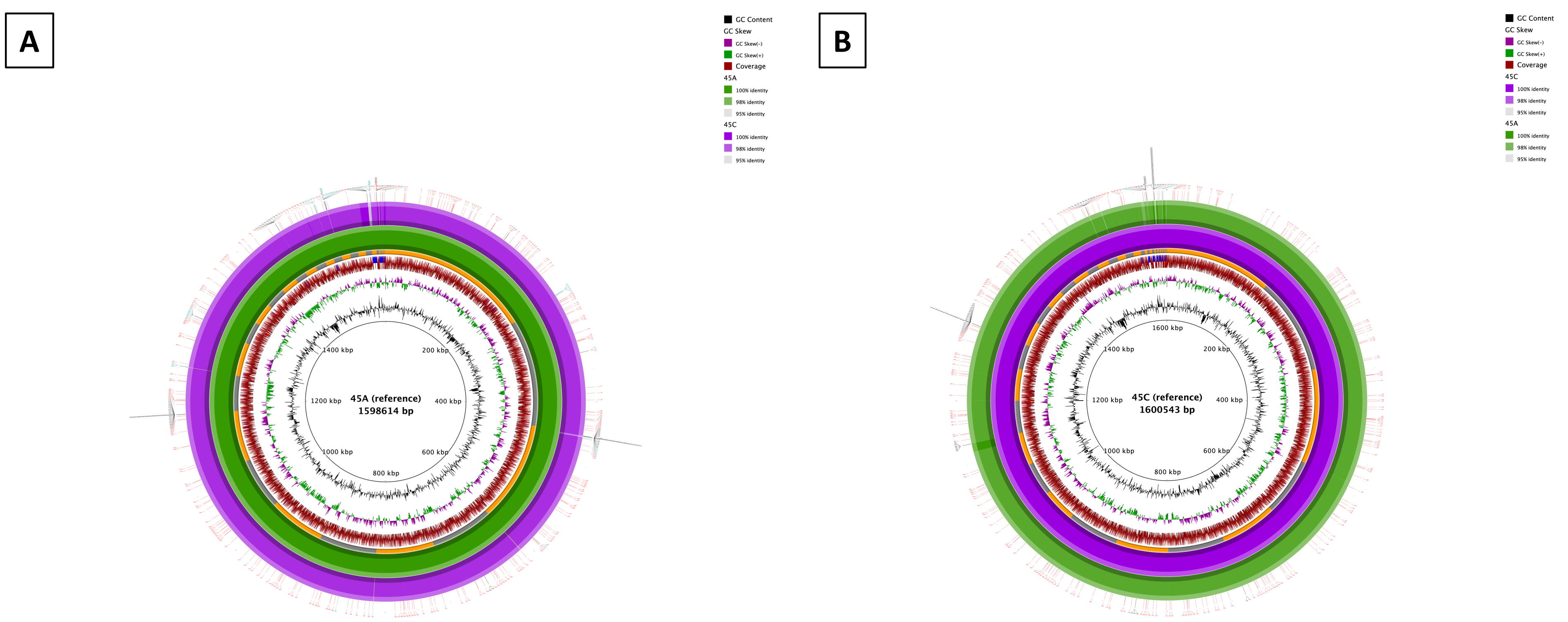


**Suppl. Fig. 2**. Representative whole genome alignment for *H. pylori* strains isolated from patient 45. Whole genome alignments of the consensus genomes generated by deep population sequencing of antrum and corpus *H. pylori* populations, using antrum (A) or corpus (B) consensus genome as the reference. Colour intensity of the outermost ring indicates percentage identity between antrum and corpus consensus genomes. Positions of contigs within the assembled reference genome are shown as a ring alternating in colour between grey and orange. Peripheral ticks indicate positions of SNPs between antrum and corpus genomes identified by whole genome alignment (black), read mapping (teal), or both methods (red) in figures A-B. Plots generated using BRIG.

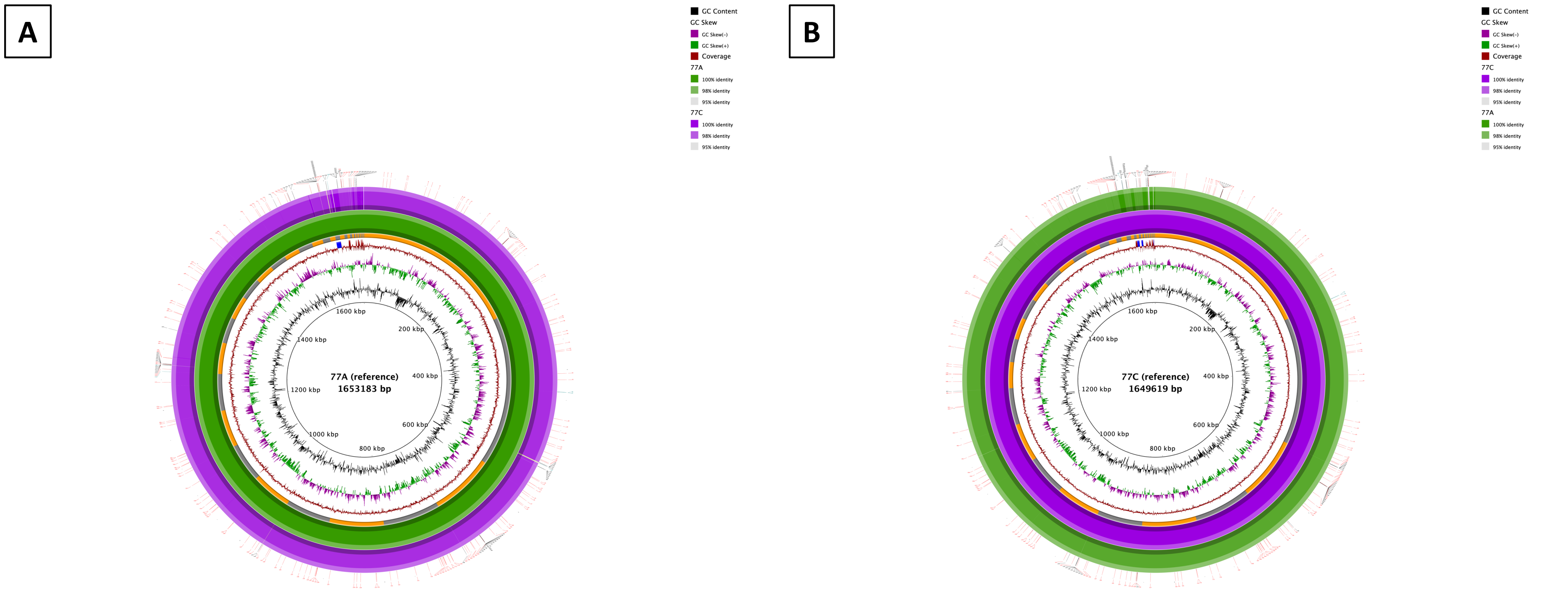


**Suppl. Fig. 3**. Representative whole genome alignment for *H. pylori* strains isolated from patient 77. Whole genome alignments of the consensus genomes generated by deep population sequencing of antrum and corpus *H. pylori* populations, using antrum (A) or corpus (B) consensus genome as the reference. Colour intensity of the outermost ring indicates percentage identity between antrum and corpus consensus genomes. Positions of contigs within the assembled reference genome are shown as a ring alternating in colour between grey and orange. Peripheral ticks indicate positions of SNPs between antrum and corpus genomes identified by whole genome alignment (black), read mapping (teal), or both methods (red) in figures A-B. Plots generated using BRIG.

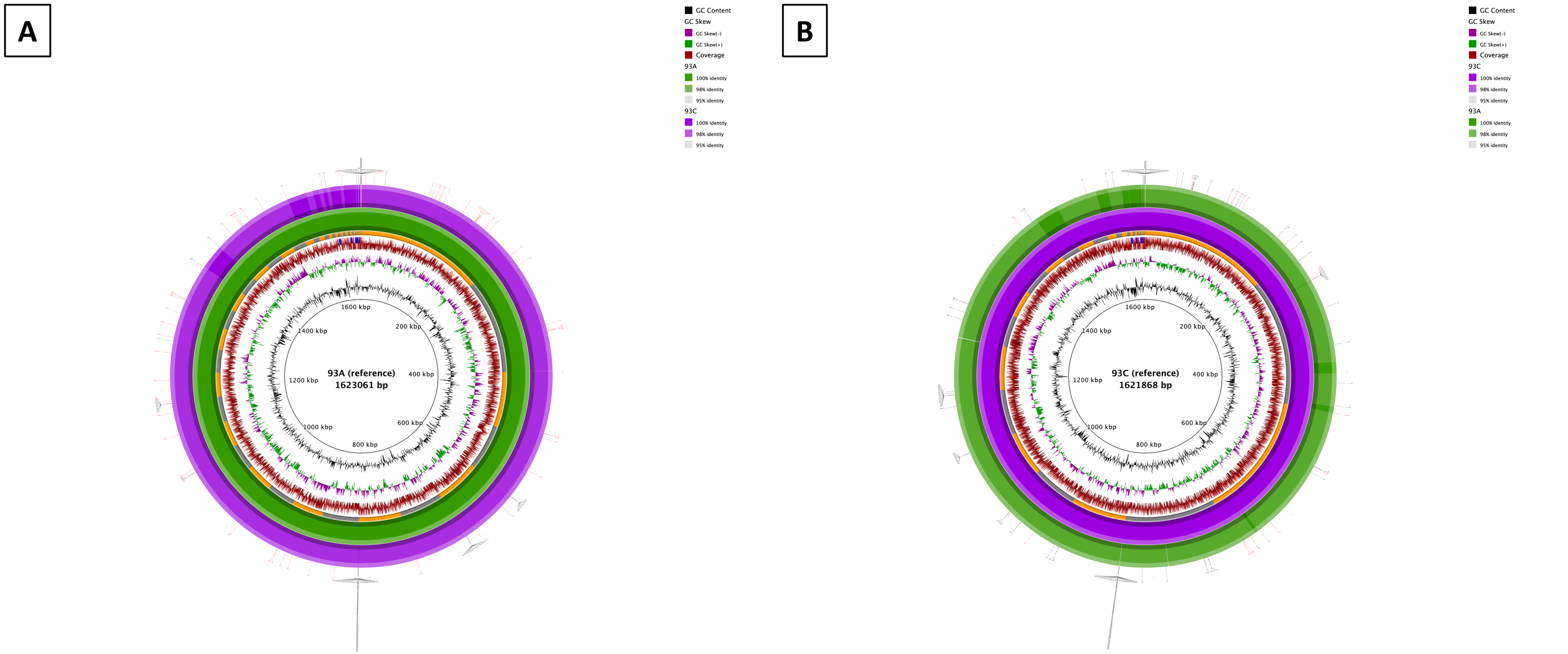


**Suppl. Fig. 4**. Representative whole genome alignment for *H. pylori* strains isolated from patient 93. Whole genome alignments of the consensus genomes generated by deep population sequencing of antrum and corpus *H. pylori* populations, using antrum (A) or corpus (B) consensus genome as the reference. Colour intensity of the outermost ring indicates percentage identity between antrum and corpus consensus genomes. Positions of contigs within the assembled reference genome are shown as a ring alternating in colour between grey and orange. Peripheral ticks indicate positions of SNPs between antrum and corpus genomes identified by whole genome alignment (black), read mapping (teal), or both methods (red) in figures A-B. Plots generated using BRIG.

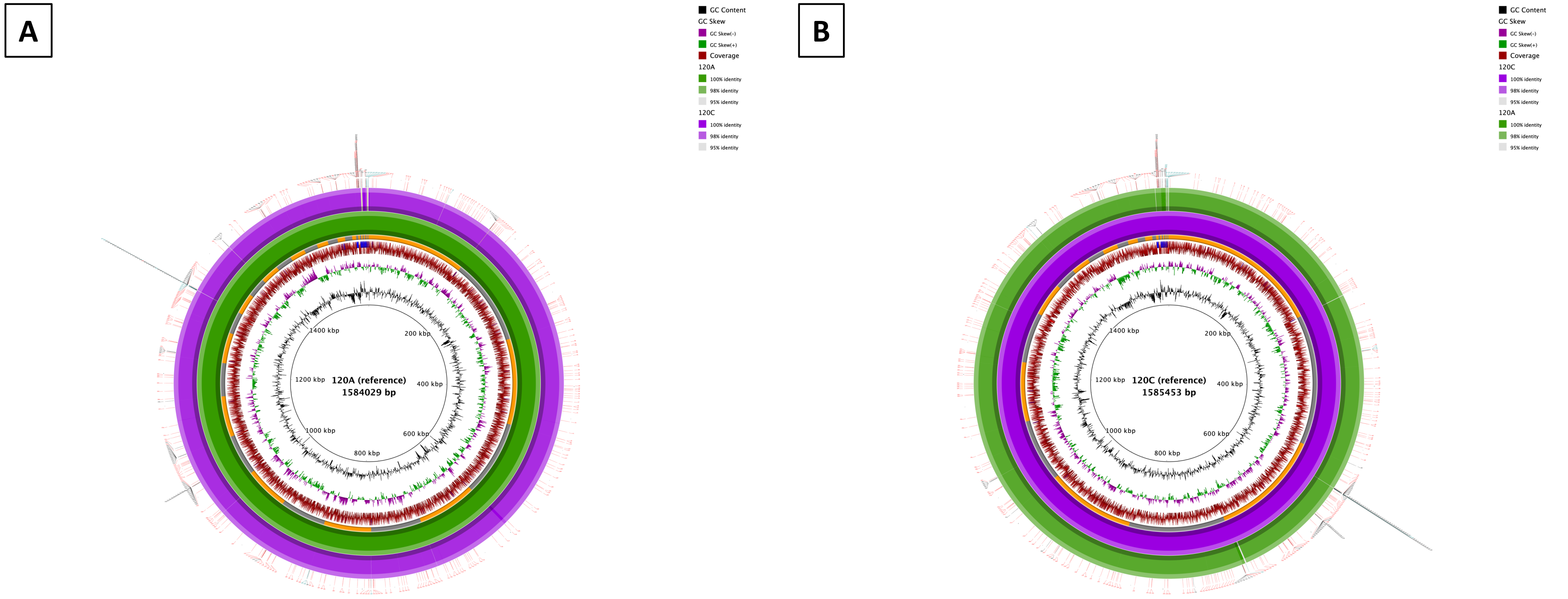


**Suppl. Fig. 5**. Representative whole genome alignment for *H. pylori* strains isolated from patient 120. Whole genome alignments of the consensus genomes generated by deep population sequencing of antrum and corpus *H. pylori* populations, using antrum (A) or corpus (B) consensus genome as the reference. Colour intensity of the outermost ring indicates percentage identity between antrum and corpus consensus genomes. Positions of contigs within the assembled reference genome are shown as a ring alternating in colour between grey and orange. Peripheral ticks indicate positions of SNPs between antrum and corpus genomes identified by whole genome alignment (black), read mapping (teal), or both methods (red) in figures A-B. Plots generated using BRIG.

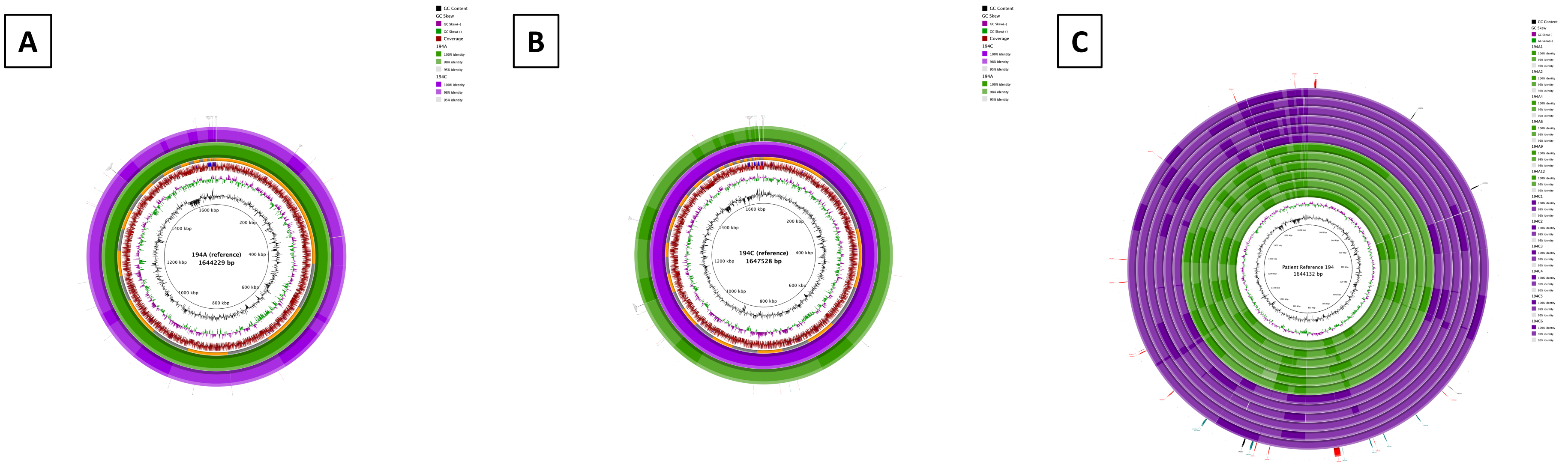


**Suppl. Fig. 6**. Representative whole genome alignment for *H. pylori* strains isolated from patient 194. Whole genome alignments of the consensus genomes generated by deep population sequencing of antrum and corpus *H. pylori* populations, using antrum (A) or corpus (B) consensus genome as the reference. Panel C depicts the alignment of colony isolate assembled genomes against the patient reference. Colour intensity of the outermost ring indicates percentage identity between antrum and corpus consensus genomes. Positions of contigs within the assembled reference genome are shown as a ring alternating in colour between grey and orange. Peripheral ticks indicate positions of SNPs between antrum and corpus genomes identified by whole genome alignment (black), read mapping (teal), or both methods (red) in figures A-B. Plots generated using BRIG.

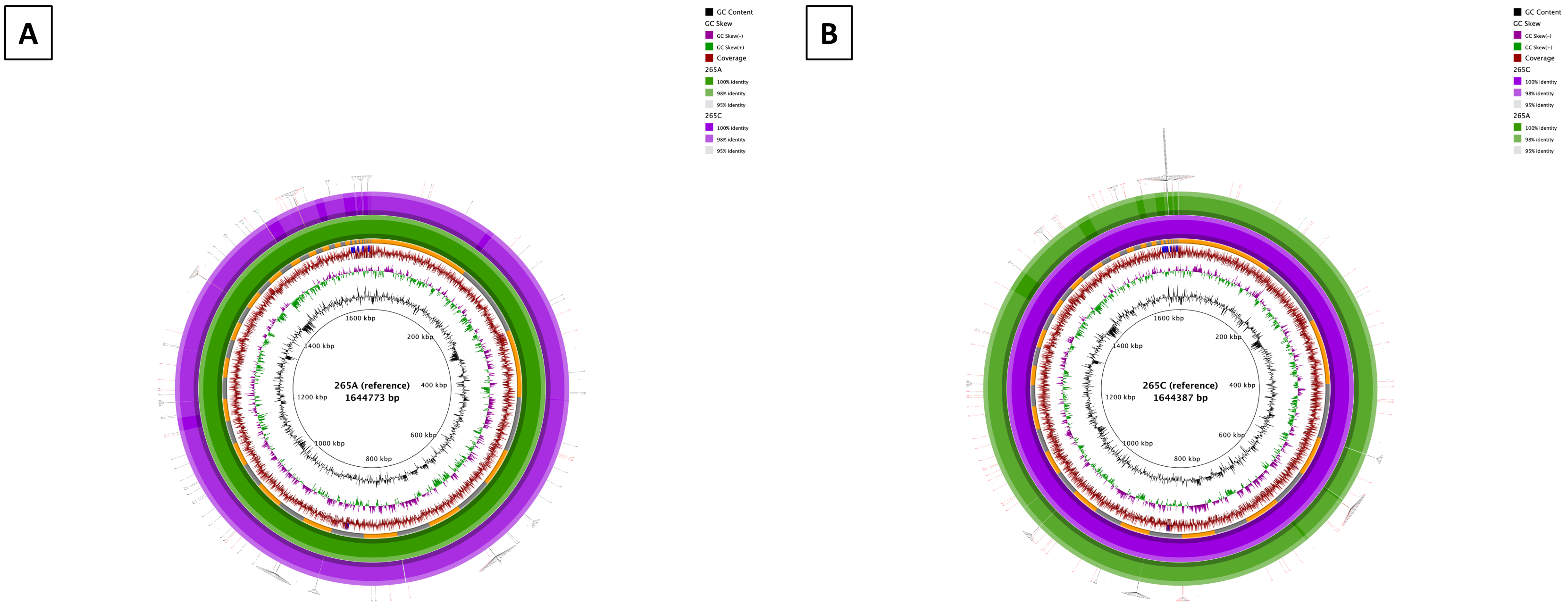


**Suppl. Fig. 7.** Representative whole genome alignment for *H. pylori* strains isolated from patient 265. Whole genome alignments of the consensus genomes generated by deep population sequencing of antrum and corpus *H. pylori* populations, using antrum (A) or corpus (B) consensus genome as the reference. Colour intensity of the outermost ring indicates percentage identity between antrum and corpus consensus genomes. Positions of contigs within the assembled reference genome are shown as a ring alternating in colour between grey and orange. Peripheral ticks indicate positions of SNPs between antrum and corpus genomes identified by whole genome alignment (black), read mapping (teal), or both methods (red) in figures A-B. Plots generated using BRIG.

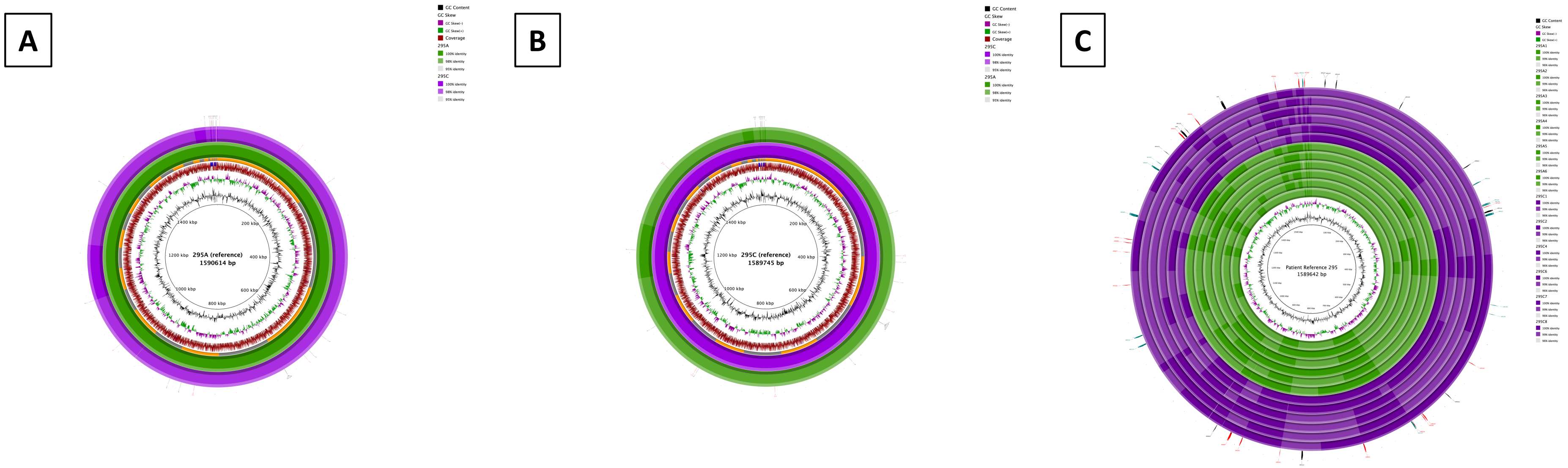


**Suppl. Fig. 8**. Representative whole genome alignment for *H. pylori* strains isolated from patient 295. Whole genome alignments of the consensus genomes generated by deep population sequencing of antrum and corpus *H. pylori* populations, using antrum (A) or corpus (B) consensus genome as the reference. Panel C depicts the alignment of colony isolate assembled genomes against the patient reference. Colour intensity of the outermost ring indicates percentage identity between antrum and corpus consensus genomes. Positions of contigs within the assembled reference genome are shown as a ring alternating in colour between grey and orange. Peripheral ticks indicate positions of SNPs between antrum and corpus genomes identified by whole genome alignment (black), read mapping (teal), or both methods (red) in figures A-B. Plots generated using BRIG.

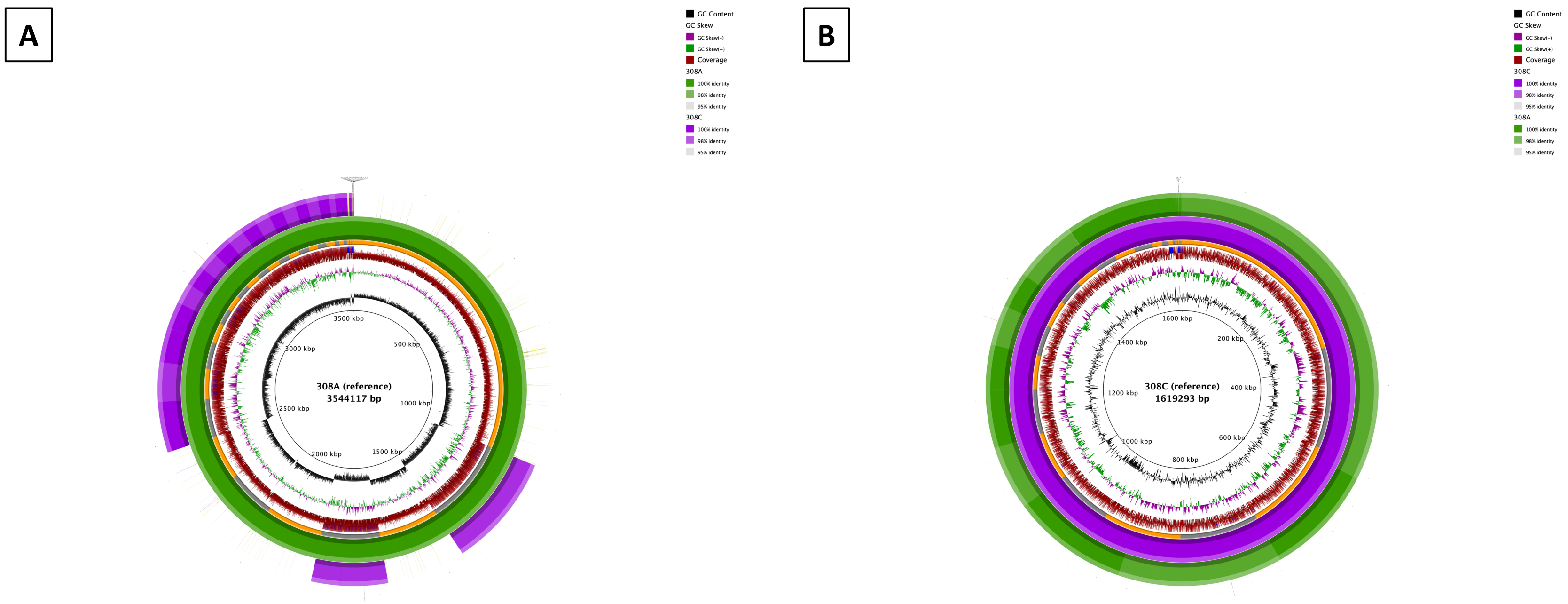


**Suppl. Fig. 9.** Representative whole genome alignment for *H. pylori* strains isolated from patient 308. Whole genome alignments of the consensus genomes generated by deep population sequencing of antrum and corpus *H. pylori* populations, using antrum (A) or corpus (B) consensus genome as the reference. Colour intensity of the outermost ring indicates percentage identity between antrum and corpus consensus genomes. Positions of contigs within the assembled reference genome are shown as a ring alternating in colour between grey and orange. Peripheral ticks indicate positions of SNPs between antrum and corpus genomes identified by whole genome alignment (black), read mapping (teal), or both methods (red) in figures A-B. Plots generated using BRIG.

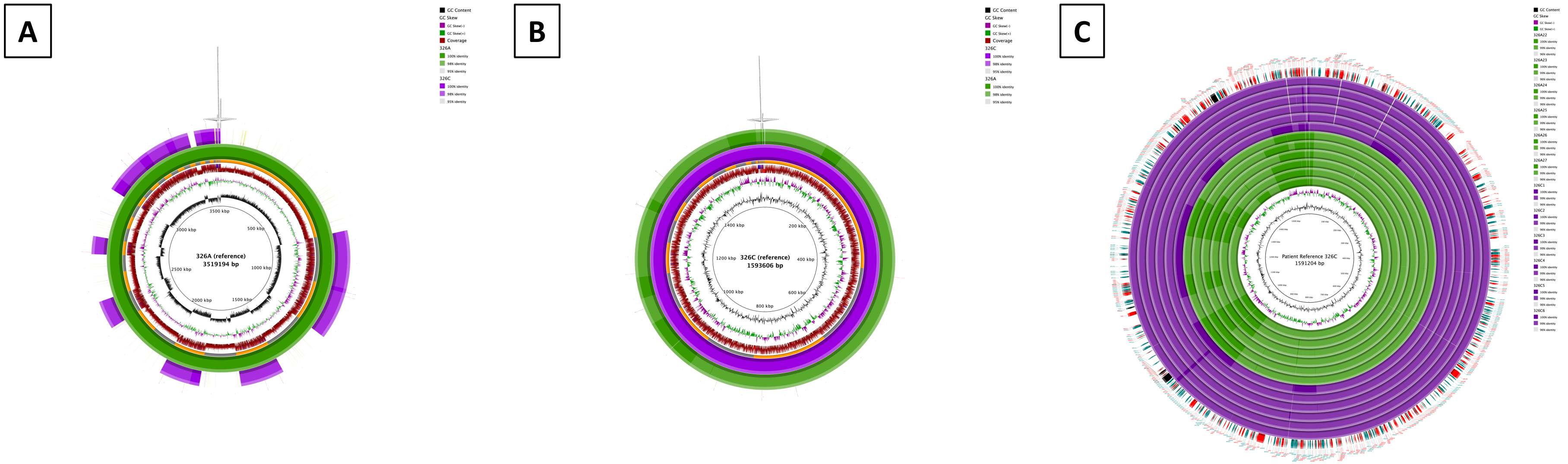


**Suppl. Fig. 10**. Representative whole genome alignment for *H. pylori* strains isolated from patient 326. Whole genome alignments of the consensus genomes generated by deep population sequencing of antrum and corpus *H. pylori* populations, using antrum (A) or corpus (B) consensus genome as the reference. Panel C depicts the alignment of colony isolate assembled genomes against the patient reference. Colour intensity of the outermost ring indicates percentage identity between antrum and corpus consensus genomes. Positions of contigs within the assembled reference genome are shown as a ring alternating in colour between grey and orange. Peripheral ticks indicate positions of SNPs between antrum and corpus genomes identified by whole genome alignment (black), read mapping (teal), or both methods (red) in figures A-B. Plots generated using BRIG.

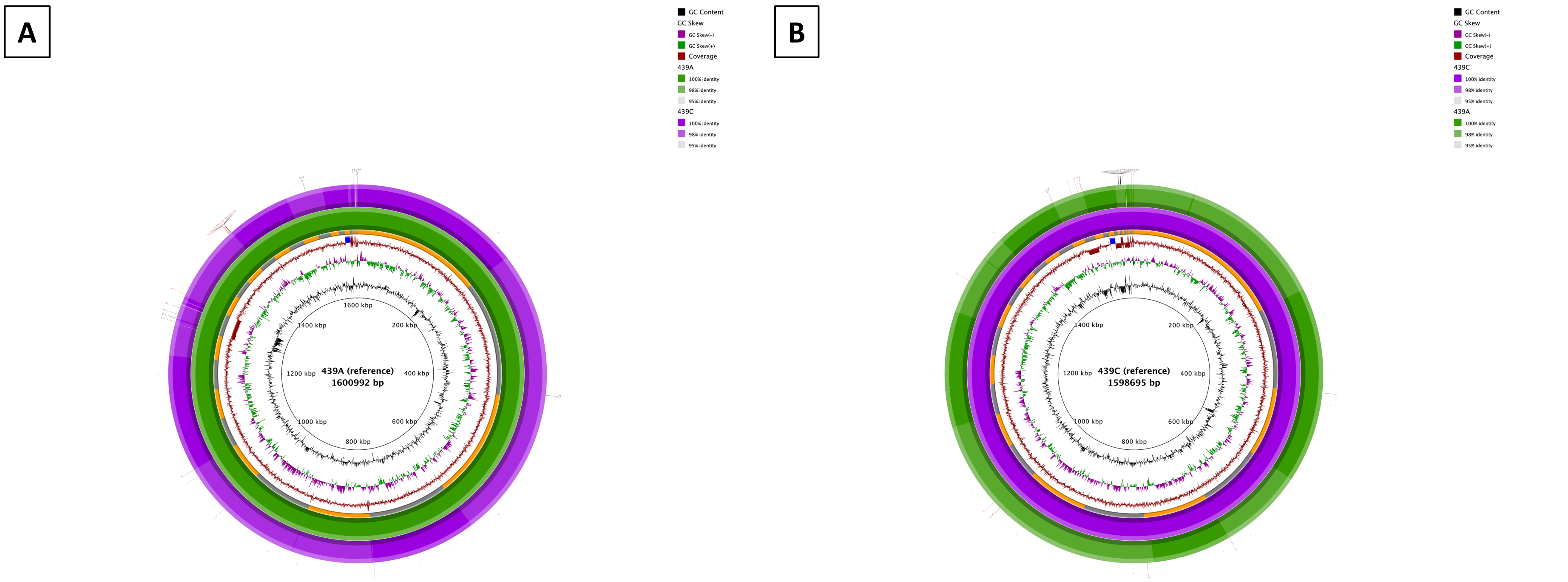


**Suppl. Fig. 11.** Representative whole genome alignment for *H. pylori* strains isolated from patient 439. Whole genome alignments of the consensus genomes generated by deep population sequencing of antrum and corpus *H. pylori* populations, using antrum (A) or corpus (B) consensus genome as the reference. Colour intensity of the outermost ring indicates percentage identity between antrum and corpus consensus genomes. Positions of contigs within the assembled reference genome are shown as a ring alternating in colour between grey and orange. Peripheral ticks indicate positions of SNPs between antrum and corpus genomes identified by whole genome alignment (black), read mapping (teal), or both methods (red) in figures A-B. Plots generated using BRIG.

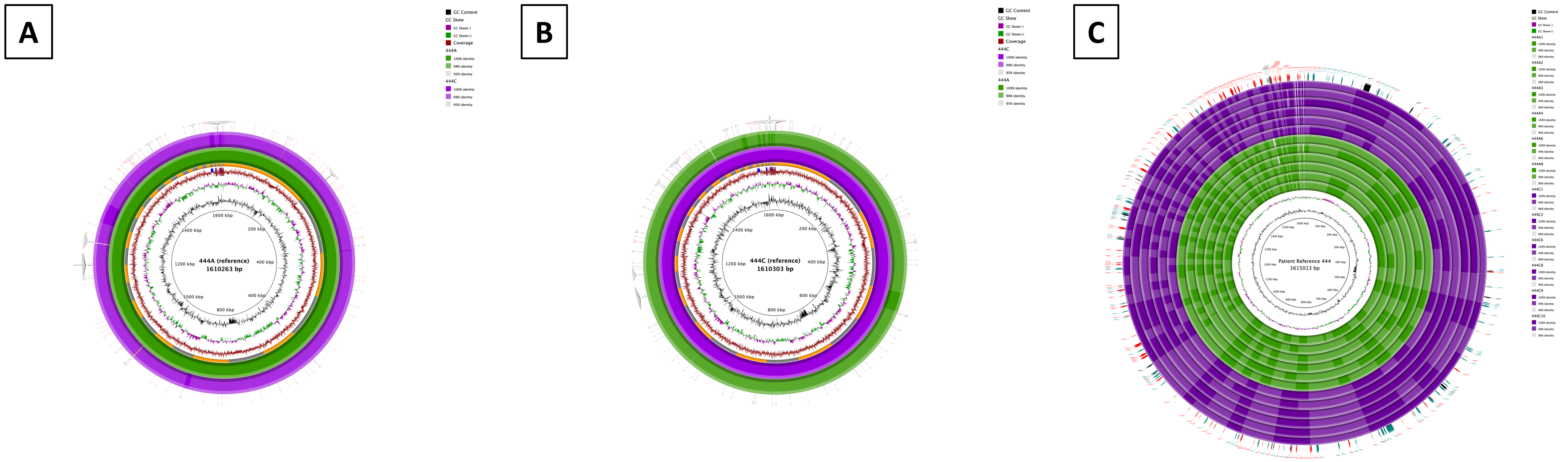


**Suppl. Fig. 12**. Representative whole genome alignment for *H. pylori* strains isolated from patient 444. Whole genome alignments of the consensus genomes generated by deep population sequencing of antrum and corpus *H. pylori* populations, using antrum (A) or corpus (B) consensus genome as the reference. Panel C depicts the alignment of colony isolate assembled genomes against the patient reference. Colour intensity of the outermost ring indicates percentage identity between antrum and corpus consensus genomes. Positions of contigs within the assembled reference genome are shown as a ring alternating in colour between grey and orange. Peripheral ticks indicate positions of SNPs between antrum and corpus genomes identified by whole genome alignment (black), read mapping (teal), or both methods (red) in figures A-B. Plots generated using BRIG.

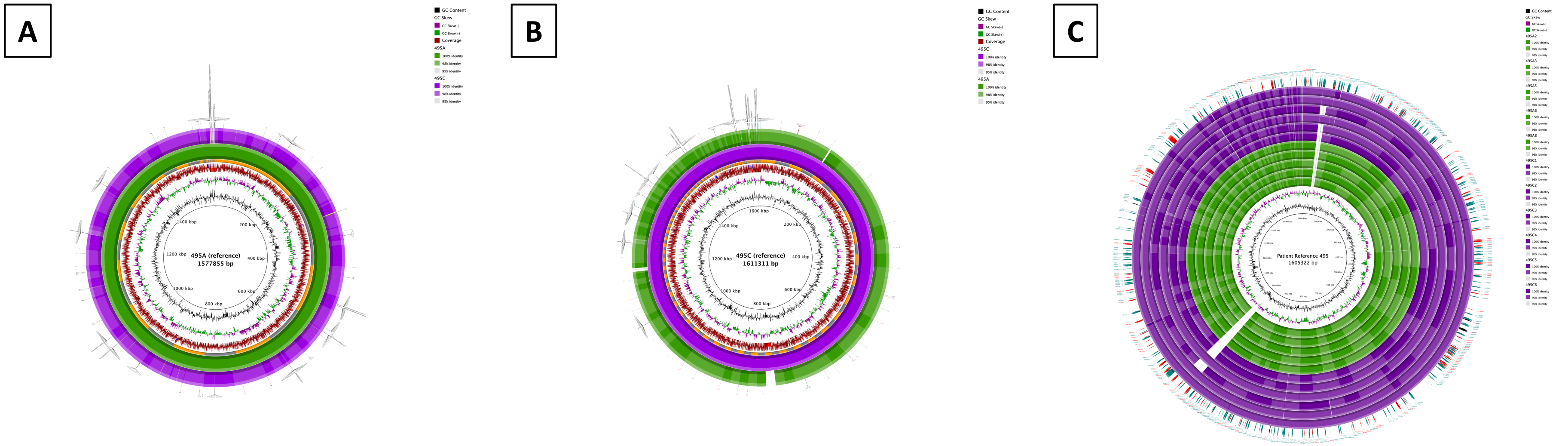


**Suppl. Fig. 13**. Representative whole genome alignment for *H. pylori* strains isolated from patient 495. Whole genome alignments of the consensus genomes generated by deep population sequencing of antrum and corpus *H. pylori* populations, using antrum (A) or corpus (B) consensus genome as the reference. Panel C depicts the alignment of colony isolate assembled genomes against the patient reference. Colour intensity of the outermost ring indicates percentage identity between antrum and corpus consensus genomes. Positions of contigs within the assembled reference genome are shown as a ring alternating in colour between grey and orange. Peripheral ticks indicate positions of SNPs between antrum and corpus genomes identified by whole genome alignment (black), read mapping (teal), or both methods (red) in figures A-B. Plots generated using BRIG.

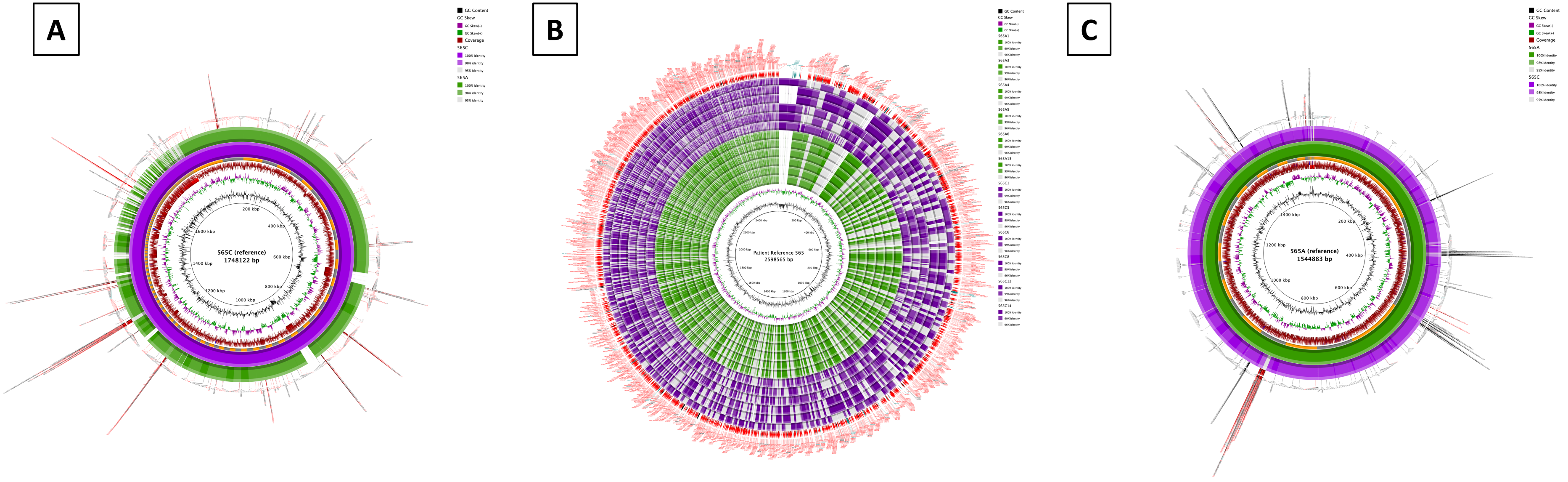


**Suppl. Fig. 14**. Representative whole genome alignment for *H. pylori* strains isolated from patient 565. Whole genome alignments of the consensus genomes generated by deep population sequencing of antrum and corpus *H. pylori* populations, using antrum (A) or corpus (B) consensus genome as the reference. Panel C depicts the alignment of colony isolate assembled genomes against the patient reference. Colour intensity of the outermost ring indicates percentage identity between antrum and corpus consensus genomes. Positions of contigs within the assembled reference genome are shown as a ring alternating in colour between grey and orange. Peripheral ticks indicate positions of SNPs between antrum and corpus genomes identified by whole genome alignment (black), read mapping (teal), or both methods (red) in figures A-B. Plots generated using BRIG.

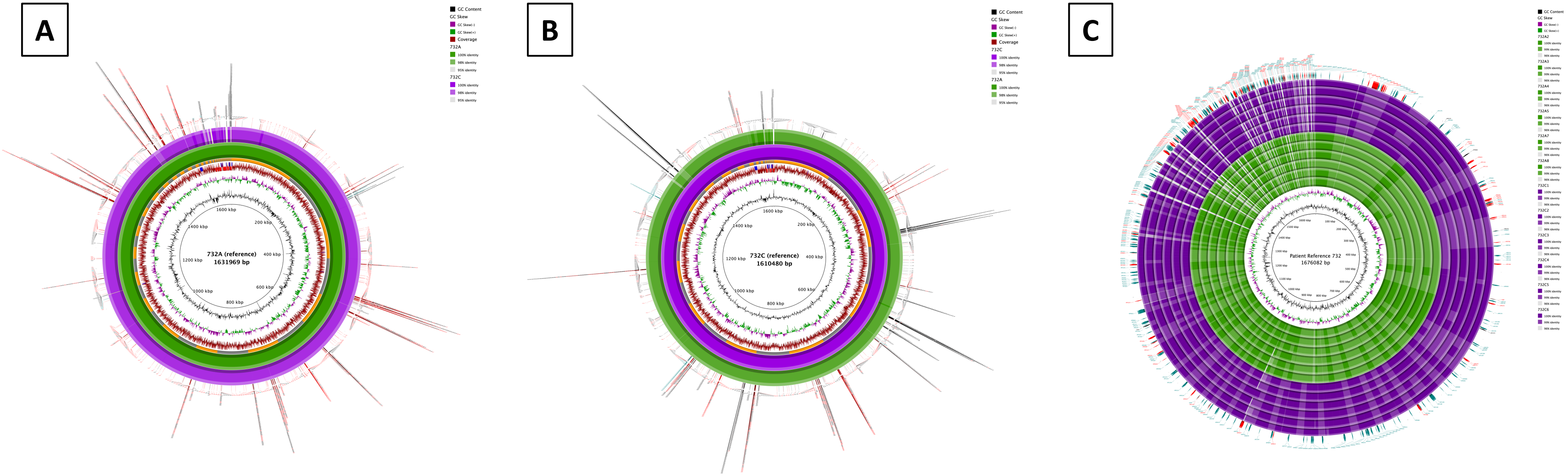


**Suppl. Fig. 15**. Representative whole genome alignment for *H. pylori* strains isolated from patient 732. Whole genome alignments of the consensus genomes generated by deep population sequencing of antrum and corpus *H. pylori* populations, using antrum (A) or corpus (B) consensus genome as the reference. Panel C depicts the alignment of colony isolate assembled genomes against the patient reference. Colour intensity of the outermost ring indicates percentage identity between antrum and corpus consensus genomes. Positions of contigs within the assembled reference genome are shown as a ring alternating in colour between grey and orange. Peripheral ticks indicate positions of SNPs between antrum and corpus genomes identified by whole genome alignment (black), read mapping (teal), or both methods (red) in figures A-B. Plots generated using BRIG.

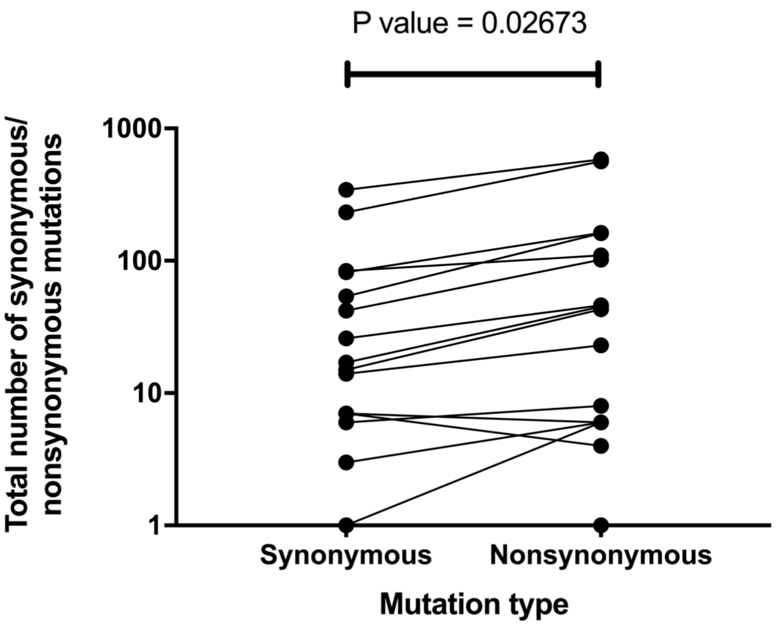


**Suppl. Fig. 16**. The validated (red) SNPs (Fig. 2A-B; Suppl. Fig 2,3,4,5,6A-B, 7, 8A-B, 9, 10A-B, 11, 12A-B, 13A-B, 14A-B, 15A-B) identified by the combined whole genome alignment and alternative stomach region mapping methodologies (depicted in Suppl. Fig. 1) were used to identify the number of synonymous or nonsynonymous mutations. A paired t-test comparing the total numbers of synonymous and nonsynonymous mutations between the paired populations was carried out.

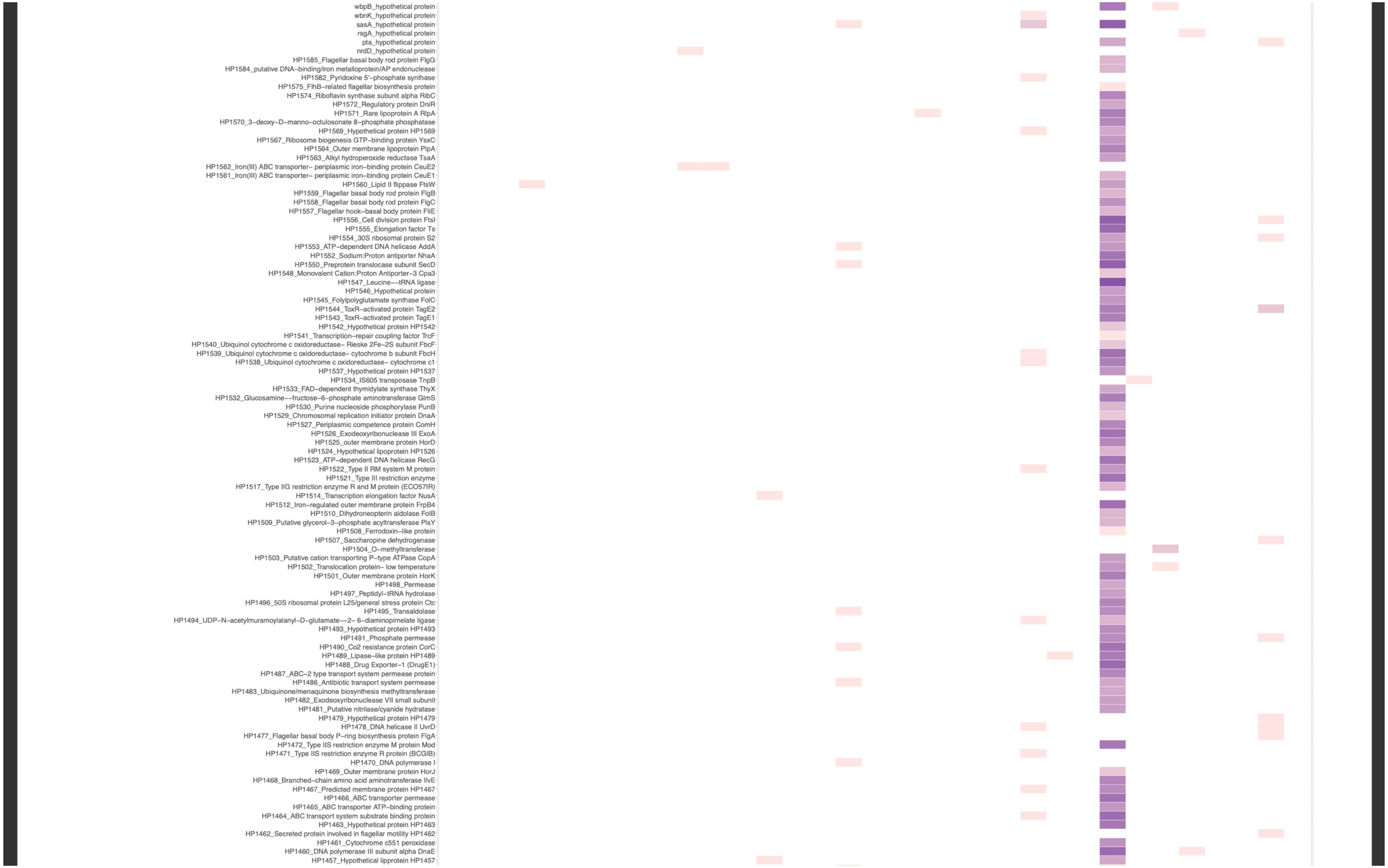
wbpB_hypothetical protein
wbnK_hypothetical protein
sasA_hypothetical protein
rsgA_hypothetical protein
pta_hypothetical protein
nrdD_hypothetical protein
HP1585_Flagellar basal body rod protein FlgG
HP1584_putative DNA−binding/iron metalloprotein/AP endonuclease
HP1582_Pyridoxine 5'−phosphate synthase
HP1575_FlhB−related flagellar biosynthesis protein
HP1574_Riboflavin synthase subunit alpha RibC
HP1572_Regulatory protein DniR
HP1571_Rare lipoprotein A RlpA
HP1570_3−deoxy−D−manno−octulosonate 8−phosphate phosphatase
HP1569_Hypothetical protein HP1569
HP1567_Ribosome biogenesis GTP−binding protein YsxC
HP1564_Outer membrane lipoprotein PlpA
HP1563_Alkyl hydroperoxide reductase TsaA
HP1562_Iron(III) ABC transporter− periplasmic iron−binding protein CeuE2
HP1561_Iron(III) ABC transporter− periplasmic iron−binding protein CeuE1
HP1560_Lipid II flippase FtsW
HP1559_Flagellar basal body rod protein FlgB
HP1558_Flagellar basal body rod protein FlgC
HP1557_Flagellar hook−basal body protein FliE
HP1556_Cell division protein FtsI
HP1555_Elongation factor Ts
HP1554_30S ribosomal protein S2
HP1553_ATP−dependent DNA helicase AddA
HP1552_Sodium:Proton antiporter NhaA
HP1550_Preprotein translocase subunit SecD
HP1548_Monovalent Cation:Proton Antiporter−3 Cpa3
HP1547_Leucine−−tRNA ligase
HP1546_Hypothetical protein
HP1545_Folylpolyglutamate synthase FolC
HP1544_ToxR−activated protein TagE2
HP1543_ToxR−activated protein TagE1
HP1542_Hypothetical protein HP1542
HP1541_Transcription−repair coupling factor TrcF
HP1540_Ubiquinol cytochrome c oxidoreductase− Rieske 2Fe−2S subunit FbcF
HP1539_Ubiquinol cytochrome c oxidoreductase− cytochrome b subunit FbcH
HP1538_Ubiquinol cytochrome c oxidoreductase− cytochrome c1
HP1537_Hypothetical protein HP1537
HP1534_IS605 transposase TnpB
HP1533_FAD−dependent thymidylate synthase ThyX
HP1532_Glucosamine−−fructose−6−phosphate aminotransferase GlmS
HP1530_Purine nucleoside phosphorylase PunB
HP1529_Chromosomal replication initiator protein DnaA
HP1527_Periplasmic competence protein ComH
HP1526_Exodeoxyribonuclease III ExoA
HP1525_outer membrane protein HorD
HP1524_Hypothetical lipoprotein HP1526
HP1523_ATP−dependent DNA helicase RecG
HP1522_Type II RM system M protein
HP1521_Type III restriction enzyme
HP1517_Type IIG restriction enzyme R and M protein (ECO57IR)
HP1514_Transcription elongation factor NusA
HP1512_Iron−regulated outer membrane protein FrpB4
HP1510_Dihydroneopterin aldolase FolB
HP1509_Putative glycerol−3−phosphate acyltransferase PlsY
HP1508_Ferrodoxin−like protein
HP1507_Saccharopine dehydrogenase
HP1504_O−methyltransferase
HP1503_Putative cation transporting P−type ATPase CopA
HP1502_Translocation protein− low temperature
HP1501_Outer membrane protein HorK
HP1498_Permease
HP1497_Peptidyl−tRNA hydrolase
HP1496_50S ribosomal protein L25/general stress protein Ctc
HP1495_Transaldolase
HP1494_UDP−N−acetylmuramoylalanyl−D−glutamate−−2− 6−diaminopimelate ligase
HP1493_Hypothetical protein HP1493
HP1491_Phosphate permease
HP1490_Co2 resistance protein CorC
HP1489_Lipase−like protein HP1489
HP1488_Drug Exporter−1 (DrugE1)
HP1487_ABC−2 type transport system permease protein
HP1486_Antibiotic transport system permease
HP1483_Ubiquinone/menaquinone biosynthesis methyltransferase
HP1482_Exodeoxyribonuclease VII small subunit
HP1481_Putative nitrilase/cyanide hydratase
HP1479_Hypothetical protein HP1479
HP1478_DNA helicase II UvrD
HP1477_Flagellar basal body P−ring biosynthesis protein FlgA
HP1472_Type IIS restriction enzyme M protein Mod
HP1471_Type IIS restriction enzyme R protein (BCGIB)
HP1470_DNA polymerase I
HP1469_Outer membrane protein HorJ
HP1468_Branched−chain amino acid aminotransferase IlvE
HP1467_Predicted membrane protein HP1467
HP1466_ABC transporter permease
HP1465_ABC transporter ATP−binding protein
HP1464_ABC transport system substrate binding protein
HP1463_Hypothetical protein HP1463
HP1462_Secreted protein involved in flagellar motility HP1462
HP1461_Cytochrome c551 peroxidase
HP1460_DNA polymerase III subunit alpha DnaE
HP1457_Hypothetical lipoprotein HP1457

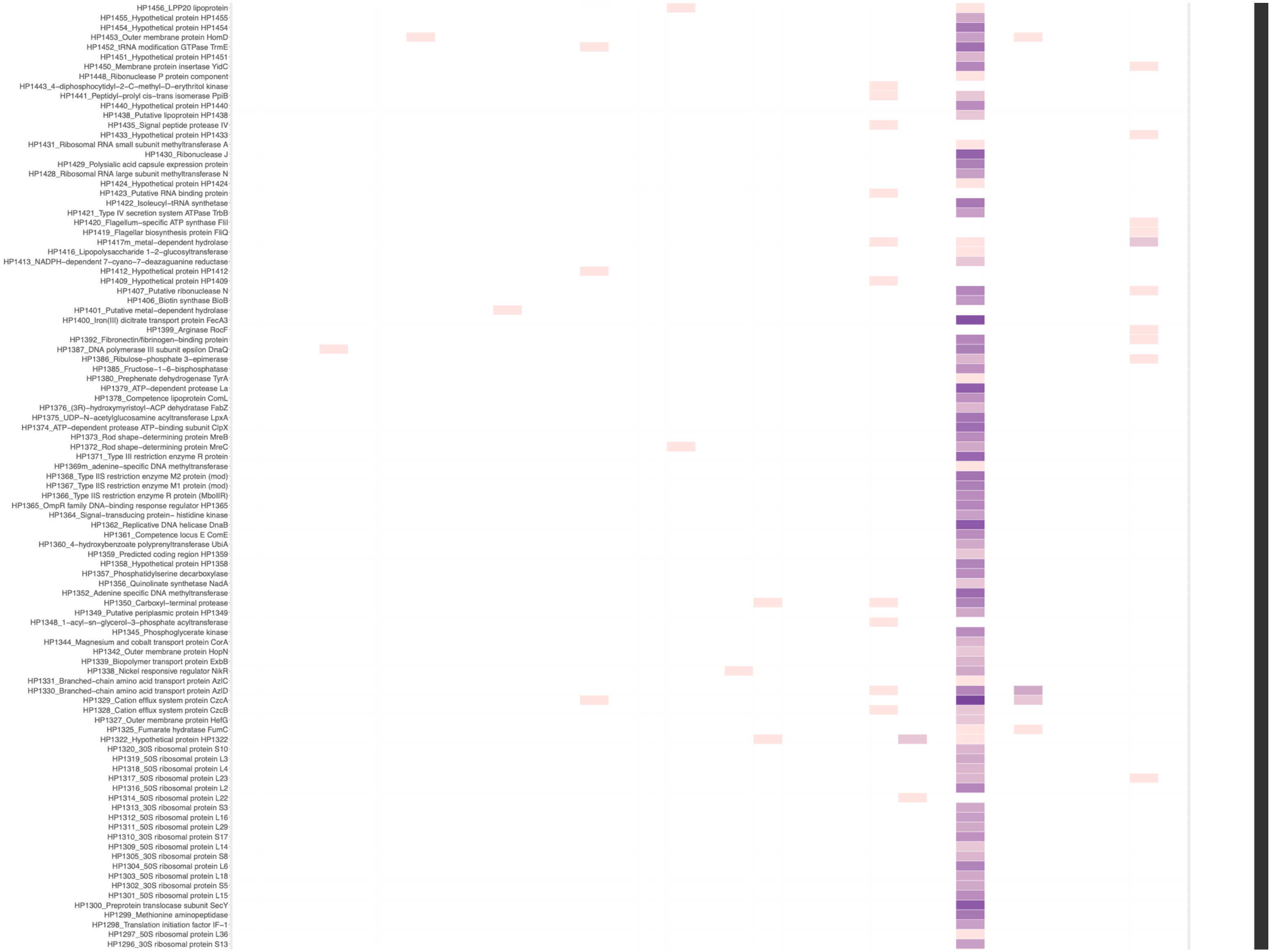

HP1456_LPP20 lipoprotein
HP1455_Hypothetical protein HP1455
HP1454_Hypothetical protein HP1454
HP1453_Outer membrane protein HomD
HP1452_tRNA modification GTPase TrmE
HP1451_Hypothetical protein HP1451
HP1450_Membrane protein insertase YidC
HP1448_Ribonuclease P protein component
HP1443_4–diphosphocytidyl–2–C–methyl–D–erythritol kinase
HP1441_Peptidyl–prolyl cis–trans isomerase PpiB
HP1440_Hypothetical protein HP1440
HP1438_Putative lipoprotein HP1438
HP1435_Signal peptide protease IV
HP1433_Hypothetical protein HP1433
HP1431_Ribosomal RNA small subunit methyltransferase A
HP1430_Ribonuclease J
HP1429_Polysialic acid capsule expression protein
HP1428_Ribosomal RNA large subunit methyltransferase N
HP1424_Hypothetical protein HP1424
HP1423_Putative RNA binding protein
HP1422_Isoleucyl–tRNA synthetase
HP1421_Type IV secretion system ATPase TrbB
HP1420_Flagellum–specific ATP synthase FliI
HP1419_Flagellar biosynthesis protein FliQ
HP1417m_metal–dependent hydrolase
HP1416_Lipopolysaccharide 1–2–glucosyltransferase
HP1413_NADPH–dependent 7–cyano–7–deazaguanine reductase
HP1412_Hypothetical protein HP1412
HP1409_Hypothetical protein HP1409
HP1407_Putative ribonuclease N
HP1406_Biotin synthase BioB
HP1401_Putative metal–dependent hydrolase
HP1400_Iron(III) dicitrate transport protein FecA3
HP1399_Arginase RocF
HP1392_Fibronectin/fibrinogen–binding protein
HP1387_DNA polymerase III subunit epsilon DnaQ
HP1386_Ribulose–phosphate 3–epimerase
HP1385_Fructose–1–6–bisphosphatase
HP1380_Prephenate dehydrogenase TyrA
HP1379_ATP–dependent protease La
HP1378_Competence lipoprotein ComL
HP1376_(3R)–hydroxymyristoyl–ACP dehydratase FabZ
HP1375_UDP–N–acetylglucosamine acyltransferase LpxA
HP1374_ATP–dependent protease ATP–binding subunit ClpX
HP1373_Rod shape–determining protein MreB
HP1372_Rod shape–determining protein MreC
HP1371_Type III restriction enzyme R protein
HP1369m_adenine–specific DNA methyltransferase
HP1368_Type IIS restriction enzyme M2 protein (mod)
HP1367_Type IIS restriction enzyme M1 protein (mod)
HP1366_Type IIS restriction enzyme R protein (MboIIR)
HP1365_OmpR family DNA–binding response regulator HP1365
HP1364_Signal–transducing protein– histidine kinase
HP1362_Replicative DNA helicase DnaB
HP1361_Competence locus E ComE
HP1360_4–hydroxybenzoate polyprenyltransferase UbiA
HP1359_Predicted coding region HP1359
HP1358_Hypothetical protein HP1358
HP1357_Phosphatidylserine decarboxylase
HP1356_Quinolinate synthetase NadA
HP1352_Adenine specific DNA methyltransferase
HP1350_Carboxyl–terminal protease
HP1349_Putative periplasmic protein HP1349
HP1348_1–acyl–sn–glycerol–3–phosphate acyltransferase
HP1345_Phosphoglycerate kinase
HP1344_Magnesium and cobalt transport protein CorA
HP1342_Outer membrane protein HopN
HP1339_Biopolymer transport protein ExbB
HP1338_Nickel responsive regulator NikR
HP1331_Branched–chain amino acid transport protein AzlC
HP1330_Branched–chain amino acid transport protein AzlD
HP1329_Cation efflux system protein CzcA
HP1328_Cation efflux system protein CzcB
HP1327_Outer membrane protein HefG
HP1325_Fumarate hydratase FumC
HP1322_Hypothetical protein HP1322
HP1320_30S ribosomal protein S10
HP1319_50S ribosomal protein L3
HP1318_50S ribosomal protein L4
HP1317_50S ribosomal protein L23
HP1316_50S ribosomal protein L2
HP1314_50S ribosomal protein L22
HP1313_30S ribosomal protein S3
HP1312_50S ribosomal protein L16
HP1311_50S ribosomal protein L29
HP1310_30S ribosomal protein S17
HP1309_50S ribosomal protein L14
HP1305_30S ribosomal protein S8
HP1304_50S ribosomal protein L6
HP1303_50S ribosomal protein L18
HP1302_30S ribosomal protein S5
HP1301_50S ribosomal protein L15
HP1300_Preprotein translocase subunit SecY
HP1299_Methionine aminopeptidase
HP1298_Translation initiation factor IF–1
HP1297_50S ribosomal protein L36
HP1296_30S ribosomal protein S13

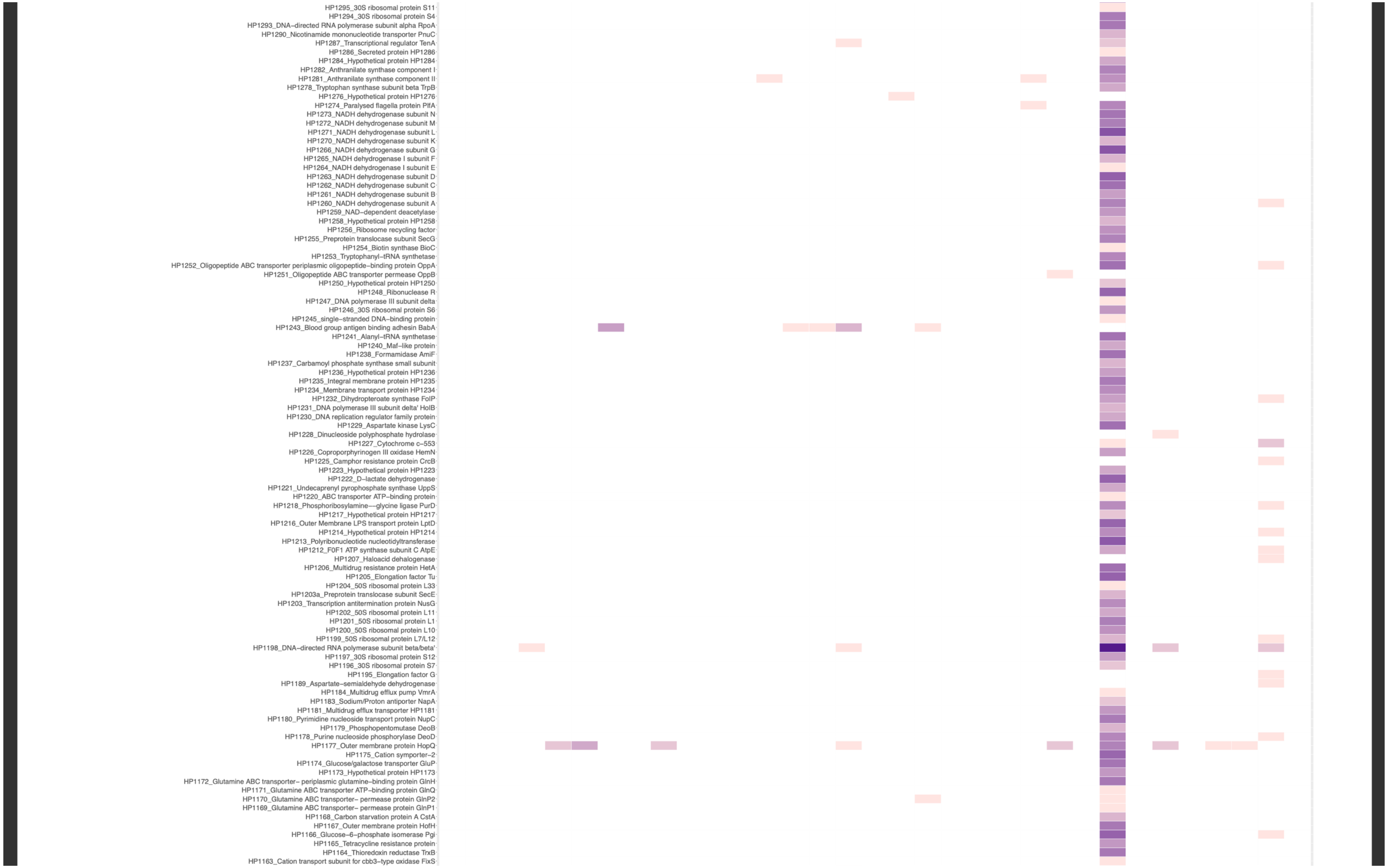
HP1295_30S ribosomal protein S11
HP1294_30S ribosomal protein S4
HP1293_DNA−directed RNA polymerase subunit alpha RpoA
HP1290_Nicotinamide mononucleotide transporter PnuC
HP1287_Transcriptional regulator TenA
HP1286_Secreted protein HP1286
HP1284_Hypothetical protein HP1284
HP1282_Anthranilate synthase component I
HP1281_Anthranilate synthase component II
HP1278_Tryptophan synthase subunit beta TrpB
HP1276_Hypothetical protein HP1276
HP1274_Paralysed flagella protein PflA
HP1273_NADH dehydrogenase subunit N
HP1272_NADH dehydrogenase subunit M
HP1271_NADH dehydrogenase subunit L
HP1270_NADH dehydrogenase subunit K
HP1266_NADH dehydrogenase subunit G
HP1265_NADH dehydrogenase I subunit F
HP1264_NADH dehydrogenase I subunit E
HP1263_NADH dehydrogenase subunit D
HP1262_NADH dehydrogenase subunit C
HP1261_NADH dehydrogenase subunit B
HP1260_NADH dehydrogenase subunit A
HP1259_NAD−dependent deacetylase
HP1258_Hypothetical protein HP1258
HP1256_Ribosome recycling factor
HP1255_Preprotein translocase subunit SecG
HP1254_Biotin synthase BioC
HP1253_Tryptophanyl−tRNA synthetase
HP1252_Oligopeptide ABC transporter periplasmic oligopeptide−binding protein OppA
HP1251_Oligopeptide ABC transporter permease OppB
HP1250_Hypothetical protein HP1250
HP1248_Ribonuclease R
HP1247_DNA polymerase III subunit delta
HP1246_30S ribosomal protein S6
HP1245_single−stranded DNA−binding protein
HP1243_Blood group antigen binding adhesin BabA
HP1241_Alanyl−tRNA synthetase
HP1240_Maf−like protein
HP1238_Formamidase AmiF
HP1237_Carbamoyl phosphate synthase small subunit
HP1236_Hypothetical protein HP1236
HP1235_Integral membrane protein HP1235
HP1234_Membrane transport protein HP1234
HP1232_Dihydropteroate synthase FolP
HP1231_DNA polymerase III subunit delta' HolB
HP1230_DNA replication regulator family protein
HP1229_Aspartate kinase LysC
HP1228_Dinucleoside polyphosphate hydrolase
HP1227_Cytochrome c−553
HP1226_Coproporphyrinogen III oxidase HemN
HP1225_Camphor resistance protein CrcB
HP1223_Hypothetical protein HP1223
HP1222_D−lactate dehydrogenase
HP1221_Undecaprenyl pyrophosphate synthase UppS
HP1220_ABC transporter ATP−binding protein
HP1218_Phosphoribosylamine−−glycine ligase PurD
HP1217_Hypothetical protein HP1217
HP1216_Outer Membrane LPS transport protein LptD
HP1214_Hypothetical protein HP1214
HP1213_Polyribonucleotide nucleotidyltransferase
HP1212_F0F1 ATP synthase subunit C AtpE
HP1207_Haloacid dehalogenase
HP1206_Multidrug resistance protein HetA
HP1205_Elongation factor Tu
HP1204_50S ribosomal protein L33
HP1203a_Preprotein translocase subunit SecE
HP1203_Transcription antitermination protein NusG
HP1202_50S ribosomal protein L11
HP1201_50S ribosomal protein L1
HP1200_50S ribosomal protein L10
HP1199_50S ribosomal protein L7/L12
HP1198_DNA−directed RNA polymerase subunit beta/beta'
HP1197_30S ribosomal protein S12
HP1196_30S ribosomal protein S7
HP1195_Elongation factor G
HP1189_Aspartate−semialdehyde dehydrogenase
HP1184_Multidrug efflux pump VmrA
HP1183_Sodium/Proton antiporter NapA
HP1181_Multidrug efflux transporter HP1181
HP1180_Pyrimidine nucleoside transport protein NupC
HP1179_Phosphopentomutase DeoB
HP1178_Purine nucleoside phosphorylase DeoD
HP1177_Outer membrane protein HopQ
HP1175_Cation symporter−2
HP1174_Glucose/galactose transporter GluP
HP1173_Hypothetical protein HP1173
HP1172_Glutamine ABC transporter− periplasmic glutamine−binding protein GlnH
HP1171_Glutamine ABC transporter ATP−binding protein GlnQ
HP1170_Glutamine ABC transporter− permease protein GlnP2
HP1169_Glutamine ABC transporter− permease protein GlnP1
HP1168_Carbon starvation protein A CstA
HP1167_Outer membrane protein HofH
HP1166_Glucose−6−phosphate isomerase Pgi
HP1165_Tetracycline resistance protein
HP1164_Thioredoxin reductase TrxB
HP1163_Cation transport subunit for cbb3−type oxidase FixS

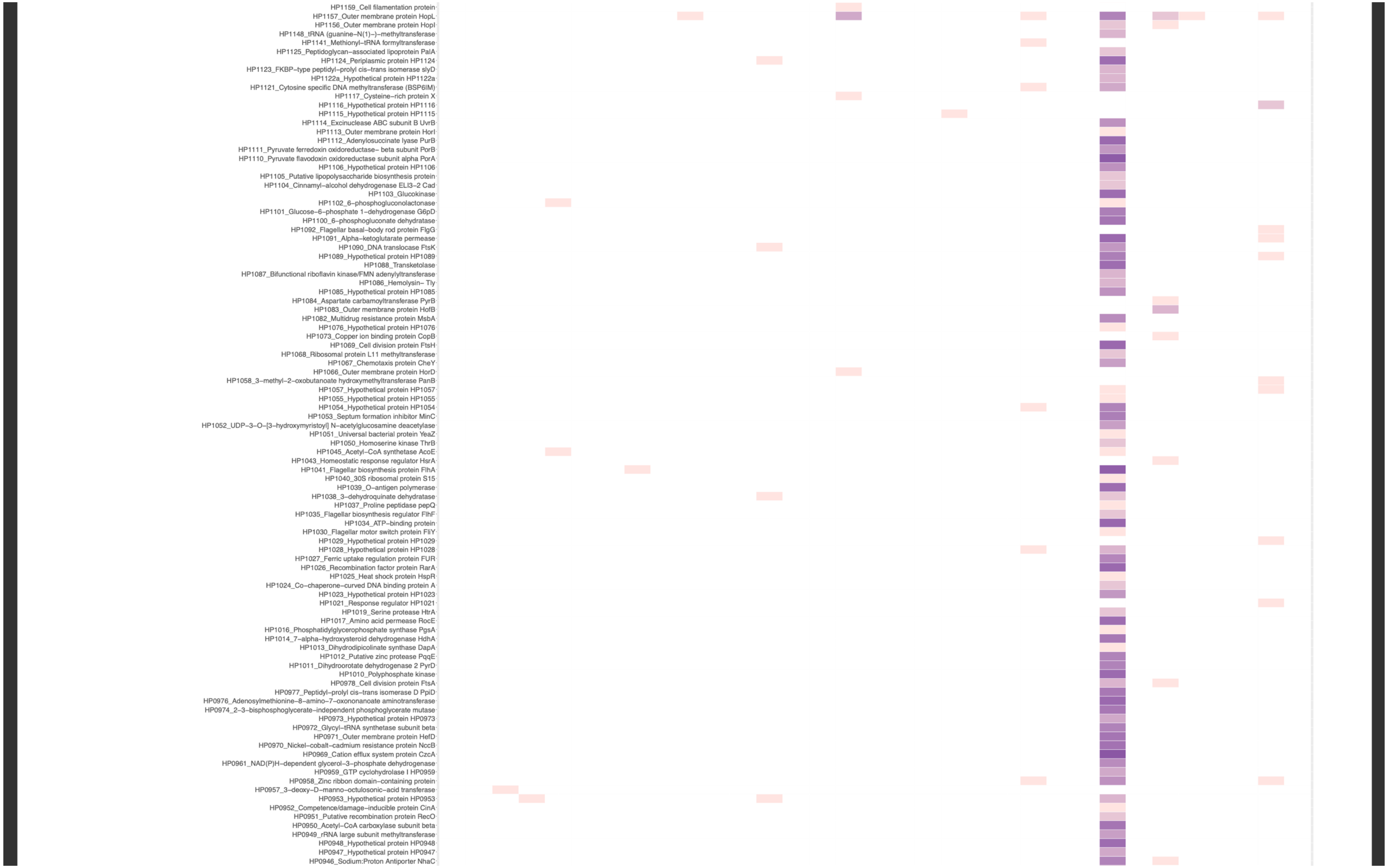

HP1159_Cell filamentation protein
HP1157_Outer membrane protein HopL
HP1156_Outer membrane protein HopI
HP1148_tRNA (guanine–N(1)–)–methyltransferase
HP1141_Methionyl–tRNA formyltransferase
HP1125_Peptidoglycan–associated lipoprotein PalA
HP1124_Periplasmic protein HP1124
HP1123_FKBP–type peptidyl–prolyl cis–trans isomerase slyD
HP1122a_Hypothetical protein HP1122a
HP1121_Cytosine specific DNA methyltransferase (BSP6IM)
HP1117_Cysteine–rich protein X
HP1116_Hypothetical protein HP1116
HP1115_Hypothetical protein HP1115
HP1114_Excinuclease ABC subunit B UvrB
HP1113_Outer membrane protein HorI
HP1112_Adenylosuccinate lyase PurB
HP1111_Pyruvate ferredoxin oxidoreductase– beta subunit PorB
HP1110_Pyruvate flavodoxin oxidoreductase subunit alpha PorA
HP1106_Hypothetical protein HP1106
HP1105_Putative lipopolysaccharide biosynthesis protein
HP1104_Cinnamyl–alcohol dehydrogenase ELI3–2 Cad
HP1103_Glucokinase
HP1102_6–phosphogluconolactonase
HP1101_Glucose–6–phosphate 1–dehydrogenase G6pD
HP1100_6–phosphogluconate dehydratase
HP1092_Flagellar basal–body rod protein FlgG
HP1091_Alpha–ketoglutarate permease
HP1090_DNA translocase FtsK
HP1089_Hypothetical protein HP1089
HP1088_Transketolase
HP1087_Bifunctional riboflavin kinase/FMN adenylyltransferase
HP1086_Hemolysin– Tly
HP1085_Hypothetical protein HP1085
HP1084_Aspartate carbamoyltransferase PyrB
HP1083_Outer membrane protein HofB
HP1082_Multidrug resistance protein MsbA
HP1076_Hypothetical protein HP1076
HP1073_Copper ion binding protein CopB
HP1069_Cell division protein FtsH
HP1068_Ribosomal protein L11 methyltransferase
HP1067_Chemotaxis protein CheY
HP1066_Outer membrane protein HorD
HP1058_3–methyl–2–oxobutanoate hydroxymethyltransferase PanB
HP1057_Hypothetical protein HP1057
HP1055_Hypothetical protein HP1055
HP1054_Hypothetical protein HP1054
HP1053_Septum formation inhibitor MinC
HP1052_UDP–3–O–[3–hydroxymyristoyl] N–acetylglucosamine deacetylase
HP1051_Universal bacterial protein YeaZ
HP1050_Homoserine kinase ThrB
HP1045_Acetyl–CoA synthetase AcoE
HP1043_Homeostatic response regulator HsrA
HP1041_Flagellar biosynthesis protein FlhA
HP1040_30S ribosomal protein S15
HP1039_O–antigen polymerase
HP1038_3–dehydroquinate dehydratase
HP1037_Proline peptidase pepQ
HP1035_Flagellar biosynthesis regulator FlhF
HP1034_ATP–binding protein
HP1030_Flagellar motor switch protein FliY
HP1029_Hypothetical protein HP1029
HP1028_Hypothetical protein HP1028
HP1027_Ferric uptake regulation protein FUR
HP1026_Recombination factor protein RarA
HP1025_Heat shock protein HspR
HP1024_Co–chaperone–curved DNA binding protein A
HP1023_Hypothetical protein HP1023
HP1021_Response regulator HP1021
HP1019_Serine protease HtrA
HP1017_Amino acid permease RocE
HP1016_Phosphatidylglycerophosphate synthase PgsA
HP1014_7–alpha–hydroxysteroid dehydrogenase HdhA
HP1013_Dihydrodipicolinate synthase DapA
HP1012_Putative zinc protease PqqE
HP1011_Dihydroorotate dehydrogenase 2 PyrD
HP1010_Polyphosphate kinase
HP0978_Cell division protein FtsA
HP0977_Peptidyl–prolyl cis–trans isomerase D PpiD
HP0976_Adenosylmethionine–8–amino–7–oxononanoate aminotransferase
HP0974_2–3–bisphosphoglycerate–independent phosphoglycerate mutase
HP0973_Hypothetical protein HP0973
HP0972_Glycyl–tRNA synthetase subunit beta
HP0971_Outer membrane protein HefD
HP0970_Nickel–cobalt–cadmium resistance protein NccB
HP0969_Cation efflux system protein CzcA
HP0961_NAD(P)H–dependent glycerol–3–phosphate dehydrogenase
HP0959_GTP cyclohydrolase I HP0959
HP0958_Zinc ribbon domain–containing protein
HP0957_3–deoxy–D–manno–octulosonic–acid transferase
HP0953_Hypothetical protein HP0953
HP0952_Competence/damage–inducible protein CinA
HP0951_Putative recombination protein RecO
HP0950_Acetyl–CoA carboxylase subunit beta
HP0949_rRNA large subunit methyltransferase
HP0948_Hypothetical protein HP0948
HP0947_Hypothetical protein HP0947
HP0946_Sodium:Proton Antiporter NhaC

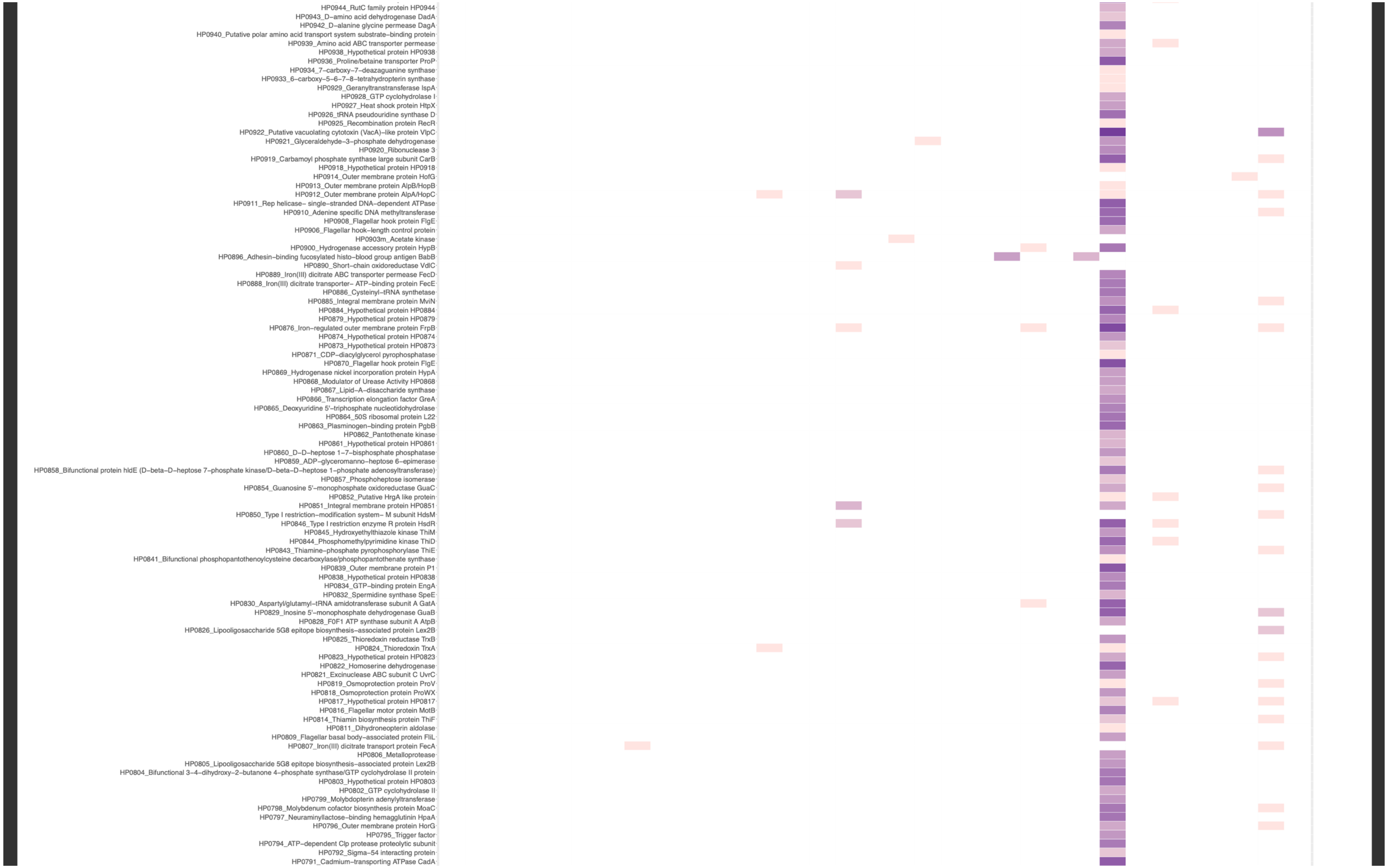
HP0944_RutC family protein HP0944
HP0943_D−amino acid dehydrogenase DadA
HP0942_D−alanine glycine permease DagA
HP0940_Putative polar amino acid transport system substrate−binding protein
HP0939_Amino acid ABC transporter permease
HP0938_Hypothetical protein HP0938
HP0936_Proline/betaine transporter ProP
HP0934_7−carboxy−7−deazaguanine synthase
HP0933_6−carboxy−5−6−7−8−tetrahydropterin synthase
HP0929_Geranyltranstransferase IspA
HP0928_GTP cyclohydrolase I
HP0927_Heat shock protein HtpX
HP0926_tRNA pseudouridine synthase D
HP0925_Recombination protein RecR
HP0922_Putative vacuolating cytotoxin (VacA)−like protein VlpC
HP0921_Glyceraldehyde−3−phosphate dehydrogenase
HP0920_Ribonuclease 3
HP0919_Carbamoyl phosphate synthase large subunit CarB
HP0918_Hypothetical protein HP0918
HP0914_Outer membrane protein HofG
HP0913_Outer membrane protein AlpB/HopB
HP0912_Outer membrane protein AlpA/HopC
HP0911_Rep helicase− single−stranded DNA−dependent ATPase
HP0910_Adenine specific DNA methyltransferase
HP0908_Flagellar hook protein FlgE
HP0906_Flagellar hook−length control protein
HP0903m_Acetate kinase
HP0900_Hydrogenase accessory protein HypB
HP0896_Adhesin−binding fucosylated histo−blood group antigen BabB
HP0890_Short−chain oxidoreductase VdlC
HP0889_Iron(III) dicitrate ABC transporter permease FecD
HP0888_Iron(III) dicitrate transporter− ATP−binding protein FecE
HP0886_Cysteinyl−tRNA synthetase
HP0885_Integral membrane protein MviN
HP0884_Hypothetical protein HP0884
HP0879_Hypothetical protein HP0879
HP0876_Iron−regulated outer membrane protein FrpB
HP0874_Hypothetical protein HP0874
HP0873_Hypothetical protein HP0873
HP0871_CDP−diacylglycerol pyrophosphatase
HP0870_Flagellar hook protein FlgE
HP0869_Hydrogenase nickel incorporation protein HypA
HP0868_Modulator of Urease Activity HP0868
HP0867_Lipid−A−disaccharide synthase
HP0866_Transcription elongation factor GreA
HP0865_Deoxyuridine 5'−triphosphate nucleotidohydrolase
HP0864_50S ribosomal protein L22
HP0863_Plasminogen−binding protein PgbB
HP0862_Pantothenate kinase
HP0861_Hypothetical protein HP0861
HP0860_D−D−heptose 1−7−bisphosphate phosphatase
HP0859_ADP−glyceromanno−heptose 6−epimerase
HP0858_Bifunctional protein hldE (D−beta−D−heptose 7−phosphate kinase/D−beta−D−heptose 1−phosphate adenosyltransferase)
HP0857_Phosphoheptose isomerase
HP0854_Guanosine 5'−monophosphate oxidoreductase GuaC
HP0852_Putative HrgA like protein
HP0851_Integral membrane protein HP0851
HP0850_Type I restriction−modification system− M subunit HdsM
HP0846_Type I restriction enzyme R protein HsdR
HP0845_Hydroxyethylthiazole kinase ThiM
HP0844_Phosphomethylpyrimidine kinase ThiD
HP0843_Thiamine−phosphate pyrophosphorylase ThiE
HP0841_Bifunctional phosphopantothenoylcysteine decarboxylase/phosphopantothenate synthase
HP0839_Outer membrane protein P1
HP0838_Hypothetical protein HP0838
HP0834_GTP−binding protein EngA
HP0832_Spermidine synthase SpeE
HP0830_Aspartyl/glutamyl−tRNA amidotransferase subunit A GatA
HP0829_Inosine 5'−monophosphate dehydrogenase GuaB
HP0828_F0F1 ATP synthase subunit A AtpB
HP0826_Lipooligosaccharide 5G8 epitope biosynthesis−associated protein Lex2B
HP0825_Thioredoxin reductase TrxB
HP0824_Thioredoxin TrxA
HP0823_Hypothetical protein HP0823
HP0822_Homoserine dehydrogenase
HP0821_Excinuclease ABC subunit C UvrC
HP0819_Osmoprotection protein ProV
HP0818_Osmoprotection protein ProWX
HP0817_Hypothetical protein HP0817
HP0816_Flagellar motor protein MotB
HP0814_Thiamin biosynthesis protein ThiF
HP0811_Dihydroneopterin aldolase
HP0809_Flagellar basal body−associated protein FliL
HP0807_Iron(III) dicitrate transport protein FecA
HP0806_Metalloprotease
HP0805_Lipooligosaccharide 5G8 epitope biosynthesis−associated protein Lex2B
HP0804_Bifunctional 3−4−dihydroxy−2−butanone 4−phosphate synthase/GTP cyclohydrolase II protein
HP0803_Hypothetical protein HP0803
HP0802_GTP cyclohydrolase II
HP0799_Molybdopterin adenylyltransferase
HP0798_Molybdenum cofactor biosynthesis protein MoaC
HP0797_Neuraminyllactose−binding hemagglutinin HpaA
HP0796_Outer membrane protein HorG
HP0795_Trigger factor
HP0794_ATP−dependent Clp protease proteolytic subunit
HP0792_Sigma−54 interacting protein
HP0791_Cadmium−transporting ATPase CadA

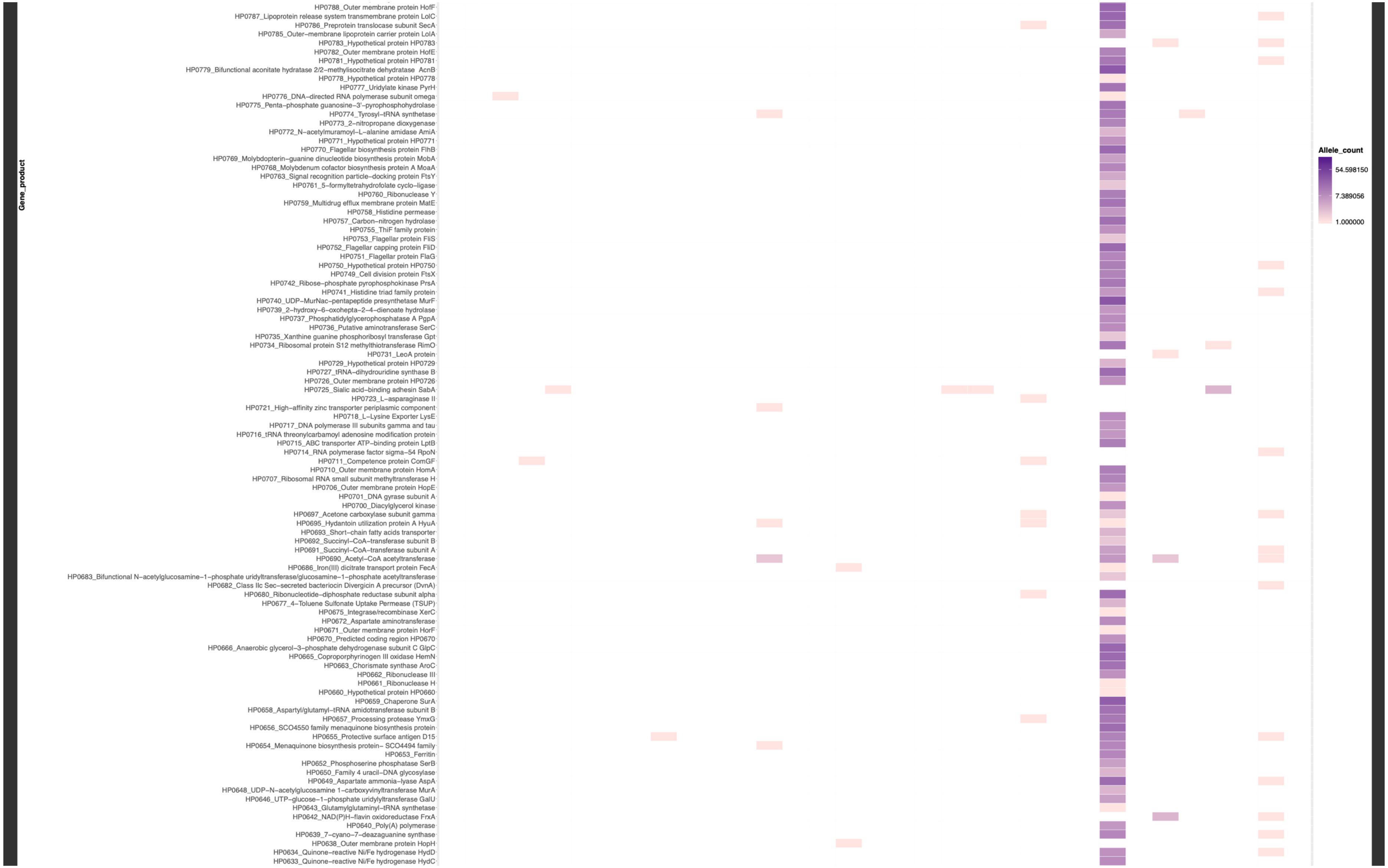
Gene_product
HP0788_Outer membrane protein HofF
HP0787_Lipoprotein release system transmembrane protein LolC
HP0786_Preprotein translocase subunit SecA
HP0785_Outer−membrane lipoprotein carrier protein LolA
HP0783_Hypothetical protein HP0783
HP0782_Outer membrane protein HofE
HP0781_Hypothetical protein HP0781
HP0779_Bifunctional aconitate hydratase 2/2−methylisocitrate dehydratase AcnB
HP0778_Hypothetical protein HP0778
HP0777_Uridylate kinase PyrH
HP0776_DNA−directed RNA polymerase subunit omega
HP0775_Penta−phosphate guanosine−3'−pyrophosphohydrolase
HP0774_Tyrosyl−tRNA synthetase
HP0773_2−nitropropane dioxygenase
HP0772_N−acetylmuramoyl−L−alanine amidase AmiA
HP0771_Hypothetical protein HP0771
HP0770_Flagellar biosynthesis protein FlhB
HP0769_Molybdopterin−guanine dinucleotide biosynthesis protein MobA
HP0768_Molybdenum cofactor biosynthesis protein A MoaA
HP0763_Signal recognition particle−docking protein FtsY
HP0761_5−formyltetrahydrofolate cyclo−ligase
HP0760_Ribonuclease Y
HP0759_Multidrug efflux membrane protein MatE
HP0758_Histidine permease
HP0757_Carbon−nitrogen hydrolase
HP0755_ThiF family protein
HP0753_Flagellar protein FliS
HP0752_Flagellar capping protein FliD
HP0751_Flagellar protein FlaG
HP0750_Hypothetical protein HP0750
HP0749_Cell division protein FtsX
HP0742_Ribose−phosphate pyrophosphokinase PrsA
HP0741_Histidine triad family protein
HP0740_UDP−MurNac−pentapeptide presynthetase MurF
HP0739_2−hydroxy−6−oxohepta−2−4−dienoate hydrolase
HP0737_Phosphatidylglycerophosphatase A PgpA
HP0736_Putative aminotransferase SerC
HP0735_Xanthine guanine phosphoribosyl transferase Gpt
HP0734_Ribosomal protein S12 methylthiotransferase RimO
HP0731_LeoA protein
HP0729_Hypothetical protein HP0729
HP0727_tRNA−dihydrouridine synthase B
HP0726_Outer membrane protein HP0726
HP0725_Sialic acid−binding adhesin SabA
HP0723_L−asparaginase II
HP0721_High−affinity zinc transporter periplasmic component
HP0718_L−Lysine Exporter LysE
HP0717_DNA polymerase III subunits gamma and tau
HP0716_tRNA threonylcarbamoyl adenosine modification protein
HP0715_ABC transporter ATP−binding protein LptB
HP0714_RNA polymerase factor sigma−54 RpoN
HP0711_Competence protein ComGF
HP0710_Outer membrane protein HomA
HP0707_Ribosomal RNA small subunit methyltransferase H
HP0706_Outer membrane protein HopE
HP0701_DNA gyrase subunit A
HP0700_Diacylglycerol kinase
HP0697_Acetone carboxylase subunit gamma
HP0695_Hydantoin utilization protein A HyuA
HP0693_Short−chain fatty acids transporter
HP0692_Succinyl−CoA−transferase subunit B
HP0691_Succinyl−CoA−transferase subunit A
HP0690_Acetyl−CoA acetyltransferase
HP0686_Iron(III) dicitrate transport protein FecA
HP0683_Bifunctional N−acetylglucosamine−1−phosphate uridyltransferase/glucosamine−1−phosphate acetyltransferase
HP0682_Class IIc Sec−secreted bacteriocin Divergicin A precursor (DvnA)
HP0680_Ribonucleotide−diphosphate reductase subunit alpha
HP0677_4−Toluene Sulfonate Uptake Permease (TSUP)
HP0675_Integrase/recombinase XerC
HP0672_Aspartate aminotransferase
HP0671_Outer membrane protein HorF
HP0670_Predicted coding region HP0670
HP0666_Anaerobic glycerol−3−phosphate dehydrogenase subunit C GlpC
HP0665_Coproporphyrinogen III oxidase HemN
HP0663_Chorismate synthase AroC
HP0662_Ribonuclease III
HP0661_Ribonuclease H
HP0660_Hypothetical protein HP0660
HP0659_Chaperone SurA
HP0658_Aspartyl/glutamyl−tRNA amidotransferase subunit B
HP0657_Processing protease YmxG
HP0656_SCO4550 family menaquinone biosynthesis protein
HP0655_Protective surface antigen D15
HP0654_Menaquinone biosynthesis protein− SCO4494 family
HP0653_Ferritin
HP0652_Phosphoserine phosphatase SerB
HP0650_Family 4 uracil−DNA glycosylase
HP0649_Aspartate ammonia−lyase AspA
HP0648_UDP−N−acetylglucosamine 1−carboxyvinyltransferase MurA
HP0646_UTP−glucose−1−phosphate uridylyltransferase GalU
HP0643_Glutamylglutaminyl−tRNA synthetase
HP0642_NAD(P)H−flavin oxidoreductase FrxA
HP0640_Poly(A) polymerase
HP0639_7−cyano−7−deazaguanine synthase
HP0638_Outer membrane protein HopH
HP0634_Quinone−reactive Ni/Fe hydrogenase HydD
HP0633_Quinone−reactive Ni/Fe hydrogenase HydC
Allele_count
54.598150
7.389056
1.000000

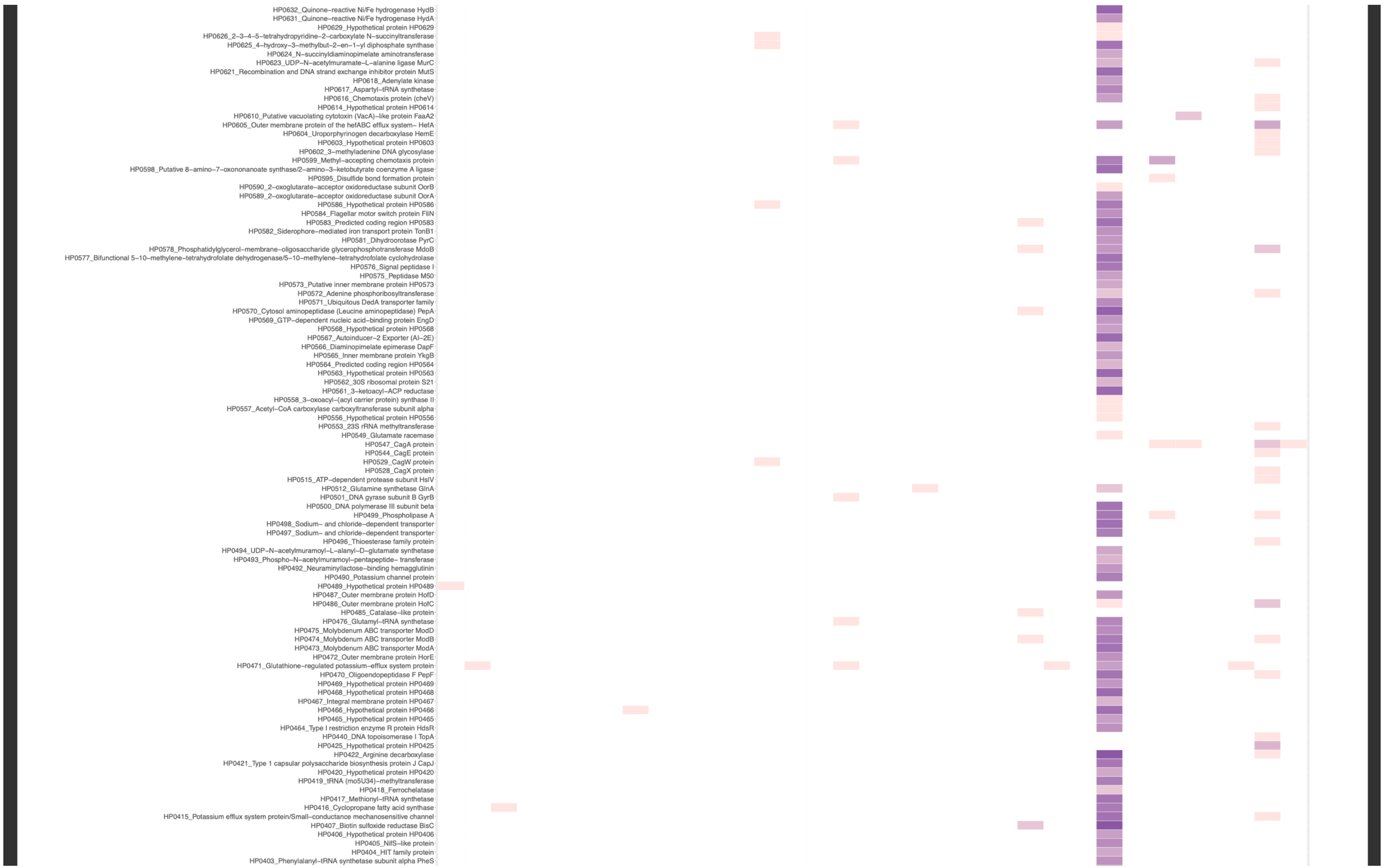
HP0632_Quinone–reactive Ni/Fe hydrogenase HydB
HP0631_Quinone–reactive Ni/Fe hydrogenase HydA
HP0629_Hypothetical protein HP0629
HP0626_2–3–4–5–tetrahydropyridine–2–carboxylate N–succinyltransferase
HP0625_4–hydroxy–3–methylbut–2–en–1–yl diphosphate synthase
HP0624_N–succinyldiaminopimelate aminotransferase
HP0623_UDP–N–acetylmuramate–L–alanine ligase MurC
HP0621_Recombination and DNA strand exchange inhibitor protein MutS
HP0618_Adenylate kinase
HP0617_Aspartyl–tRNA synthetase
HP0616_Chemotaxis protein (cheV)
HP0614_Hypothetical protein HP0614
HP0610_Putative vacuolating cytotoxin (VacA)–like protein FaaA2
HP0605_Outer membrane protein of the hefABC efflux system– HefA
HP0604_Uroporphyrinogen decarboxylase HemE
HP0603_Hypothetical protein HP0603
HP0602_3–methyladenine DNA glycosylase
HP0599_Methyl–accepting chemotaxis protein
HP0598_Putative 8–amino–7–oxononanoate synthase/2–amino–3–ketobutyrate coenzyme A ligase
HP0595_Disulfide bond formation protein
HP0590_2–oxoglutarate–acceptor oxidoreductase subunit OorB
HP0589_2–oxoglutarate–acceptor oxidoreductase subunit OorA
HP0586_Hypothetical protein HP0586
HP0584_Flagellar motor switch protein FliN
HP0583_Predicted coding region HP0583
HP0582_Siderophore–mediated iron transport protein TonB1
HP0581_Dihydroorotase PyrC
HP0578_Phosphatidylglycerol–membrane–oligosaccharide glycerophosphotransferase MdoB
HP0577_Bifunctional 5–10–methylene–tetrahydrofolate dehydrogenase/5–10–methylene–tetrahydrofolate cyclohydrolase
HP0576_Signal peptidase I
HP0575_Peptidase M50
HP0573_Putative inner membrane protein HP0573
HP0572_Adenine phosphoribosyltransferase
HP0571_Ubiquitous DedA transporter family
HP0570_Cytosol aminopeptidase (Leucine aminopeptidase) PepA
HP0569_GTP–dependent nucleic acid–binding protein EngD
HP0568_Hypothetical protein HP0568
HP0567_Autoinducer–2 Exporter (AI–2E)
HP0566_Diaminopimelate epimerase DapF
HP0565_Inner membrane protein YkgB
HP0564_Predicted coding region HP0564
HP0563_Hypothetical protein HP0563
HP0562_30S ribosomal protein S21
HP0561_3–ketoacyl–ACP reductase
HP0558_3–oxoacyl–(acyl carrier protein) synthase II
HP0557_Acetyl–CoA carboxylase carboxyltransferase subunit alpha
HP0556_Hypothetical protein HP0556
HP0553_23S rRNA methyltransferase
HP0549_Glutamate racemase
HP0547_CagA protein
HP0544_CagE protein
HP0529_CagW protein
HP0528_CagX protein
HP0515_ATP–dependent protease subunit HslV
HP0512_Glutamine synthetase GlnA
HP0501_DNA gyrase subunit B GyrB
HP0500_DNA polymerase III subunit beta
HP0499_Phospholipase A
HP0498_Sodium– and chloride–dependent transporter
HP0497_Sodium– and chloride–dependent transporter
HP0496_Thioesterase family protein
HP0494_UDP–N–acetylmuramoyl–L–alanyl–D–glutamate synthetase
HP0493_Phospho–N–acetylmuramoyl–pentapeptide– transferase
HP0492_Neuraminyllactose–binding hemagglutinin
HP0490_Potassium channel protein
HP0489_Hypothetical protein HP0489
HP0487_Outer membrane protein HofD
HP0486_Outer membrane protein HofC
HP0485_Catalase–like protein
HP0476_Glutamyl–tRNA synthetase
HP0475_Molybdenum ABC transporter ModD
HP0474_Molybdenum ABC transporter ModB
HP0473_Molybdenum ABC transporter ModA
HP0472_Outer membrane protein HorE
HP0471_Glutathione–regulated potassium–efflux system protein
HP0470_Oligoendopeptidase F PepF
HP0469_Hypothetical protein HP0469
HP0468_Hypothetical protein HP0468
HP0467_Integral membrane protein HP0467
HP0466_Hypothetical protein HP0466
HP0465_Hypothetical protein HP0465
HP0464_Type I restriction enzyme R protein HdsR
HP0440_DNA topoisomerase I TopA
HP0425_Hypothetical protein HP0425
HP0422_Arginine decarboxylase
HP0421_Type 1 capsular polysaccharide biosynthesis protein J CapJ
HP0420_Hypothetical protein HP0420
HP0419_tRNA (mo5U34)–methyltransferase
HP0418_Ferrochelatase
HP0417_Methionyl–tRNA synthetase
HP0416_Cyclopropane fatty acid synthase
HP0415_Potassium efflux system protein/Small–conductance mechanosensitive channel
HP0407_Biotin sulfoxide reductase BisC
HP0406_Hypothetical protein HP0406
HP0405_NifS–like protein
HP0404_HIT family protein
HP0403_Phenylalanyl–tRNA synthetase subunit alpha PheS

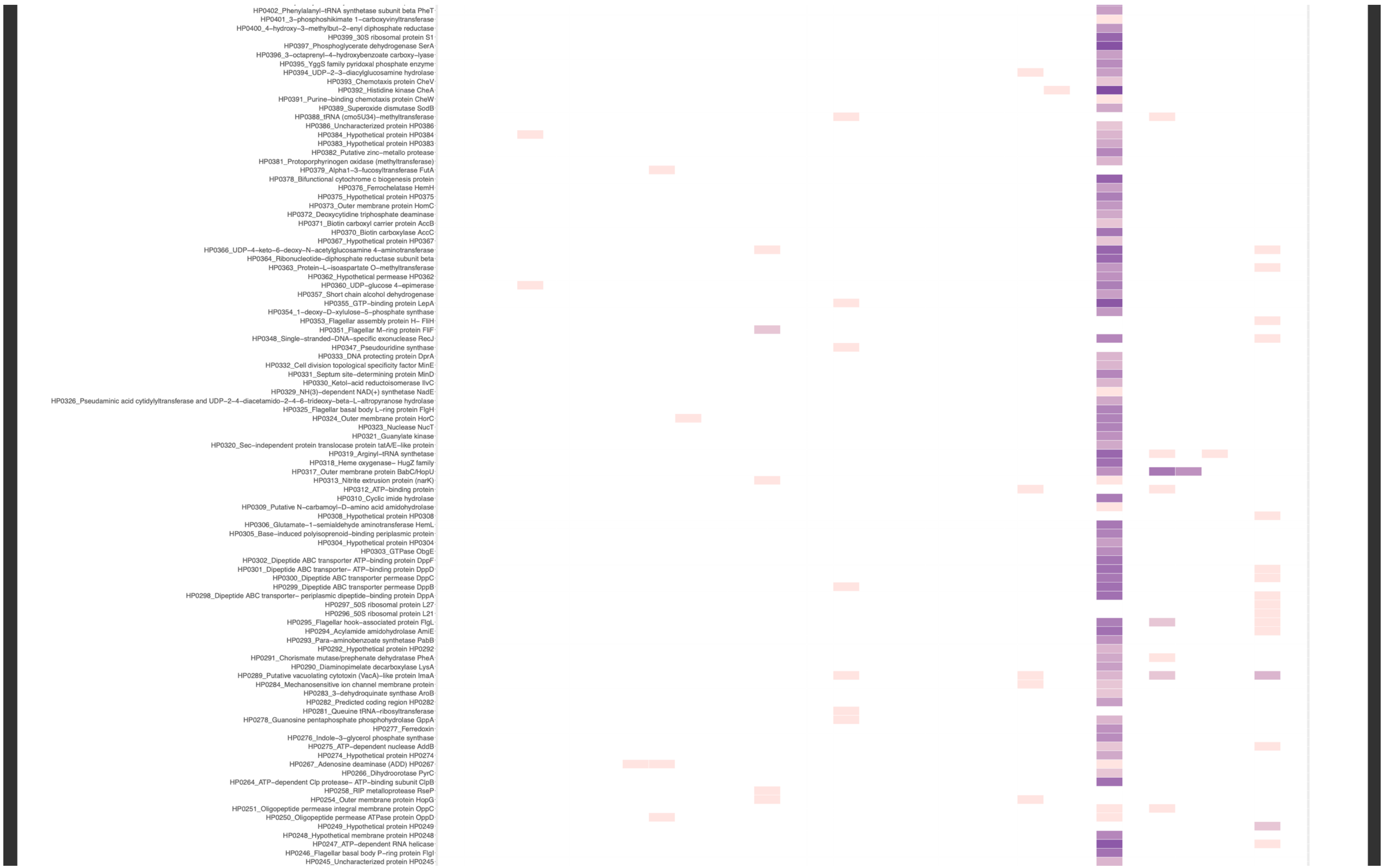

HP0402_Phenylalanyl−tRNA synthetase subunit beta PheT
HP0401_3−phosphoshikimate 1−carboxyvinyltransferase
HP0400_4−hydroxy−3−methylbut−2−enyl diphosphate reductase
HP0399_30S ribosomal protein S1
HP0397_Phosphoglycerate dehydrogenase SerA
HP0396_3−octaprenyl−4−hydroxybenzoate carboxy−lyase
HP0395_YggS family pyridoxal phosphate enzyme
HP0394_UDP−2−3−diacylglucosamine hydrolase
HP0393_Chemotaxis protein CheV
HP0392_Histidine kinase CheA
HP0391_Purine−binding chemotaxis protein CheW
HP0389_Superoxide dismutase SodB
HP0388_tRNA (cmo5U34)−methyltransferase
HP0386_Uncharacterized protein HP0386
HP0384_Hypothetical protein HP0384
HP0383_Hypothetical protein HP0383
HP0382_Putative zinc−metallo protease
HP0381_Protoporphyrinogen oxidase (methyltransferase)
HP0379_Alpha1−3−fucosyltransferase FutA
HP0378_Bifunctional cytochrome c biogenesis protein
HP0376_Ferrochelatase HemH
HP0375_Hypothetical protein HP0375
HP0373_Outer membrane protein HomC
HP0372_Deoxycytidine triphosphate deaminase
HP0371_Biotin carboxyl carrier protein AccB
HP0370_Biotin carboxylase AccC
HP0367_Hypothetical protein HP0367
HP0366_UDP−4−keto−6−deoxy−N−acetylglucosamine 4−aminotransferase
HP0364_Ribonucleotide−diphosphate reductase subunit beta
HP0363_Protein−L−isoaspartate O−methyltransferase
HP0362_Hypothetical permease HP0362
HP0360_UDP−glucose 4−epimerase
HP0357_Short chain alcohol dehydrogenase
HP0355_GTP−binding protein LepA
HP0354_1−deoxy−D−xylulose−5−phosphate synthase
HP0353_Flagellar assembly protein H− FliH
HP0351_Flagellar M−ring protein FliF
HP0348_Single−stranded−DNA−specific exonuclease RecJ
HP0347_Pseudouridine synthase
HP0333_DNA protecting protein DprA
HP0332_Cell division topological specificity factor MinE
HP0331_Septum site−determining protein MinD
HP0330_Ketol−acid reductoisomerase IlvC
HP0329_NH(3)−dependent NAD(+) synthetase NadE
HP0326_Pseudaminic acid cytidylyltransferase and UDP−2−4−diacetamido−2−4−6−trideoxy−beta−L−altropyranose hydrolase
HP0325_Flagellar basal body L−ring protein FlgH
HP0324_Outer membrane protein HorC
HP0323_Nuclease NucT
HP0321_Guanylate kinase
HP0320_Sec−independent protein translocase protein tatA/E−like protein
HP0319_Arginyl−tRNA synthetase
HP0318_Heme oxygenase− HugZ family
HP0317_Outer membrane protein BabC/HopU
HP0313_Nitrite extrusion protein (narK)
HP0312_ATP−binding protein
HP0310_Cyclic imide hydrolase
HP0309_Putative N−carbamoyl−D−amino acid amidohydrolase
HP0308_Hypothetical protein HP0308
HP0306_Glutamate−1−semialdehyde aminotransferase HemL
HP0305_Base−induced polyisoprenoid−binding periplasmic protein
HP0304_Hypothetical protein HP0304
HP0303_GTPase ObgE
HP0302_Dipeptide ABC transporter ATP−binding protein DppF
HP0301_Dipeptide ABC transporter− ATP−binding protein DppD
HP0300_Dipeptide ABC transporter permease DppC
HP0299_Dipeptide ABC transporter permease DppB
HP0298_Dipeptide ABC transporter− periplasmic dipeptide−binding protein DppA
HP0297_50S ribosomal protein L27
HP0296_50S ribosomal protein L21
HP0295_Flagellar hook−associated protein FlgL
HP0294_Acylamide amidohydrolase AmiE
HP0293_Para−aminobenzoate synthetase PabB
HP0292_Hypothetical protein HP0292
HP0291_Chorismate mutase/prephenate dehydratase PheA
HP0290_Diaminopimelate decarboxylase LysA
HP0289_Putative vacuolating cytotoxin (VacA)−like protein ImaA
HP0284_Mechanosensitive ion channel membrane protein
HP0283_3−dehydroquinate synthase AroB
HP0282_Predicted coding region HP0282
HP0281_Queuine tRNA−ribosyltransferase
HP0278_Guanosine pentaphosphate phosphohydrolase GppA
HP0277_Ferredoxin
HP0276_Indole−3−glycerol phosphate synthase
HP0275_ATP−dependent nuclease AddB
HP0274_Hypothetical protein HP0274
HP0267_Adenosine deaminase (ADD) HP0267
HP0266_Dihydroorotase PyrC
HP0264_ATP−dependent Clp protease− ATP−binding subunit ClpB
HP0258_RIP metalloprotease RseP
HP0254_Outer membrane protein HopG
HP0251_Oligopeptide permease integral membrane protein OppC
HP0250_Oligopeptide permease ATPase protein OppD
HP0249_Hypothetical protein HP0249
HP0248_Hypothetical membrane protein HP0248
HP0247_ATP−dependent RNA helicase
HP0246_Flagellar basal body P−ring protein FlgI
HP0245_Uncharacterized protein HP0245

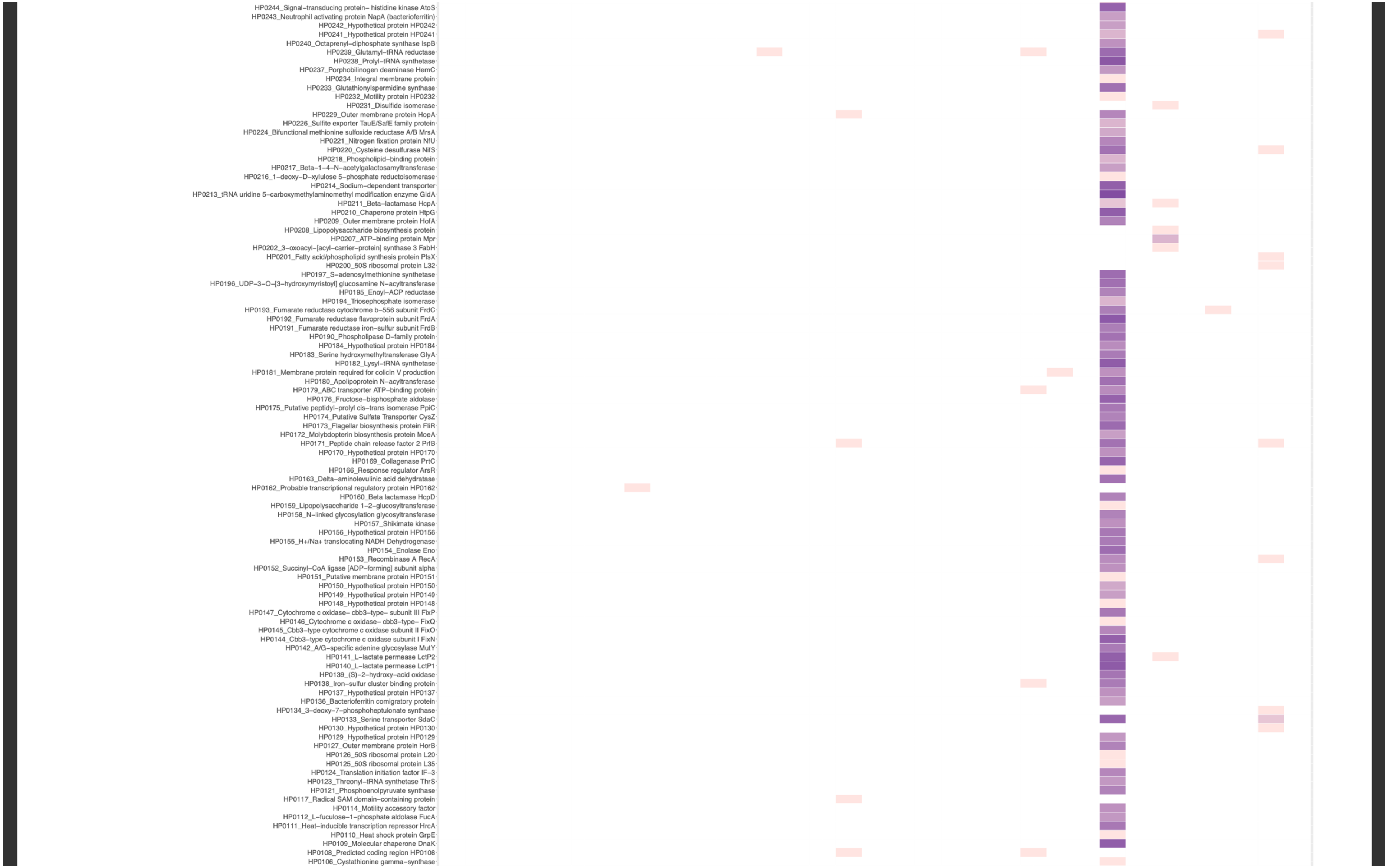
HP0244_Signal−transducing protein− histidine kinase AtoS
HP0243_Neutrophil activating protein NapA (bacterioferritin)
HP0242_Hypothetical protein HP0242
HP0241_Hypothetical protein HP0241
HP0240_Octaprenyl−diphosphate synthase IspB
HP0239_Glutamyl−tRNA reductase
HP0238_Prolyl−tRNA synthetase
HP0237_Porphobilinogen deaminase HemC
HP0234_Integral membrane protein
HP0233_Glutathionylspermidine synthase
HP0232_Motility protein HP0232
HP0231_Disulfide isomerase
HP0229_Outer membrane protein HopA
HP0226_Sulfite exporter TauE/SafE family protein
HP0224_Bifunctional methionine sulfoxide reductase A/B MrsA
HP0221_Nitrogen fixation protein NfU
HP0220_Cysteine desulfurase NifS
HP0218_Phospholipid−binding protein
HP0217_Beta−1−4−N−acetylgalactosamyltransferase
HP0216_1−deoxy−D−xylulose 5−phosphate reductoisomerase
HP0214_Sodium−dependent transporter
HP0213_tRNA uridine 5−carboxymethylaminomethyl modification enzyme GidA
HP0211_Beta−lactamase HcpA
HP0210_Chaperone protein HtpG
HP0209_Outer membrane protein HofA
HP0208_Lipopolysaccharide biosynthesis protein
HP0207_ATP−binding protein Mpr
HP0202_3−oxoacyl−[acyl−carrier−protein] synthase 3 FabH
HP0201_Fatty acid/phospholipid synthesis protein PlsX
HP0200_50S ribosomal protein L32
HP0197_S−adenosylmethionine synthetase
HP0196_UDP−3−O−[3−hydroxymyristoyl] glucosamine N−acyltransferase
HP0195_Enoyl−ACP reductase
HP0194_Triosephosphate isomerase
HP0193_Fumarate reductase cytochrome b−556 subunit FrdC
HP0192_Fumarate reductase flavoprotein subunit FrdA
HP0191_Fumarate reductase iron−sulfur subunit FrdB
HP0190_Phospholipase D−family protein
HP0184_Hypothetical protein HP0184
HP0183_Serine hydroxymethyltransferase GlyA
HP0182_Lysyl−tRNA synthetase
HP0181_Membrane protein required for colicin V production
HP0180_Apolipoprotein N−acyltransferase
HP0179_ABC transporter ATP−binding protein
HP0176_Fructose−bisphosphate aldolase
HP0175_Putative peptidyl−prolyl cis−trans isomerase PpiC
HP0174_Putative Sulfate Transporter CysZ
HP0173_Flagellar biosynthesis protein FliR
HP0172_Molybdopterin biosynthesis protein MoeA
HP0171_Peptide chain release factor 2 PrfB
HP0170_Hypothetical protein HP0170
HP0169_Collagenase PrtC
HP0166_Response regulator ArsR
HP0163_Delta−aminolevulinic acid dehydratase
HP0162_Probable transcriptional regulatory protein HP0162
HP0160_Beta lactamase HcpD
HP0159_Lipopolysaccharide 1−2−glucosyltransferase
HP0158_N−linked glycosylation glycosyltransferase
HP0157_Shikimate kinase
HP0156_Hypothetical protein HP0156
HP0155_H+/Na+ translocating NADH Dehydrogenase
HP0154_Enolase Eno
HP0153_Recombinase A RecA
HP0152_Succinyl−CoA ligase [ADP−forming] subunit alpha
HP0151_Putative membrane protein HP0151
HP0150_Hypothetical protein HP0150
HP0149_Hypothetical protein HP0149
HP0148_Hypothetical protein HP0148
HP0147_Cytochrome c oxidase− cbb3−type− subunit III FixP
HP0146_Cytochrome c oxidase− cbb3−type− FixQ
HP0145_Cbb3−type cytochrome c oxidase subunit II FixO
HP0144_Cbb3−type cytochrome c oxidase subunit I FixN
HP0142_A/G−specific adenine glycosylase MutY
HP0141_L−lactate permease LctP2
HP0140_L−lactate permease LctP1
HP0139_(S)−2−hydroxy−acid oxidase
HP0138_Iron−sulfur cluster binding protein
HP0137_Hypothetical protein HP0137
HP0136_Bacterioferritin comigratory protein
HP0134_3−deoxy−7−phosphoheptulonate synthase
HP0133_Serine transporter SdaC
HP0130_Hypothetical protein HP0130
HP0129_Hypothetical protein HP0129
HP0127_Outer membrane protein HorB
HP0126_50S ribosomal protein L20
HP0125_50S ribosomal protein L35
HP0124_Translation initiation factor IF−3
HP0123_Threonyl−tRNA synthetase ThrS
HP0121_Phosphoenolpyruvate synthase
HP0117_Radical SAM domain−containing protein
HP0114_Motility accessory factor
HP0112_L−fuculose−1−phosphate aldolase FucA
HP0111_Heat−inducible transcription repressor HrcA
HP0110_Heat shock protein GrpE
HP0109_Molecular chaperone DnaK
HP0108_Predicted coding region HP0108
HP0106_Cystathionine gamma−synthase

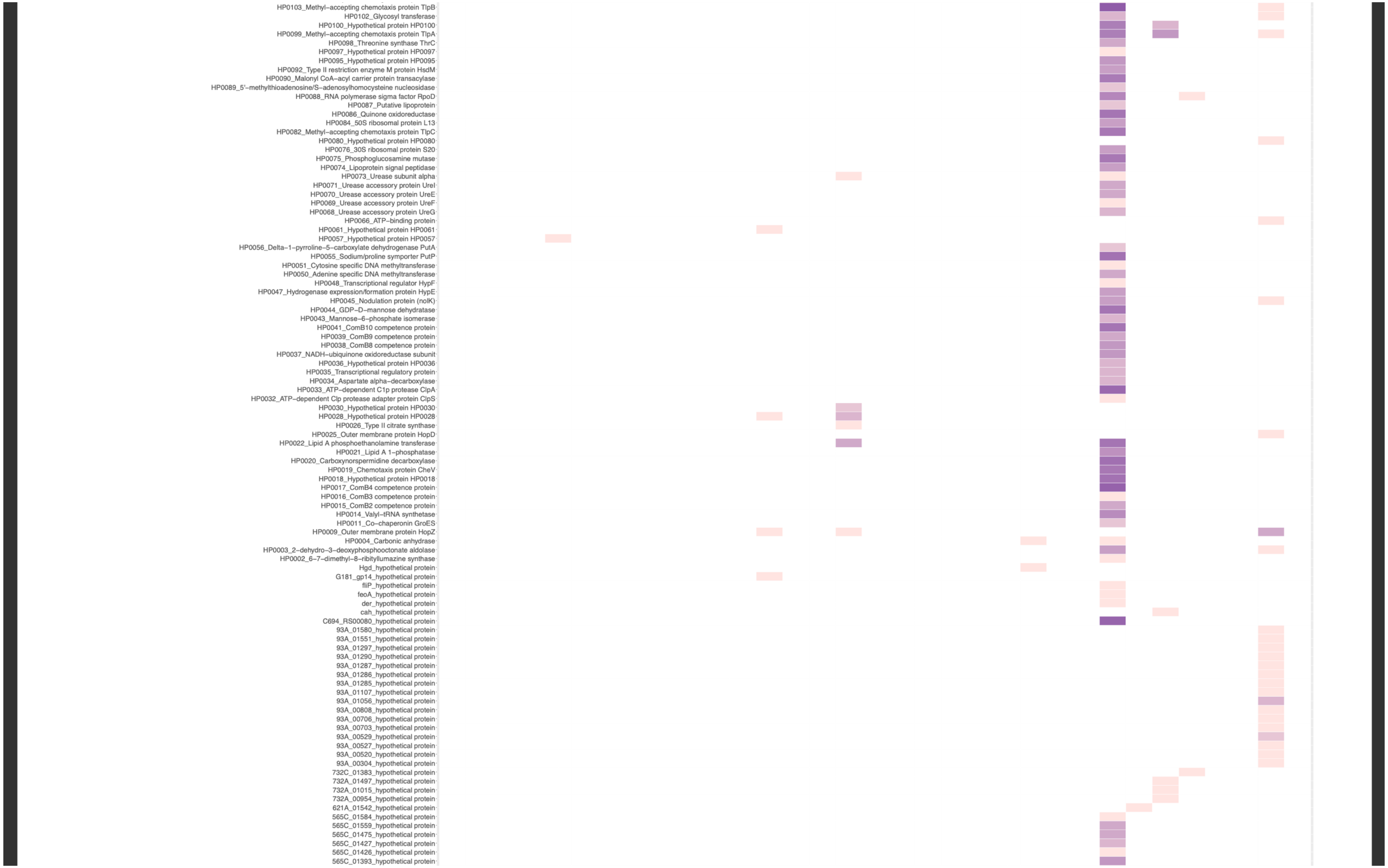
HP0103_Methyl–accepting chemotaxis protein TlpB
HP0102_Glycosyl transferase
HP0100_Hypothetical protein HP0100
HP0099_Methyl–accepting chemotaxis protein TlpA
HP0098_Threonine synthase ThrC
HP0097_Hypothetical protein HP0097
HP0095_Hypothetical protein HP0095
HP0092_Type II restriction enzyme M protein HsdM
HP0090_Malonyl CoA–acyl carrier protein transacylase
HP0089_5'–methylthioadenosine/S–adenosylhomocysteine nucleosidase
HP0088_RNA polymerase sigma factor RpoD
HP0087_Putative lipoprotein
HP0086_Quinone oxidoreductase
HP0084_50S ribosomal protein L13
HP0082_Methyl–accepting chemotaxis protein TlpC
HP0080_Hypothetical protein HP0080
HP0076_30S ribosomal protein S20
HP0075_Phosphoglucosamine mutase
HP0074_Lipoprotein signal peptidase
HP0073_Urease subunit alpha
HP0071_Urease accessory protein UreI
HP0070_Urease accessory protein UreE
HP0069_Urease accessory protein UreF
HP0068_Urease accessory protein UreG
HP0066_ATP–binding protein
HP0061_Hypothetical protein HP0061
HP0057_Hypothetical protein HP0057
HP0056_Delta–1–pyrroline–5–carboxylate dehydrogenase PutA
HP0055_Sodium/proline symporter PutP
HP0051_Cytosine specific DNA methyltransferase
HP0050_Adenine specific DNA methyltransferase
HP0048_Transcriptional regulator HypF
HP0047_Hydrogenase expression/formation protein HypE
HP0045_Nodulation protein (nolK)
HP0044_GDP–D–mannose dehydratase
HP0043_Mannose–6–phosphate isomerase
HP0041_ComB10 competence protein
HP0039_ComB9 competence protein
HP0038_ComB8 competence protein
HP0037_NADH–ubiquinone oxidoreductase subunit
HP0036_Hypothetical protein HP0036
HP0035_Transcriptional regulatory protein
HP0034_Aspartate alpha–decarboxylase
HP0033_ATP–dependent C1p protease ClpA
HP0032_ATP–dependent Clp protease adapter protein ClpS
HP0030_Hypothetical protein HP0030
HP0028_Hypothetical protein HP0028
HP0026_Type II citrate synthase
HP0025_Outer membrane protein HopD
HP0022_Lipid A phosphoethanolamine transferase
HP0021_Lipid A 1–phosphatase
HP0020_Carboxynorspermidine decarboxylase
HP0019_Chemotaxis protein CheV
HP0018_Hypothetical protein HP0018
HP0017_ComB4 competence protein
HP0016_ComB3 competence protein
HP0015_ComB2 competence protein
HP0014_Valyl–tRNA synthetase
HP0011_Co–chaperonin GroES
HP0009_Outer membrane protein HopZ
HP0004_Carbonic anhydrase
HP0003_2–dehydro–3–deoxyphosphooctonate aldolase
HP0002_6–7–dimethyl–8–ribityllumazine synthase
Hgd_hypothetical protein
G181_gp14_hypothetical protein
fliP_hypothetical protein
feoA_hypothetical protein
der_hypothetical protein
cah_hypothetical protein
C694_RS00080_hypothetical protein
93A_01580_hypothetical protein
93A_01551_hypothetical protein
93A_01297_hypothetical protein
93A_01290_hypothetical protein
93A_01287_hypothetical protein
93A_01286_hypothetical protein
93A_01285_hypothetical protein
93A_01107_hypothetical protein
93A_01056_hypothetical protein
93A_00808_hypothetical protein
93A_00706_hypothetical protein
93A_00703_hypothetical protein
93A_00529_hypothetical protein
93A_00527_hypothetical protein
93A_00520_hypothetical protein
93A_00304_hypothetical protein
732C_01383_hypothetical protein
732A_01497_hypothetical protein
732A_01015_hypothetical protein
732A_00954_hypothetical protein
621A_01542_hypothetical protein
565C_01584_hypothetical protein
565C_01559_hypothetical protein
565C_01475_hypothetical protein
565C_01427_hypothetical protein
565C_01426_hypothetical protein
565C_01393_hypothetical protein

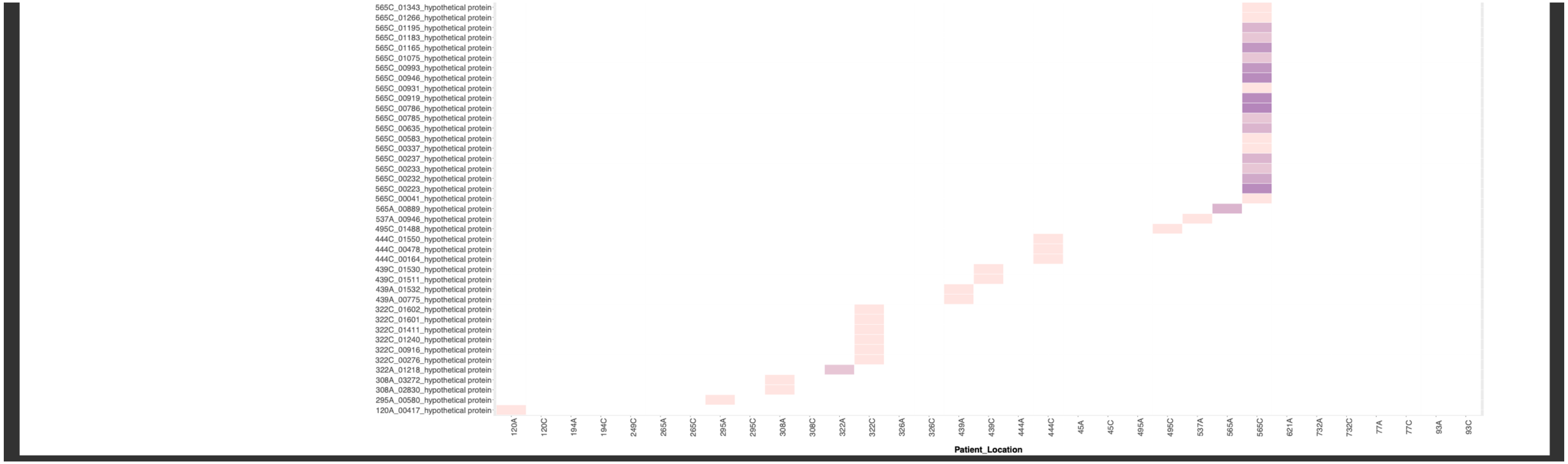


**Suppl. Fig. 17.** Heat maps showing all common allelic variation across genes/gene products for each antrum- and corpus-derived *H. pylori* population, derived from deep sequencing of the population from each biopsy and read-mapping back to the consensus genome to identify variant bases at each locus. Higher colour intensity indicates a larger number of variant bases within that gene/gene product.

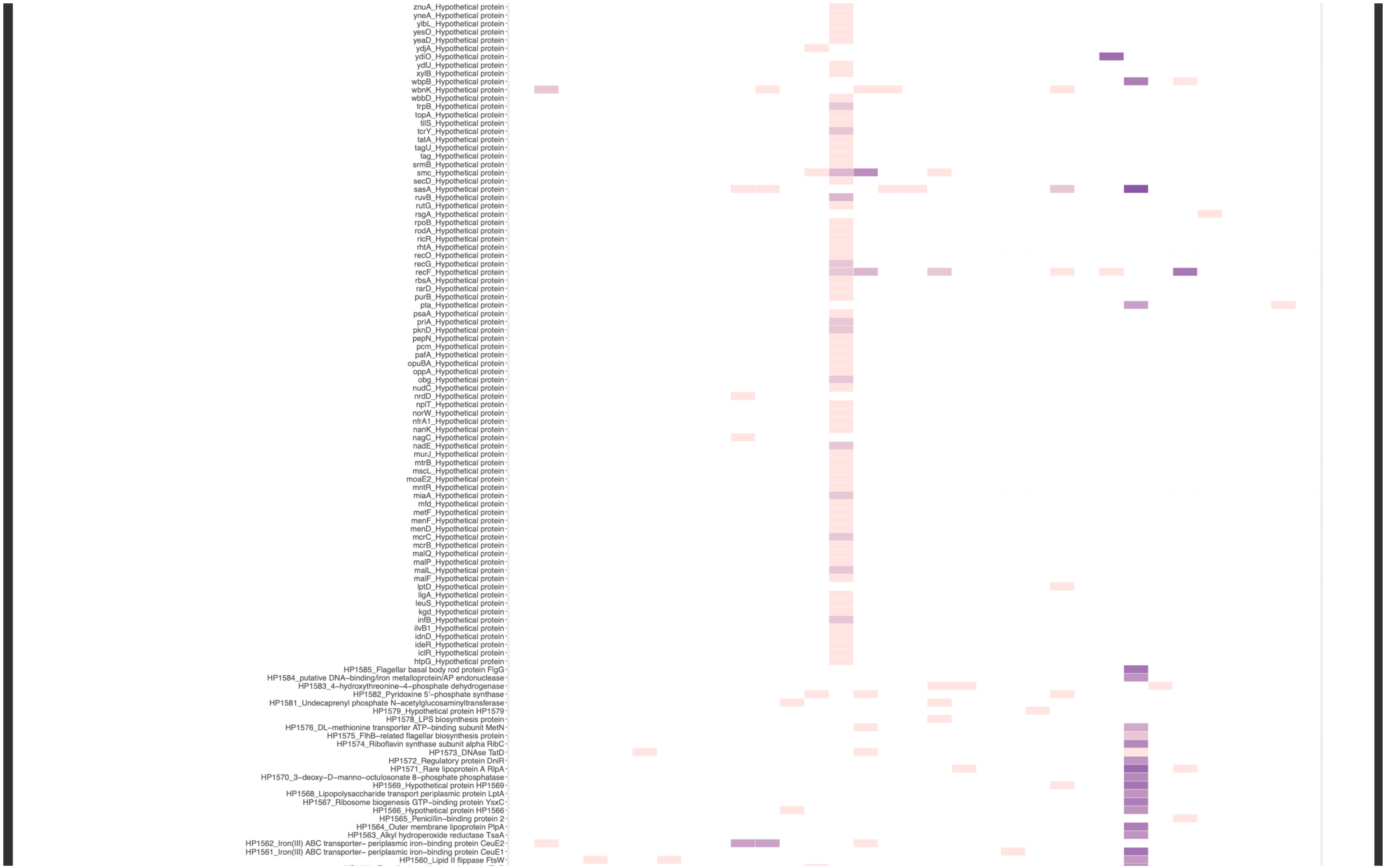
znuA_Hypothetical protein
yneA_Hypothetical protein
ylbL_Hypothetical protein
yesO_Hypothetical protein
yeaD_Hypothetical protein
ydjA_Hypothetical protein
ydiO_Hypothetical protein
ydfJ_Hypothetical protein
xylB_Hypothetical protein
wbpB_Hypothetical protein
wbnK_Hypothetical protein
wbbD_Hypothetical protein
trpB_Hypothetical protein
topA_Hypothetical protein
tilS_Hypothetical protein
tcrY_Hypothetical protein
tatA_Hypothetical protein
tagU_Hypothetical protein
tag_Hypothetical protein
srmB_Hypothetical protein
smc_Hypothetical protein
secD_Hypothetical protein
sasA_Hypothetical protein
ruvB_Hypothetical protein
rutG_Hypothetical protein
rsgA_Hypothetical protein
rpoB_Hypothetical protein
rodA_Hypothetical protein
ricR_Hypothetical protein
rhtA_Hypothetical protein
recO_Hypothetical protein
recG_Hypothetical protein
recF_Hypothetical protein
rbsA_Hypothetical protein
rarD_Hypothetical protein
purB_Hypothetical protein
pta_Hypothetical protein
psaA_Hypothetical protein
priA_Hypothetical protein
pknD_Hypothetical protein
pepN_Hypothetical protein
pcm_Hypothetical protein
pafA_Hypothetical protein
opuBA_Hypothetical protein
oppA_Hypothetical protein
obg_Hypothetical protein
nudC_Hypothetical protein
nrdD_Hypothetical protein
nplT_Hypothetical protein
norW_Hypothetical protein
nfrA1_Hypothetical protein
nanK_Hypothetical protein
nagC_Hypothetical protein
nadE_Hypothetical protein
murJ_Hypothetical protein
mtrB_Hypothetical protein
mscL_Hypothetical protein
moaE2_Hypothetical protein
mntR_Hypothetical protein
miaA_Hypothetical protein
mfd_Hypothetical protein
metF_Hypothetical protein
menF_Hypothetical protein
menD_Hypothetical protein
mcrC_Hypothetical protein
mcrB_Hypothetical protein
malQ_Hypothetical protein
malP_Hypothetical protein
malL_Hypothetical protein
malF_Hypothetical protein
lptD_Hypothetical protein
ligA_Hypothetical protein
leuS_Hypothetical protein
kgd_Hypothetical protein
infB_Hypothetical protein
ilvB1_Hypothetical protein
idnD_Hypothetical protein
ideR_Hypothetical protein
iclR_Hypothetical protein
htpG_Hypothetical protein
HP1585_Flagellar basal body rod protein FlgG
HP1584_putative DNA–binding/iron metalloprotein/AP endonuclease
HP1583_4–hydroxythreonine–4–phosphate dehydrogenase
HP1582_Pyridoxine 5'–phosphate synthase
HP1581_Undecaprenyl phosphate N–acetylglucosaminyltransferase
HP1579_Hypothetical protein HP1579
HP1578_LPS biosynthesis protein
HP1576_DL–methionine transporter ATP–binding subunit MetN
HP1575_FlhB–related flagellar biosynthesis protein
HP1574_Riboflavin synthase subunit alpha RibC
HP1573_DNAse TatD
HP1572_Regulatory protein DniR
HP1571_Rare lipoprotein A RlpA
HP1570_3–deoxy–D–manno–octulosonate 8–phosphate phosphatase
HP1569_Hypothetical protein HP1569
HP1568_Lipopolysaccharide transport periplasmic protein LptA
HP1567_Ribosome biogenesis GTP–binding protein YsxC
HP1566_Hypothetical protein HP1566
HP1565_Penicillin–binding protein 2
HP1564_Outer membrane lipoprotein PlpA
HP1563_Alkyl hydroperoxide reductase TsaA
HP1562_Iron(III) ABC transporter– periplasmic iron–binding protein CeuE2
HP1561_Iron(III) ABC transporter– periplasmic iron–binding protein CeuE1
HP1560_Lipid II flippase FtsW

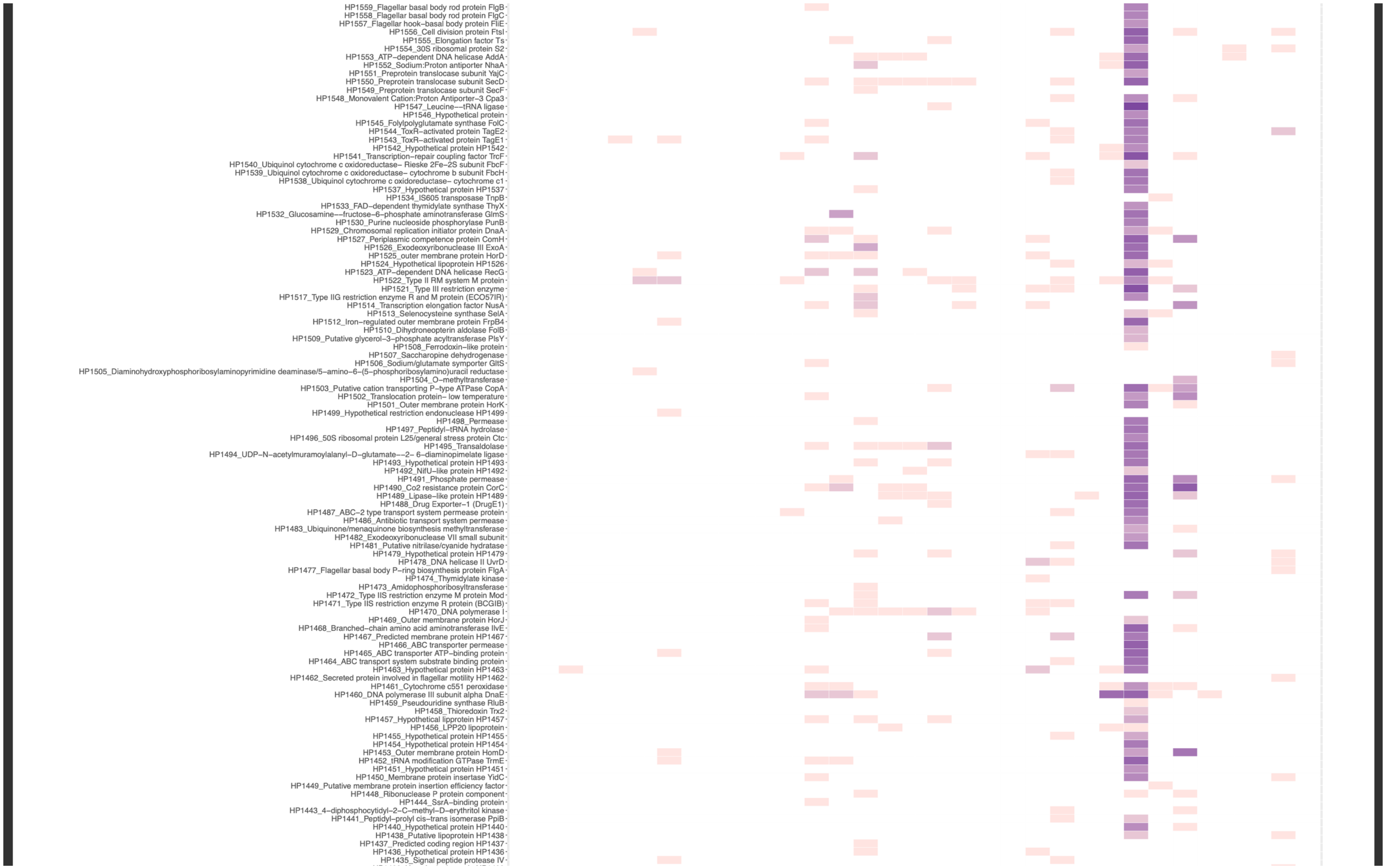
HP1559_Flagellar basal body rod protein FlgB
HP1558_Flagellar basal body rod protein FlgC
HP1557_Flagellar hook–basal body protein FliE
HP1556_Cell division protein FtsI
HP1555_Elongation factor Ts
HP1554_30S ribosomal protein S2
HP1553_ATP–dependent DNA helicase AddA
HP1552_Sodium:Proton antiporter NhaA
HP1551_Preprotein translocase subunit YajC
HP1550_Preprotein translocase subunit SecD
HP1549_Preprotein translocase subunit SecF
HP1548_Monovalent Cation:Proton Antiporter–3 Cpa3
HP1547_Leucine––tRNA ligase
HP1546_Hypothetical protein
HP1545_Folylpolyglutamate synthase FolC
HP1544_ToxR–activated protein TagE2
HP1543_ToxR–activated protein TagE1
HP1542_Hypothetical protein HP1542
HP1541_Transcription–repair coupling factor TrcF
HP1540_Ubiquinol cytochrome c oxidoreductase– Rieske 2Fe–2S subunit FbcF
HP1539_Ubiquinol cytochrome c oxidoreductase– cytochrome b subunit FbcH
HP1538_Ubiquinol cytochrome c oxidoreductase– cytochrome c1
HP1537_Hypothetical protein HP1537
HP1534_IS605 transposase TnpB
HP1533_FAD–dependent thymidylate synthase ThyX
HP1532_Glucosamine––fructose–6–phosphate aminotransferase GlmS
HP1530_Purine nucleoside phosphorylase PunB
HP1529_Chromosomal replication initiator protein DnaA
HP1527_Periplasmic competence protein ComH
HP1526_Exodeoxyribonuclease III ExoA
HP1525_outer membrane protein HorD
HP1524_Hypothetical lipoprotein HP1526
HP1523_ATP–dependent DNA helicase RecG
HP1522_Type II RM system M protein
HP1521_Type III restriction enzyme
HP1517_Type IIG restriction enzyme R and M protein (ECO57IR)
HP1514_Transcription elongation factor NusA
HP1513_Selenocysteine synthase SelA
HP1512_Iron–regulated outer membrane protein FrpB4
HP1510_Dihydroneopterin aldolase FolB
HP1509_Putative glycerol–3–phosphate acyltransferase PlsY
HP1508_Ferrodoxin–like protein
HP1507_Saccharopine dehydrogenase
HP1506_Sodium/glutamate symporter GltS
HP1505_Diaminohydroxyphosphoribosylaminopyrimidine deaminase/5–amino–6–(5–phosphoribosylamino)uracil reductase
HP1504_O–methyltransferase
HP1503_Putative cation transporting P–type ATPase CopA
HP1502_Translocation protein– low temperature
HP1501_Outer membrane protein HorK
HP1499_Hypothetical restriction endonuclease HP1499
HP1498_Permease
HP1497_Peptidyl–tRNA hydrolase
HP1496_50S ribosomal protein L25/general stress protein Ctc
HP1495_Transaldolase
HP1494_UDP–N–acetylmuramoylalanyl–D–glutamate––2– 6–diaminopimelate ligase
HP1493_Hypothetical protein HP1493
HP1492_NifU–like protein HP1492
HP1491_Phosphate permease
HP1490_Co2 resistance protein CorC
HP1489_Lipase–like protein HP1489
HP1488_Drug Exporter–1 (DrugE1)
HP1487_ABC–2 type transport system permease protein
HP1486_Antibiotic transport system permease
HP1483_Ubiquinone/menaquinone biosynthesis methyltransferase
HP1482_Exodeoxyribonuclease VII small subunit
HP1481_Putative nitrilase/cyanide hydratase
HP1479_Hypothetical protein HP1479
HP1478_DNA helicase II UvrD
HP1477_Flagellar basal body P–ring biosynthesis protein FlgA
HP1474_Thymidylate kinase
HP1473_Amidophosphoribosyltransferase
HP1472_Type IIS restriction enzyme M protein Mod
HP1471_Type IIS restriction enzyme R protein (BCGIB)
HP1470_DNA polymerase I
HP1469_Outer membrane protein HorJ
HP1468_Branched–chain amino acid aminotransferase IlvE
HP1467_Predicted membrane protein HP1467
HP1466_ABC transporter permease
HP1465_ABC transporter ATP–binding protein
HP1464_ABC transport system substrate binding protein
HP1463_Hypothetical protein HP1463
HP1462_Secreted protein involved in flagellar motility HP1462
HP1461_Cytochrome c551 peroxidase
HP1460_DNA polymerase III subunit alpha DnaE
HP1459_Pseudouridine synthase RluB
HP1458_Thioredoxin Trx2
HP1457_Hypothetical lipprotein HP1457
HP1456_LPP20 lipoprotein
HP1455_Hypothetical protein HP1455
HP1454_Hypothetical protein HP1454
HP1453_Outer membrane protein HomD
HP1452_tRNA modification GTPase TrmE
HP1451_Hypothetical protein HP1451
HP1450_Membrane protein insertase YidC
HP1449_Putative membrane protein insertion efficiency factor
HP1448_Ribonuclease P protein component
HP1444_SsrA–binding protein
HP1443_4–diphosphocytidyl–2–C–methyl–D–erythritol kinase
HP1441_Peptidyl–prolyl cis–trans isomerase PpiB
HP1440_Hypothetical protein HP1440
HP1438_Putative lipoprotein HP1438
HP1437_Predicted coding region HP1437
HP1436_Hypothetical protein HP1436
HP1435_Signal peptide protease IV

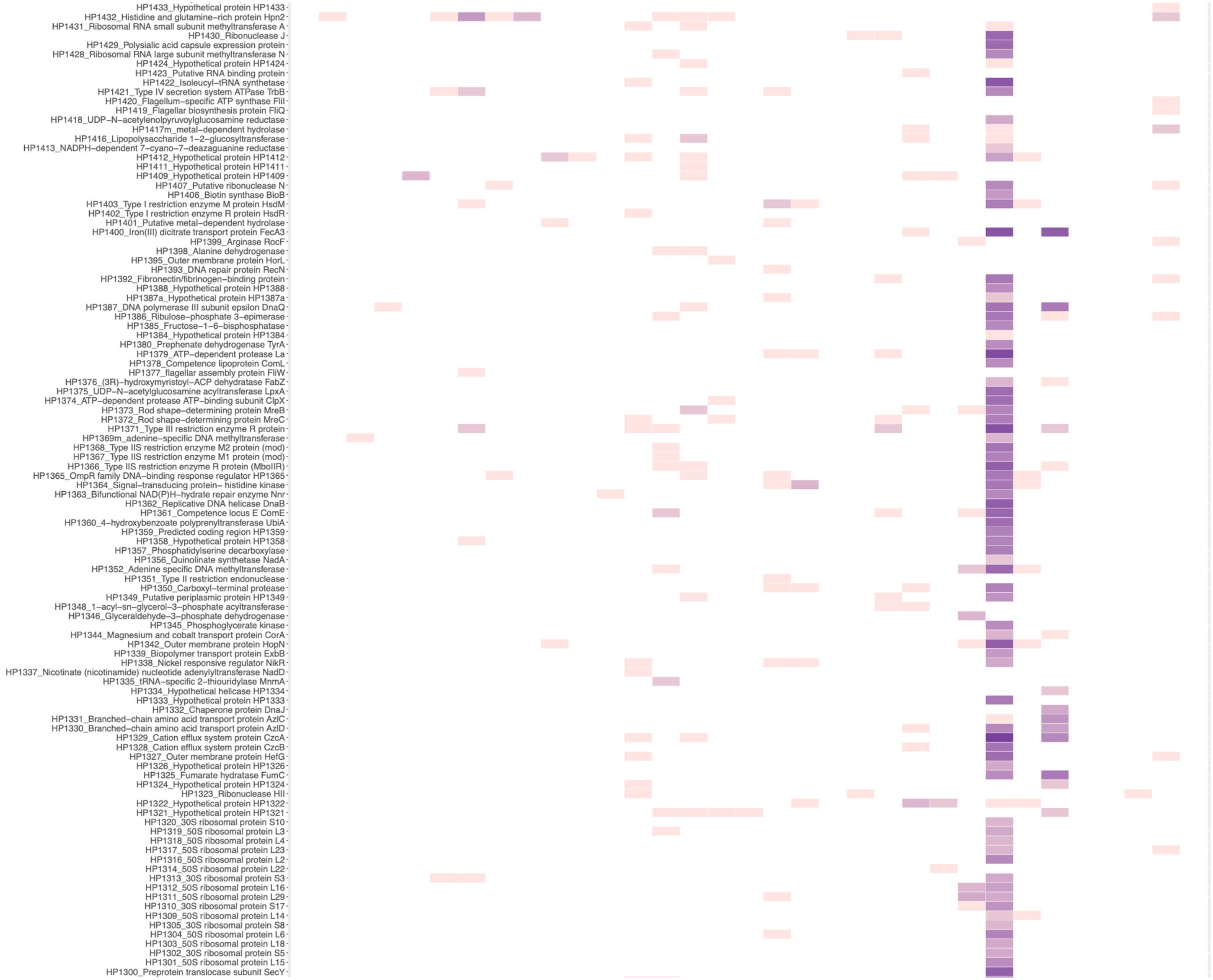

HP1433_Hypothetical protein HP1433
HP1432_Histidine and glutamine–rich protein Hpn2
HP1431_Ribosomal RNA small subunit methyltransferase A
HP1430_Ribonuclease J
HP1429_Polysialic acid capsule expression protein
HP1428_Ribosomal RNA large subunit methyltransferase N
HP1424_Hypothetical protein HP1424
HP1423_Putative RNA binding protein
HP1422_Isoleucyl–tRNA synthetase
HP1421_Type IV secretion system ATPase TrbB
HP1420_Flagellum–specific ATP synthase FliI
HP1419_Flagellar biosynthesis protein FliQ
HP1418_UDP–N–acetylenolpyruvoylglucosamine reductase
HP1417m_metal–dependent hydrolase
HP1416_Lipopolysaccharide 1–2–glucosyltransferase
HP1413_NADPH–dependent 7–cyano–7–deazaguanine reductase
HP1412_Hypothetical protein HP1412
HP1411_Hypothetical protein HP1411
HP1409_Hypothetical protein HP1409
HP1407_Putative ribonuclease N
HP1406_Biotin synthase BioB
HP1403_Type I restriction enzyme M protein HsdM
HP1402_Type I restriction enzyme R protein HsdR
HP1401_Putative metal–dependent hydrolase
HP1400_Iron(III) dicitrate transport protein FecA3
HP1399_Arginase RocF
HP1398_Alanine dehydrogenase
HP1395_Outer membrane protein HorL
HP1393_DNA repair protein RecN
HP1392_Fibronectin/fibrinogen–binding protein
HP1388_Hypothetical protein HP1388
HP1387a_Hypothetical protein HP1387a
HP1387_DNA polymerase III subunit epsilon DnaQ
HP1386_Ribulose–phosphate 3–epimerase
HP1385_Fructose–1–6–bisphosphatase
HP1384_Hypothetical protein HP1384
HP1380_Prephenate dehydrogenase TyrA
HP1379_ATP–dependent protease La
HP1378_Competence lipoprotein ComL
HP1377_flagellar assembly protein FliW
HP1376_(3R)–hydroxymyristoyl–ACP dehydratase FabZ
HP1375_UDP–N–acetylglucosamine acyltransferase LpxA
HP1374_ATP–dependent protease ATP–binding subunit ClpX
HP1373_Rod shape–determining protein MreB
HP1372_Rod shape–determining protein MreC
HP1371_Type III restriction enzyme R protein
HP1369m_adenine–specific DNA methyltransferase
HP1368_Type IIS restriction enzyme M2 protein (mod)
HP1367_Type IIS restriction enzyme M1 protein (mod)
HP1366_Type IIS restriction enzyme R protein (MboIIR)
HP1365_OmpR family DNA–binding response regulator HP1365
HP1364_Signal–transducing protein– histidine kinase
HP1363_Bifunctional NAD(P)H–hydrate repair enzyme Nnr
HP1362_Replicative DNA helicase DnaB
HP1361_Competence locus E ComE
HP1360_4–hydroxybenzoate polyprenyltransferase UbiA
HP1359_Predicted coding region HP1359
HP1358_Hypothetical protein HP1358
HP1357_Phosphatidylserine decarboxylase
HP1356_Quinolinate synthetase NadA
HP1352_Adenine specific DNA methyltransferase
HP1351_Type II restriction endonuclease
HP1350_Carboxyl–terminal protease
HP1349_Putative periplasmic protein HP1349
HP1348_1–acyl–sn–glycerol–3–phosphate acyltransferase
HP1346_Glyceraldehyde–3–phosphate dehydrogenase
HP1345_Phosphoglycerate kinase
HP1344_Magnesium and cobalt transport protein CorA
HP1342_Outer membrane protein HopN
HP1339_Biopolymer transport protein ExbB
HP1338_Nickel responsive regulator NikR
HP1337_Nicotinate (nicotinamide) nucleotide adenylyltransferase NadD
HP1335_tRNA–specific 2–thiouridylase MnmA
HP1334_Hypothetical helicase HP1334
HP1333_Hypothetical protein HP1333
HP1332_Chaperone protein DnaJ
HP1331_Branched–chain amino acid transport protein AzlC
HP1330_Branched–chain amino acid transport protein AzlD
HP1329_Cation efflux system protein CzcA
HP1328_Cation efflux system protein CzcB
HP1327_Outer membrane protein HefG
HP1326_Hypothetical protein HP1326
HP1325_Fumarate hydratase FumC
HP1324_Hypothetical protein HP1324
HP1323_Ribonuclease HII
HP1322_Hypothetical protein HP1322
HP1321_Hypothetical protein HP1321
HP1320_30S ribosomal protein S10
HP1319_50S ribosomal protein L3
HP1318_50S ribosomal protein L4
HP1317_50S ribosomal protein L23
HP1316_50S ribosomal protein L2
HP1314_50S ribosomal protein L22
HP1313_30S ribosomal protein S3
HP1312_50S ribosomal protein L16
HP1311_50S ribosomal protein L29
HP1310_30S ribosomal protein S17
HP1309_50S ribosomal protein L14
HP1305_30S ribosomal protein S8
HP1304_50S ribosomal protein L6
HP1303_50S ribosomal protein L18
HP1302_30S ribosomal protein S5
HP1301_50S ribosomal protein L15
HP1300_Preprotein translocase subunit SecY

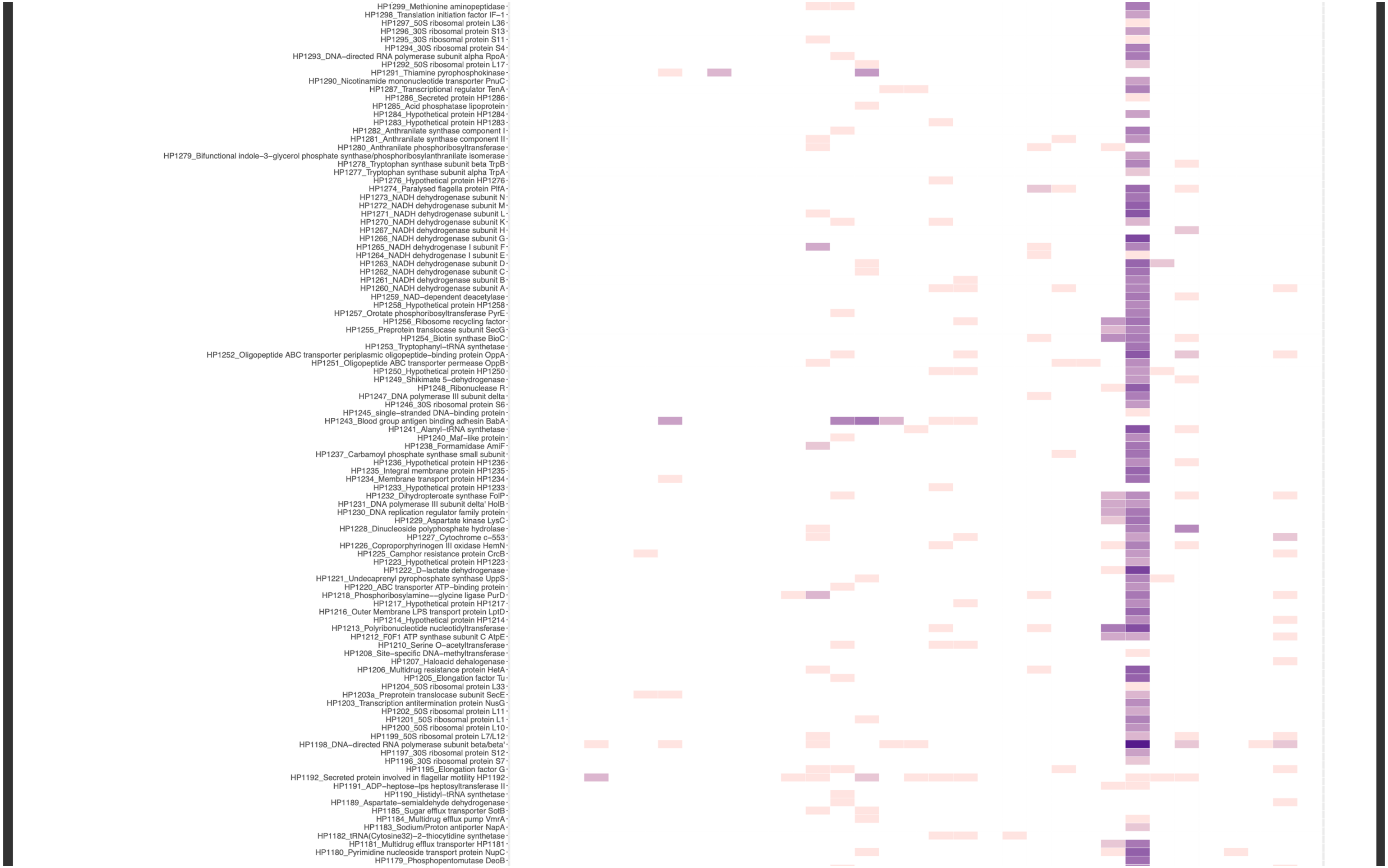
HP1299_Methionine aminopeptidase
HP1298_Translation initiation factor IF−1
HP1297_50S ribosomal protein L36
HP1296_30S ribosomal protein S13
HP1295_30S ribosomal protein S11
HP1294_30S ribosomal protein S4
HP1293_DNA−directed RNA polymerase subunit alpha RpoA
HP1292_50S ribosomal protein L17
HP1291_Thiamine pyrophosphokinase
HP1290_Nicotinamide mononucleotide transporter PnuC
HP1287_Transcriptional regulator TenA
HP1286_Secreted protein HP1286
HP1285_Acid phosphatase lipoprotein
HP1284_Hypothetical protein HP1284
HP1283_Hypothetical protein HP1283
HP1282_Anthranilate synthase component I
HP1281_Anthranilate synthase component II
HP1280_Anthranilate phosphoribosyltransferase
HP1279_Bifunctional indole−3−glycerol phosphate synthase/phosphoribosylanthranilate isomerase
HP1278_Tryptophan synthase subunit beta TrpB
HP1277_Tryptophan synthase subunit alpha TrpA
HP1276_Hypothetical protein HP1276
HP1274_Paralysed flagella protein PflA
HP1273_NADH dehydrogenase subunit N
HP1272_NADH dehydrogenase subunit M
HP1271_NADH dehydrogenase subunit L
HP1270_NADH dehydrogenase subunit K
HP1267_NADH dehydrogenase subunit H
HP1266_NADH dehydrogenase subunit G
HP1265_NADH dehydrogenase I subunit F
HP1264_NADH dehydrogenase I subunit E
HP1263_NADH dehydrogenase subunit D
HP1262_NADH dehydrogenase subunit C
HP1261_NADH dehydrogenase subunit B
HP1260_NADH dehydrogenase subunit A
HP1259_NAD−dependent deacetylase
HP1258_Hypothetical protein HP1258
HP1257_Orotate phosphoribosyltransferase PyrE
HP1256_Ribosome recycling factor
HP1255_Preprotein translocase subunit SecG
HP1254_Biotin synthase BioC
HP1253_Tryptophanyl−tRNA synthetase
HP1252_Oligopeptide ABC transporter periplasmic oligopeptide−binding protein OppA
HP1251_Oligopeptide ABC transporter permease OppB
HP1250_Hypothetical protein HP1250
HP1249_Shikimate 5−dehydrogenase
HP1248_Ribonuclease R
HP1247_DNA polymerase III subunit delta
HP1246_30S ribosomal protein S6
HP1245_single−stranded DNA−binding protein
HP1243_Blood group antigen binding adhesin BabA
HP1241_Alanyl−tRNA synthetase
HP1240_Maf−like protein
HP1238_Formamidase AmiF
HP1237_Carbamoyl phosphate synthase small subunit
HP1236_Hypothetical protein HP1236
HP1235_Integral membrane protein HP1235
HP1234_Membrane transport protein HP1234
HP1233_Hypothetical protein HP1233
HP1232_Dihydropteroate synthase FolP
HP1231_DNA polymerase III subunit delta' HolB
HP1230_DNA replication regulator family protein
HP1229_Aspartate kinase LysC
HP1228_Dinucleoside polyphosphate hydrolase
HP1227_Cytochrome c−553
HP1226_Coproporphyrinogen III oxidase HemN
HP1225_Camphor resistance protein CrcB
HP1223_Hypothetical protein HP1223
HP1222_D−lactate dehydrogenase
HP1221_Undecaprenyl pyrophosphate synthase UppS
HP1220_ABC transporter ATP−binding protein
HP1218_Phosphoribosylamine−−glycine ligase PurD
HP1217_Hypothetical protein HP1217
HP1216_Outer Membrane LPS transport protein LptD
HP1214_Hypothetical protein HP1214
HP1213_Polyribonucleotide nucleotidyltransferase
HP1212_F0F1 ATP synthase subunit C AtpE
HP1210_Serine O−acetyltransferase
HP1208_Site−specific DNA−methyltransferase
HP1207_Haloacid dehalogenase
HP1206_Multidrug resistance protein HetA
HP1205_Elongation factor Tu
HP1204_50S ribosomal protein L33
HP1203a_Preprotein translocase subunit SecE
HP1203_Transcription antitermination protein NusG
HP1202_50S ribosomal protein L11
HP1201_50S ribosomal protein L1
HP1200_50S ribosomal protein L10
HP1199_50S ribosomal protein L7/L12
HP1198_DNA−directed RNA polymerase subunit beta/beta'
HP1197_30S ribosomal protein S12
HP1196_30S ribosomal protein S7
HP1195_Elongation factor G
HP1192_Secreted protein involved in flagellar motility HP1192
HP1191_ADP−heptose−lps heptosyltransferase II
HP1190_Histidyl−tRNA synthetase
HP1189_Aspartate−semialdehyde dehydrogenase
HP1185_Sugar efflux transporter SotB
HP1184_Multidrug efflux pump VmrA
HP1183_Sodium/Proton antiporter NapA
HP1182_tRNA(Cytosine32)−2−thiocytidine synthetase
HP1181_Multidrug efflux transporter HP1181
HP1180_Pyrimidine nucleoside transport protein NupC
HP1179_Phosphopentomutase DeoB

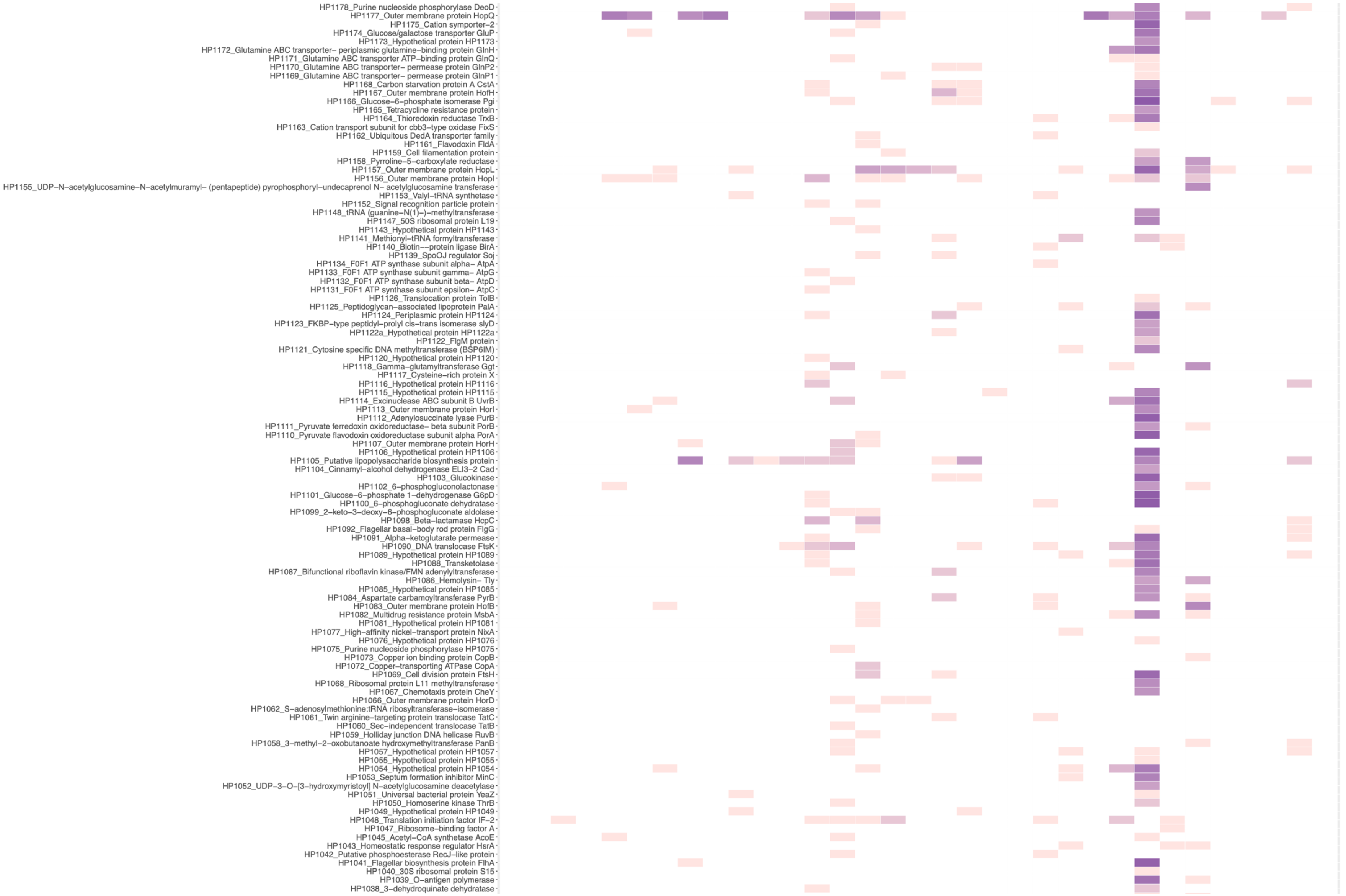
HP1178_Purine nucleoside phosphorylase DeoD
HP1177_Outer membrane protein HopQ
HP1175_Cation symporter–2
HP1174_Glucose/galactose transporter GluP
HP1173_Hypothetical protein HP1173
HP1172_Glutamine ABC transporter– periplasmic glutamine–binding protein GlnH
HP1171_Glutamine ABC transporter ATP–binding protein GlnQ
HP1170_Glutamine ABC transporter– permease protein GlnP2
HP1169_Glutamine ABC transporter– permease protein GlnP1
HP1168_Carbon starvation protein A CstA
HP1167_Outer membrane protein HofH
HP1166_Glucose–6–phosphate isomerase Pgi
HP1165_Tetracycline resistance protein
HP1164_Thioredoxin reductase TrxB
HP1163_Cation transport subunit for cbb3–type oxidase FixS
HP1162_Ubiquitous DedA transporter family
HP1161_Flavodoxin FldA
HP1159_Cell filamentation protein
HP1158_Pyrroline–5–carboxylate reductase
HP1157_Outer membrane protein HopL
HP1156_Outer membrane protein HopI
HP1155_UDP–N–acetylglucosamine–N–acetylmuramyl– (pentapeptide) pyrophosphoryl–undecaprenol N– acetylglucosamine transferase
HP1153_Valyl–tRNA synthetase
HP1152_Signal recognition particle protein
HP1148_tRNA (guanine–N(1)–)–methyltransferase
HP1147_50S ribosomal protein L19
HP1143_Hypothetical protein HP1143
HP1141_Methionyl–tRNA formyltransferase
HP1140_Biotin––protein ligase BirA
HP1139_SpoOJ regulator Soj
HP1134_F0F1 ATP synthase subunit alpha– AtpA
HP1133_F0F1 ATP synthase subunit gamma– AtpG
HP1132_F0F1 ATP synthase subunit beta– AtpD
HP1131_F0F1 ATP synthase subunit epsilon– AtpC
HP1126_Translocation protein TolB
HP1125_Peptidoglycan–associated lipoprotein PalA
HP1124_Periplasmic protein HP1124
HP1123_FKBP–type peptidyl–prolyl cis–trans isomerase slyD
HP1122a_Hypothetical protein HP1122a
HP1122_FlgM protein
HP1121_Cytosine specific DNA methyltransferase (BSP6IM)
HP1120_Hypothetical protein HP1120
HP1118_Gamma–glutamyltransferase Ggt
HP1117_Cysteine–rich protein X
HP1116_Hypothetical protein HP1116
HP1115_Hypothetical protein HP1115
HP1114_Excinuclease ABC subunit B UvrB
HP1113_Outer membrane protein HorI
HP1112_Adenylosuccinate lyase PurB
HP1111_Pyruvate ferredoxin oxidoreductase– beta subunit PorB
HP1110_Pyruvate flavodoxin oxidoreductase subunit alpha PorA
HP1107_Outer membrane protein HorH
HP1106_Hypothetical protein HP1106
HP1105_Putative lipopolysaccharide biosynthesis protein
HP1104_Cinnamyl–alcohol dehydrogenase ELI3–2 Cad
HP1103_Glucokinase
HP1102_6–phosphogluconolactonase
HP1101_Glucose–6–phosphate 1–dehydrogenase G6pD
HP1100_6–phosphogluconate dehydratase
HP1099_2–keto–3–deoxy–6–phosphogluconate aldolase
HP1098_Beta–lactamase HcpC
HP1092_Flagellar basal–body rod protein FlgG
HP1091_Alpha–ketoglutarate permease
HP1090_DNA translocase FtsK
HP1089_Hypothetical protein HP1089
HP1088_Transketolase
HP1087_Bifunctional riboflavin kinase/FMN adenylyltransferase
HP1086_Hemolysin– Tly
HP1085_Hypothetical protein HP1085
HP1084_Aspartate carbamoyltransferase PyrB
HP1083_Outer membrane protein HofB
HP1082_Multidrug resistance protein MsbA
HP1081_Hypothetical protein HP1081
HP1077_High–affinity nickel–transport protein NixA
HP1076_Hypothetical protein HP1076
HP1075_Purine nucleoside phosphorylase HP1075
HP1073_Copper ion binding protein CopB
HP1072_Copper–transporting ATPase CopA
HP1069_Cell division protein FtsH
HP1068_Ribosomal protein L11 methyltransferase
HP1067_Chemotaxis protein CheY
HP1066_Outer membrane protein HorD
HP1062_S–adenosylmethionine:tRNA ribosyltransferase–isomerase
HP1061_Twin arginine–targeting protein translocase TatC
HP1060_Sec–independent translocase TatB
HP1059_Holliday junction DNA helicase RuvB
HP1058_3–methyl–2–oxobutanoate hydroxymethyltransferase PanB
HP1057_Hypothetical protein HP1057
HP1055_Hypothetical protein HP1055
HP1054_Hypothetical protein HP1054
HP1053_Septum formation inhibitor MinC
HP1052_UDP–3–O–[3–hydroxymyristoyl] N–acetylglucosamine deacetylase
HP1051_Universal bacterial protein YeaZ
HP1050_Homoserine kinase ThrB
HP1049_Hypothetical protein HP1049
HP1048_Translation initiation factor IF–2
HP1047_Ribosome–binding factor A
HP1045_Acetyl–CoA synthetase AcoE
HP1043_Homeostatic response regulator HsrA
HP1042_Putative phosphoesterase RecJ–like protein
HP1041_Flagellar biosynthesis protein FlhA
HP1040_30S ribosomal protein S15
HP1039_O–antigen polymerase
HP1038_3–dehydroquinate dehydratase

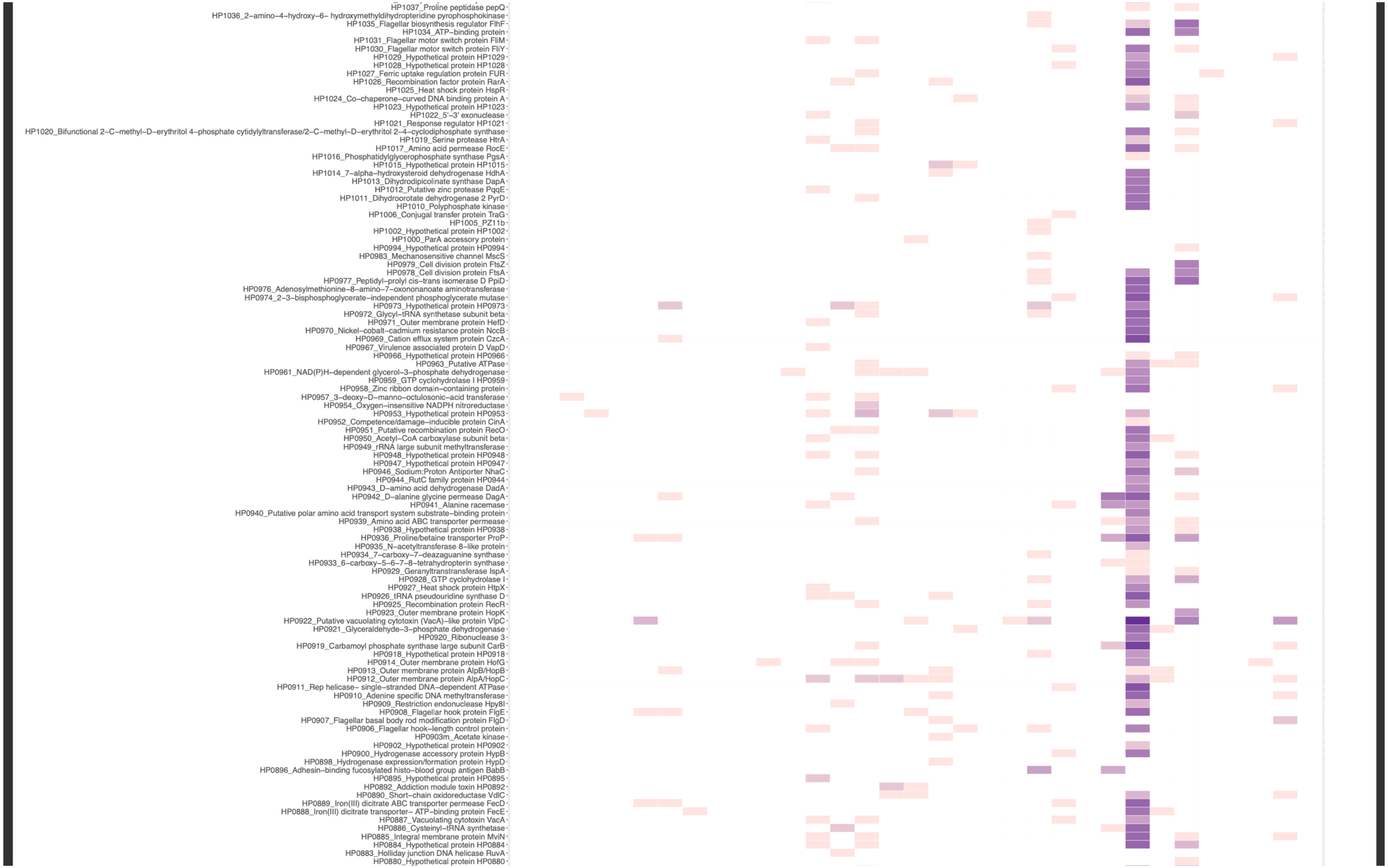

HP1037_Proline peptidase pepQ
HP1036_2–amino–4–hydroxy–6– hydroxymethyldihydropteridine pyrophosphokinase
HP1035_Flagellar biosynthesis regulator FlhF
HP1034_ATP–binding protein
HP1031_Flagellar motor switch protein FliM
HP1030_Flagellar motor switch protein FliY
HP1029_Hypothetical protein HP1029
HP1028_Hypothetical protein HP1028
HP1027_Ferric uptake regulation protein FUR
HP1026_Recombination factor protein RarA
HP1025_Heat shock protein HspR
HP1024_Co–chaperone–curved DNA binding protein A
HP1023_Hypothetical protein HP1023
HP1022_5'–3' exonuclease
HP1021_Response regulator HP1021
HP1020_Bifunctional 2–C–methyl–D–erythritol 4–phosphate cytidylyltransferase/2–C–methyl–D–erythritol 2–4–cyclodiphosphate synthase
HP1019_Serine protease HtrA
HP1017_Amino acid permease RocE
HP1016_Phosphatidylglycerophosphate synthase PgsA
HP1015_Hypothetical protein HP1015
HP1014_7–alpha–hydroxysteroid dehydrogenase HdhA
HP1013_Dihydrodipicolinate synthase DapA
HP1012_Putative zinc protease PqqE
HP1011_Dihydroorotate dehydrogenase 2 PyrD
HP1010_Polyphosphate kinase
HP1006_Conjugal transfer protein TraG
HP1005_PZ11b
HP1002_Hypothetical protein HP1002
HP1000_ParA accessory protein
HP0994_Hypothetical protein HP0994
HP0983_Mechanosensitive channel MscS
HP0979_Cell division protein FtsZ
HP0978_Cell division protein FtsA
HP0977_Peptidyl–prolyl cis–trans isomerase D PpiD
HP0976_Adenosylmethionine–8–amino–7–oxononanoate aminotransferase
HP0974_2–3–bisphosphoglycerate–independent phosphoglycerate mutase
HP0973_Hypothetical protein HP0973
HP0972_Glycyl–tRNA synthetase subunit beta
HP0971_Outer membrane protein HefD
HP0970_Nickel–cobalt–cadmium resistance protein NccB
HP0969_Cation efflux system protein CzcA
HP0967_Virulence associated protein D VapD
HP0966_Hypothetical protein HP0966
HP0963_Putative ATPase
HP0961_NAD(P)H–dependent glycerol–3–phosphate dehydrogenase
HP0959_GTP cyclohydrolase I HP0959
HP0958_Zinc ribbon domain–containing protein
HP0957_3–deoxy–D–manno–octulosonic–acid transferase
HP0954_Oxygen–insensitive NADPH nitroreductase
HP0953_Hypothetical protein HP0953
HP0952_Competence/damage–inducible protein CinA
HP0951_Putative recombination protein RecO
HP0950_Acetyl–CoA carboxylase subunit beta
HP0949_rRNA large subunit methyltransferase
HP0948_Hypothetical protein HP0948
HP0947_Hypothetical protein HP0947
HP0946_Sodium:Proton Antiporter NhaC
HP0944_RutC family protein HP0944
HP0943_D–amino acid dehydrogenase DadA
HP0942_D–alanine glycine permease DagA
HP0941_Alanine racemase
HP0940_Putative polar amino acid transport system substrate–binding protein
HP0939_Amino acid ABC transporter permease
HP0938_Hypothetical protein HP0938
HP0936_Proline/betaine transporter ProP
HP0935_N–acetyltransferase 8–like protein
HP0934_7–carboxy–7–deazaguanine synthase
HP0933_6–carboxy–5–6–7–8–tetrahydropterin synthase
HP0929_Geranyltranstransferase IspA
HP0928_GTP cyclohydrolase I
HP0927_Heat shock protein HtpX
HP0926_tRNA pseudouridine synthase D
HP0925_Recombination protein RecR
HP0923_Outer membrane protein HopK
HP0922_Putative vacuolating cytotoxin (VacA)–like protein VlpC
HP0921_Glyceraldehyde–3–phosphate dehydrogenase
HP0920_Ribonuclease 3
HP0919_Carbamoyl phosphate synthase large subunit CarB
HP0918_Hypothetical protein HP0918
HP0914_Outer membrane protein HofG
HP0913_Outer membrane protein AlpB/HopB
HP0912_Outer membrane protein AlpA/HopC
HP0911_Rep helicase– single–stranded DNA–dependent ATPase
HP0910_Adenine specific DNA methyltransferase
HP0909_Restriction endonuclease Hpy8I
HP0908_Flagellar hook protein FlgE
HP0907_Flagellar basal body rod modification protein FlgD
HP0906_Flagellar hook–length control protein
HP0903m_Acetate kinase
HP0902_Hypothetical protein HP0902
HP0900_Hydrogenase accessory protein HypB
HP0898_Hydrogenase expression/formation protein HypD
HP0896_Adhesin–binding fucosylated histo–blood group antigen BabB
HP0895_Hypothetical protein HP0895
HP0892_Addiction module toxin HP0892
HP0890_Short–chain oxidoreductase VdlC
HP0889_Iron(III) dicitrate ABC transporter permease FecD
HP0888_Iron(III) dicitrate transporter– ATP–binding protein FecE
HP0887_Vacuolating cytotoxin VacA
HP0886_Cysteinyl–tRNA synthetase
HP0885_Integral membrane protein MviN
HP0884_Hypothetical protein HP0884
HP0883_Holliday junction DNA helicase RuvA
HP0880_Hypothetical protein HP0880

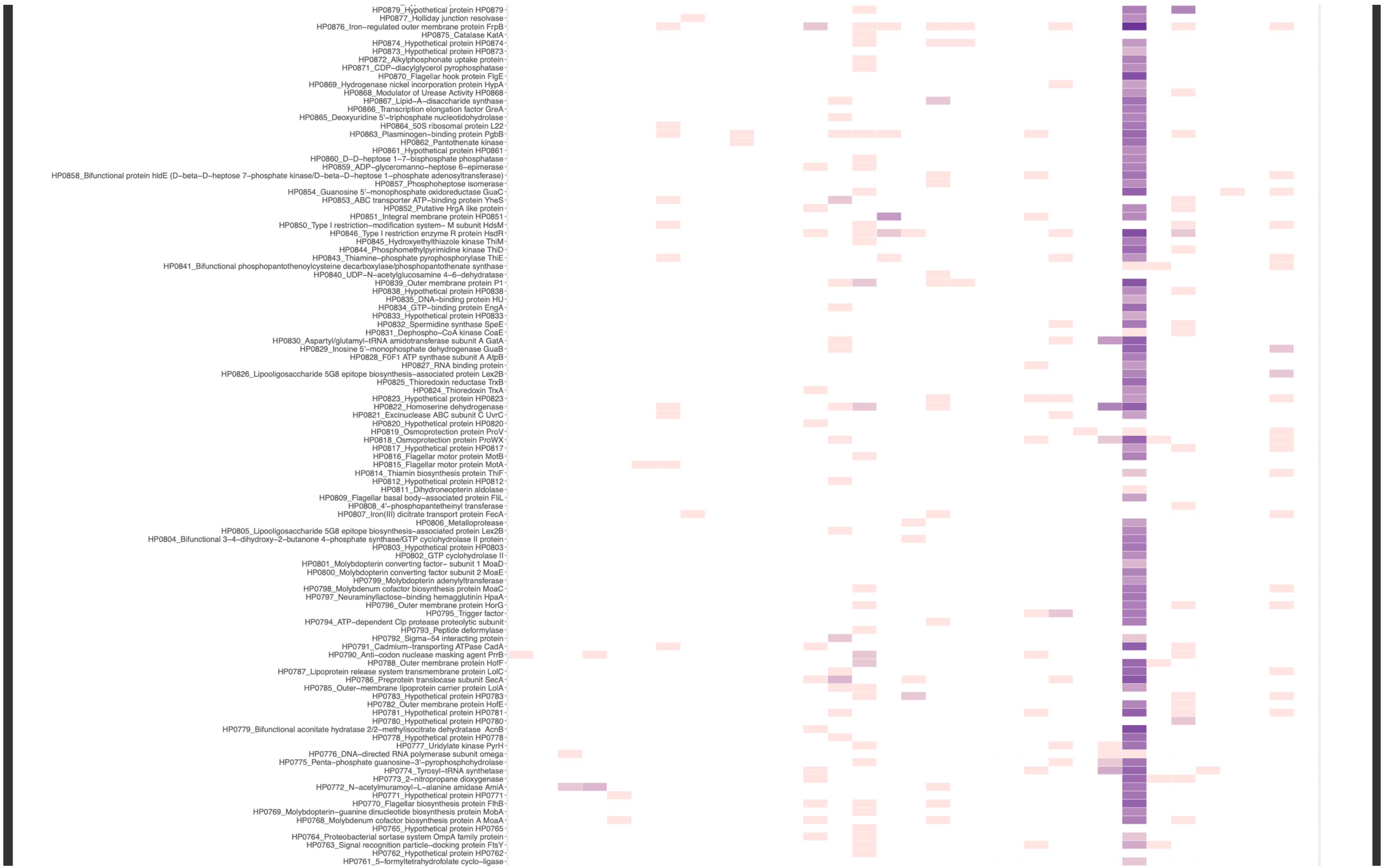

HP0879_Hypothetical protein HP0879
HP0877_Holliday junction resolvase
HP0876_Iron–regulated outer membrane protein FrpB
HP0875_Catalase KatA
HP0874_Hypothetical protein HP0874
HP0873_Hypothetical protein HP0873
HP0872_Alkylphosphonate uptake protein
HP0871_CDP–diacylglycerol pyrophosphatase
HP0870_Flagellar hook protein FlgE
HP0869_Hydrogenase nickel incorporation protein HypA
HP0868_Modulator of Urease Activity HP0868
HP0867_Lipid–A–disaccharide synthase
HP0866_Transcription elongation factor GreA
HP0865_Deoxyuridine 5'–triphosphate nucleotidohydrolase
HP0864_50S ribosomal protein L22
HP0863_Plasminogen–binding protein PgbB
HP0862_Pantothenate kinase
HP0861_Hypothetical protein HP0861
HP0860_D–D–heptose 1–7–bisphosphate phosphatase
HP0859_ADP–glyceromanno–heptose 6–epimerase
HP0858_Bifunctional protein hldE (D–beta–D–heptose 7–phosphate kinase/D–beta–D–heptose 1–phosphate adenosyltransferase)
HP0857_Phosphoheptose isomerase
HP0854_Guanosine 5'–monophosphate oxidoreductase GuaC
HP0853_ABC transporter ATP–binding protein YheS
HP0852_Putative HrgA like protein
HP0851_Integral membrane protein HP0851
HP0850_Type I restriction–modification system– M subunit HdsM
HP0846_Type I restriction enzyme R protein HsdR
HP0845_Hydroxyethylthiazole kinase ThiM
HP0844_Phosphomethylpyrimidine kinase ThiD
HP0843_Thiamine–phosphate pyrophosphorylase ThiE
HP0841_Bifunctional phosphopantothenoylcysteine decarboxylase/phosphopantothenate synthase
HP0840_UDP–N–acetylglucosamine 4–6–dehydratase
HP0839_Outer membrane protein P1
HP0838_Hypothetical protein HP0838
HP0835_DNA–binding protein HU
HP0834_GTP–binding protein EngA
HP0833_Hypothetical protein HP0833
HP0832_Spermidine synthase SpeE
HP0831_Dephospho–CoA kinase CoaE
HP0830_Aspartyl/glutamyl–tRNA amidotransferase subunit A GatA
HP0829_Inosine 5'–monophosphate dehydrogenase GuaB
HP0828_F0F1 ATP synthase subunit A AtpB
HP0827_RNA binding protein
HP0826_Lipooligosaccharide 5G8 epitope biosynthesis–associated protein Lex2B
HP0825_Thioredoxin reductase TrxB
HP0824_Thioredoxin TrxA
HP0823_Hypothetical protein HP0823
HP0822_Homoserine dehydrogenase
HP0821_Excinuclease ABC subunit C UvrC
HP0820_Hypothetical protein HP0820
HP0819_Osmoprotection protein ProV
HP0818_Osmoprotection protein ProWX
HP0817_Hypothetical protein HP0817
HP0816_Flagellar motor protein MotB
HP0815_Flagellar motor protein MotA
HP0814_Thiamin biosynthesis protein ThiF
HP0812_Hypothetical protein HP0812
HP0811_Dihydroneopterin aldolase
HP0809_Flagellar basal body–associated protein FliL
HP0808_4'–phosphopantetheinyl transferase
HP0807_Iron(III) dicitrate transport protein FecA
HP0806_Metalloprotease
HP0805_Lipooligosaccharide 5G8 epitope biosynthesis–associated protein Lex2B
HP0804_Bifunctional 3–4–dihydroxy–2–butanone 4–phosphate synthase/GTP cyclohydrolase II protein
HP0803_Hypothetical protein HP0803
HP0802_GTP cyclohydrolase II
HP0801_Molybdopterin converting factor– subunit 1 MoaD
HP0800_Molybdopterin converting factor subunit 2 MoaE
HP0799_Molybdopterin adenylyltransferase
HP0798_Molybdenum cofactor biosynthesis protein MoaC
HP0797_Neuraminyllactose–binding hemagglutinin HpaA
HP0796_Outer membrane protein HorG
HP0795_Trigger factor
HP0794_ATP–dependent Clp protease proteolytic subunit
HP0793_Peptide deformylase
HP0792_Sigma–54 interacting protein
HP0791_Cadmium–transporting ATPase CadA
HP0790_Anti–codon nuclease masking agent PrrB
HP0788_Outer membrane protein HofF
HP0787_Lipoprotein release system transmembrane protein LolC
HP0786_Preprotein translocase subunit SecA
HP0785_Outer–membrane lipoprotein carrier protein LolA
HP0783_Hypothetical protein HP0783
HP0782_Outer membrane protein HofE
HP0781_Hypothetical protein HP0781
HP0780_Hypothetical protein HP0780
HP0779_Bifunctional aconitate hydratase 2/2–methylisocitrate dehydratase AcnB
HP0778_Hypothetical protein HP0778
HP0777_Uridylate kinase PyrH
HP0776_DNA–directed RNA polymerase subunit omega
HP0775_Penta–phosphate guanosine–3'–pyrophosphohydrolase
HP0774_Tyrosyl–tRNA synthetase
HP0773_2–nitropropane dioxygenase
HP0772_N–acetylmuramoyl–L–alanine amidase AmiA
HP0771_Hypothetical protein HP0771
HP0770_Flagellar biosynthesis protein FlhB
HP0769_Molybdopterin–guanine dinucleotide biosynthesis protein MobA
HP0768_Molybdenum cofactor biosynthesis protein A MoaA
HP0765_Hypothetical protein HP0765
HP0764_Proteobacterial sortase system OmpA family protein
HP0763_Signal recognition particle–docking protein FtsY
HP0762_Hypothetical protein HP0762
HP0761_5–formyltetrahydrofolate cyclo–ligase

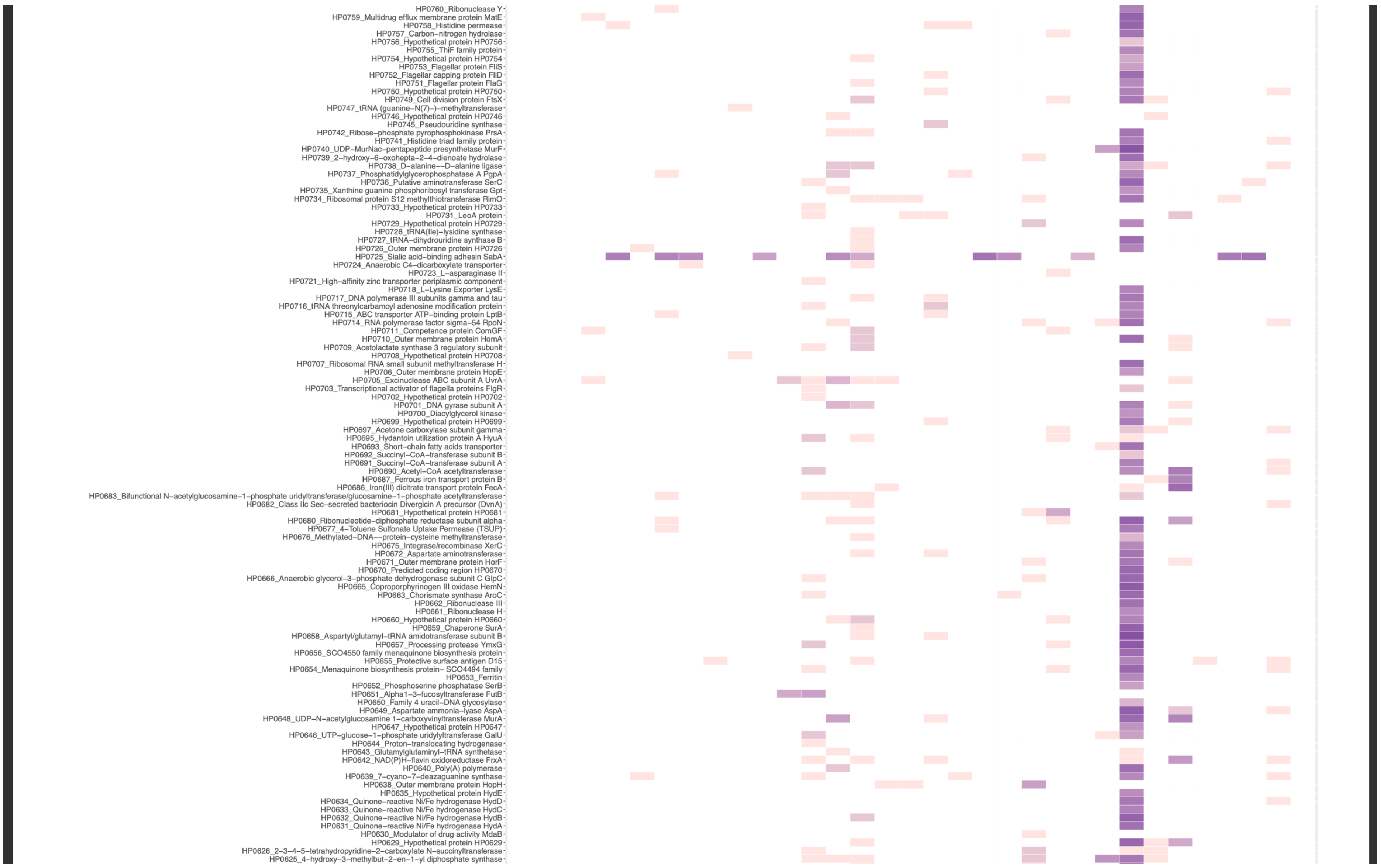
HP0760_Ribonuclease Y
HP0759_Multidrug efflux membrane protein MatE
HP0758_Histidine permease
HP0757_Carbon–nitrogen hydrolase
HP0756_Hypothetical protein HP0756
HP0755_ThiF family protein
HP0754_Hypothetical protein HP0754
HP0753_Flagellar protein FliS
HP0752_Flagellar capping protein FliD
HP0751_Flagellar protein FlaG
HP0750_Hypothetical protein HP0750
HP0749_Cell division protein FtsX
HP0747_tRNA (guanine–N(7)–)–methyltransferase
HP0746_Hypothetical protein HP0746
HP0745_Pseudouridine synthase
HP0742_Ribose–phosphate pyrophosphokinase PrsA
HP0741_Histidine triad family protein
HP0740_UDP–MurNac–pentapeptide presynthetase MurF
HP0739_2–hydroxy–6–oxohepta–2–4–dienoate hydrolase
HP0738_D–alanine––D–alanine ligase
HP0737_Phosphatidylglycerophosphatase A PgpA
HP0736_Putative aminotransferase SerC
HP0735_Xanthine guanine phosphoribosyl transferase Gpt
HP0734_Ribosomal protein S12 methylthiotransferase RimO
HP0733_Hypothetical protein HP0733
HP0731_LeoA protein
HP0729_Hypothetical protein HP0729
HP0728_tRNA(Ile)–lysidine synthase
HP0727_tRNA–dihydrouridine synthase B
HP0726_Outer membrane protein HP0726
HP0725_Sialic acid–binding adhesin SabA
HP0724_Anaerobic C4–dicarboxylate transporter
HP0723_L–asparaginase II
HP0721_High–affinity zinc transporter periplasmic component
HP0718_L–Lysine Exporter LysE
HP0717_DNA polymerase III subunits gamma and tau
HP0716_tRNA threonylcarbamoyl adenosine modification protein
HP0715_ABC transporter ATP–binding protein LptB
HP0714_RNA polymerase factor sigma–54 RpoN
HP0711_Competence protein ComGF
HP0710_Outer membrane protein HomA
HP0709_Acetolactate synthase 3 regulatory subunit
HP0708_Hypothetical protein HP0708
HP0707_Ribosomal RNA small subunit methyltransferase H
HP0706_Outer membrane protein HopE
HP0705_Excinuclease ABC subunit A UvrA
HP0703_Transcriptional activator of flagella proteins FlgR
HP0702_Hypothetical protein HP0702
HP0701_DNA gyrase subunit A
HP0700_Diacylglycerol kinase
HP0699_Hypothetical protein HP0699
HP0697_Acetone carboxylase subunit gamma
HP0695_Hydantoin utilization protein A HyuA
HP0693_Short–chain fatty acids transporter
HP0692_Succinyl–CoA–transferase subunit B
HP0691_Succinyl–CoA–transferase subunit A
HP0690_Acetyl–CoA acetyltransferase
HP0687_Ferrous iron transport protein B
HP0686_Iron(III) dicitrate transport protein FecA
HP0683_Bifunctional N–acetylglucosamine–1–phosphate uridyltransferase/glucosamine–1–phosphate acetyltransferase
HP0682_Class IIc Sec–secreted bacteriocin Divergicin A precursor (DvnA)
HP0681_Hypothetical protein HP0681
HP0680_Ribonucleotide–diphosphate reductase subunit alpha
HP0677_4–Toluene Sulfonate Uptake Permease (TSUP)
HP0676_Methylated–DNA––protein–cysteine methyltransferase
HP0675_Integrase/recombinase XerC
HP0672_Aspartate aminotransferase
HP0671_Outer membrane protein HorF
HP0670_Predicted coding region HP0670
HP0666_Anaerobic glycerol–3–phosphate dehydrogenase subunit C GlpC
HP0665_Coproporphyrinogen III oxidase HemN
HP0663_Chorismate synthase AroC
HP0662_Ribonuclease III
HP0661_Ribonuclease H
HP0660_Hypothetical protein HP0660
HP0659_Chaperone SurA
HP0658_Aspartyl/glutamyl–tRNA amidotransferase subunit B
HP0657_Processing protease YmxG
HP0656_SCO4550 family menaquinone biosynthesis protein
HP0655_Protective surface antigen D15
HP0654_Menaquinone biosynthesis protein– SCO4494 family
HP0653_Ferritin
HP0652_Phosphoserine phosphatase SerB
HP0651_Alpha1–3–fucosyltransferase FutB
HP0650_Family 4 uracil–DNA glycosylase
HP0649_Aspartate ammonia–lyase AspA
HP0648_UDP–N–acetylglucosamine 1–carboxyvinyltransferase MurA
HP0647_Hypothetical protein HP0647
HP0646_UTP–glucose–1–phosphate uridylyltransferase GalU
HP0644_Proton–translocating hydrogenase
HP0643_Glutamylglutaminyl–tRNA synthetase
HP0642_NAD(P)H–flavin oxidoreductase FrxA
HP0640_Poly(A) polymerase
HP0639_7–cyano–7–deazaguanine synthase
HP0638_Outer membrane protein HopH
HP0635_Hypothetical protein HydE
HP0634_Quinone–reactive Ni/Fe hydrogenase HydD
HP0633_Quinone–reactive Ni/Fe hydrogenase HydC
HP0632_Quinone–reactive Ni/Fe hydrogenase HydB
HP0631_Quinone–reactive Ni/Fe hydrogenase HydA
HP0630_Modulator of drug activity MdaB
HP0629_Hypothetical protein HP0629
HP0626_2–3–4–5–tetrahydropyridine–2–carboxylate N–succinyltransferase
HP0625_4–hydroxy–3–methylbut–2–en–1–yl diphosphate synthase

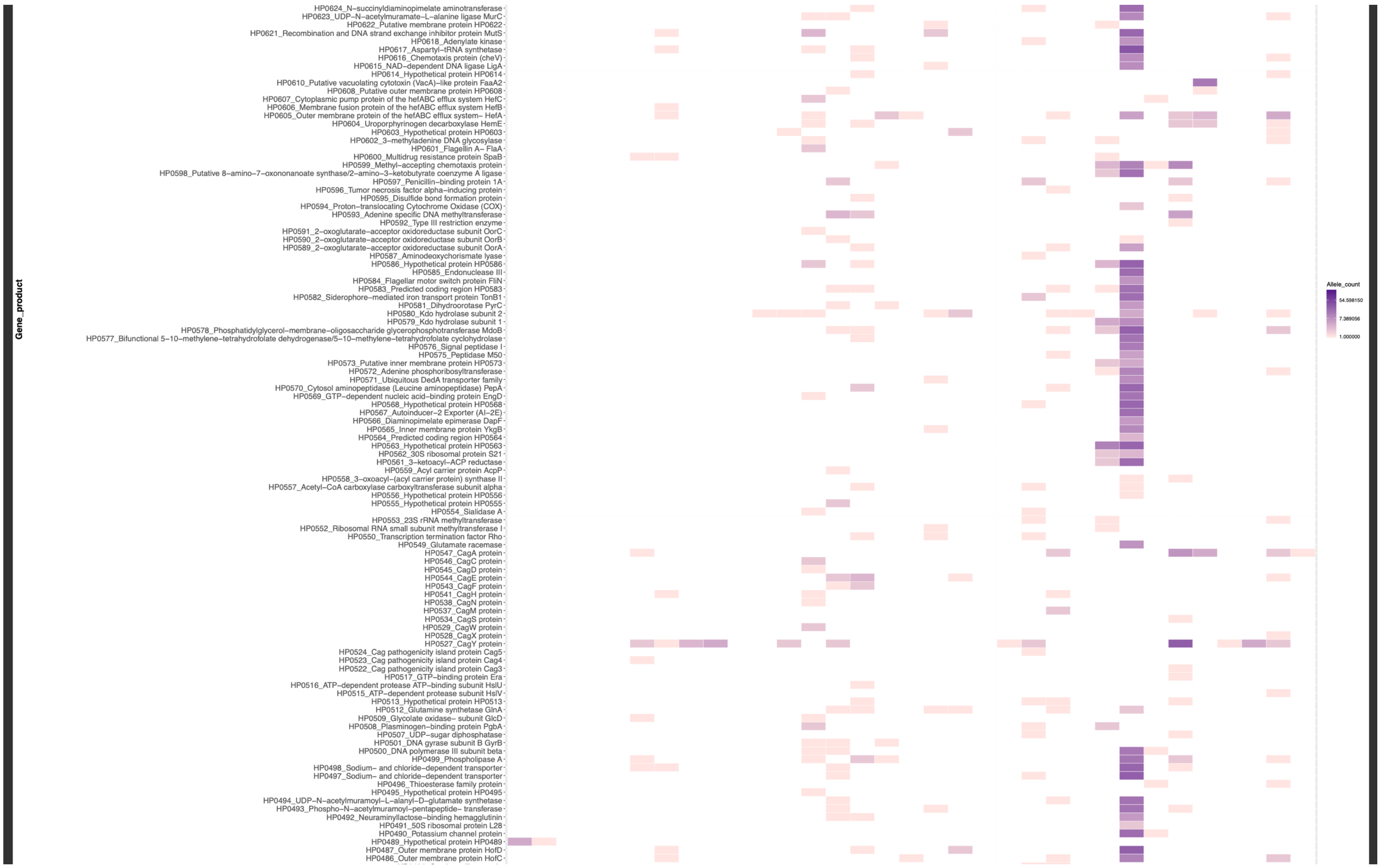

Gene_product
HP0624_N–succinyldiaminopimelate aminotransferase
HP0623_UDP–N–acetylmuramate–L–alanine ligase MurC
HP0622_Putative membrane protein HP0622
HP0621_Recombination and DNA strand exchange inhibitor protein MutS
HP0618_Adenylate kinase
HP0617_Aspartyl–tRNA synthetase
HP0616_Chemotaxis protein (cheV)
HP0615_NAD–dependent DNA ligase LigA
HP0614_Hypothetical protein HP0614
HP0610_Putative vacuolating cytotoxin (VacA)–like protein FaaA2
HP0608_Putative outer membrane protein HP0608
HP0607_Cytoplasmic pump protein of the hefABC efflux system HefC
HP0606_Membrane fusion protein of the hefABC efflux system HefB
HP0605_Outer membrane protein of the hefABC efflux system– HefA
HP0604_Uroporphyrinogen decarboxylase HemE
HP0603_Hypothetical protein HP0603
HP0602_3–methyladenine DNA glycosylase
HP0601_Flagellin A– FlaA
HP0600_Multidrug resistance protein SpaB
HP0599_Methyl–accepting chemotaxis protein
HP0598_Putative 8–amino–7–oxononanoate synthase/2–amino–3–ketobutyrate coenzyme A ligase
HP0597_Penicillin–binding protein 1A
HP0596_Tumor necrosis factor alpha–inducing protein
HP0595_Disulfide bond formation protein
HP0594_Proton–translocating Cytochrome Oxidase (COX)
HP0593_Adenine specific DNA methyltransferase
HP0592_Type III restriction enzyme
HP0591_2–oxoglutarate–acceptor oxidoreductase subunit OorC
HP0590_2–oxoglutarate–acceptor oxidoreductase subunit OorB
HP0589_2–oxoglutarate–acceptor oxidoreductase subunit OorA
HP0587_Aminodeoxychorismate lyase
HP0586_Hypothetical protein HP0586
HP0585_Endonuclease III
HP0584_Flagellar motor switch protein FliN
HP0583_Predicted coding region HP0583
HP0582_Siderophore–mediated iron transport protein TonB1
HP0581_Dihydroorotase PyrC
HP0580_Kdo hydrolase subunit 2
HP0579_Kdo hydrolase subunit 1
HP0578_Phosphatidylglycerol–membrane–oligosaccharide glycerophosphotransferase MdoB
HP0577_Bifunctional 5–10–methylene–tetrahydrofolate dehydrogenase/5–10–methylene–tetrahydrofolate cyclohydrolase
HP0576_Signal peptidase I
HP0575_Peptidase M50
HP0573_Putative inner membrane protein HP0573
HP0572_Adenine phosphoribosyltransferase
HP0571_Ubiquitous DedA transporter family
HP0570_Cytosol aminopeptidase (Leucine aminopeptidase) PepA
HP0569_GTP–dependent nucleic acid–binding protein EngD
HP0568_Hypothetical protein HP0568
HP0567_Autoinducer–2 Exporter (AI–2E)
HP0566_Diaminopimelate epimerase DapF
HP0565_Inner membrane protein YkgB
HP0564_Predicted coding region HP0564
HP0563_Hypothetical protein HP0563
HP0562_30S ribosomal protein S21
HP0561_3–ketoacyl–ACP reductase
HP0559_Acyl carrier protein AcpP
HP0558_3–oxoacyl–(acyl carrier protein) synthase II
HP0557_Acetyl–CoA carboxylase carboxyltransferase subunit alpha
HP0556_Hypothetical protein HP0556
HP0555_Hypothetical protein HP0555
HP0554_Sialidase A
HP0553_23S rRNA methyltransferase
HP0552_Ribosomal RNA small subunit methyltransferase I
HP0550_Transcription termination factor Rho
HP0549_Glutamate racemase
HP0547_CagA protein
HP0546_CagC protein
HP0545_CagD protein
HP0544_CagE protein
HP0543_CagF protein
HP0541_CagH protein
HP0538_CagN protein
HP0537_CagM protein
HP0534_CagS protein
HP0529_CagW protein
HP0528_CagX protein
HP0527_CagY protein
HP0524_Cag pathogenicity island protein Cag5
HP0523_Cag pathogenicity island protein Cag4
HP0522_Cag pathogenicity island protein Cag3
HP0517_GTP–binding protein Era
HP0516_ATP–dependent protease ATP–binding subunit HslU
HP0515_ATP–dependent protease subunit HslV
HP0513_Hypothetical protein HP0513
HP0512_Glutamine synthetase GlnA
HP0509_Glycolate oxidase– subunit GlcD
HP0508_Plasminogen–binding protein PgbA
HP0507_UDP–sugar diphosphatase
HP0501_DNA gyrase subunit B GyrB
HP0500_DNA polymerase III subunit beta
HP0499_Phospholipase A
HP0498_Sodium– and chloride–dependent transporter
HP0497_Sodium– and chloride–dependent transporter
HP0496_Thioesterase family protein
HP0495_Hypothetical protein HP0495
HP0494_UDP–N–acetylmuramoyl–L–alanyl–D–glutamate synthetase
HP0493_Phospho–N–acetylmuramoyl–pentapeptide– transferase
HP0492_Neuraminyllactose–binding hemagglutinin
HP0491_50S ribosomal protein L28
HP0490_Potassium channel protein
HP0489_Hypothetical protein HP0489
HP0487_Outer membrane protein HofD
HP0486_Outer membrane protein HofC
Allele_count
54.598150
7.389056
1.000000

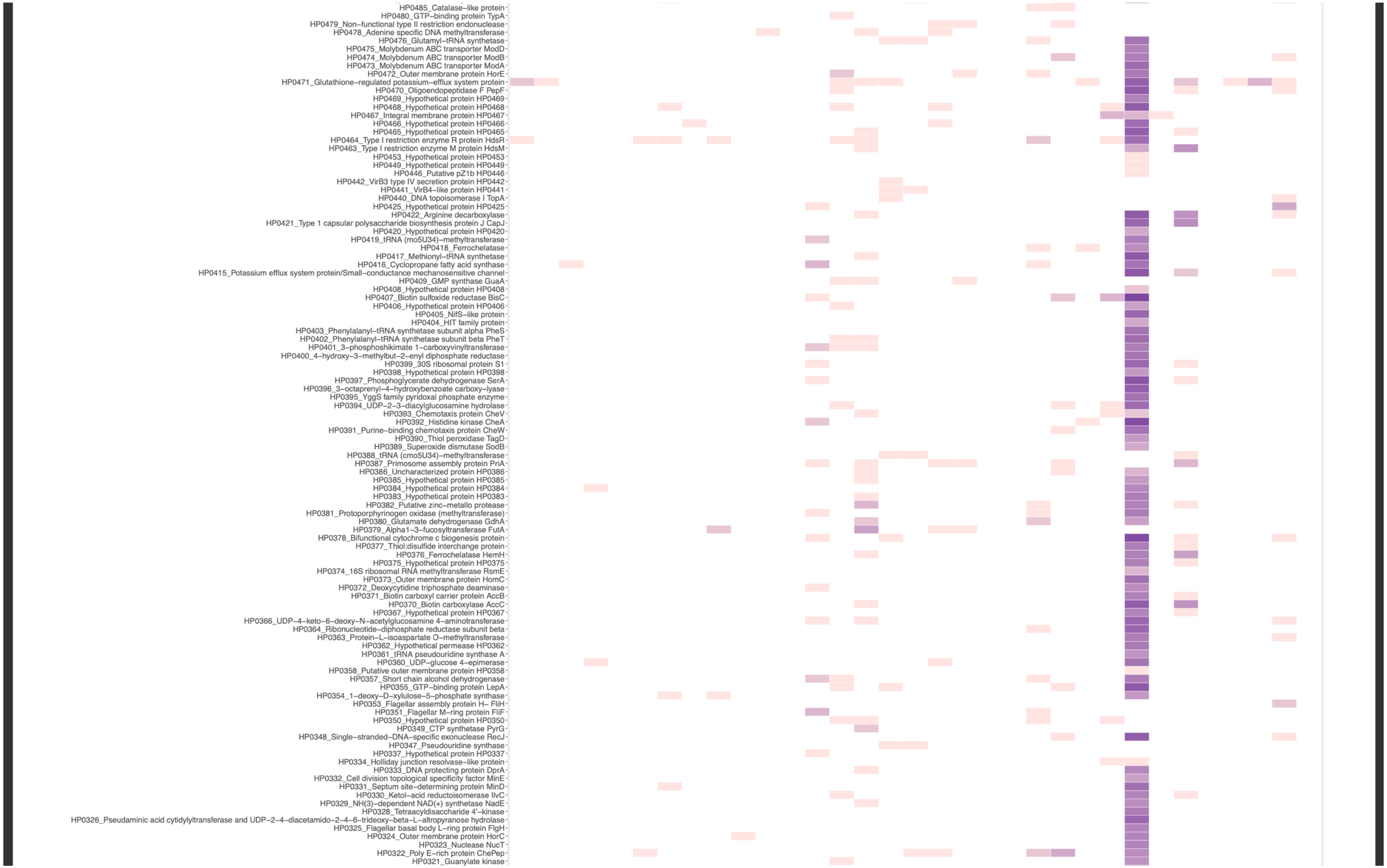
HP0485_Catalase–like protein
HP0480_GTP–binding protein TypA
HP0479_Non–functional type II restriction endonuclease
HP0478_Adenine specific DNA methyltransferase
HP0476_Glutamyl–tRNA synthetase
HP0475_Molybdenum ABC transporter ModD
HP0474_Molybdenum ABC transporter ModB
HP0473_Molybdenum ABC transporter ModA
HP0472_Outer membrane protein HorE
HP0471_Glutathione–regulated potassium–efflux system protein
HP0470_Oligoendopeptidase F PepF
HP0469_Hypothetical protein HP0469
HP0468_Hypothetical protein HP0468
HP0467_Integral membrane protein HP0467
HP0466_Hypothetical protein HP0466
HP0465_Hypothetical protein HP0465
HP0464_Type I restriction enzyme R protein HdsR
HP0463_Type I restriction enzyme M protein HdsM
HP0453_Hypothetical protein HP0453
HP0449_Hypothetical protein HP0449
HP0446_Putative pZ1b HP0446
HP0442_VirB3 type IV secretion protein HP0442
HP0441_VirB4–like protein HP0441
HP0440_DNA topoisomerase I TopA
HP0425_Hypothetical protein HP0425
HP0422_Arginine decarboxylase
HP0421_Type 1 capsular polysaccharide biosynthesis protein J CapJ
HP0420_Hypothetical protein HP0420
HP0419_tRNA (mo5U34)–methyltransferase
HP0418_Ferrochelatase
HP0417_Methionyl–tRNA synthetase
HP0416_Cyclopropane fatty acid synthase
HP0415_Potassium efflux system protein/Small–conductance mechanosensitive channel
HP0409_GMP synthase GuaA
HP0408_Hypothetical protein HP0408
HP0407_Biotin sulfoxide reductase BisC
HP0406_Hypothetical protein HP0406
HP0405_NifS–like protein
HP0404_HIT family protein
HP0403_Phenylalanyl–tRNA synthetase subunit alpha PheS
HP0402_Phenylalanyl–tRNA synthetase subunit beta PheT
HP0401_3–phosphoshikimate 1–carboxyvinyltransferase
HP0400_4–hydroxy–3–methylbut–2–enyl diphosphate reductase
HP0399_30S ribosomal protein S1
HP0398_Hypothetical protein HP0398
HP0397_Phosphoglycerate dehydrogenase SerA
HP0396_3–octaprenyl–4–hydroxybenzoate carboxy–lyase
HP0395_YggS family pyridoxal phosphate enzyme
HP0394_UDP–2–3–diacylglucosamine hydrolase
HP0393_Chemotaxis protein CheV
HP0392_Histidine kinase CheA
HP0391_Purine–binding chemotaxis protein CheW
HP0390_Thiol peroxidase TagD
HP0389_Superoxide dismutase SodB
HP0388_tRNA (cmo5U34)–methyltransferase
HP0387_Primosome assembly protein PriA
HP0386_Uncharacterized protein HP0386
HP0385_Hypothetical protein HP0385
HP0384_Hypothetical protein HP0384
HP0383_Hypothetical protein HP0383
HP0382_Putative zinc–metallo protease
HP0381_Protoporphyrinogen oxidase (methyltransferase)
HP0380_Glutamate dehydrogenase GdhA
HP0379_Alpha1–3–fucosyltransferase FutA
HP0378_Bifunctional cytochrome c biogenesis protein
HP0377_Thiol:disulfide interchange protein
HP0376_Ferrochelatase HemH
HP0375_Hypothetical protein HP0375
HP0374_16S ribosomal RNA methyltransferase RsmE
HP0373_Outer membrane protein HomC
HP0372_Deoxycytidine triphosphate deaminase
HP0371_Biotin carboxyl carrier protein AccB
HP0370_Biotin carboxylase AccC
HP0367_Hypothetical protein HP0367
HP0366_UDP–4–keto–6–deoxy–N–acetylglucosamine 4–aminotransferase
HP0364_Ribonucleotide–diphosphate reductase subunit beta
HP0363_Protein–L–isoaspartate O–methyltransferase
HP0362_Hypothetical permease HP0362
HP0361_tRNA pseudouridine synthase A
HP0360_UDP–glucose 4–epimerase
HP0358_Putative outer membrane protein HP0358
HP0357_Short chain alcohol dehydrogenase
HP0355_GTP–binding protein LepA
HP0354_1–deoxy–D–xylulose–5–phosphate synthase
HP0353_Flagellar assembly protein H– FliH
HP0351_Flagellar M–ring protein FliF
HP0350_Hypothetical protein HP0350
HP0349_CTP synthetase PyrG
HP0348_Single–stranded–DNA–specific exonuclease RecJ
HP0347_Pseudouridine synthase
HP0337_Hypothetical protein HP0337
HP0334_Holliday junction resolvase–like protein
HP0333_DNA protecting protein DprA
HP0332_Cell division topological specificity factor MinE
HP0331_Septum site–determining protein MinD
HP0330_Ketol–acid reductoisomerase IlvC
HP0329_NH(3)–dependent NAD(+) synthetase NadE
HP0328_Tetraacyldisaccharide 4'–kinase
HP0326_Pseudaminic acid cytidylyltransferase and UDP–2–4–diacetamido–2–4–6–trideoxy–beta–L–altropyranose hydrolase
HP0325_Flagellar basal body L–ring protein FlgH
HP0324_Outer membrane protein HorC
HP0323_Nuclease NucT
HP0322_Poly E–rich protein ChePep
HP0321_Guanylate kinase

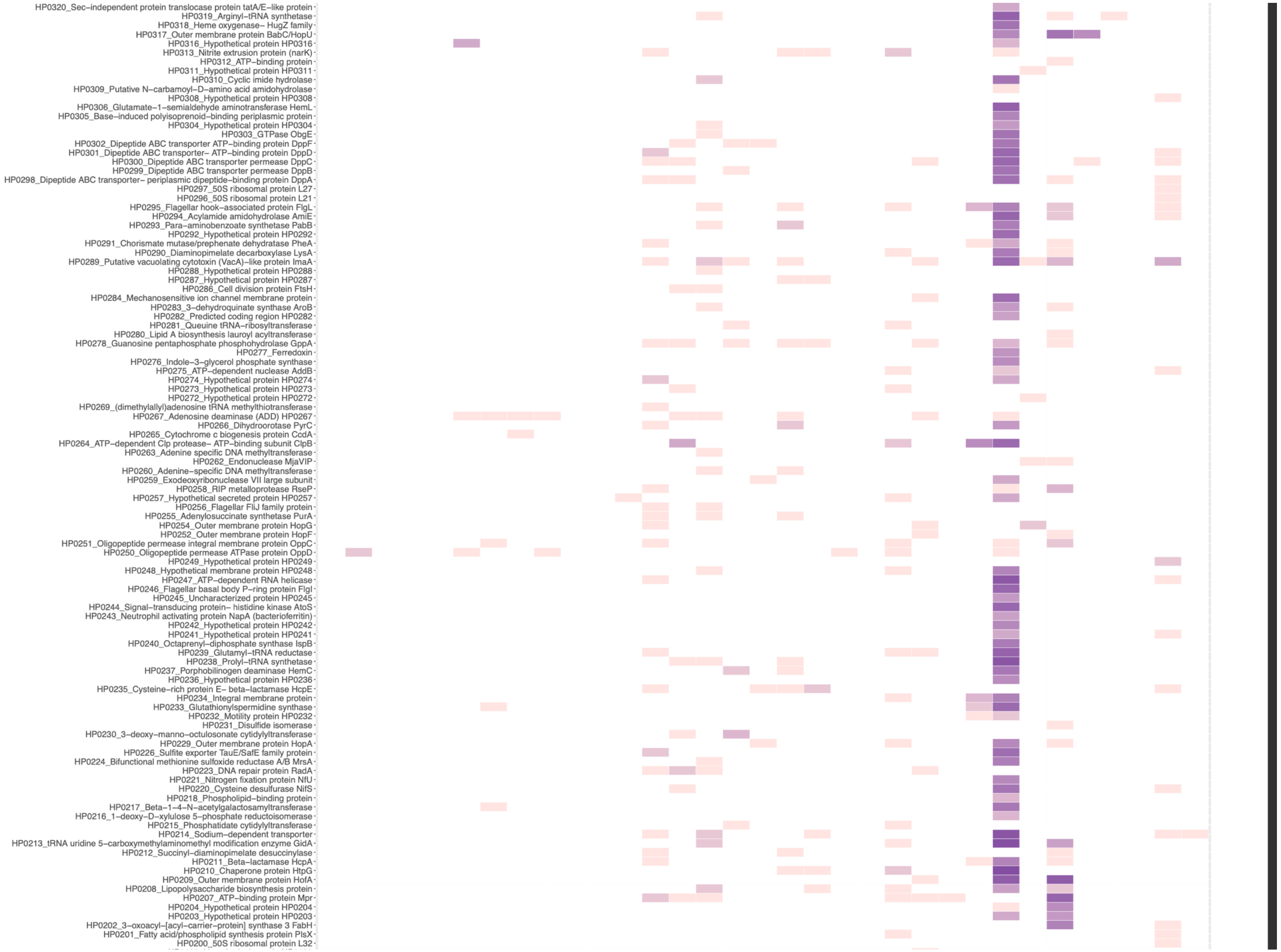

HP0320_Sec–independent protein translocase protein tatA/E–like protein
HP0319_Arginyl–tRNA synthetase
HP0318_Heme oxygenase– HugZ family
HP0317_Outer membrane protein BabC/HopU
HP0316_Hypothetical protein HP0316
HP0313_Nitrite extrusion protein (narK)
HP0312_ATP–binding protein
HP0311_Hypothetical protein HP0311
HP0310_Cyclic imide hydrolase
HP0309_Putative N–carbamoyl–D–amino acid amidohydrolase
HP0308_Hypothetical protein HP0308
HP0306_Glutamate–1–semialdehyde aminotransferase HemL
HP0305_Base–induced polyisoprenoid–binding periplasmic protein
HP0304_Hypothetical protein HP0304
HP0303_GTPase ObgE
HP0302_Dipeptide ABC transporter ATP–binding protein DppF
HP0301_Dipeptide ABC transporter– ATP–binding protein DppD
HP0300_Dipeptide ABC transporter permease DppC
HP0299_Dipeptide ABC transporter permease DppB
HP0298_Dipeptide ABC transporter– periplasmic dipeptide–binding protein DppA
HP0297_50S ribosomal protein L27
HP0296_50S ribosomal protein L21
HP0295_Flagellar hook–associated protein FlgL
HP0294_Acylamide amidohydrolase AmiE
HP0293_Para–aminobenzoate synthetase PabB
HP0292_Hypothetical protein HP0292
HP0291_Chorismate mutase/prephenate dehydratase PheA
HP0290_Diaminopimelate decarboxylase LysA
HP0289_Putative vacuolating cytotoxin (VacA)–like protein ImaA
HP0288_Hypothetical protein HP0288
HP0287_Hypothetical protein HP0287
HP0286_Cell division protein FtsH
HP0284_Mechanosensitive ion channel membrane protein
HP0283_3–dehydroquinate synthase AroB
HP0282_Predicted coding region HP0282
HP0281_Queuine tRNA–ribosyltransferase
HP0280_Lipid A biosynthesis lauroyl acyltransferase
HP0278_Guanosine pentaphosphate phosphohydrolase GppA
HP0277_Ferredoxin
HP0276_Indole–3–glycerol phosphate synthase
HP0275_ATP–dependent nuclease AddB
HP0274_Hypothetical protein HP0274
HP0273_Hypothetical protein HP0273
HP0272_Hypothetical protein HP0272
HP0269_(dimethylallyl)adenosine tRNA methylthiotransferase
HP0267_Adenosine deaminase (ADD) HP0267
HP0266_Dihydroorotase PyrC
HP0265_Cytochrome c biogenesis protein CcdA
HP0264_ATP–dependent Clp protease– ATP–binding subunit ClpB
HP0263_Adenine specific DNA methyltransferase
HP0262_Endonuclease MjaVIP
HP0260_Adenine–specific DNA methyltransferase
HP0259_Exodeoxyribonuclease VII large subunit
HP0258_RIP metalloprotease RseP
HP0257_Hypothetical secreted protein HP0257
HP0256_Flagellar FliJ family protein
HP0255_Adenylosuccinate synthetase PurA
HP0254_Outer membrane protein HopG
HP0252_Outer membrane protein HopF
HP0251_Oligopeptide permease integral membrane protein OppC
HP0250_Oligopeptide permease ATPase protein OppD
HP0249_Hypothetical protein HP0249
HP0248_Hypothetical membrane protein HP0248
HP0247_ATP–dependent RNA helicase
HP0246_Flagellar basal body P–ring protein FlgI
HP0245_Uncharacterized protein HP0245
HP0244_Signal–transducing protein– histidine kinase AtoS
HP0243_Neutrophil activating protein NapA (bacterioferritin)
HP0242_Hypothetical protein HP0242
HP0241_Hypothetical protein HP0241
HP0240_Octaprenyl–diphosphate synthase IspB
HP0239_Glutamyl–tRNA reductase
HP0238_Prolyl–tRNA synthetase
HP0237_Porphobilinogen deaminase HemC
HP0236_Hypothetical protein HP0236
HP0235_Cysteine–rich protein E– beta–lactamase HcpE
HP0234_Integral membrane protein
HP0233_Glutathionylspermidine synthase
HP0232_Motility protein HP0232
HP0231_Disulfide isomerase
HP0230_3–deoxy–manno–octulosonate cytidylyltransferase
HP0229_Outer membrane protein HopA
HP0226_Sulfite exporter TauE/SafE family protein
HP0224_Bifunctional methionine sulfoxide reductase A/B MrsA
HP0223_DNA repair protein RadA
HP0221_Nitrogen fixation protein NfU
HP0220_Cysteine desulfurase NifS
HP0218_Phospholipid–binding protein
HP0217_Beta–1–4–N–acetylgalactosamyltransferase
HP0216_1–deoxy–D–xylulose 5–phosphate reductoisomerase
HP0215_Phosphatidate cytidylyltransferase
HP0214_Sodium–dependent transporter
HP0213_tRNA uridine 5–carboxymethylaminomethyl modification enzyme GidA
HP0212_Succinyl–diaminopimelate desuccinylase
HP0211_Beta–lactamase HcpA
HP0210_Chaperone protein HtpG
HP0209_Outer membrane protein HofA
HP0208_Lipopolysaccharide biosynthesis protein
HP0207_ATP–binding protein Mpr
HP0204_Hypothetical protein HP0204
HP0203_Hypothetical protein HP0203
HP0202_3–oxoacyl–[acyl–carrier–protein] synthase 3 FabH
HP0201_Fatty acid/phospholipid synthesis protein PlsX
HP0200_50S ribosomal protein L32

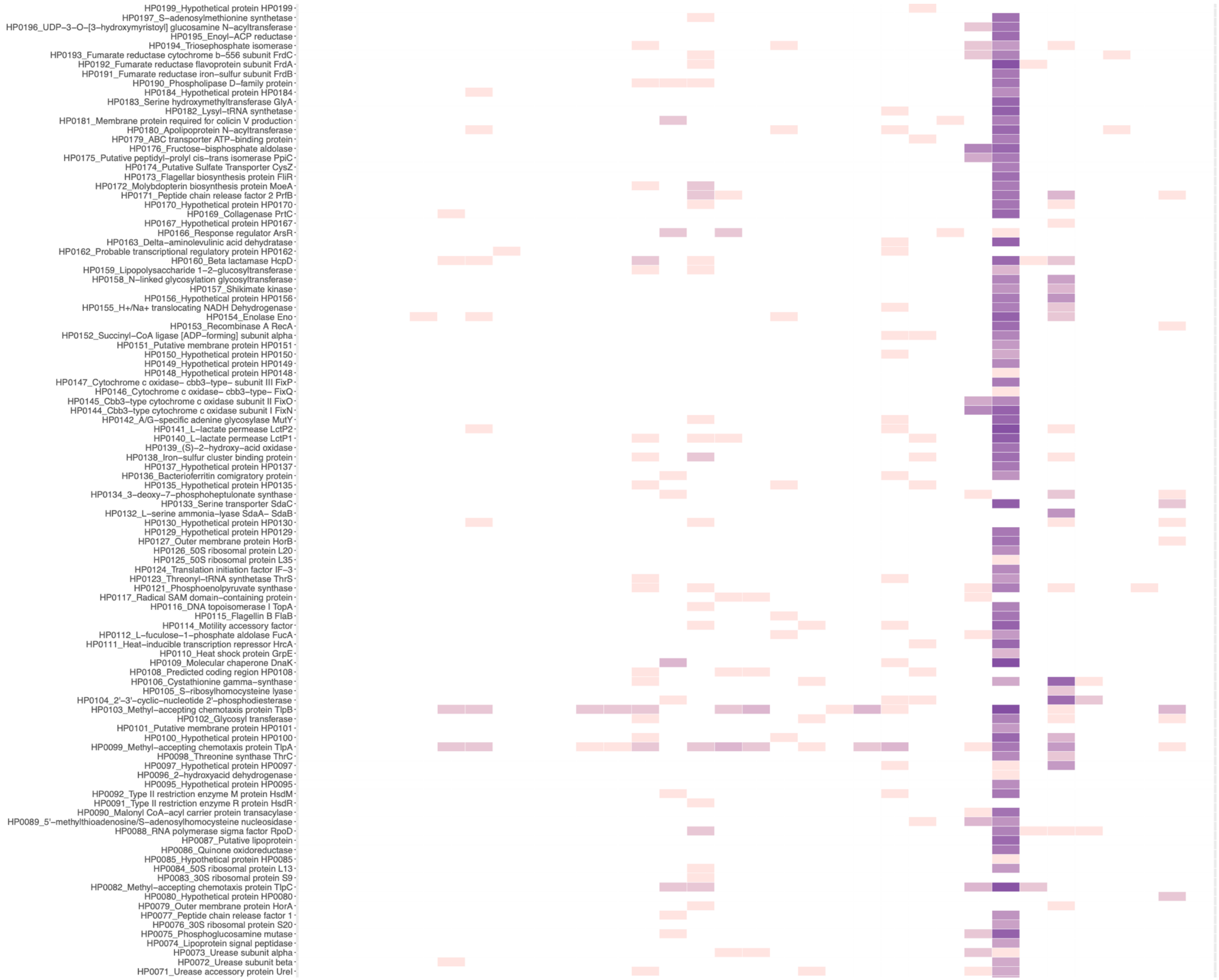
HP0199_Hypothetical protein HP0199
HP0197_S–adenosylmethionine synthetase
HP0196_UDP–3–O–[3–hydroxymyristoyl] glucosamine N–acyltransferase
HP0195_Enoyl–ACP reductase
HP0194_Triosephosphate isomerase
HP0193_Fumarate reductase cytochrome b–556 subunit FrdC
HP0192_Fumarate reductase flavoprotein subunit FrdA
HP0191_Fumarate reductase iron–sulfur subunit FrdB
HP0190_Phospholipase D–family protein
HP0184_Hypothetical protein HP0184
HP0183_Serine hydroxymethyltransferase GlyA
HP0182_Lysyl–tRNA synthetase
HP0181_Membrane protein required for colicin V production
HP0180_Apolipoprotein N–acyltransferase
HP0179_ABC transporter ATP–binding protein
HP0176_Fructose–bisphosphate aldolase
HP0175_Putative peptidyl–prolyl cis–trans isomerase PpiC
HP0174_Putative Sulfate Transporter CysZ
HP0173_Flagellar biosynthesis protein FliR
HP0172_Molybdopterin biosynthesis protein MoeA
HP0171_Peptide chain release factor 2 PrfB
HP0170_Hypothetical protein HP0170
HP0169_Collagenase PrtC
HP0167_Hypothetical protein HP0167
HP0166_Response regulator ArsR
HP0163_Delta–aminolevulinic acid dehydratase
HP0162_Probable transcriptional regulatory protein HP0162
HP0160_Beta lactamase HcpD
HP0159_Lipopolysaccharide 1–2–glucosyltransferase
HP0158_N–linked glycosylation glycosyltransferase
HP0157_Shikimate kinase
HP0156_Hypothetical protein HP0156
HP0155_H+/Na+ translocating NADH Dehydrogenase
HP0154_Enolase Eno
HP0153_Recombinase A RecA
HP0152_Succinyl–CoA ligase [ADP–forming] subunit alpha
HP0151_Putative membrane protein HP0151
HP0150_Hypothetical protein HP0150
HP0149_Hypothetical protein HP0149
HP0148_Hypothetical protein HP0148
HP0147_Cytochrome c oxidase– cbb3–type– subunit III FixP
HP0146_Cytochrome c oxidase– cbb3–type– FixQ
HP0145_Cbb3–type cytochrome c oxidase subunit II FixO
HP0144_Cbb3–type cytochrome c oxidase subunit I FixN
HP0142_A/G–specific adenine glycosylase MutY
HP0141_L–lactate permease LctP2
HP0140_L–lactate permease LctP1
HP0139_(S)–2–hydroxy–acid oxidase
HP0138_Iron–sulfur cluster binding protein
HP0137_Hypothetical protein HP0137
HP0136_Bacterioferritin comigratory protein
HP0135_Hypothetical protein HP0135
HP0134_3–deoxy–7–phosphoheptulonate synthase
HP0133_Serine transporter SdaC
HP0132_L–serine ammonia–lyase SdaA– SdaB
HP0130_Hypothetical protein HP0130
HP0129_Hypothetical protein HP0129
HP0127_Outer membrane protein HorB
HP0126_50S ribosomal protein L20
HP0125_50S ribosomal protein L35
HP0124_Translation initiation factor IF–3
HP0123_Threonyl–tRNA synthetase ThrS
HP0121_Phosphoenolpyruvate synthase
HP0117_Radical SAM domain–containing protein
HP0116_DNA topoisomerase I TopA
HP0115_Flagellin B FlaB
HP0114_Motility accessory factor
HP0112_L–fuculose–1–phosphate aldolase FucA
HP0111_Heat–inducible transcription repressor HrcA
HP0110_Heat shock protein GrpE
HP0109_Molecular chaperone DnaK
HP0108_Predicted coding region HP0108
HP0106_Cystathionine gamma–synthase
HP0105_S–ribosylhomocysteine lyase
HP0104_2'–3'–cyclic–nucleotide 2'–phosphodiesterase
HP0103_Methyl–accepting chemotaxis protein TlpB
HP0102_Glycosyl transferase
HP0101_Putative membrane protein HP0101
HP0100_Hypothetical protein HP0100
HP0099_Methyl–accepting chemotaxis protein TlpA
HP0098_Threonine synthase ThrC
HP0097_Hypothetical protein HP0097
HP0096_2–hydroxyacid dehydrogenase
HP0095_Hypothetical protein HP0095
HP0092_Type II restriction enzyme M protein HsdM
HP0091_Type II restriction enzyme R protein HsdR
HP0090_Malonyl CoA–acyl carrier protein transacylase
HP0089_5'–methylthioadenosine/S–adenosylhomocysteine nucleosidase
HP0088_RNA polymerase sigma factor RpoD
HP0087_Putative lipoprotein
HP0086_Quinone oxidoreductase
HP0085_Hypothetical protein HP0085
HP0084_50S ribosomal protein L13
HP0083_30S ribosomal protein S9
HP0082_Methyl–accepting chemotaxis protein TlpC
HP0080_Hypothetical protein HP0080
HP0079_Outer membrane protein HorA
HP0077_Peptide chain release factor 1
HP0076_30S ribosomal protein S20
HP0075_Phosphoglucosamine mutase
HP0074_Lipoprotein signal peptidase
HP0073_Urease subunit alpha
HP0072_Urease subunit beta
HP0071_Urease accessory protein UreI

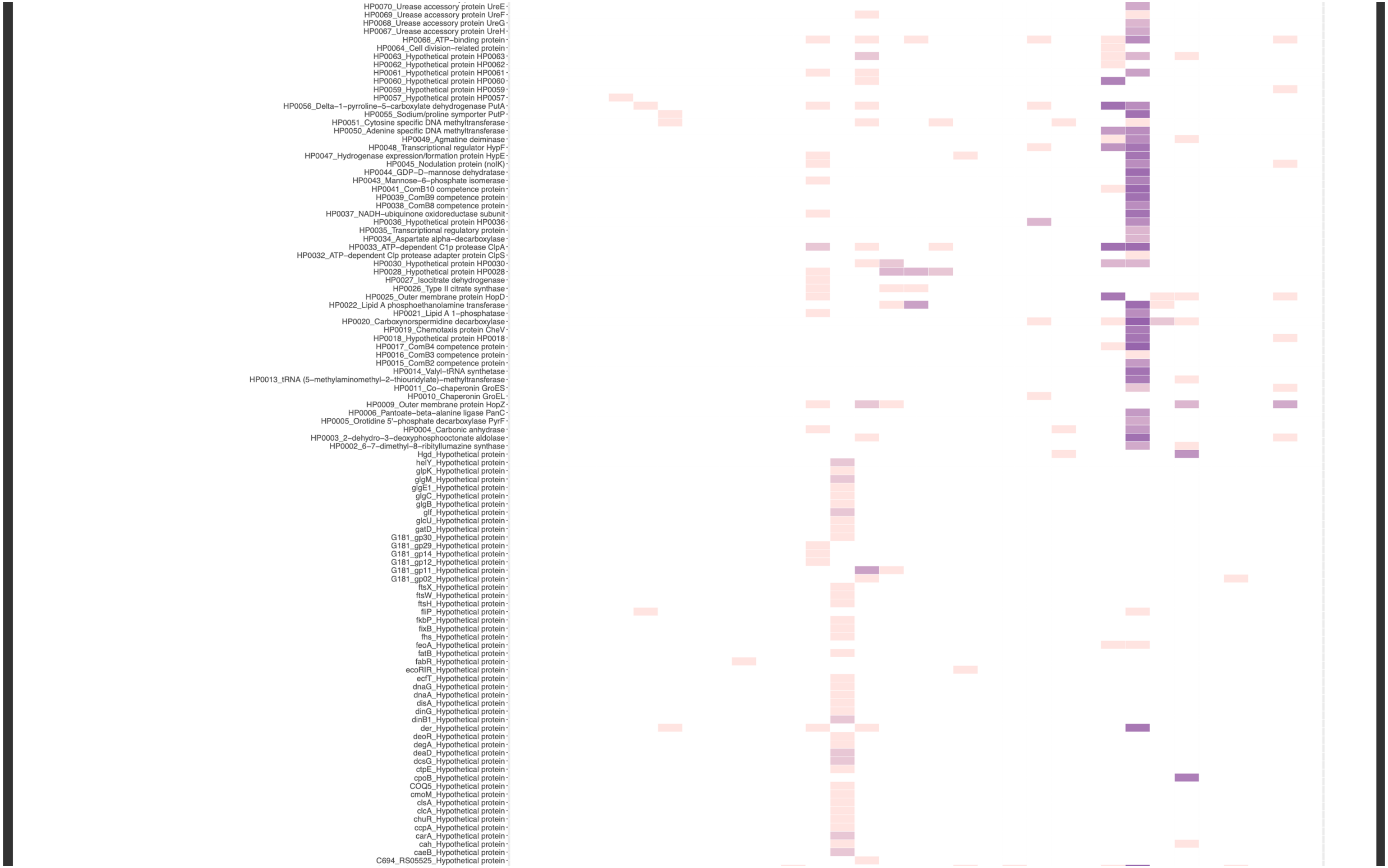
HP0070_Urease accessory protein UreE
HP0069_Urease accessory protein UreF
HP0068_Urease accessory protein UreG
HP0067_Urease accessory protein UreH
HP0066_ATP–binding protein
HP0064_Cell division–related protein
HP0063_Hypothetical protein HP0063
HP0062_Hypothetical protein HP0062
HP0061_Hypothetical protein HP0061
HP0060_Hypothetical protein HP0060
HP0059_Hypothetical protein HP0059
HP0057_Hypothetical protein HP0057
HP0056_Delta–1–pyrroline–5–carboxylate dehydrogenase PutA
HP0055_Sodium/proline symporter PutP
HP0051_Cytosine specific DNA methyltransferase
HP0050_Adenine specific DNA methyltransferase
HP0049_Agmatine deiminase
HP0048_Transcriptional regulator HypF
HP0047_Hydrogenase expression/formation protein HypE
HP0045_Nodulation protein (nolK)
HP0044_GDP–D–mannose dehydratase
HP0043_Mannose–6–phosphate isomerase
HP0041_ComB10 competence protein
HP0039_ComB9 competence protein
HP0038_ComB8 competence protein
HP0037_NADH–ubiquinone oxidoreductase subunit
HP0036_Hypothetical protein HP0036
HP0035_Transcriptional regulatory protein
HP0034_Aspartate alpha–decarboxylase
HP0033_ATP–dependent C1p protease ClpA
HP0032_ATP–dependent Clp protease adapter protein ClpS
HP0030_Hypothetical protein HP0030
HP0028_Hypothetical protein HP0028
HP0027_Isocitrate dehydrogenase
HP0026_Type II citrate synthase
HP0025_Outer membrane protein HopD
HP0022_Lipid A phosphoethanolamine transferase
HP0021_Lipid A 1–phosphatase
HP0020_Carboxynorspermidine decarboxylase
HP0019_Chemotaxis protein CheV
HP0018_Hypothetical protein HP0018
HP0017_ComB4 competence protein
HP0016_ComB3 competence protein
HP0015_ComB2 competence protein
HP0014_Valyl–tRNA synthetase
HP0013_tRNA (5–methylaminomethyl–2–thiouridylate)–methyltransferase
HP0011_Co–chaperonin GroES
HP0010_Chaperonin GroEL
HP0009_Outer membrane protein HopZ
HP0006_Pantoate–beta–alanine ligase PanC
HP0005_Orotidine 5'–phosphate decarboxylase PyrF
HP0004_Carbonic anhydrase
HP0003_2–dehydro–3–deoxyphosphooctonate aldolase
HP0002_6–7–dimethyl–8–ribityllumazine synthase
Hgd_Hypothetical protein
helY_Hypothetical protein
glpK_Hypothetical protein
glgM_Hypothetical protein
glgE1_Hypothetical protein
glgC_Hypothetical protein
glgB_Hypothetical protein
glf_Hypothetical protein
glcU_Hypothetical protein
gatD_Hypothetical protein
G181_gp30_Hypothetical protein
G181_gp29_Hypothetical protein
G181_gp14_Hypothetical protein
G181_gp12_Hypothetical protein
G181_gp11_Hypothetical protein
G181_gp02_Hypothetical protein
ftsX_Hypothetical protein
ftsW_Hypothetical protein
ftsH_Hypothetical protein
fliP_Hypothetical protein
fkbP_Hypothetical protein
fixB_Hypothetical protein
fhs_Hypothetical protein
feoA_Hypothetical protein
fatB_Hypothetical protein
fabR_Hypothetical protein
ecoRIR_Hypothetical protein
ecfT_Hypothetical protein
dnaG_Hypothetical protein
dnaA_Hypothetical protein
disA_Hypothetical protein
dinG_Hypothetical protein
dinB1_Hypothetical protein
der_Hypothetical protein
deoR_Hypothetical protein
degA_Hypothetical protein
deaD_Hypothetical protein
dcsG_Hypothetical protein
ctpE_Hypothetical protein
cpoB_Hypothetical protein
COQ5_Hypothetical protein
cmoM_Hypothetical protein
clsA_Hypothetical protein
clcA_Hypothetical protein
chuR_Hypothetical protein
ccpA_Hypothetical protein
carA_Hypothetical protein
cah_Hypothetical protein
caeB_Hypothetical protein
C694_RS05525_Hypothetical protein

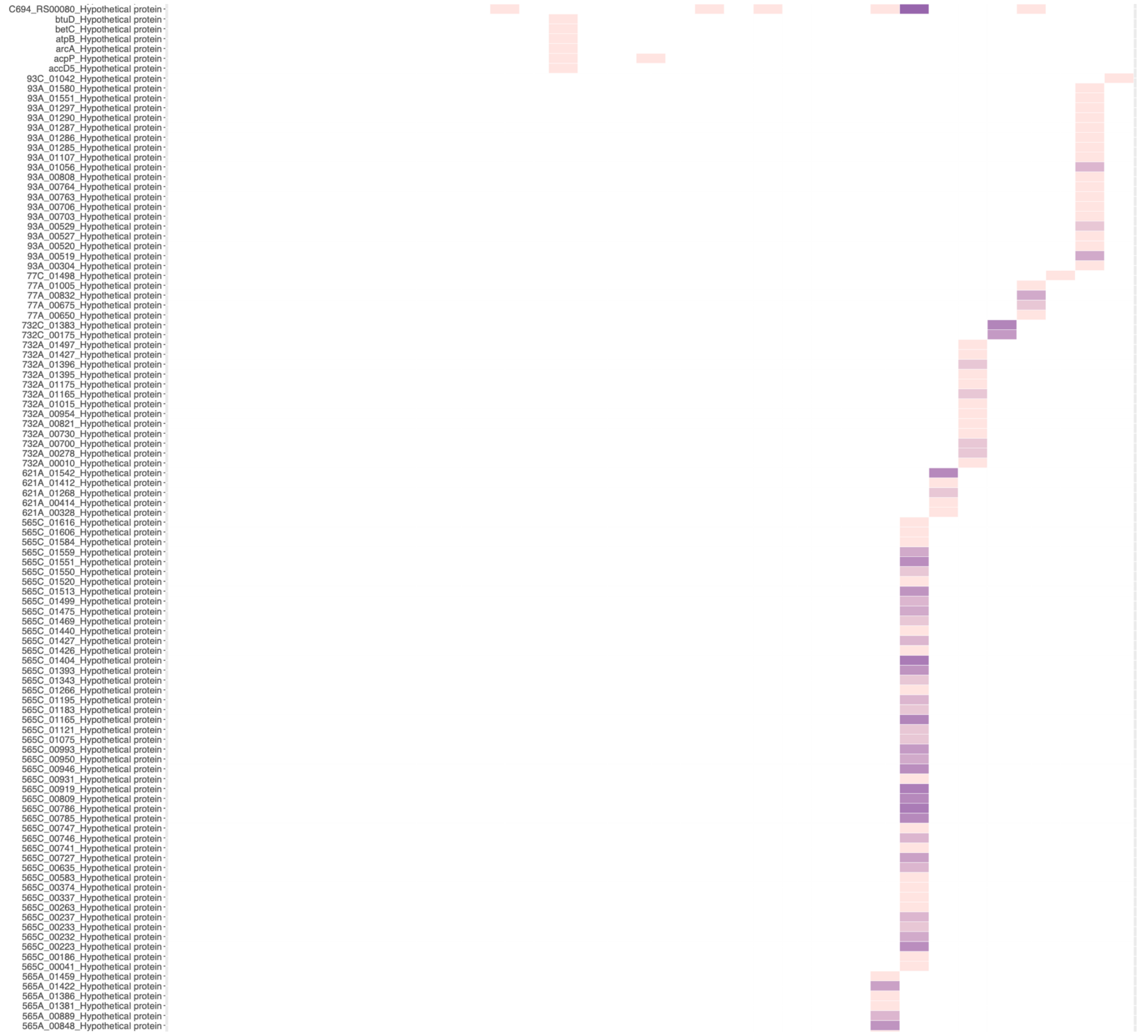
C694_RS00080_Hypothetical protein
btuD_Hypothetical protein
betC_Hypothetical protein
atpB_Hypothetical protein
arcA_Hypothetical protein
acpP_Hypothetical protein
accD5_Hypothetical protein
93C_01042_Hypothetical protein
93A_01580_Hypothetical protein
93A_01551_Hypothetical protein
93A_01297_Hypothetical protein
93A_01290_Hypothetical protein
93A_01287_Hypothetical protein
93A_01286_Hypothetical protein
93A_01285_Hypothetical protein
93A_01107_Hypothetical protein
93A_01056_Hypothetical protein
93A_00808_Hypothetical protein
93A_00764_Hypothetical protein
93A_00763_Hypothetical protein
93A_00706_Hypothetical protein
93A_00703_Hypothetical protein
93A_00529_Hypothetical protein
93A_00527_Hypothetical protein
93A_00520_Hypothetical protein
93A_00519_Hypothetical protein
93A_00304_Hypothetical protein
77C_01498_Hypothetical protein
77A_01005_Hypothetical protein
77A_00832_Hypothetical protein
77A_00675_Hypothetical protein
77A_00650_Hypothetical protein
732C_01383_Hypothetical protein
732C_00175_Hypothetical protein
732A_01497_Hypothetical protein
732A_01427_Hypothetical protein
732A_01396_Hypothetical protein
732A_01395_Hypothetical protein
732A_01175_Hypothetical protein
732A_01165_Hypothetical protein
732A_01015_Hypothetical protein
732A_00954_Hypothetical protein
732A_00821_Hypothetical protein
732A_00730_Hypothetical protein
732A_00700_Hypothetical protein
732A_00278_Hypothetical protein
732A_00010_Hypothetical protein
621A_01542_Hypothetical protein
621A_01412_Hypothetical protein
621A_01268_Hypothetical protein
621A_00414_Hypothetical protein
621A_00328_Hypothetical protein
565C_01616_Hypothetical protein
565C_01606_Hypothetical protein
565C_01584_Hypothetical protein
565C_01559_Hypothetical protein
565C_01551_Hypothetical protein
565C_01550_Hypothetical protein
565C_01520_Hypothetical protein
565C_01513_Hypothetical protein
565C_01499_Hypothetical protein
565C_01475_Hypothetical protein
565C_01469_Hypothetical protein
565C_01440_Hypothetical protein
565C_01427_Hypothetical protein
565C_01426_Hypothetical protein
565C_01404_Hypothetical protein
565C_01393_Hypothetical protein
565C_01343_Hypothetical protein
565C_01266_Hypothetical protein
565C_01195_Hypothetical protein
565C_01183_Hypothetical protein
565C_01165_Hypothetical protein
565C_01121_Hypothetical protein
565C_01075_Hypothetical protein
565C_00993_Hypothetical protein
565C_00950_Hypothetical protein
565C_00946_Hypothetical protein
565C_00931_Hypothetical protein
565C_00919_Hypothetical protein
565C_00809_Hypothetical protein
565C_00786_Hypothetical protein
565C_00785_Hypothetical protein
565C_00747_Hypothetical protein
565C_00746_Hypothetical protein
565C_00741_Hypothetical protein
565C_00727_Hypothetical protein
565C_00635_Hypothetical protein
565C_00583_Hypothetical protein
565C_00374_Hypothetical protein
565C_00337_Hypothetical protein
565C_00263_Hypothetical protein
565C_00237_Hypothetical protein
565C_00233_Hypothetical protein
565C_00232_Hypothetical protein
565C_00223_Hypothetical protein
565C_00186_Hypothetical protein
565C_00041_Hypothetical protein
565A_01459_Hypothetical protein
565A_01422_Hypothetical protein
565A_01386_Hypothetical protein
565A_01381_Hypothetical protein
565A_00889_Hypothetical protein
565A_00848_Hypothetical protein

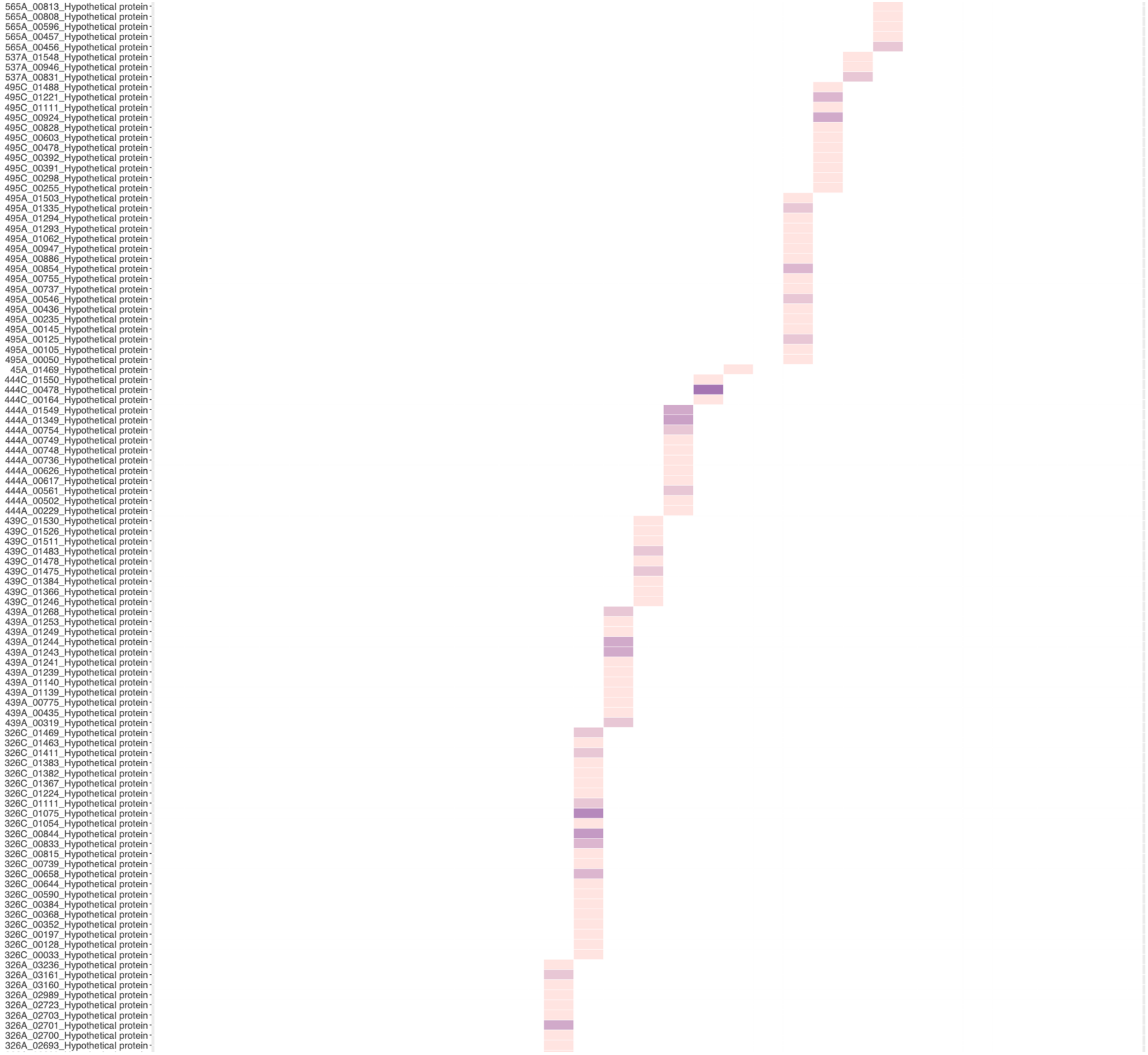
565A_00813_Hypothetical protein
565A_00808_Hypothetical protein
565A_00596_Hypothetical protein
565A_00457_Hypothetical protein
565A_00456_Hypothetical protein
537A_01548_Hypothetical protein
537A_00946_Hypothetical protein
537A_00831_Hypothetical protein
495C_01488_Hypothetical protein
495C_01221_Hypothetical protein
495C_01111_Hypothetical protein
495C_00924_Hypothetical protein
495C_00828_Hypothetical protein
495C_00603_Hypothetical protein
495C_00478_Hypothetical protein
495C_00392_Hypothetical protein
495C_00391_Hypothetical protein
495C_00298_Hypothetical protein
495C_00255_Hypothetical protein
495A_01503_Hypothetical protein
495A_01335_Hypothetical protein
495A_01294_Hypothetical protein
495A_01293_Hypothetical protein
495A_01062_Hypothetical protein
495A_00947_Hypothetical protein
495A_00886_Hypothetical protein
495A_00854_Hypothetical protein
495A_00755_Hypothetical protein
495A_00737_Hypothetical protein
495A_00546_Hypothetical protein
495A_00436_Hypothetical protein
495A_00235_Hypothetical protein
495A_00145_Hypothetical protein
495A_00125_Hypothetical protein
495A_00105_Hypothetical protein
495A_00050_Hypothetical protein
45A_01469_Hypothetical protein
444C_01550_Hypothetical protein
444C_00478_Hypothetical protein
444C_00164_Hypothetical protein
444A_01549_Hypothetical protein
444A_01349_Hypothetical protein
444A_00754_Hypothetical protein
444A_00749_Hypothetical protein
444A_00748_Hypothetical protein
444A_00736_Hypothetical protein
444A_00626_Hypothetical protein
444A_00617_Hypothetical protein
444A_00561_Hypothetical protein
444A_00502_Hypothetical protein
444A_00229_Hypothetical protein
439C_01530_Hypothetical protein
439C_01526_Hypothetical protein
439C_01511_Hypothetical protein
439C_01483_Hypothetical protein
439C_01478_Hypothetical protein
439C_01475_Hypothetical protein
439C_01384_Hypothetical protein
439C_01366_Hypothetical protein
439C_01246_Hypothetical protein
439A_01268_Hypothetical protein
439A_01253_Hypothetical protein
439A_01249_Hypothetical protein
439A_01244_Hypothetical protein
439A_01243_Hypothetical protein
439A_01241_Hypothetical protein
439A_01239_Hypothetical protein
439A_01140_Hypothetical protein
439A_01139_Hypothetical protein
439A_00775_Hypothetical protein
439A_00435_Hypothetical protein
439A_00319_Hypothetical protein
326C_01469_Hypothetical protein
326C_01463_Hypothetical protein
326C_01411_Hypothetical protein
326C_01383_Hypothetical protein
326C_01382_Hypothetical protein
326C_01367_Hypothetical protein
326C_01224_Hypothetical protein
326C_01111_Hypothetical protein
326C_01075_Hypothetical protein
326C_01054_Hypothetical protein
326C_00844_Hypothetical protein
326C_00833_Hypothetical protein
326C_00815_Hypothetical protein
326C_00739_Hypothetical protein
326C_00658_Hypothetical protein
326C_00644_Hypothetical protein
326C_00590_Hypothetical protein
326C_00384_Hypothetical protein
326C_00368_Hypothetical protein
326C_00352_Hypothetical protein
326C_00197_Hypothetical protein
326C_00128_Hypothetical protein
326C_00033_Hypothetical protein
326A_03236_Hypothetical protein
326A_03161_Hypothetical protein
326A_03160_Hypothetical protein
326A_02989_Hypothetical protein
326A_02723_Hypothetical protein
326A_02703_Hypothetical protein
326A_02701_Hypothetical protein
326A_02700_Hypothetical protein
326A_02693_Hypothetical protein

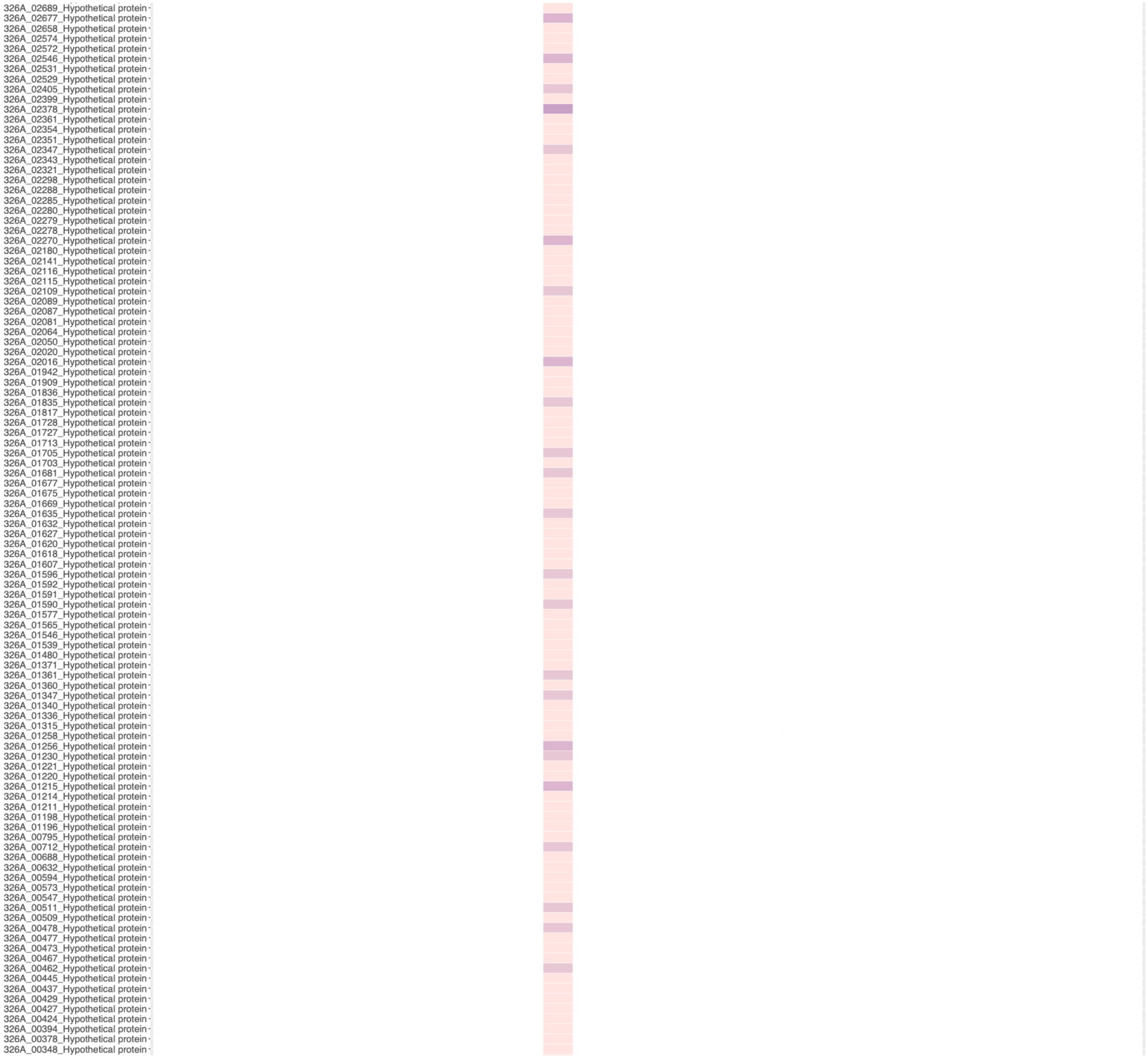
326A_02689_Hypothetical protein
326A_02677_Hypothetical protein
326A_02658_Hypothetical protein
326A_02574_Hypothetical protein
326A_02572_Hypothetical protein
326A_02546_Hypothetical protein
326A_02531_Hypothetical protein
326A_02529_Hypothetical protein
326A_02405_Hypothetical protein
326A_02399_Hypothetical protein
326A_02378_Hypothetical protein
326A_02361_Hypothetical protein
326A_02354_Hypothetical protein
326A_02351_Hypothetical protein
326A_02347_Hypothetical protein
326A_02343_Hypothetical protein
326A_02321_Hypothetical protein
326A_02298_Hypothetical protein
326A_02288_Hypothetical protein
326A_02285_Hypothetical protein
326A_02280_Hypothetical protein
326A_02279_Hypothetical protein
326A_02278_Hypothetical protein
326A_02270_Hypothetical protein
326A_02180_Hypothetical protein
326A_02141_Hypothetical protein
326A_02116_Hypothetical protein
326A_02115_Hypothetical protein
326A_02109_Hypothetical protein
326A_02089_Hypothetical protein
326A_02087_Hypothetical protein
326A_02081_Hypothetical protein
326A_02064_Hypothetical protein
326A_02050_Hypothetical protein
326A_02020_Hypothetical protein
326A_02016_Hypothetical protein
326A_01942_Hypothetical protein
326A_01909_Hypothetical protein
326A_01836_Hypothetical protein
326A_01835_Hypothetical protein
326A_01817_Hypothetical protein
326A_01728_Hypothetical protein
326A_01727_Hypothetical protein
326A_01713_Hypothetical protein
326A_01705_Hypothetical protein
326A_01703_Hypothetical protein
326A_01681_Hypothetical protein
326A_01677_Hypothetical protein
326A_01675_Hypothetical protein
326A_01669_Hypothetical protein
326A_01635_Hypothetical protein
326A_01632_Hypothetical protein
326A_01627_Hypothetical protein
326A_01620_Hypothetical protein
326A_01618_Hypothetical protein
326A_01607_Hypothetical protein
326A_01596_Hypothetical protein
326A_01592_Hypothetical protein
326A_01591_Hypothetical protein
326A_01590_Hypothetical protein
326A_01577_Hypothetical protein
326A_01565_Hypothetical protein
326A_01546_Hypothetical protein
326A_01539_Hypothetical protein
326A_01480_Hypothetical protein
326A_01371_Hypothetical protein
326A_01361_Hypothetical protein
326A_01360_Hypothetical protein
326A_01347_Hypothetical protein
326A_01340_Hypothetical protein
326A_01336_Hypothetical protein
326A_01315_Hypothetical protein
326A_01258_Hypothetical protein
326A_01256_Hypothetical protein
326A_01230_Hypothetical protein
326A_01221_Hypothetical protein
326A_01220_Hypothetical protein
326A_01215_Hypothetical protein
326A_01214_Hypothetical protein
326A_01211_Hypothetical protein
326A_01198_Hypothetical protein
326A_01196_Hypothetical protein
326A_00795_Hypothetical protein
326A_00712_Hypothetical protein
326A_00688_Hypothetical protein
326A_00632_Hypothetical protein
326A_00594_Hypothetical protein
326A_00573_Hypothetical protein
326A_00547_Hypothetical protein
326A_00511_Hypothetical protein
326A_00509_Hypothetical protein
326A_00478_Hypothetical protein
326A_00477_Hypothetical protein
326A_00473_Hypothetical protein
326A_00467_Hypothetical protein
326A_00462_Hypothetical protein
326A_00445_Hypothetical protein
326A_00437_Hypothetical protein
326A_00429_Hypothetical protein
326A_00427_Hypothetical protein
326A_00424_Hypothetical protein
326A_00394_Hypothetical protein
326A_00378_Hypothetical protein
326A_00348_Hypothetical protein

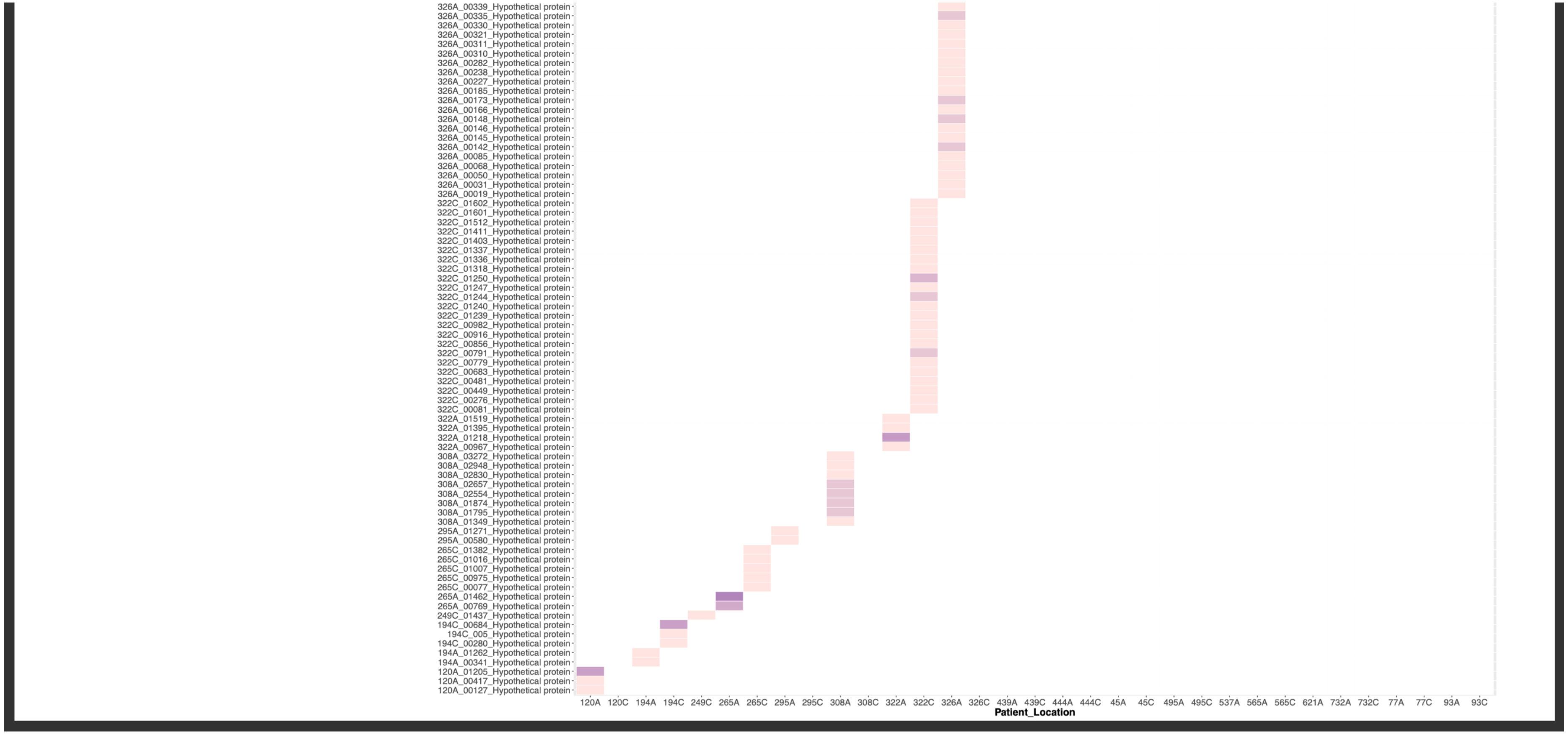


**Suppl. Fig. 18.** Heat maps showing all minor allelic variation across genes/gene products for each antrum- and corpus-derived *H. pylori* population, derived from deep sequencing of the population from each biopsy and read-mapping back to the consensus genome to identify variant bases at each locus. Higher colour intensity indicates a larger number of variant bases within that gene/gene product.

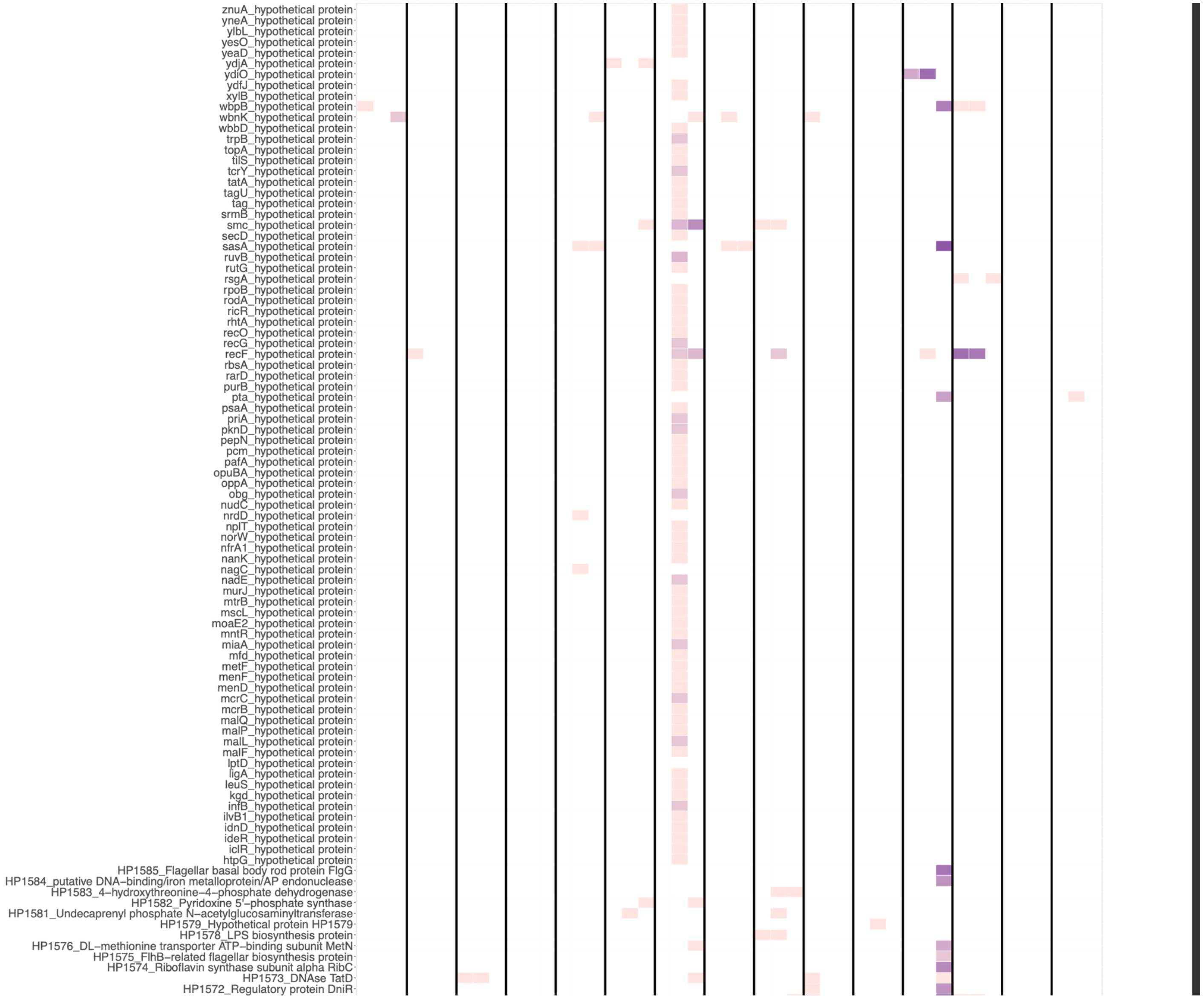
znuA_hypothetical protein
yneA_hypothetical protein
ylbL_hypothetical protein
yesO_hypothetical protein
yeaD_hypothetical protein
ydjA_hypothetical protein
ydiO_hypothetical protein
ydfJ_hypothetical protein
xylB_hypothetical protein
wbpB_hypothetical protein
wbnK_hypothetical protein
wbbD_hypothetical protein
trpB_hypothetical protein
topA_hypothetical protein
tilS_hypothetical protein
tcrY_hypothetical protein
tatA_hypothetical protein
tagU_hypothetical protein
tag_hypothetical protein
srmB_hypothetical protein
smc_hypothetical protein
secD_hypothetical protein
sasA_hypothetical protein
ruvB_hypothetical protein
rutG_hypothetical protein
rsgA_hypothetical protein
rpoB_hypothetical protein
rodA_hypothetical protein
ricR_hypothetical protein
rhtA_hypothetical protein
recO_hypothetical protein
recG_hypothetical protein
recF_hypothetical protein
rbsA_hypothetical protein
rarD_hypothetical protein
purB_hypothetical protein
pta_hypothetical protein
psaA_hypothetical protein
priA_hypothetical protein
pknD_hypothetical protein
pepN_hypothetical protein
pcm_hypothetical protein
pafA_hypothetical protein
opuBA_hypothetical protein
oppA_hypothetical protein
obg_hypothetical protein
nudC_hypothetical protein
nrdD_hypothetical protein
nplT_hypothetical protein
norW_hypothetical protein
nfrA1_hypothetical protein
nanK_hypothetical protein
nagC_hypothetical protein
nadE_hypothetical protein
murJ_hypothetical protein
mtrB_hypothetical protein
mscL_hypothetical protein
moaE2_hypothetical protein
mntR_hypothetical protein
miaA_hypothetical protein
mfd_hypothetical protein
metF_hypothetical protein
menF_hypothetical protein
menD_hypothetical protein
mcrC_hypothetical protein
mcrB_hypothetical protein
malQ_hypothetical protein
malP_hypothetical protein
malL_hypothetical protein
malF_hypothetical protein
lptD_hypothetical protein
ligA_hypothetical protein
leuS_hypothetical protein
kgd_hypothetical protein
infB_hypothetical protein
ilvB1_hypothetical protein
idnD_hypothetical protein
ideR_hypothetical protein
iclR_hypothetical protein
htpG_hypothetical protein
HP1585_Flagellar basal body rod protein FlgG
HP1584_putative DNA−binding/iron metalloprotein/AP endonuclease
HP1583_4−hydroxythreonine−4−phosphate dehydrogenase
HP1582_Pyridoxine 5'−phosphate synthase
HP1581_Undecaprenyl phosphate N−acetylglucosaminyltransferase
HP1579_Hypothetical protein HP1579
HP1578_LPS biosynthesis protein
HP1576_DL−methionine transporter ATP−binding subunit MetN
HP1575_FlhB−related flagellar biosynthesis protein
HP1574_Riboflavin synthase subunit alpha RibC
HP1573_DNAse TatD
HP1572_Regulatory protein DniR

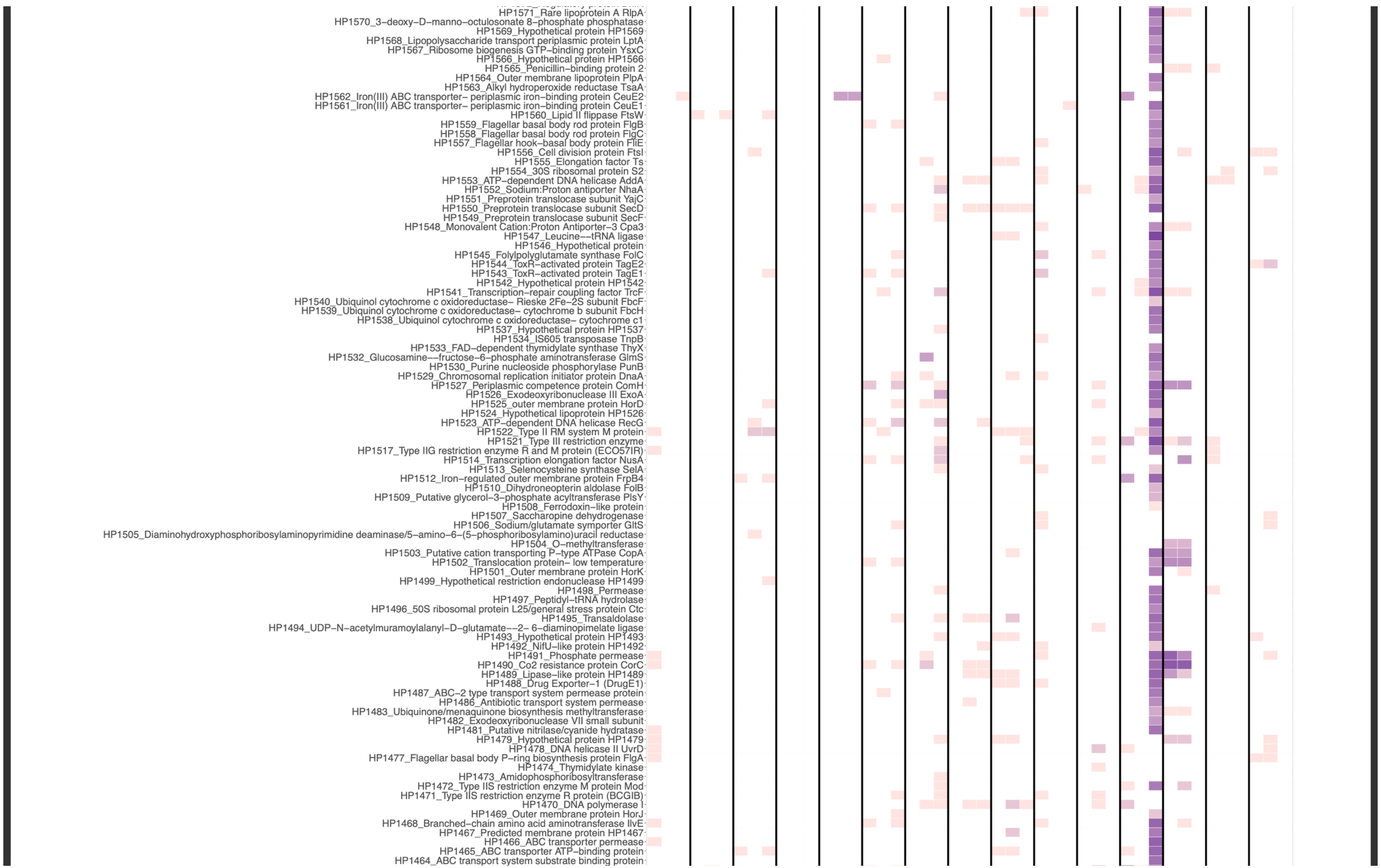
HP1571_Rare lipoprotein A RlpA
HP1570_3-deoxy-D-manno-octulosonate 8-phosphate phosphatase
HP1569_Hypothetical protein HP1569
HP1568_Lipopolysaccharide transport periplasmic protein LptA
HP1567_Ribosome biogenesis GTP-binding protein YsxC
HP1566_Hypothetical protein HP1566
HP1565_Penicillin-binding protein 2
HP1564_Outer membrane lipoprotein PlpA
HP1563_Alkyl hydroperoxide reductase TsaA
HP1562_Iron(III) ABC transporter- periplasmic iron-binding protein CeuE2
HP1561_Iron(III) ABC transporter- periplasmic iron-binding protein CeuE1
HP1560_Lipid II flippase FtsW
HP1559_Flagellar basal body rod protein FlgB
HP1558_Flagellar basal body rod protein FlgC
HP1557_Flagellar hook-basal body protein FliE
HP1556_Cell division protein FtsI
HP1555_Elongation factor Ts
HP1554_30S ribosomal protein S2
HP1553_ATP-dependent DNA helicase AddA
HP1552_Sodium:Proton antiporter NhaA
HP1551_Preprotein translocase subunit YajC
HP1550_Preprotein translocase subunit SecD
HP1549_Preprotein translocase subunit SecF
HP1548_Monovalent Cation:Proton Antiporter-3 Cpa3
HP1547_Leucine--tRNA ligase
HP1546_Hypothetical protein
HP1545_Folylpolyglutamate synthase FolC
HP1544_ToxR-activated protein TagE2
HP1543_ToxR-activated protein TagE1
HP1542_Hypothetical protein HP1542
HP1541_Transcription-repair coupling factor TrcF
HP1540_Ubiquinol cytochrome c oxidoreductase- Rieske 2Fe-2S subunit FbcF
HP1539_Ubiquinol cytochrome c oxidoreductase- cytochrome b subunit FbcH
HP1538_Ubiquinol cytochrome c oxidoreductase- cytochrome c1
HP1537_Hypothetical protein HP1537
HP1534_IS605 transposase TnpB
HP1533_FAD-dependent thymidylate synthase ThyX
HP1532_Glucosamine--fructose-6-phosphate aminotransferase GlmS
HP1530_Purine nucleoside phosphorylase PunB
HP1529_Chromosomal replication initiator protein DnaA
HP1527_Periplasmic competence protein ComH
HP1526_Exodeoxyribonuclease III ExoA
HP1525_outer membrane protein HorD
HP1524_Hypothetical lipoprotein HP1526
HP1523_ATP-dependent DNA helicase RecG
HP1522_Type II RM system M protein
HP1521_Type III restriction enzyme
HP1517_Type IIG restriction enzyme R and M protein (ECO57IR)
HP1514_Transcription elongation factor NusA
HP1513_Selenocysteine synthase SelA
HP1512_Iron-regulated outer membrane protein FrpB4
HP1510_Dihydroneopterin aldolase FolB
HP1509_Putative glycerol-3-phosphate acyltransferase PlsY
HP1508_Ferrodoxin-like protein
HP1507_Saccharopine dehydrogenase
HP1506_Sodium/glutamate symporter GltS
HP1505_Diaminohydroxyphosphoribosylaminopyrimidine deaminase/5-amino-6-(5-phosphoribosylamino)uracil reductase
HP1504_O-methyltransferase
HP1503_Putative cation transporting P-type ATPase CopA
HP1502_Translocation protein- low temperature
HP1501_Outer membrane protein HorK
HP1499_Hypothetical restriction endonuclease HP1499
HP1498_Permease
HP1497_Peptidyl-tRNA hydrolase
HP1496_50S ribosomal protein L25/general stress protein Ctc
HP1495_Transaldolase
HP1494_UDP-N-acetylmuramoylalanyl-D-glutamate--2- 6-diaminopimelate ligase
HP1493_Hypothetical protein HP1493
HP1492_NifU-like protein HP1492
HP1491_Phosphate permease
HP1490_Co2 resistance protein CorC
HP1489_Lipase-like protein HP1489
HP1488_Drug Exporter-1 (DrugE1)
HP1487_ABC-2 type transport system permease protein
HP1486_Antibiotic transport system permease
HP1483_Ubiquinone/menaquinone biosynthesis methyltransferase
HP1482_Exodeoxyribonuclease VII small subunit
HP1481_Putative nitrilase/cyanide hydratase
HP1479_Hypothetical protein HP1479
HP1478_DNA helicase II UvrD
HP1477_Flagellar basal body P-ring biosynthesis protein FlgA
HP1474_Thymidylate kinase
HP1473_Amidophosphoribosyltransferase
HP1472_Type IIS restriction enzyme M protein Mod
HP1471_Type IIS restriction enzyme R protein (BCGIB)
HP1470_DNA polymerase I
HP1469_Outer membrane protein HorJ
HP1468_Branched-chain amino acid aminotransferase IlvE
HP1467_Predicted membrane protein HP1467
HP1466_ABC transporter permease
HP1465_ABC transporter ATP-binding protein
HP1464_ABC transport system substrate binding protein

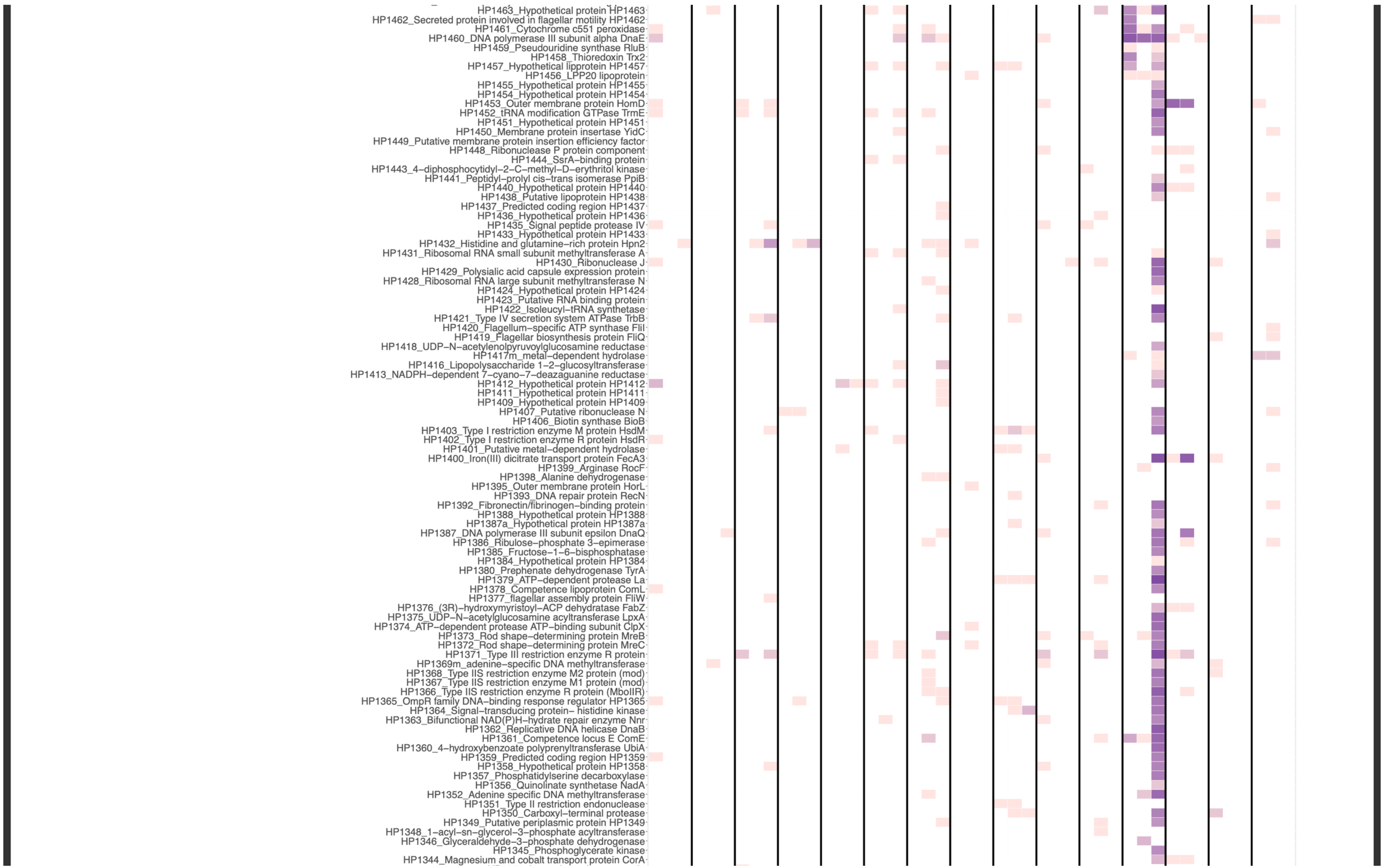
HP1463_Hypothetical protein HP1463
HP1462_Secreted protein involved in flagellar motility HP1462
HP1461_Cytochrome c551 peroxidase
HP1460_DNA polymerase III subunit alpha DnaE
HP1459_Pseudouridine synthase RluB
HP1458_Thioredoxin Trx2
HP1457_Hypothetical lipprotein HP1457
HP1456_LPP20 lipoprotein
HP1455_Hypothetical protein HP1455
HP1454_Hypothetical protein HP1454
HP1453_Outer membrane protein HomD
HP1452_tRNA modification GTPase TrmE
HP1451_Hypothetical protein HP1451
HP1450_Membrane protein insertase YidC
HP1449_Putative membrane protein insertion efficiency factor
HP1448_Ribonuclease P protein component
HP1444_SsrA–binding protein
HP1443_4–diphosphocytidyl–2–C–methyl–D–erythritol kinase
HP1441_Peptidyl–prolyl cis–trans isomerase PpiB
HP1440_Hypothetical protein HP1440
HP1438_Putative lipoprotein HP1438
HP1437_Predicted coding region HP1437
HP1436_Hypothetical protein HP1436
HP1435_Signal peptide protease IV
HP1433_Hypothetical protein HP1433
HP1432_Histidine and glutamine–rich protein Hpn2
HP1431_Ribosomal RNA small subunit methyltransferase A
HP1430_Ribonuclease J
HP1429_Polysialic acid capsule expression protein
HP1428_Ribosomal RNA large subunit methyltransferase N
HP1424_Hypothetical protein HP1424
HP1423_Putative RNA binding protein
HP1422_Isoleucyl–tRNA synthetase
HP1421_Type IV secretion system ATPase TrbB
HP1420_Flagellum–specific ATP synthase FliI
HP1419_Flagellar biosynthesis protein FliQ
HP1418_UDP–N–acetylenolpyruvoylglucosamine reductase
HP1417m_metal–dependent hydrolase
HP1416_Lipopolysaccharide 1–2–glucosyltransferase
HP1413_NADPH–dependent 7–cyano–7–deazaguanine reductase
HP1412_Hypothetical protein HP1412
HP1411_Hypothetical protein HP1411
HP1409_Hypothetical protein HP1409
HP1407_Putative ribonuclease N
HP1406_Biotin synthase BioB
HP1403_Type I restriction enzyme M protein HsdM
HP1402_Type I restriction enzyme R protein HsdR
HP1401_Putative metal–dependent hydrolase
HP1400_Iron(III) dicitrate transport protein FecA3
HP1399_Arginase RocF
HP1398_Alanine dehydrogenase
HP1395_Outer membrane protein HorL
HP1393_DNA repair protein RecN
HP1392_Fibronectin/fibrinogen–binding protein
HP1388_Hypothetical protein HP1388
HP1387a_Hypothetical protein HP1387a
HP1387_DNA polymerase III subunit epsilon DnaQ
HP1386_Ribulose–phosphate 3–epimerase
HP1385_Fructose–1–6–bisphosphatase
HP1384_Hypothetical protein HP1384
HP1380_Prephenate dehydrogenase TyrA
HP1379_ATP–dependent protease La
HP1378_Competence lipoprotein ComL
HP1377_flagellar assembly protein FliW
HP1376_(3R)–hydroxymyristoyl–ACP dehydratase FabZ
HP1375_UDP–N–acetylglucosamine acyltransferase LpxA
HP1374_ATP–dependent protease ATP–binding subunit ClpX
HP1373_Rod shape–determining protein MreB
HP1372_Rod shape–determining protein MreC
HP1371_Type III restriction enzyme R protein
HP1369m_adenine–specific DNA methyltransferase
HP1368_Type IIS restriction enzyme M2 protein (mod)
HP1367_Type IIS restriction enzyme M1 protein (mod)
HP1366_Type IIS restriction enzyme R protein (MboIIR)
HP1365_OmpR family DNA–binding response regulator HP1365
HP1364_Signal–transducing protein– histidine kinase
HP1363_Bifunctional NAD(P)H–hydrate repair enzyme Nnr
HP1362_Replicative DNA helicase DnaB
HP1361_Competence locus E ComE
HP1360_4–hydroxybenzoate polyprenyltransferase UbiA
HP1359_Predicted coding region HP1359
HP1358_Hypothetical protein HP1358
HP1357_Phosphatidylserine decarboxylase
HP1356_Quinolinate synthetase NadA
HP1352_Adenine specific DNA methyltransferase
HP1351_Type II restriction endonuclease
HP1350_Carboxyl–terminal protease
HP1349_Putative periplasmic protein HP1349
HP1348_1–acyl–sn–glycerol–3–phosphate acyltransferase
HP1346_Glyceraldehyde–3–phosphate dehydrogenase
HP1345_Phosphoglycerate kinase
HP1344_Magnesium and cobalt transport protein CorA

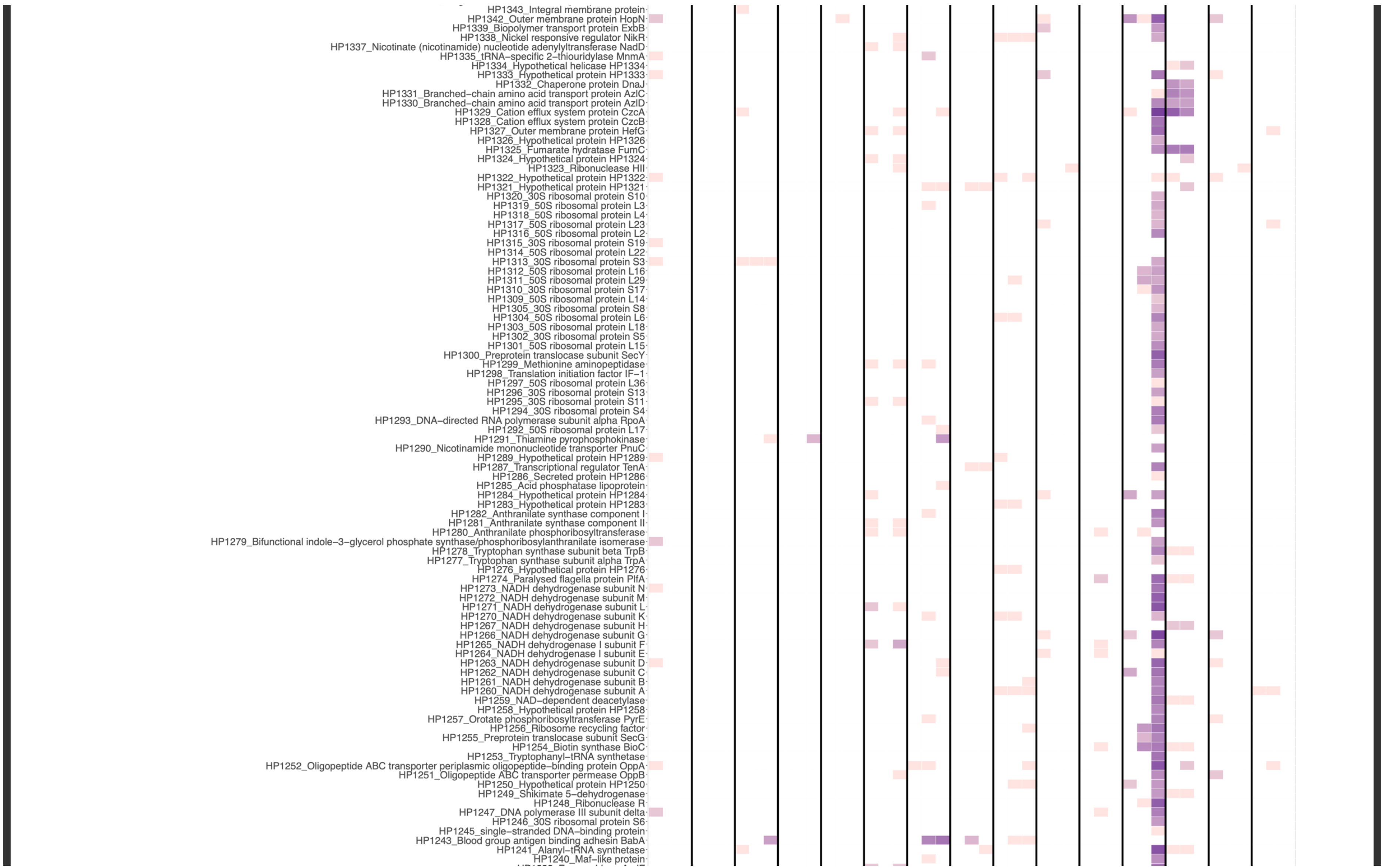
HP1343_Integral membrane protein
HP1342_Outer membrane protein HopN
HP1339_Biopolymer transport protein ExbB
HP1338_Nickel responsive regulator NikR
HP1337_Nicotinate (nicotinamide) nucleotide adenylyltransferase NadD
HP1335_tRNA-specific 2-thiouridylase MnmA
HP1334_Hypothetical helicase HP1334
HP1333_Hypothetical protein HP1333
HP1332_Chaperone protein DnaJ
HP1331_Branched-chain amino acid transport protein AzlC
HP1330_Branched-chain amino acid transport protein AzlD
HP1329_Cation efflux system protein CzcA
HP1328_Cation efflux system protein CzcB
HP1327_Outer membrane protein HefG
HP1326_Hypothetical protein HP1326
HP1325_Fumarate hydratase FumC
HP1324_Hypothetical protein HP1324
HP1323_Ribonuclease HII
HP1322_Hypothetical protein HP1322
HP1321_Hypothetical protein HP1321
HP1320_30S ribosomal protein S10
HP1319_50S ribosomal protein L3
HP1318_50S ribosomal protein L4
HP1317_50S ribosomal protein L23
HP1316_50S ribosomal protein L2
HP1315_30S ribosomal protein S19
HP1314_50S ribosomal protein L22
HP1313_30S ribosomal protein S3
HP1312_50S ribosomal protein L16
HP1311_50S ribosomal protein L29
HP1310_30S ribosomal protein S17
HP1309_50S ribosomal protein L14
HP1305_30S ribosomal protein S8
HP1304_50S ribosomal protein L6
HP1303_50S ribosomal protein L18
HP1302_30S ribosomal protein S5
HP1301_50S ribosomal protein L15
HP1300_Preprotein translocase subunit SecY
HP1299_Methionine aminopeptidase
HP1298_Translation initiation factor IF-1
HP1297_50S ribosomal protein L36
HP1296_30S ribosomal protein S13
HP1295_30S ribosomal protein S11
HP1294_30S ribosomal protein S4
HP1293_DNA-directed RNA polymerase subunit alpha RpoA
HP1292_50S ribosomal protein L17
HP1291_Thiamine pyrophosphokinase
HP1290_Nicotinamide mononucleotide transporter PnuC
HP1289_Hypothetical protein HP1289
HP1287_Transcriptional regulator TenA
HP1286_Secreted protein HP1286
HP1285_Acid phosphatase lipoprotein
HP1284_Hypothetical protein HP1284
HP1283_Hypothetical protein HP1283
HP1282_Anthranilate synthase component I
HP1281_Anthranilate synthase component II
HP1280_Anthranilate phosphoribosyltransferase
HP1279_Bifunctional indole-3-glycerol phosphate synthase/phosphoribosylanthranilate isomerase
HP1278_Tryptophan synthase subunit beta TrpB
HP1277_Tryptophan synthase subunit alpha TrpA
HP1276_Hypothetical protein HP1276
HP1274_Paralysed flagella protein PflA
HP1273_NADH dehydrogenase subunit N
HP1272_NADH dehydrogenase subunit M
HP1271_NADH dehydrogenase subunit L
HP1270_NADH dehydrogenase subunit K
HP1267_NADH dehydrogenase subunit H
HP1266_NADH dehydrogenase subunit G
HP1265_NADH dehydrogenase I subunit F
HP1264_NADH dehydrogenase I subunit E
HP1263_NADH dehydrogenase subunit D
HP1262_NADH dehydrogenase subunit C
HP1261_NADH dehydrogenase subunit B
HP1260_NADH dehydrogenase subunit A
HP1259_NAD-dependent deacetylase
HP1258_Hypothetical protein HP1258
HP1257_Orotate phosphoribosyltransferase PyrE
HP1256_Ribosome recycling factor
HP1255_Preprotein translocase subunit SecG
HP1254_Biotin synthase BioC
HP1253_Tryptophanyl-tRNA synthetase
HP1252_Oligopeptide ABC transporter periplasmic oligopeptide-binding protein OppA
HP1251_Oligopeptide ABC transporter permease OppB
HP1250_Hypothetical protein HP1250
HP1249_Shikimate 5-dehydrogenase
HP1248_Ribonuclease R
HP1247_DNA polymerase III subunit delta
HP1246_30S ribosomal protein S6
HP1245_single-stranded DNA-binding protein
HP1243_Blood group antigen binding adhesin BabA
HP1241_Alanyl-tRNA synthetase
HP1240_Maf-like protein

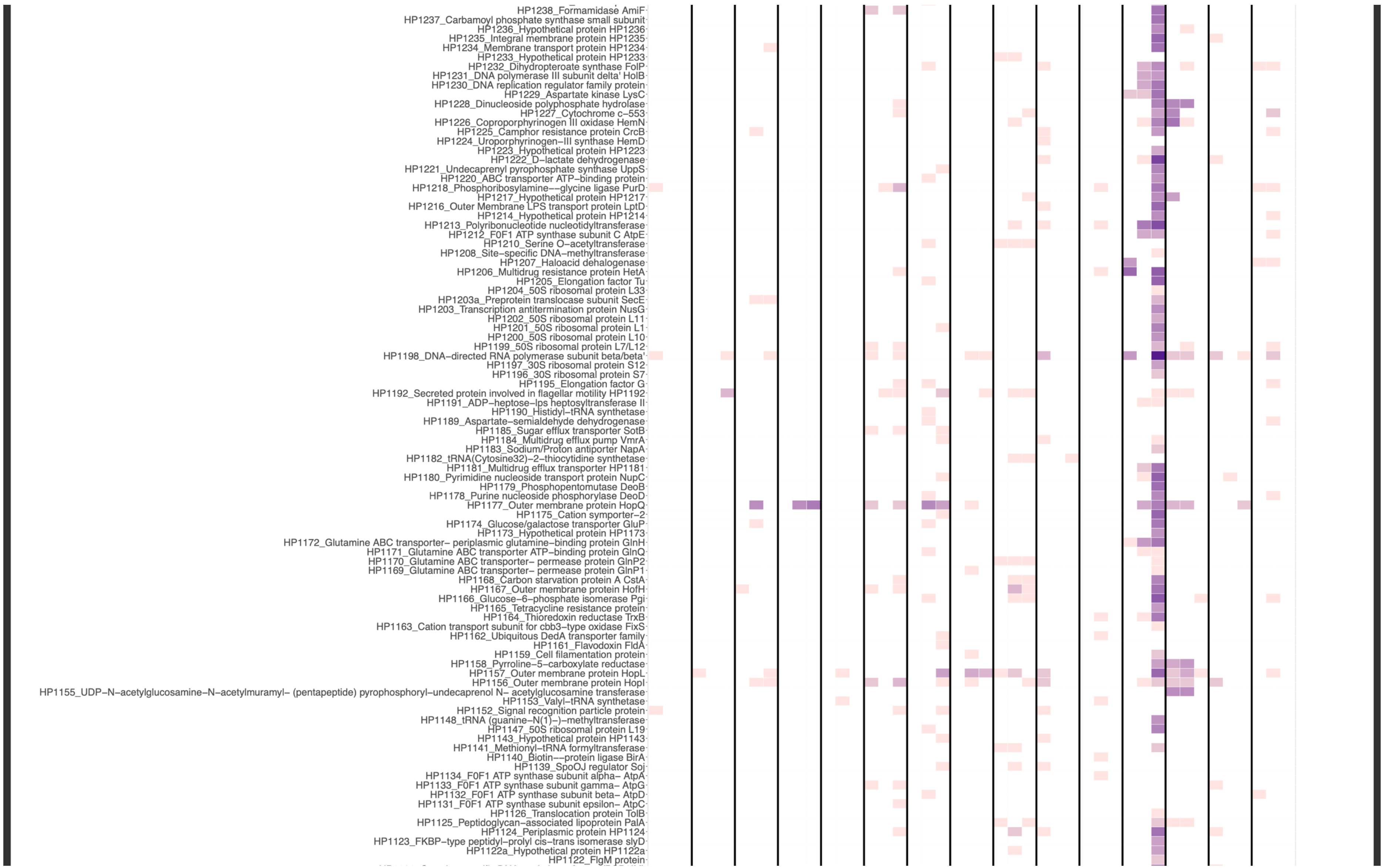
HP1238_Formamidase AmiF
HP1237_Carbamoyl phosphate synthase small subunit
HP1236_Hypothetical protein HP1236
HP1235_Integral membrane protein HP1235
HP1234_Membrane transport protein HP1234
HP1233_Hypothetical protein HP1233
HP1232_Dihydropteroate synthase FolP
HP1231_DNA polymerase III subunit delta' HolB
HP1230_DNA replication regulator family protein
HP1229_Aspartate kinase LysC
HP1228_Dinucleoside polyphosphate hydrolase
HP1227_Cytochrome c−553
HP1226_Coproporphyrinogen III oxidase HemN
HP1225_Camphor resistance protein CrcB
HP1224_Uroporphyrinogen−III synthase HemD
HP1223_Hypothetical protein HP1223
HP1222_D−lactate dehydrogenase
HP1221_Undecaprenyl pyrophosphate synthase UppS
HP1220_ABC transporter ATP−binding protein
HP1218_Phosphoribosylamine−−glycine ligase PurD
HP1217_Hypothetical protein HP1217
HP1216_Outer Membrane LPS transport protein LptD
HP1214_Hypothetical protein HP1214
HP1213_Polyribonucleotide nucleotidyltransferase
HP1212_F0F1 ATP synthase subunit C AtpE
HP1210_Serine O−acetyltransferase
HP1208_Site−specific DNA−methyltransferase
HP1207_Haloacid dehalogenase
HP1206_Multidrug resistance protein HetA
HP1205_Elongation factor Tu
HP1204_50S ribosomal protein L33
HP1203a_Preprotein translocase subunit SecE
HP1203_Transcription antitermination protein NusG
HP1202_50S ribosomal protein L11
HP1201_50S ribosomal protein L1
HP1200_50S ribosomal protein L10
HP1199_50S ribosomal protein L7/L12
HP1198_DNA−directed RNA polymerase subunit beta/beta'
HP1197_30S ribosomal protein S12
HP1196_30S ribosomal protein S7
HP1195_Elongation factor G
HP1192_Secreted protein involved in flagellar motility HP1192
HP1191_ADP−heptose−lps heptosyltransferase II
HP1190_Histidyl−tRNA synthetase
HP1189_Aspartate−semialdehyde dehydrogenase
HP1185_Sugar efflux transporter SotB
HP1184_Multidrug efflux pump VmrA
HP1183_Sodium/Proton antiporter NapA
HP1182_tRNA(Cytosine32)−2−thiocytidine synthetase
HP1181_Multidrug efflux transporter HP1181
HP1180_Pyrimidine nucleoside transport protein NupC
HP1179_Phosphopentomutase DeoB
HP1178_Purine nucleoside phosphorylase DeoD
HP1177_Outer membrane protein HopQ
HP1175_Cation symporter−2
HP1174_Glucose/galactose transporter GluP
HP1173_Hypothetical protein HP1173
HP1172_Glutamine ABC transporter− periplasmic glutamine−binding protein GlnH
HP1171_Glutamine ABC transporter ATP−binding protein GlnQ
HP1170_Glutamine ABC transporter− permease protein GlnP2
HP1169_Glutamine ABC transporter− permease protein GlnP1
HP1168_Carbon starvation protein A CstA
HP1167_Outer membrane protein HofH
HP1166_Glucose−6−phosphate isomerase Pgi
HP1165_Tetracycline resistance protein
HP1164_Thioredoxin reductase TrxB
HP1163_Cation transport subunit for cbb3−type oxidase FixS
HP1162_Ubiquitous DedA transporter family
HP1161_Flavodoxin FldA
HP1159_Cell filamentation protein
HP1158_Pyrroline−5−carboxylate reductase
HP1157_Outer membrane protein HopL
HP1156_Outer membrane protein HopI
HP1155_UDP−N−acetylglucosamine−N−acetylmuramyl− (pentapeptide) pyrophosphoryl−undecaprenol N− acetylglucosamine transferase
HP1153_Valyl−tRNA synthetase
HP1152_Signal recognition particle protein
HP1148_tRNA (guanine−N(1)−)−methyltransferase
HP1147_50S ribosomal protein L19
HP1143_Hypothetical protein HP1143
HP1141_Methionyl−tRNA formyltransferase
HP1140_Biotin−−protein ligase BirA
HP1139_SpoOJ regulator Soj
HP1134_F0F1 ATP synthase subunit alpha− AtpA
HP1133_F0F1 ATP synthase subunit gamma− AtpG
HP1132_F0F1 ATP synthase subunit beta− AtpD
HP1131_F0F1 ATP synthase subunit epsilon− AtpC
HP1126_Translocation protein TolB
HP1125_Peptidoglycan−associated lipoprotein PalA
HP1124_Periplasmic protein HP1124
HP1123_FKBP−type peptidyl−prolyl cis−trans isomerase slyD
HP1122a_Hypothetical protein HP1122a
HP1122_FlgM protein

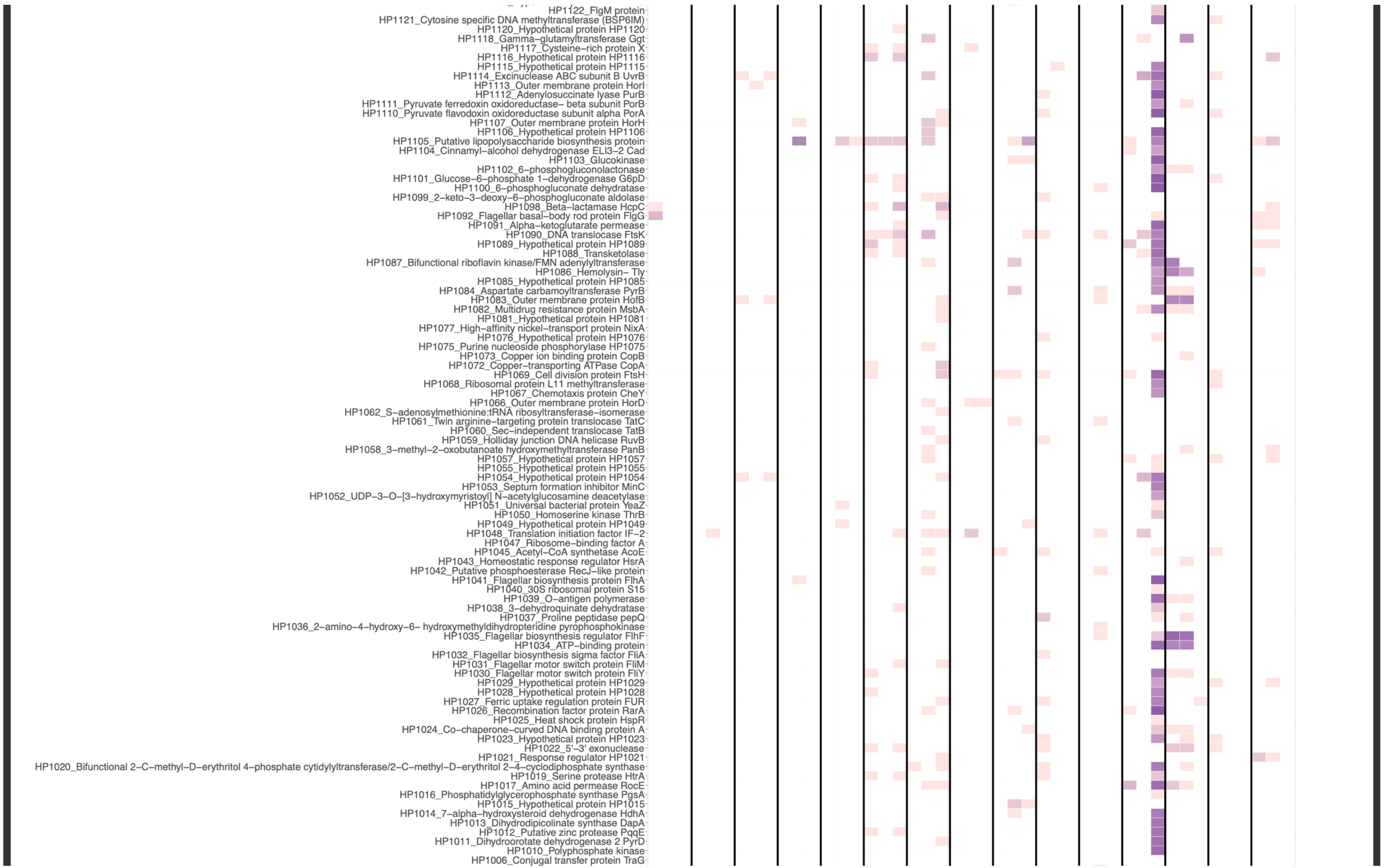
HP1122_FlgM protein
HP1121_Cytosine specific DNA methyltransferase (BSP6IM)
HP1120_Hypothetical protein HP1120
HP1118_Gamma–glutamyltransferase Ggt
HP1117_Cysteine–rich protein X
HP1116_Hypothetical protein HP1116
HP1115_Hypothetical protein HP1115
HP1114_Excinuclease ABC subunit B UvrB
HP1113_Outer membrane protein HorI
HP1112_Adenylosuccinate lyase PurB
HP1111_Pyruvate ferredoxin oxidoreductase– beta subunit PorB
HP1110_Pyruvate flavodoxin oxidoreductase subunit alpha PorA
HP1107_Outer membrane protein HorH
HP1106_Hypothetical protein HP1106
HP1105_Putative lipopolysaccharide biosynthesis protein
HP1104_Cinnamyl–alcohol dehydrogenase ELI3–2 Cad
HP1103_Glucokinase
HP1102_6–phosphogluconolactonase
HP1101_Glucose–6–phosphate 1–dehydrogenase G6pD
HP1100_6–phosphogluconate dehydratase
HP1099_2–keto–3–deoxy–6–phosphogluconate aldolase
HP1098_Beta–lactamase HcpC
HP1092_Flagellar basal–body rod protein FlgG
HP1091_Alpha–ketoglutarate permease
HP1090_DNA translocase FtsK
HP1089_Hypothetical protein HP1089
HP1088_Transketolase
HP1087_Bifunctional riboflavin kinase/FMN adenylyltransferase
HP1086_Hemolysin– Tly
HP1085_Hypothetical protein HP1085
HP1084_Aspartate carbamoyltransferase PyrB
HP1083_Outer membrane protein HofB
HP1082_Multidrug resistance protein MsbA
HP1081_Hypothetical protein HP1081
HP1077_High–affinity nickel–transport protein NixA
HP1076_Hypothetical protein HP1076
HP1075_Purine nucleoside phosphorylase HP1075
HP1073_Copper ion binding protein CopB
HP1072_Copper–transporting ATPase CopA
HP1069_Cell division protein FtsH
HP1068_Ribosomal protein L11 methyltransferase
HP1067_Chemotaxis protein CheY
HP1066_Outer membrane protein HorD
HP1062_S–adenosylmethionine:tRNA ribosyltransferase–isomerase
HP1061_Twin arginine–targeting protein translocase TatC
HP1060_Sec–independent translocase TatB
HP1059_Holliday junction DNA helicase RuvB
HP1058_3–methyl–2–oxobutanoate hydroxymethyltransferase PanB
HP1057_Hypothetical protein HP1057
HP1055_Hypothetical protein HP1055
HP1054_Hypothetical protein HP1054
HP1053_Septum formation inhibitor MinC
HP1052_UDP–3–O–[3–hydroxymyristoyl] N–acetylglucosamine deacetylase
HP1051_Universal bacterial protein YeaZ
HP1050_Homoserine kinase ThrB
HP1049_Hypothetical protein HP1049
HP1048_Translation initiation factor IF–2
HP1047_Ribosome–binding factor A
HP1045_Acetyl–CoA synthetase AcoE
HP1043_Homeostatic response regulator HsrA
HP1042_Putative phosphoesterase RecJ–like protein
HP1041_Flagellar biosynthesis protein FlhA
HP1040_30S ribosomal protein S15
HP1039_O–antigen polymerase
HP1038_3–dehydroquinate dehydratase
HP1037_Proline peptidase pepQ
HP1036_2–amino–4–hydroxy–6– hydroxymethyldihydropteridine pyrophosphokinase
HP1035_Flagellar biosynthesis regulator FlhF
HP1034_ATP–binding protein
HP1032_Flagellar biosynthesis sigma factor FliA
HP1031_Flagellar motor switch protein FliM
HP1030_Flagellar motor switch protein FliY
HP1029_Hypothetical protein HP1029
HP1028_Hypothetical protein HP1028
HP1027_Ferric uptake regulation protein FUR
HP1026_Recombination factor protein RarA
HP1025_Heat shock protein HspR
HP1024_Co–chaperone–curved DNA binding protein A
HP1023_Hypothetical protein HP1023
HP1022_5'–3' exonuclease
HP1021_Response regulator HP1021
HP1020_Bifunctional 2–C–methyl–D–erythritol 4–phosphate cytidylyltransferase/2–C–methyl–D–erythritol 2–4–cyclodiphosphate synthase
HP1019_Serine protease HtrA
HP1017_Amino acid permease RocE
HP1016_Phosphatidylglycerophosphate synthase PgsA
HP1015_Hypothetical protein HP1015
HP1014_7–alpha–hydroxysteroid dehydrogenase HdhA
HP1013_Dihydrodipicolinate synthase DapA
HP1012_Putative zinc protease PqqE
HP1011_Dihydroorotate dehydrogenase 2 PyrD
HP1010_Polyphosphate kinase
HP1006_Conjugal transfer protein TraG

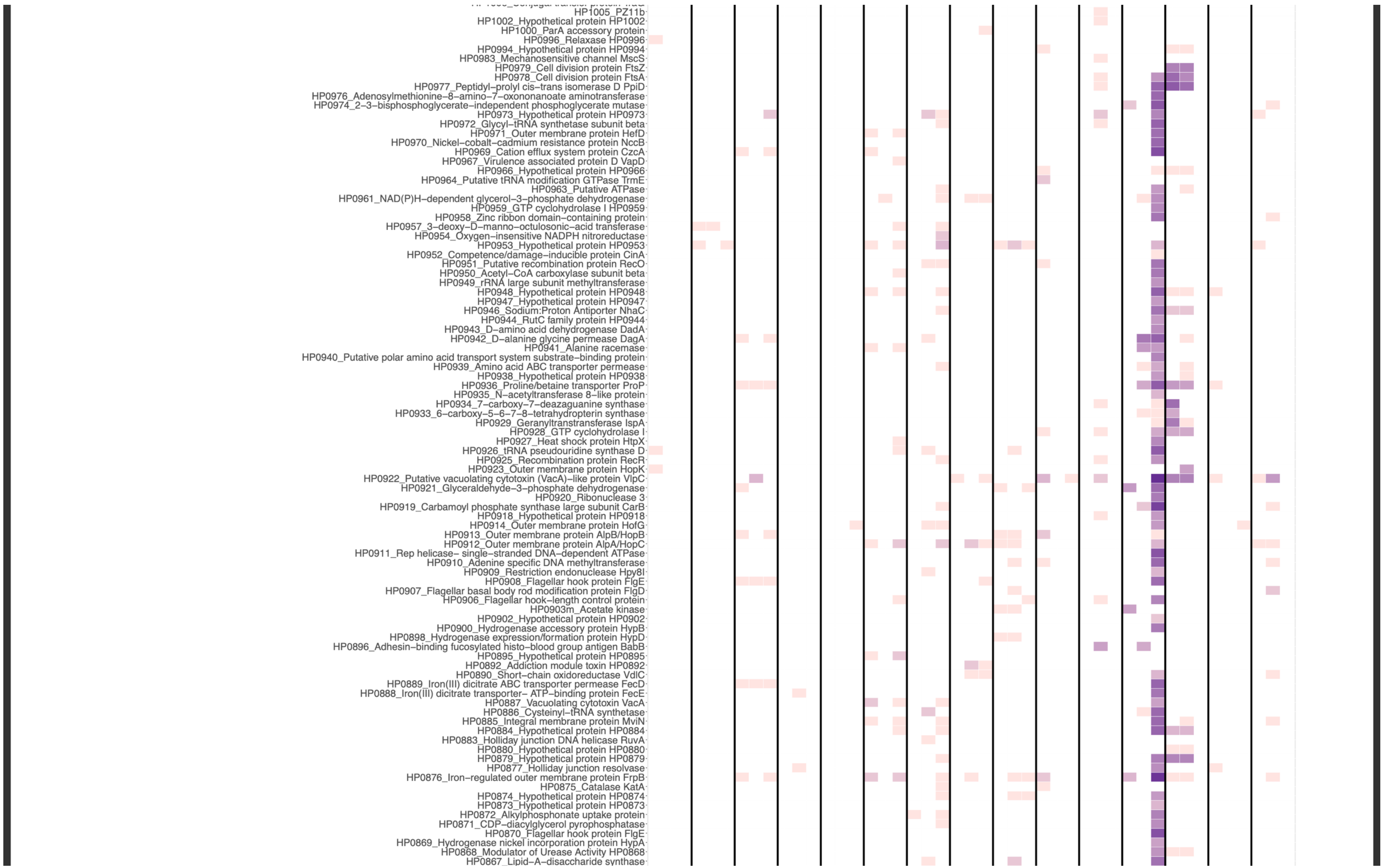
HP1005_PZ11b
HP1002_Hypothetical protein HP1002
HP1000_ParA accessory protein
HP0996_Relaxase HP0996
HP0994_Hypothetical protein HP0994
HP0983_Mechanosensitive channel MscS
HP0979_Cell division protein FtsZ
HP0978_Cell division protein FtsA
HP0977_Peptidyl–prolyl cis–trans isomerase D PpiD
HP0976_Adenosylmethionine–8–amino–7–oxononanoate aminotransferase
HP0974_2–3–bisphosphoglycerate–independent phosphoglycerate mutase
HP0973_Hypothetical protein HP0973
HP0972_Glycyl–tRNA synthetase subunit beta
HP0971_Outer membrane protein HefD
HP0970_Nickel–cobalt–cadmium resistance protein NccB
HP0969_Cation efflux system protein CzcA
HP0967_Virulence associated protein D VapD
HP0966_Hypothetical protein HP0966
HP0964_Putative tRNA modification GTPase TrmE
HP0963_Putative ATPase
HP0961_NAD(P)H–dependent glycerol–3–phosphate dehydrogenase
HP0959_GTP cyclohydrolase I HP0959
HP0958_Zinc ribbon domain–containing protein
HP0957_3–deoxy–D–manno–octulosonic–acid transferase
HP0954_Oxygen–insensitive NADPH nitroreductase
HP0953_Hypothetical protein HP0953
HP0952_Competence/damage–inducible protein CinA
HP0951_Putative recombination protein RecO
HP0950_Acetyl–CoA carboxylase subunit beta
HP0949_rRNA large subunit methyltransferase
HP0948_Hypothetical protein HP0948
HP0947_Hypothetical protein HP0947
HP0946_Sodium:Proton Antiporter NhaC
HP0944_RutC family protein HP0944
HP0943_D–amino acid dehydrogenase DadA
HP0942_D–alanine glycine permease DagA
HP0941_Alanine racemase
HP0940_Putative polar amino acid transport system substrate–binding protein
HP0939_Amino acid ABC transporter permease
HP0938_Hypothetical protein HP0938
HP0936_Proline/betaine transporter ProP
HP0935_N–acetyltransferase 8–like protein
HP0934_7–carboxy–7–deazaguanine synthase
HP0933_6–carboxy–5–6–7–8–tetrahydropterin synthase
HP0929_Geranyltranstransferase IspA
HP0928_GTP cyclohydrolase I
HP0927_Heat shock protein HtpX
HP0926_tRNA pseudouridine synthase D
HP0925_Recombination protein RecR
HP0923_Outer membrane protein HopK
HP0922_Putative vacuolating cytotoxin (VacA)–like protein VlpC
HP0921_Glyceraldehyde–3–phosphate dehydrogenase
HP0920_Ribonuclease 3
HP0919_Carbamoyl phosphate synthase large subunit CarB
HP0918_Hypothetical protein HP0918
HP0914_Outer membrane protein HofG
HP0913_Outer membrane protein AlpB/HopB
HP0912_Outer membrane protein AlpA/HopC
HP0911_Rep helicase– single–stranded DNA–dependent ATPase
HP0910_Adenine specific DNA methyltransferase
HP0909_Restriction endonuclease Hpy8I
HP0908_Flagellar hook protein FlgE
HP0907_Flagellar basal body rod modification protein FlgD
HP0906_Flagellar hook–length control protein
HP0903m_Acetate kinase
HP0902_Hypothetical protein HP0902
HP0900_Hydrogenase accessory protein HypB
HP0898_Hydrogenase expression/formation protein HypD
HP0896_Adhesin–binding fucosylated histo–blood group antigen BabB
HP0895_Hypothetical protein HP0895
HP0892_Addiction module toxin HP0892
HP0890_Short–chain oxidoreductase VdlC
HP0889_Iron(III) dicitrate ABC transporter permease FecD
HP0888_Iron(III) dicitrate transporter– ATP–binding protein FecE
HP0887_Vacuolating cytotoxin VacA
HP0886_Cysteinyl–tRNA synthetase
HP0885_Integral membrane protein MviN
HP0884_Hypothetical protein HP0884
HP0883_Holliday junction DNA helicase RuvA
HP0880_Hypothetical protein HP0880
HP0879_Hypothetical protein HP0879
HP0877_Holliday junction resolvase
HP0876_Iron–regulated outer membrane protein FrpB
HP0875_Catalase KatA
HP0874_Hypothetical protein HP0874
HP0873_Hypothetical protein HP0873
HP0872_Alkylphosphonate uptake protein
HP0871_CDP–diacylglycerol pyrophosphatase
HP0870_Flagellar hook protein FlgE
HP0869_Hydrogenase nickel incorporation protein HypA
HP0868_Modulator of Urease Activity HP0868
HP0867_Lipid–A–disaccharide synthase

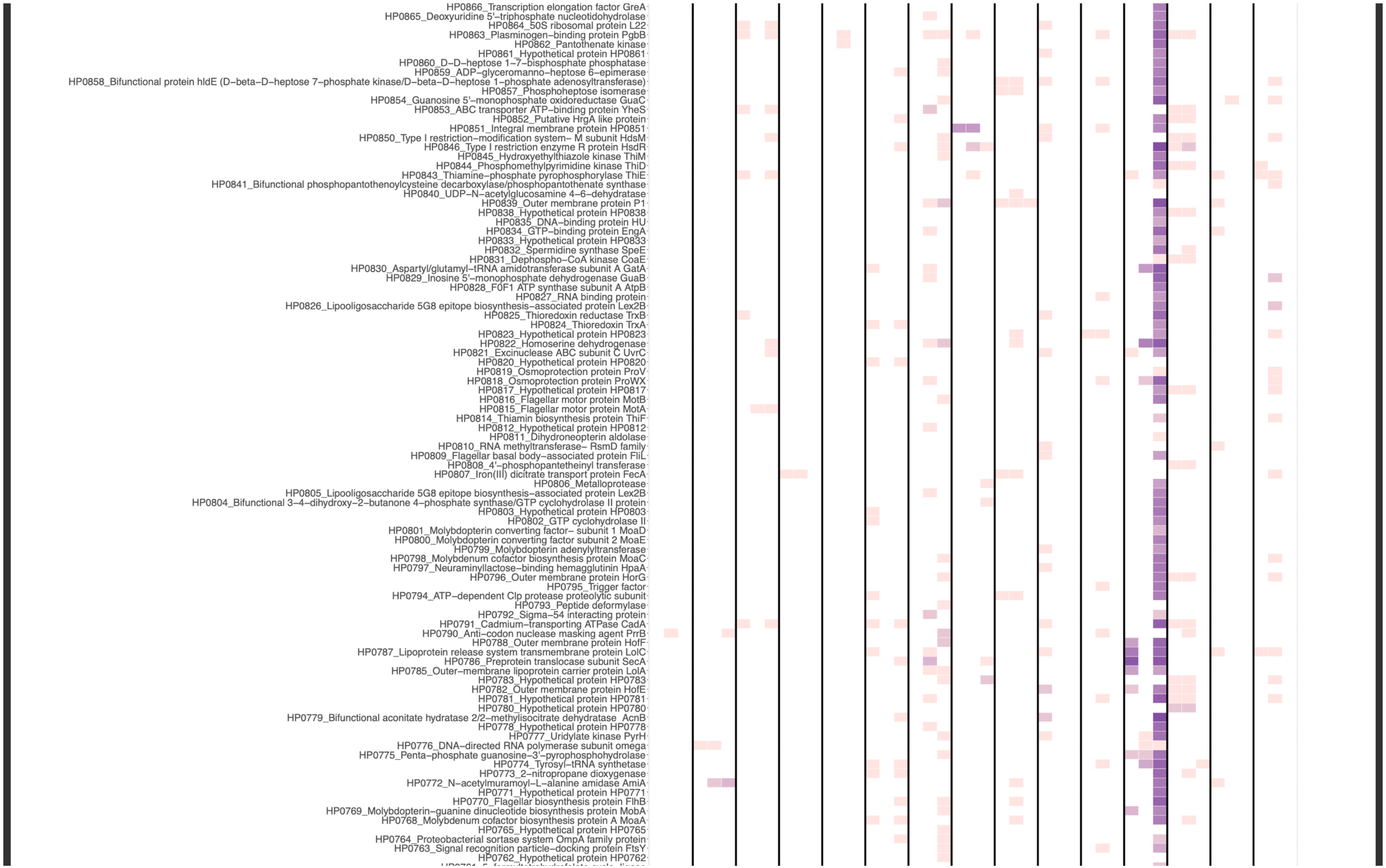

HP0866_Transcription elongation factor GreA
HP0865_Deoxyuridine 5'–triphosphate nucleotidohydrolase
HP0864_50S ribosomal protein L22
HP0863_Plasminogen–binding protein PgbB
HP0862_Pantothenate kinase
HP0861_Hypothetical protein HP0861
HP0860_D–D–heptose 1–7–bisphosphate phosphatase
HP0859_ADP–glyceromanno–heptose 6–epimerase
HP0858_Bifunctional protein hldE (D–beta–D–heptose 7–phosphate kinase/D–beta–D–heptose 1–phosphate adenosyltransferase)
HP0857_Phosphoheptose isomerase
HP0854_Guanosine 5'–monophosphate oxidoreductase GuaC
HP0853_ABC transporter ATP–binding protein YheS
HP0852_Putative HrgA like protein
HP0851_Integral membrane protein HP0851
HP0850_Type I restriction–modification system– M subunit HdsM
HP0846_Type I restriction enzyme R protein HsdR
HP0845_Hydroxyethylthiazole kinase ThiM
HP0844_Phosphomethylpyrimidine kinase ThiD
HP0843_Thiamine–phosphate pyrophosphorylase ThiE
HP0841_Bifunctional phosphopantothenoylcysteine decarboxylase/phosphopantothenate synthase
HP0840_UDP–N–acetylglucosamine 4–6–dehydratase
HP0839_Outer membrane protein P1
HP0838_Hypothetical protein HP0838
HP0835_DNA–binding protein HU
HP0834_GTP–binding protein EngA
HP0833_Hypothetical protein HP0833
HP0832_Spermidine synthase SpeE
HP0831_Dephospho–CoA kinase CoaE
HP0830_Aspartyl/glutamyl–tRNA amidotransferase subunit A GatA
HP0829_Inosine 5'–monophosphate dehydrogenase GuaB
HP0828_F0F1 ATP synthase subunit A AtpB
HP0827_RNA binding protein
HP0826_Lipooligosaccharide 5G8 epitope biosynthesis–associated protein Lex2B
HP0825_Thioredoxin reductase TrxB
HP0824_Thioredoxin TrxA
HP0823_Hypothetical protein HP0823
HP0822_Homoserine dehydrogenase
HP0821_Excinuclease ABC subunit C UvrC
HP0820_Hypothetical protein HP0820
HP0819_Osmoprotection protein ProV
HP0818_Osmoprotection protein ProWX
HP0817_Hypothetical protein HP0817
HP0816_Flagellar motor protein MotB
HP0815_Flagellar motor protein MotA
HP0814_Thiamin biosynthesis protein ThiF
HP0812_Hypothetical protein HP0812
HP0811_Dihydroneopterin aldolase
HP0810_RNA methyltransferase– RsmD family
HP0809_Flagellar basal body–associated protein FliL
HP0808_4'–phosphopantetheinyl transferase
HP0807_Iron(III) dicitrate transport protein FecA
HP0806_Metalloprotease
HP0805_Lipooligosaccharide 5G8 epitope biosynthesis–associated protein Lex2B
HP0804_Bifunctional 3–4–dihydroxy–2–butanone 4–phosphate synthase/GTP cyclohydrolase II protein
HP0803_Hypothetical protein HP0803
HP0802_GTP cyclohydrolase II
HP0801_Molybdopterin converting factor– subunit 1 MoaD
HP0800_Molybdopterin converting factor subunit 2 MoaE
HP0799_Molybdopterin adenylyltransferase
HP0798_Molybdenum cofactor biosynthesis protein MoaC
HP0797_Neuraminyllactose–binding hemagglutinin HpaA
HP0796_Outer membrane protein HorG
HP0795_Trigger factor
HP0794_ATP–dependent Clp protease proteolytic subunit
HP0793_Peptide deformylase
HP0792_Sigma–54 interacting protein
HP0791_Cadmium–transporting ATPase CadA
HP0790_Anti–codon nuclease masking agent PrrB
HP0788_Outer membrane protein HofF
HP0787_Lipoprotein release system transmembrane protein LolC
HP0786_Preprotein translocase subunit SecA
HP0785_Outer–membrane lipoprotein carrier protein LolA
HP0783_Hypothetical protein HP0783
HP0782_Outer membrane protein HofE
HP0781_Hypothetical protein HP0781
HP0780_Hypothetical protein HP0780
HP0779_Bifunctional aconitate hydratase 2/2–methylisocitrate dehydratase AcnB
HP0778_Hypothetical protein HP0778
HP0777_Uridylate kinase PyrH
HP0776_DNA–directed RNA polymerase subunit omega
HP0775_Penta–phosphate guanosine–3'–pyrophosphohydrolase
HP0774_Tyrosyl–tRNA synthetase
HP0773_2–nitropropane dioxygenase
HP0772_N–acetylmuramoyl–L–alanine amidase AmiA
HP0771_Hypothetical protein HP0771
HP0770_Flagellar biosynthesis protein FlhB
HP0769_Molybdopterin–guanine dinucleotide biosynthesis protein MobA
HP0768_Molybdenum cofactor biosynthesis protein A MoaA
HP0765_Hypothetical protein HP0765
HP0764_Proteobacterial sortase system OmpA family protein
HP0763_Signal recognition particle–docking protein FtsY
HP0762_Hypothetical protein HP0762

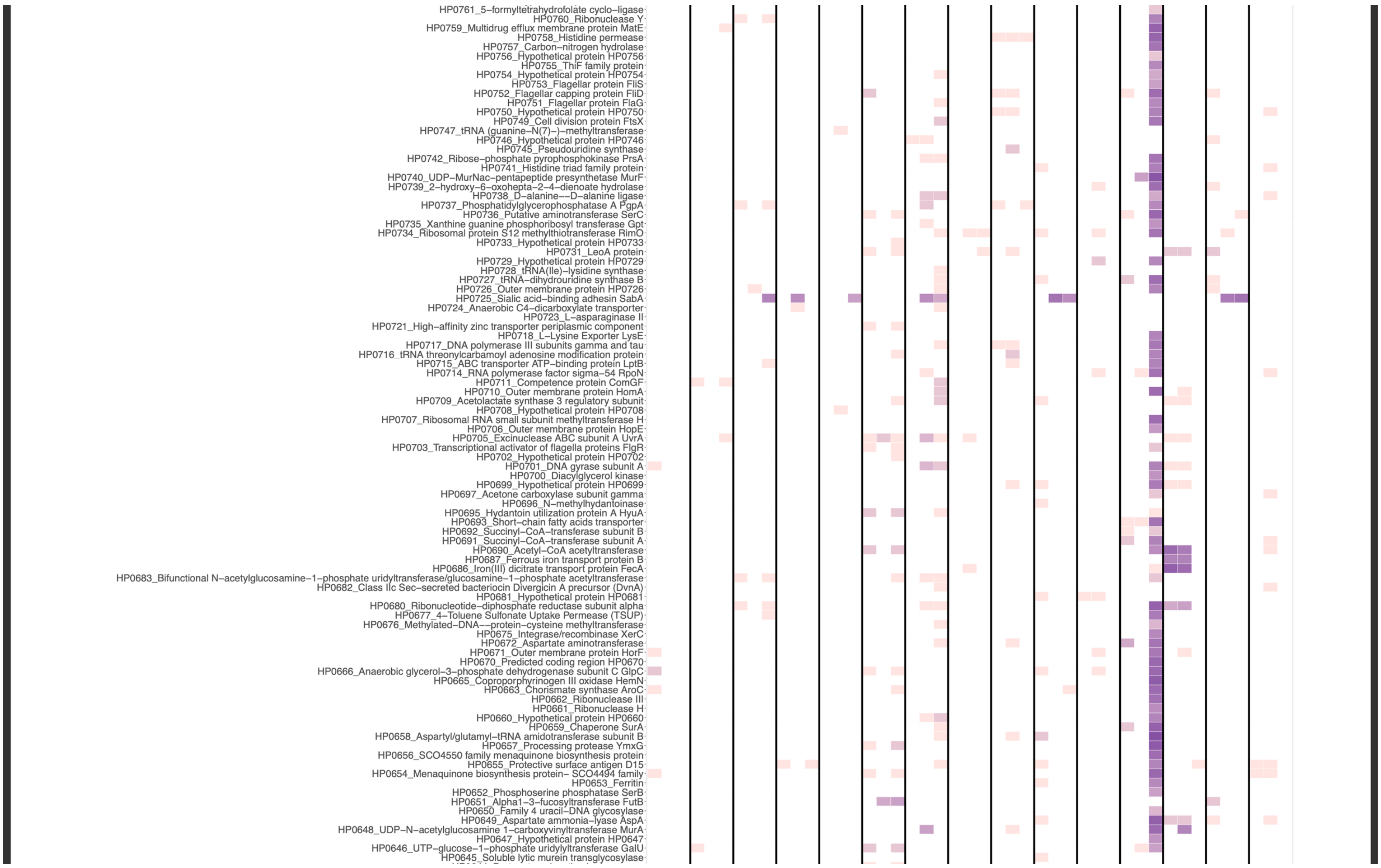
HP0761_5-formyltetrahydrofolate cyclo-ligase
HP0760_Ribonuclease Y
HP0759_Multidrug efflux membrane protein MatE
HP0758_Histidine permease
HP0757_Carbon-nitrogen hydrolase
HP0756_Hypothetical protein HP0756
HP0755_ThiF family protein
HP0754_Hypothetical protein HP0754
HP0753_Flagellar protein FliS
HP0752_Flagellar capping protein FliD
HP0751_Flagellar protein FlaG
HP0750_Hypothetical protein HP0750
HP0749_Cell division protein FtsX
HP0747_tRNA (guanine-N(7)-)-methyltransferase
HP0746_Hypothetical protein HP0746
HP0745_Pseudouridine synthase
HP0742_Ribose-phosphate pyrophosphokinase PrsA
HP0741_Histidine triad family protein
HP0740_UDP-MurNac-pentapeptide presynthetase MurF
HP0739_2-hydroxy-6-oxohepta-2-4-dienoate hydrolase
HP0738_D-alanine--D-alanine ligase
HP0737_Phosphatidylglycerophosphatase A PgpA
HP0736_Putative aminotransferase SerC
HP0735_Xanthine guanine phosphoribosyl transferase Gpt
HP0734_Ribosomal protein S12 methylthiotransferase RimO
HP0733_Hypothetical protein HP0733
HP0731_LeoA protein
HP0729_Hypothetical protein HP0729
HP0728_tRNA(Ile)-lysidine synthase
HP0727_tRNA-dihydrouridine synthase B
HP0726_Outer membrane protein HP0726
HP0725_Sialic acid-binding adhesin SabA
HP0724_Anaerobic C4-dicarboxylate transporter
HP0723_L-asparaginase II
HP0721_High-affinity zinc transporter periplasmic component
HP0718_L-Lysine Exporter LysE
HP0717_DNA polymerase III subunits gamma and tau
HP0716_tRNA threonylcarbamoyl adenosine modification protein
HP0715_ABC transporter ATP-binding protein LptB
HP0714_RNA polymerase factor sigma-54 RpoN
HP0711_Competence protein ComGF
HP0710_Outer membrane protein HomA
HP0709_Acetolactate synthase 3 regulatory subunit
HP0708_Hypothetical protein HP0708
HP0707_Ribosomal RNA small subunit methyltransferase H
HP0706_Outer membrane protein HopE
HP0705_Excinuclease ABC subunit A UvrA
HP0703_Transcriptional activator of flagella proteins FlgR
HP0702_Hypothetical protein HP0702
HP0701_DNA gyrase subunit A
HP0700_Diacylglycerol kinase
HP0699_Hypothetical protein HP0699
HP0697_Acetone carboxylase subunit gamma
HP0696_N-methylhydantoinase
HP0695_Hydantoin utilization protein A HyuA
HP0693_Short-chain fatty acids transporter
HP0692_Succinyl-CoA-transferase subunit B
HP0691_Succinyl-CoA-transferase subunit A
HP0690_Acetyl-CoA acetyltransferase
HP0687_Ferrous iron transport protein B
HP0686_Iron(III) dicitrate transport protein FecA
HP0683_Bifunctional N-acetylglucosamine-1-phosphate uridyltransferase/glucosamine-1-phosphate acetyltransferase
HP0682_Class IIc Sec-secreted bacteriocin Divergicin A precursor (DvnA)
HP0681_Hypothetical protein HP0681
HP0680_Ribonucleotide-diphosphate reductase subunit alpha
HP0677_4-Toluene Sulfonate Uptake Permease (TSUP)
HP0676_Methylated-DNA--protein-cysteine methyltransferase
HP0675_Integrase/recombinase XerC
HP0672_Aspartate aminotransferase
HP0671_Outer membrane protein HorF
HP0670_Predicted coding region HP0670
HP0666_Anaerobic glycerol-3-phosphate dehydrogenase subunit C GlpC
HP0665_Coproporphyrinogen III oxidase HemN
HP0663_Chorismate synthase AroC
HP0662_Ribonuclease III
HP0661_Ribonuclease H
HP0660_Hypothetical protein HP0660
HP0659_Chaperone SurA
HP0658_Aspartyl/glutamyl-tRNA amidotransferase subunit B
HP0657_Processing protease YmxG
HP0656_SCO4550 family menaquinone biosynthesis protein
HP0655_Protective surface antigen D15
HP0654_Menaquinone biosynthesis protein- SCO4494 family
HP0653_Ferritin
HP0652_Phosphoserine phosphatase SerB
HP0651_Alpha1-3-fucosyltransferase FutB
HP0650_Family 4 uracil-DNA glycosylase
HP0649_Aspartate ammonia-lyase AspA
HP0648_UDP-N-acetylglucosamine 1-carboxyvinyltransferase MurA
HP0647_Hypothetical protein HP0647
HP0646_UTP-glucose-1-phosphate uridylyltransferase GalU
HP0645_Soluble lytic murein transglycosylase

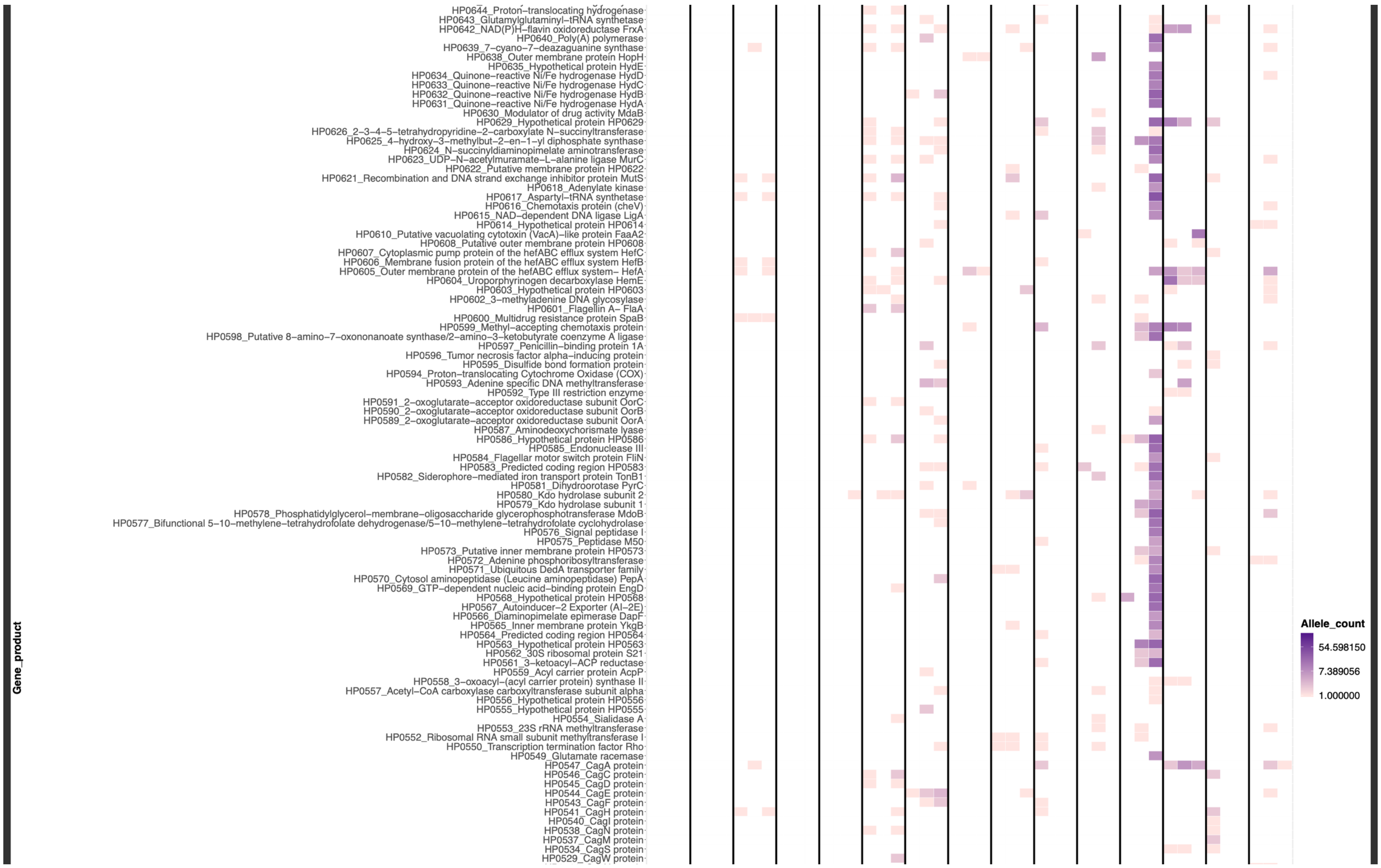
Gene_product
HP0644_Proton–translocating hydrogenase
HP0643_Glutamylglutaminyl–tRNA synthetase
HP0642_NAD(P)H–flavin oxidoreductase FrxA
HP0640_Poly(A) polymerase
HP0639_7–cyano–7–deazaguanine synthase
HP0638_Outer membrane protein HopH
HP0635_Hypothetical protein HydE
HP0634_Quinone–reactive Ni/Fe hydrogenase HydD
HP0633_Quinone–reactive Ni/Fe hydrogenase HydC
HP0632_Quinone–reactive Ni/Fe hydrogenase HydB
HP0631_Quinone–reactive Ni/Fe hydrogenase HydA
HP0630_Modulator of drug activity MdaB
HP0629_Hypothetical protein HP0629
HP0626_2–3–4–5–tetrahydropyridine–2–carboxylate N–succinyltransferase
HP0625_4–hydroxy–3–methylbut–2–en–1–yl diphosphate synthase
HP0624_N–succinyldiaminopimelate aminotransferase
HP0623_UDP–N–acetylmuramate–L–alanine ligase MurC
HP0622_Putative membrane protein HP0622
HP0621_Recombination and DNA strand exchange inhibitor protein MutS
HP0618_Adenylate kinase
HP0617_Aspartyl–tRNA synthetase
HP0616_Chemotaxis protein (cheV)
HP0615_NAD–dependent DNA ligase LigA
HP0614_Hypothetical protein HP0614
HP0610_Putative vacuolating cytotoxin (VacA)–like protein FaaA2
HP0608_Putative outer membrane protein HP0608
HP0607_Cytoplasmic pump protein of the hefABC efflux system HefC
HP0606_Membrane fusion protein of the hefABC efflux system HefB
HP0605_Outer membrane protein of the hefABC efflux system– HefA
HP0604_Uroporphyrinogen decarboxylase HemE
HP0603_Hypothetical protein HP0603
HP0602_3–methyladenine DNA glycosylase
HP0601_Flagellin A– FlaA
HP0600_Multidrug resistance protein SpaB
HP0599_Methyl–accepting chemotaxis protein
HP0598_Putative 8–amino–7–oxononanoate synthase/2–amino–3–ketobutyrate coenzyme A ligase
HP0597_Penicillin–binding protein 1A
HP0596_Tumor necrosis factor alpha–inducing protein
HP0595_Disulfide bond formation protein
HP0594_Proton–translocating Cytochrome Oxidase (COX)
HP0593_Adenine specific DNA methyltransferase
HP0592_Type III restriction enzyme
HP0591_2–oxoglutarate–acceptor oxidoreductase subunit OorC
HP0590_2–oxoglutarate–acceptor oxidoreductase subunit OorB
HP0589_2–oxoglutarate–acceptor oxidoreductase subunit OorA
HP0587_Aminodeoxychorismate lyase
HP0586_Hypothetical protein HP0586
HP0585_Endonuclease III
HP0584_Flagellar motor switch protein FliN
HP0583_Predicted coding region HP0583
HP0582_Siderophore–mediated iron transport protein TonB1
HP0581_Dihydroorotase PyrC
HP0580_Kdo hydrolase subunit 2
HP0579_Kdo hydrolase subunit 1
HP0578_Phosphatidylglycerol–membrane–oligosaccharide glycerophosphotransferase MdoB
HP0577_Bifunctional 5–10–methylene–tetrahydrofolate dehydrogenase/5–10–methylene–tetrahydrofolate cyclohydrolase
HP0576_Signal peptidase I
HP0575_Peptidase M50
HP0573_Putative inner membrane protein HP0573
HP0572_Adenine phosphoribosyltransferase
HP0571_Ubiquitous DedA transporter family
HP0570_Cytosol aminopeptidase (Leucine aminopeptidase) PepA
HP0569_GTP–dependent nucleic acid–binding protein EngD
HP0568_Hypothetical protein HP0568
HP0567_Autoinducer–2 Exporter (AI–2E)
HP0566_Diaminopimelate epimerase DapF
HP0565_Inner membrane protein YkgB
HP0564_Predicted coding region HP0564
HP0563_Hypothetical protein HP0563
HP0562_30S ribosomal protein S21
HP0561_3–ketoacyl–ACP reductase
HP0559_Acyl carrier protein AcpP
HP0558_3–oxoacyl–(acyl carrier protein) synthase II
HP0557_Acetyl–CoA carboxylase carboxyltransferase subunit alpha
HP0556_Hypothetical protein HP0556
HP0555_Hypothetical protein HP0555
HP0554_Sialidase A
HP0553_23S rRNA methyltransferase
HP0552_Ribosomal RNA small subunit methyltransferase I
HP0550_Transcription termination factor Rho
HP0549_Glutamate racemase
HP0547_CagA protein
HP0546_CagC protein
HP0545_CagD protein
HP0544_CagE protein
HP0543_CagF protein
HP0541_CagH protein
HP0540_CagI protein
HP0538_CagN protein
HP0537_CagM protein
HP0534_CagS protein
HP0529_CagW protein
Allele_count
54.598150
7.389056
1.000000

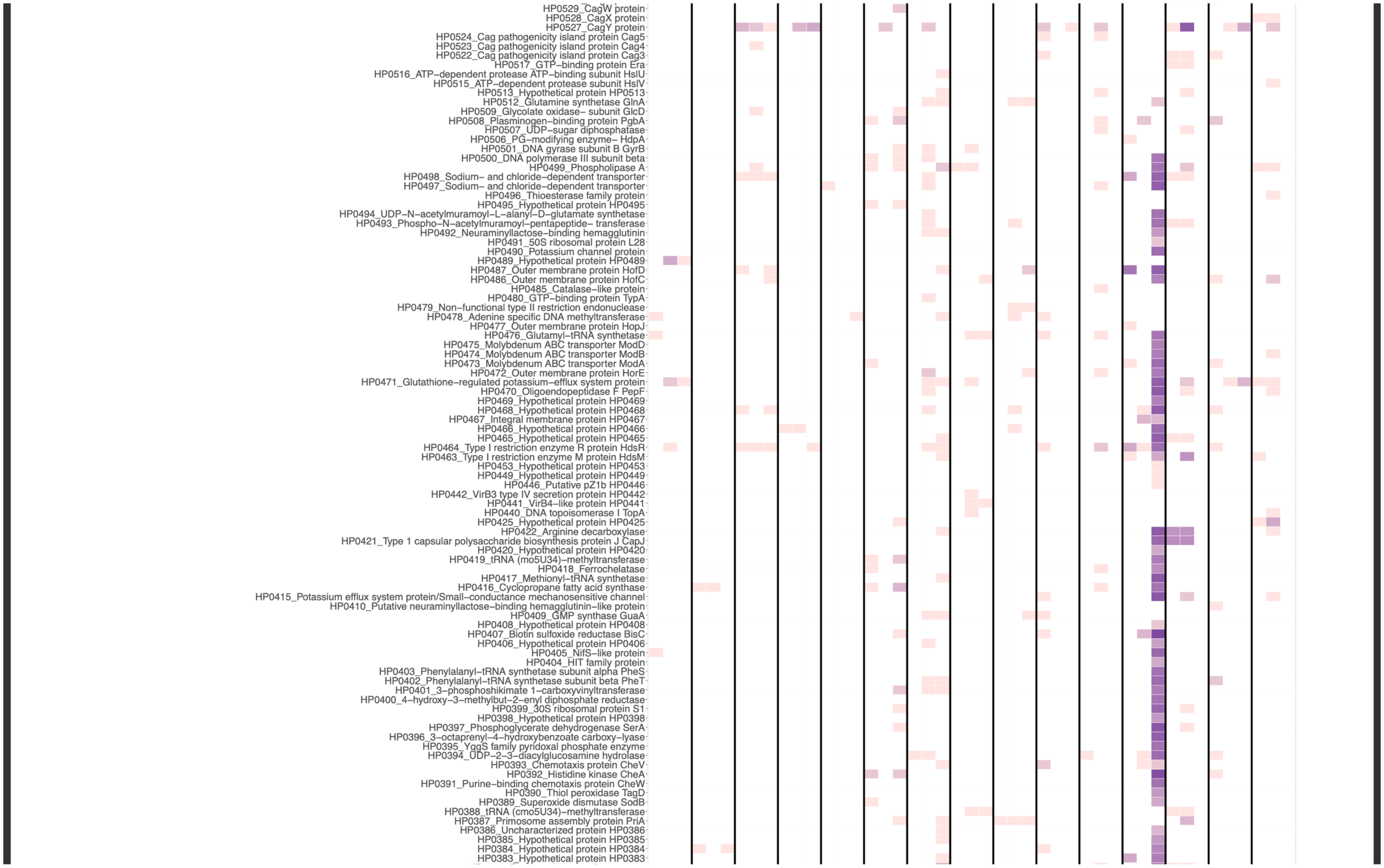
HP0529_CagW protein
HP0528_CagX protein
HP0527_CagY protein
HP0524_Cag pathogenicity island protein Cag5
HP0523_Cag pathogenicity island protein Cag4
HP0522_Cag pathogenicity island protein Cag3
HP0517_GTP–binding protein Era
HP0516_ATP–dependent protease ATP–binding subunit HslU
HP0515_ATP–dependent protease subunit HslV
HP0513_Hypothetical protein HP0513
HP0512_Glutamine synthetase GlnA
HP0509_Glycolate oxidase– subunit GlcD
HP0508_Plasminogen–binding protein PgbA
HP0507_UDP–sugar diphosphatase
HP0506_PG–modifying enzyme– HdpA
HP0501_DNA gyrase subunit B GyrB
HP0500_DNA polymerase III subunit beta
HP0499_Phospholipase A
HP0498_Sodium– and chloride–dependent transporter
HP0497_Sodium– and chloride–dependent transporter
HP0496_Thioesterase family protein
HP0495_Hypothetical protein HP0495
HP0494_UDP–N–acetylmuramoyl–L–alanyl–D–glutamate synthetase
HP0493_Phospho–N–acetylmuramoyl–pentapeptide– transferase
HP0492_Neuraminyllactose–binding hemagglutinin
HP0491_50S ribosomal protein L28
HP0490_Potassium channel protein
HP0489_Hypothetical protein HP0489
HP0487_Outer membrane protein HofD
HP0486_Outer membrane protein HofC
HP0485_Catalase–like protein
HP0480_GTP–binding protein TypA
HP0479_Non–functional type II restriction endonuclease
HP0478_Adenine specific DNA methyltransferase
HP0477_Outer membrane protein HopJ
HP0476_Glutamyl–tRNA synthetase
HP0475_Molybdenum ABC transporter ModD
HP0474_Molybdenum ABC transporter ModB
HP0473_Molybdenum ABC transporter ModA
HP0472_Outer membrane protein HorE
HP0471_Glutathione–regulated potassium–efflux system protein
HP0470_Oligoendopeptidase F PepF
HP0469_Hypothetical protein HP0469
HP0468_Hypothetical protein HP0468
HP0467_Integral membrane protein HP0467
HP0466_Hypothetical protein HP0466
HP0465_Hypothetical protein HP0465
HP0464_Type I restriction enzyme R protein HdsR
HP0463_Type I restriction enzyme M protein HdsM
HP0453_Hypothetical protein HP0453
HP0449_Hypothetical protein HP0449
HP0446_Putative pZ1b HP0446
HP0442_VirB3 type IV secretion protein HP0442
HP0441_VirB4–like protein HP0441
HP0440_DNA topoisomerase I TopA
HP0425_Hypothetical protein HP0425
HP0422_Arginine decarboxylase
HP0421_Type 1 capsular polysaccharide biosynthesis protein J CapJ
HP0420_Hypothetical protein HP0420
HP0419_tRNA (mo5U34)–methyltransferase
HP0418_Ferrochelatase
HP0417_Methionyl–tRNA synthetase
HP0416_Cyclopropane fatty acid synthase
HP0415_Potassium efflux system protein/Small–conductance mechanosensitive channel
HP0410_Putative neuraminyllactose–binding hemagglutinin–like protein
HP0409_GMP synthase GuaA
HP0408_Hypothetical protein HP0408
HP0407_Biotin sulfoxide reductase BisC
HP0406_Hypothetical protein HP0406
HP0405_NifS–like protein
HP0404_HIT family protein
HP0403_Phenylalanyl–tRNA synthetase subunit alpha PheS
HP0402_Phenylalanyl–tRNA synthetase subunit beta PheT
HP0401_3–phosphoshikimate 1–carboxyvinyltransferase
HP0400_4–hydroxy–3–methylbut–2–enyl diphosphate reductase
HP0399_30S ribosomal protein S1
HP0398_Hypothetical protein HP0398
HP0397_Phosphoglycerate dehydrogenase SerA
HP0396_3–octaprenyl–4–hydroxybenzoate carboxy–lyase
HP0395_YggS family pyridoxal phosphate enzyme
HP0394_UDP–2–3–diacylglucosamine hydrolase
HP0393_Chemotaxis protein CheV
HP0392_Histidine kinase CheA
HP0391_Purine–binding chemotaxis protein CheW
HP0390_Thiol peroxidase TagD
HP0389_Superoxide dismutase SodB
HP0388_tRNA (cmo5U34)–methyltransferase
HP0387_Primosome assembly protein PriA
HP0386_Uncharacterized protein HP0386
HP0385_Hypothetical protein HP0385
HP0384_Hypothetical protein HP0384
HP0383_Hypothetical protein HP0383

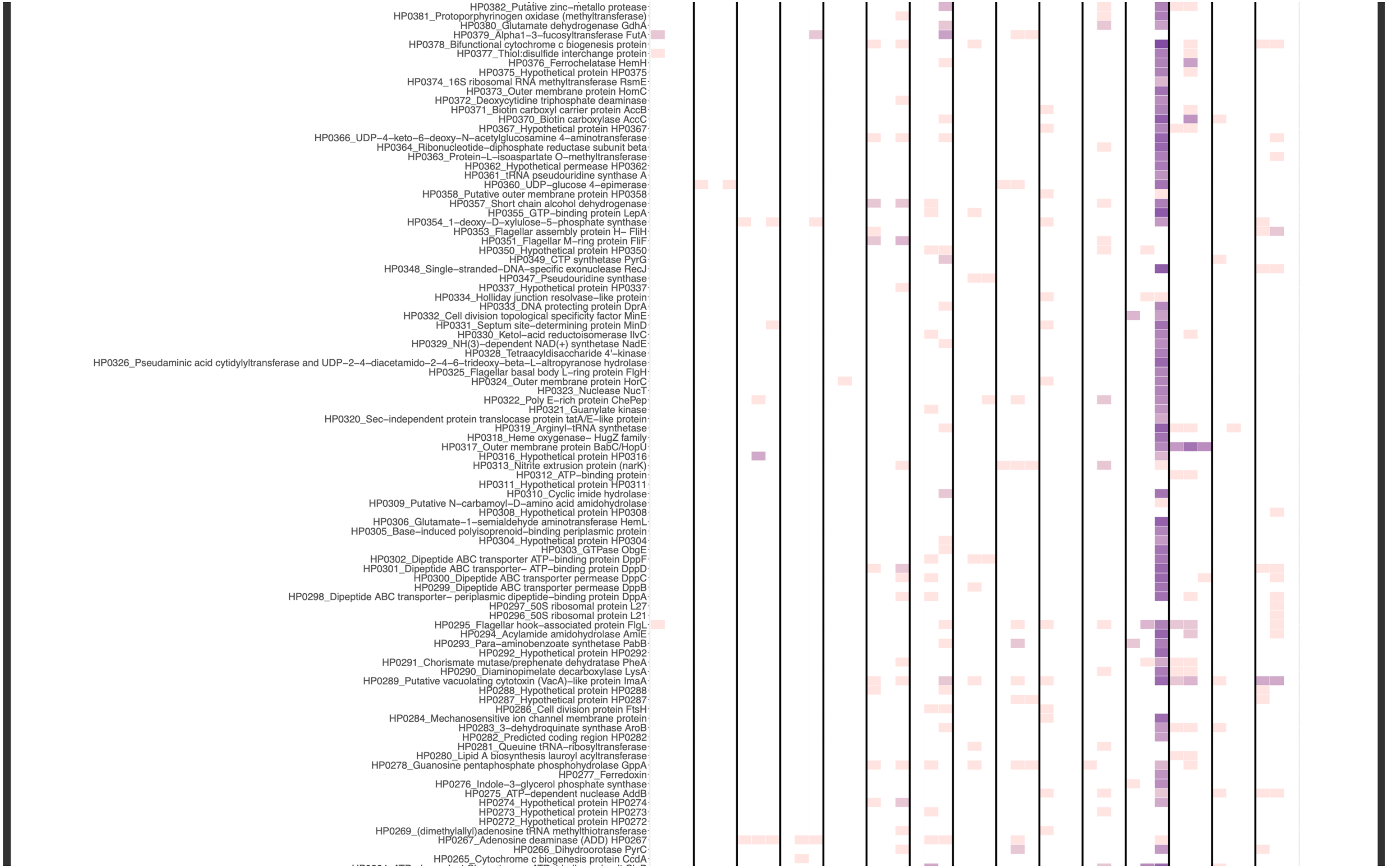
HP0382_Putative zinc–metallo protease
HP0381_Protoporphyrinogen oxidase (methyltransferase)
HP0380_Glutamate dehydrogenase GdhA
HP0379_Alpha1–3–fucosyltransferase FutA
HP0378_Bifunctional cytochrome c biogenesis protein
HP0377_Thiol:disulfide interchange protein
HP0376_Ferrochelatase HemH
HP0375_Hypothetical protein HP0375
HP0374_16S ribosomal RNA methyltransferase RsmE
HP0373_Outer membrane protein HomC
HP0372_Deoxycytidine triphosphate deaminase
HP0371_Biotin carboxyl carrier protein AccB
HP0370_Biotin carboxylase AccC
HP0367_Hypothetical protein HP0367
HP0366_UDP–4–keto–6–deoxy–N–acetylglucosamine 4–aminotransferase
HP0364_Ribonucleotide–diphosphate reductase subunit beta
HP0363_Protein–L–isoaspartate O–methyltransferase
HP0362_Hypothetical permease HP0362
HP0361_tRNA pseudouridine synthase A
HP0360_UDP–glucose 4–epimerase
HP0358_Putative outer membrane protein HP0358
HP0357_Short chain alcohol dehydrogenase
HP0355_GTP–binding protein LepA
HP0354_1–deoxy–D–xylulose–5–phosphate synthase
HP0353_Flagellar assembly protein H– FliH
HP0351_Flagellar M–ring protein FliF
HP0350_Hypothetical protein HP0350
HP0349_CTP synthetase PyrG
HP0348_Single–stranded–DNA–specific exonuclease RecJ
HP0347_Pseudouridine synthase
HP0337_Hypothetical protein HP0337
HP0334_Holliday junction resolvase–like protein
HP0333_DNA protecting protein DprA
HP0332_Cell division topological specificity factor MinE
HP0331_Septum site–determining protein MinD
HP0330_Ketol–acid reductoisomerase IlvC
HP0329_NH(3)–dependent NAD(+) synthetase NadE
HP0328_Tetraacyldisaccharide 4'–kinase
HP0326_Pseudaminic acid cytidylyltransferase and UDP–2–4–diacetamido–2–4–6–trideoxy–beta–L–altropyranose hydrolase
HP0325_Flagellar basal body L–ring protein FlgH
HP0324_Outer membrane protein HorC
HP0323_Nuclease NucT
HP0322_Poly E–rich protein ChePep
HP0321_Guanylate kinase
HP0320_Sec–independent protein translocase protein tatA/E–like protein
HP0319_Arginyl–tRNA synthetase
HP0318_Heme oxygenase– HugZ family
HP0317_Outer membrane protein BabC/HopU
HP0316_Hypothetical protein HP0316
HP0313_Nitrite extrusion protein (narK)
HP0312_ATP–binding protein
HP0311_Hypothetical protein HP0311
HP0310_Cyclic imide hydrolase
HP0309_Putative N–carbamoyl–D–amino acid amidohydrolase
HP0308_Hypothetical protein HP0308
HP0306_Glutamate–1–semialdehyde aminotransferase HemL
HP0305_Base–induced polyisoprenoid–binding periplasmic protein
HP0304_Hypothetical protein HP0304
HP0303_GTPase ObgE
HP0302_Dipeptide ABC transporter ATP–binding protein DppF
HP0301_Dipeptide ABC transporter– ATP–binding protein DppD
HP0300_Dipeptide ABC transporter permease DppC
HP0299_Dipeptide ABC transporter permease DppB
HP0298_Dipeptide ABC transporter– periplasmic dipeptide–binding protein DppA
HP0297_50S ribosomal protein L27
HP0296_50S ribosomal protein L21
HP0295_Flagellar hook–associated protein FlgL
HP0294_Acylamide amidohydrolase AmiE
HP0293_Para–aminobenzoate synthetase PabB
HP0292_Hypothetical protein HP0292
HP0291_Chorismate mutase/prephenate dehydratase PheA
HP0290_Diaminopimelate decarboxylase LysA
HP0289_Putative vacuolating cytotoxin (VacA)–like protein ImaA
HP0288_Hypothetical protein HP0288
HP0287_Hypothetical protein HP0287
HP0286_Cell division protein FtsH
HP0284_Mechanosensitive ion channel membrane protein
HP0283_3–dehydroquinate synthase AroB
HP0282_Predicted coding region HP0282
HP0281_Queuine tRNA–ribosyltransferase
HP0280_Lipid A biosynthesis lauroyl acyltransferase
HP0278_Guanosine pentaphosphate phosphohydrolase GppA
HP0277_Ferredoxin
HP0276_Indole–3–glycerol phosphate synthase
HP0275_ATP–dependent nuclease AddB
HP0274_Hypothetical protein HP0274
HP0273_Hypothetical protein HP0273
HP0272_Hypothetical protein HP0272
HP0269_(dimethylallyl)adenosine tRNA methylthiotransferase
HP0267_Adenosine deaminase (ADD) HP0267
HP0266_Dihydroorotase PyrC
HP0265_Cytochrome c biogenesis protein CcdA

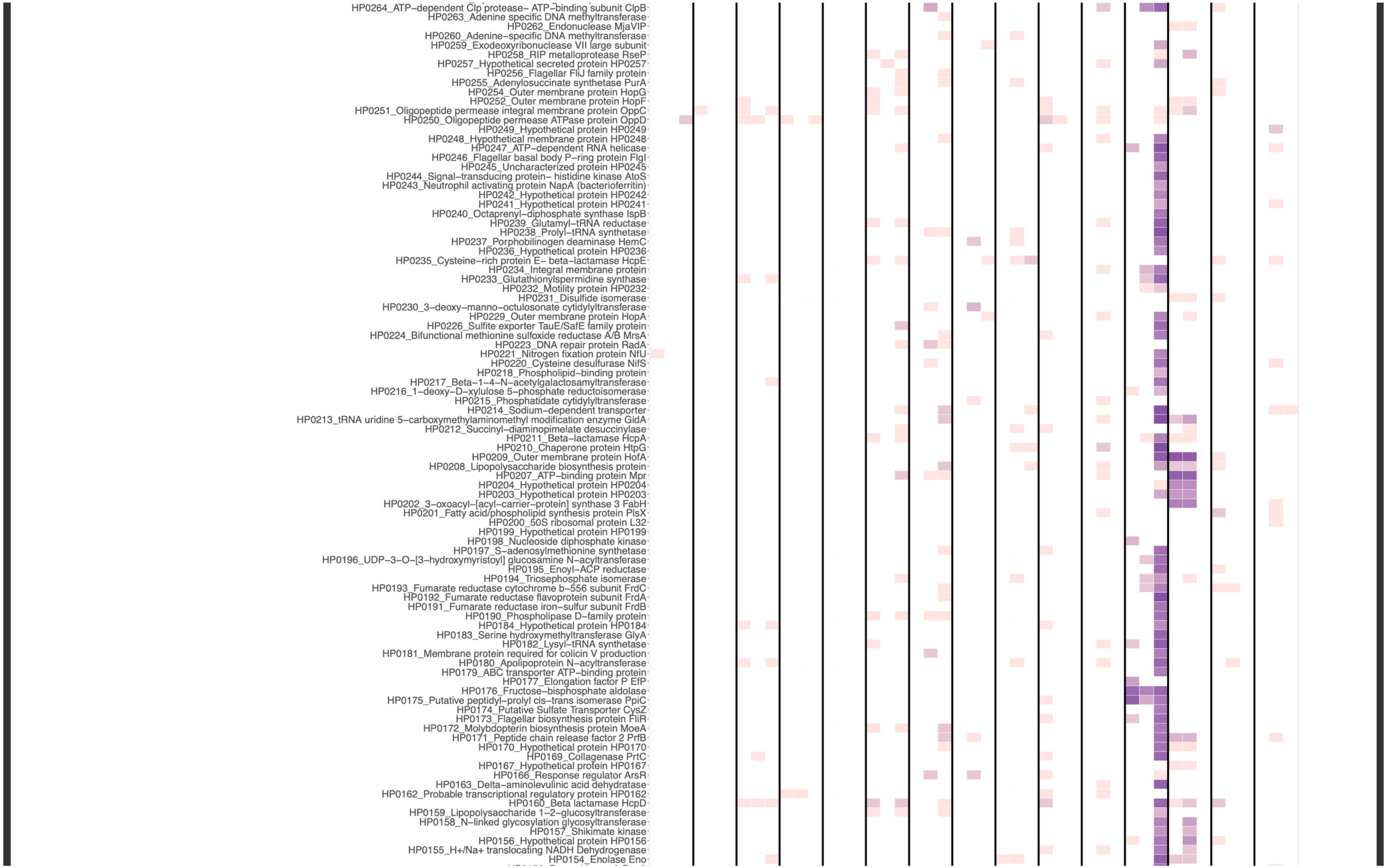
HP0264_ATP-dependent Clp protease- ATP-binding subunit ClpB
HP0263_Adenine specific DNA methyltransferase
HP0262_Endonuclease MjaVIP
HP0260_Adenine-specific DNA methyltransferase
HP0259_Exodeoxyribonuclease VII large subunit
HP0258_RIP metalloprotease RseP
HP0257_Hypothetical secreted protein HP0257
HP0256_Flagellar FliJ family protein
HP0255_Adenylosuccinate synthetase PurA
HP0254_Outer membrane protein HopG
HP0252_Outer membrane protein HopF
HP0251_Oligopeptide permease integral membrane protein OppC
HP0250_Oligopeptide permease ATPase protein OppD
HP0249_Hypothetical protein HP0249
HP0248_Hypothetical membrane protein HP0248
HP0247_ATP-dependent RNA helicase
HP0246_Flagellar basal body P-ring protein FlgI
HP0245_Uncharacterized protein HP0245
HP0244_Signal-transducing protein- histidine kinase AtoS
HP0243_Neutrophil activating protein NapA (bacterioferritin)
HP0242_Hypothetical protein HP0242
HP0241_Hypothetical protein HP0241
HP0240_Octaprenyl-diphosphate synthase IspB
HP0239_Glutamyl-tRNA reductase
HP0238_Prolyl-tRNA synthetase
HP0237_Porphobilinogen deaminase HemC
HP0236_Hypothetical protein HP0236
HP0235_Cysteine-rich protein E- beta-lactamase HcpE
HP0234_Integral membrane protein
HP0233_Glutathionylspermidine synthase
HP0232_Motility protein HP0232
HP0231_Disulfide isomerase
HP0230_3-deoxy-manno-octulosonate cytidylyltransferase
HP0229_Outer membrane protein HopA
HP0226_Sulfite exporter TauE/SafE family protein
HP0224_Bifunctional methionine sulfoxide reductase A/B MrsA
HP0223_DNA repair protein RadA
HP0221_Nitrogen fixation protein NfU
HP0220_Cysteine desulfurase NifS
HP0218_Phospholipid-binding protein
HP0217_Beta-1-4-N-acetylgalactosamyltransferase
HP0216_1-deoxy-D-xylulose 5-phosphate reductoisomerase
HP0215_Phosphatidate cytidylyltransferase
HP0214_Sodium-dependent transporter
HP0213_tRNA uridine 5-carboxymethylaminomethyl modification enzyme GidA
HP0212_Succinyl-diaminopimelate desuccinylase
HP0211_Beta-lactamase HcpA
HP0210_Chaperone protein HtpG
HP0209_Outer membrane protein HofA
HP0208_Lipopolysaccharide biosynthesis protein
HP0207_ATP-binding protein Mpr
HP0204_Hypothetical protein HP0204
HP0203_Hypothetical protein HP0203
HP0202_3-oxoacyl-[acyl-carrier-protein] synthase 3 FabH
HP0201_Fatty acid/phospholipid synthesis protein PlsX
HP0200_50S ribosomal protein L32
HP0199_Hypothetical protein HP0199
HP0198_Nucleoside diphosphate kinase
HP0197_S-adenosylmethionine synthetase
HP0196_UDP-3-O-[3-hydroxymyristoyl] glucosamine N-acyltransferase
HP0195_Enoyl-ACP reductase
HP0194_Triosephosphate isomerase
HP0193_Fumarate reductase cytochrome b-556 subunit FrdC
HP0192_Fumarate reductase flavoprotein subunit FrdA
HP0191_Fumarate reductase iron-sulfur subunit FrdB
HP0190_Phospholipase D-family protein
HP0184_Hypothetical protein HP0184
HP0183_Serine hydroxymethyltransferase GlyA
HP0182_Lysyl-tRNA synthetase
HP0181_Membrane protein required for colicin V production
HP0180_Apolipoprotein N-acyltransferase
HP0179_ABC transporter ATP-binding protein
HP0177_Elongation factor P EfP
HP0176_Fructose-bisphosphate aldolase
HP0175_Putative peptidyl-prolyl cis-trans isomerase PpiC
HP0174_Putative Sulfate Transporter CysZ
HP0173_Flagellar biosynthesis protein FliR
HP0172_Molybdopterin biosynthesis protein MoeA
HP0171_Peptide chain release factor 2 PrfB
HP0170_Hypothetical protein HP0170
HP0169_Collagenase PrtC
HP0167_Hypothetical protein HP0167
HP0166_Response regulator ArsR
HP0163_Delta-aminolevulinic acid dehydratase
HP0162_Probable transcriptional regulatory protein HP0162
HP0160_Beta lactamase HcpD
HP0159_Lipopolysaccharide 1-2-glucosyltransferase
HP0158_N-linked glycosylation glycosyltransferase
HP0157_Shikimate kinase
HP0156_Hypothetical protein HP0156
HP0155_H+/Na+ translocating NADH Dehydrogenase
HP0154_Enolase Eno

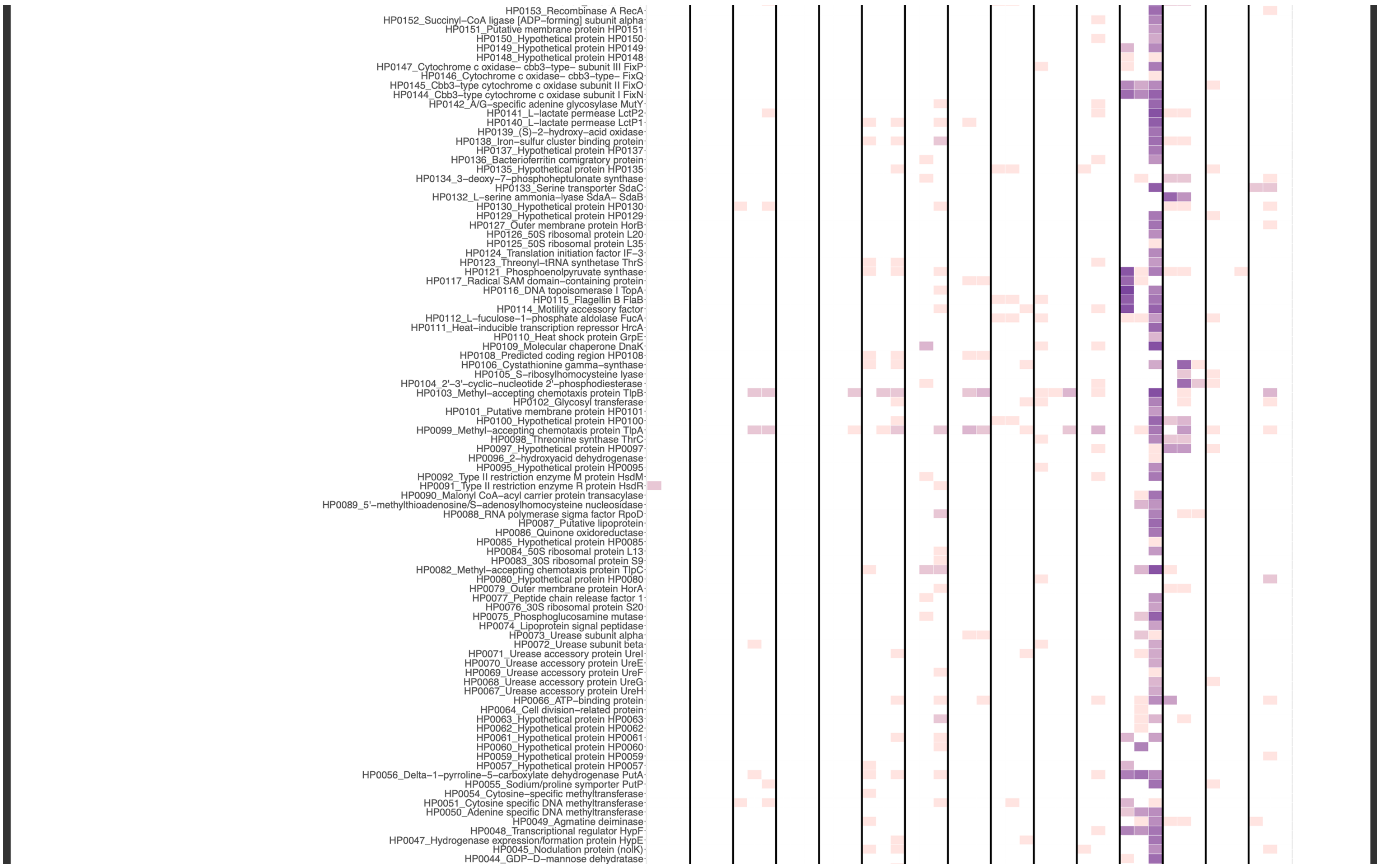
HP0153_Recombinase A RecA
HP0152_Succinyl−CoA ligase [ADP−forming] subunit alpha
HP0151_Putative membrane protein HP0151
HP0150_Hypothetical protein HP0150
HP0149_Hypothetical protein HP0149
HP0148_Hypothetical protein HP0148
HP0147_Cytochrome c oxidase− cbb3−type− subunit III FixP
HP0146_Cytochrome c oxidase− cbb3−type− FixQ
HP0145_Cbb3−type cytochrome c oxidase subunit II FixO
HP0144_Cbb3−type cytochrome c oxidase subunit I FixN
HP0142_A/G−specific adenine glycosylase MutY
HP0141_L−lactate permease LctP2
HP0140_L−lactate permease LctP1
HP0139_(S)−2−hydroxy−acid oxidase
HP0138_Iron−sulfur cluster binding protein
HP0137_Hypothetical protein HP0137
HP0136_Bacterioferritin comigratory protein
HP0135_Hypothetical protein HP0135
HP0134_3−deoxy−7−phosphoheptulonate synthase
HP0133_Serine transporter SdaC
HP0132_L−serine ammonia−lyase SdaA− SdaB
HP0130_Hypothetical protein HP0130
HP0129_Hypothetical protein HP0129
HP0127_Outer membrane protein HorB
HP0126_50S ribosomal protein L20
HP0125_50S ribosomal protein L35
HP0124_Translation initiation factor IF−3
HP0123_Threonyl−tRNA synthetase ThrS
HP0121_Phosphoenolpyruvate synthase
HP0117_Radical SAM domain−containing protein
HP0116_DNA topoisomerase I TopA
HP0115_Flagellin B FlaB
HP0114_Motility accessory factor
HP0112_L−fuculose−1−phosphate aldolase FucA
HP0111_Heat−inducible transcription repressor HrcA
HP0110_Heat shock protein GrpE
HP0109_Molecular chaperone DnaK
HP0108_Predicted coding region HP0108
HP0106_Cystathionine gamma−synthase
HP0105_S−ribosylhomocysteine lyase
HP0104_2'−3'−cyclic−nucleotide 2'−phosphodiesterase
HP0103_Methyl−accepting chemotaxis protein TlpB
HP0102_Glycosyl transferase
HP0101_Putative membrane protein HP0101
HP0100_Hypothetical protein HP0100
HP0099_Methyl−accepting chemotaxis protein TlpA
HP0098_Threonine synthase ThrC
HP0097_Hypothetical protein HP0097
HP0096_2−hydroxyacid dehydrogenase
HP0095_Hypothetical protein HP0095
HP0092_Type II restriction enzyme M protein HsdM
HP0091_Type II restriction enzyme R protein HsdR
HP0090_Malonyl CoA−acyl carrier protein transacylase
HP0089_5'−methylthioadenosine/S−adenosylhomocysteine nucleosidase
HP0088_RNA polymerase sigma factor RpoD
HP0087_Putative lipoprotein
HP0086_Quinone oxidoreductase
HP0085_Hypothetical protein HP0085
HP0084_50S ribosomal protein L13
HP0083_30S ribosomal protein S9
HP0082_Methyl−accepting chemotaxis protein TlpC
HP0080_Hypothetical protein HP0080
HP0079_Outer membrane protein HorA
HP0077_Peptide chain release factor 1
HP0076_30S ribosomal protein S20
HP0075_Phosphoglucosamine mutase
HP0074_Lipoprotein signal peptidase
HP0073_Urease subunit alpha
HP0072_Urease subunit beta
HP0071_Urease accessory protein UreI
HP0070_Urease accessory protein UreE
HP0069_Urease accessory protein UreF
HP0068_Urease accessory protein UreG
HP0067_Urease accessory protein UreH
HP0066_ATP−binding protein
HP0064_Cell division−related protein
HP0063_Hypothetical protein HP0063
HP0062_Hypothetical protein HP0062
HP0061_Hypothetical protein HP0061
HP0060_Hypothetical protein HP0060
HP0059_Hypothetical protein HP0059
HP0057_Hypothetical protein HP0057
HP0056_Delta−1−pyrroline−5−carboxylate dehydrogenase PutA
HP0055_Sodium/proline symporter PutP
HP0054_Cytosine−specific methyltransferase
HP0051_Cytosine specific DNA methyltransferase
HP0050_Adenine specific DNA methyltransferase
HP0049_Agmatine deiminase
HP0048_Transcriptional regulator HypF
HP0047_Hydrogenase expression/formation protein HypE
HP0045_Nodulation protein (nolK)
HP0044_GDP−D−mannose dehydratase

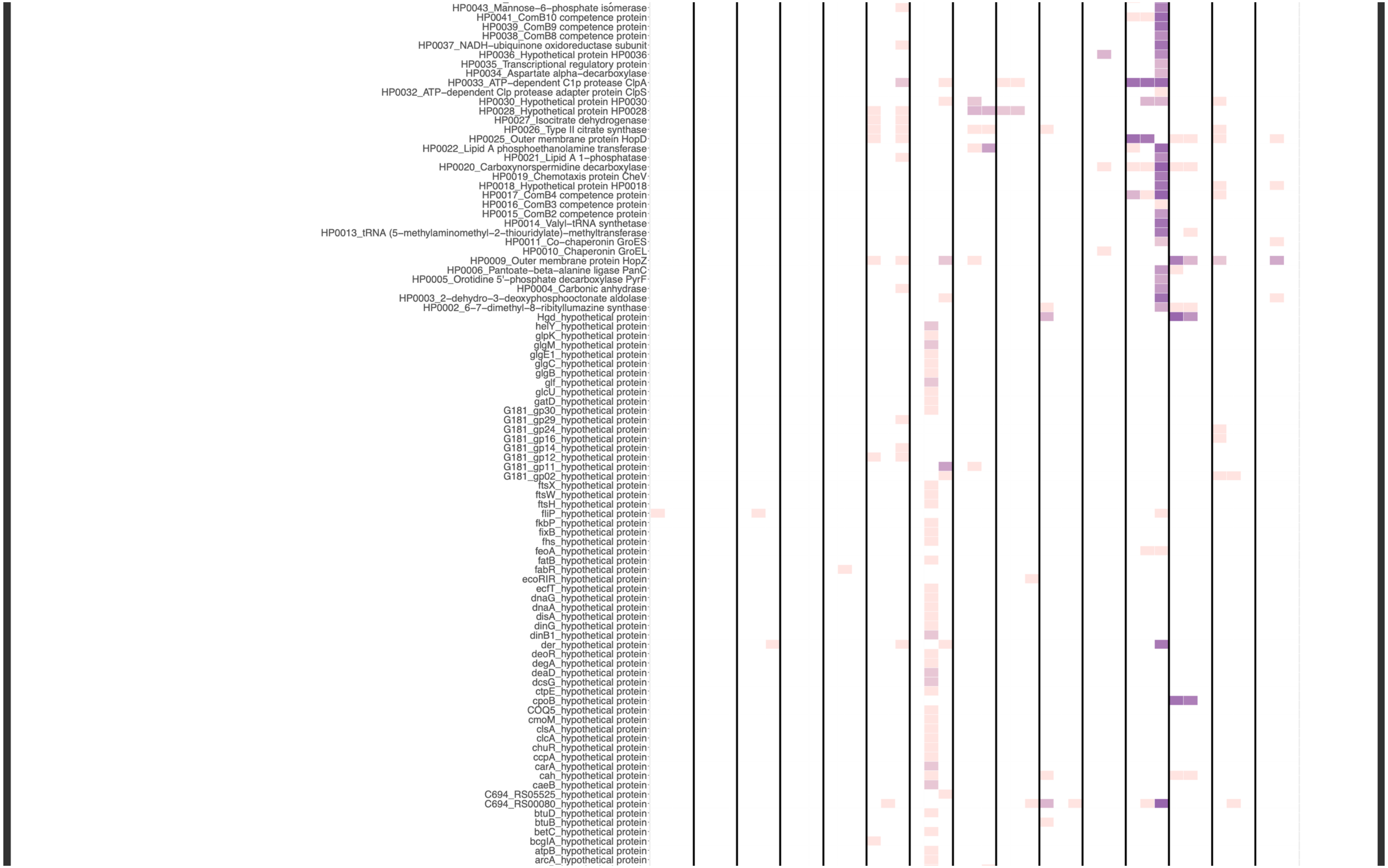
HP0043_Mannose–6–phosphate isomerase
HP0041_ComB10 competence protein
HP0039_ComB9 competence protein
HP0038_ComB8 competence protein
HP0037_NADH–ubiquinone oxidoreductase subunit
HP0036_Hypothetical protein HP0036
HP0035_Transcriptional regulatory protein
HP0034_Aspartate alpha–decarboxylase
HP0033_ATP–dependent C1p protease ClpA
HP0032_ATP–dependent Clp protease adapter protein ClpS
HP0030_Hypothetical protein HP0030
HP0028_Hypothetical protein HP0028
HP0027_Isocitrate dehydrogenase
HP0026_Type II citrate synthase
HP0025_Outer membrane protein HopD
HP0022_Lipid A phosphoethanolamine transferase
HP0021_Lipid A 1–phosphatase
HP0020_Carboxynorspermidine decarboxylase
HP0019_Chemotaxis protein CheV
HP0018_Hypothetical protein HP0018
HP0017_ComB4 competence protein
HP0016_ComB3 competence protein
HP0015_ComB2 competence protein
HP0014_Valyl–tRNA synthetase
HP0013_tRNA (5–methylaminomethyl–2–thiouridylate)–methyltransferase
HP0011_Co–chaperonin GroES
HP0010_Chaperonin GroEL
HP0009_Outer membrane protein HopZ
HP0006_Pantoate–beta–alanine ligase PanC
HP0005_Orotidine 5'–phosphate decarboxylase PyrF
HP0004_Carbonic anhydrase
HP0003_2–dehydro–3–deoxyphosphooctonate aldolase
HP0002_6–7–dimethyl–8–ribityllumazine synthase
Hgd_hypothetical protein
helY_hypothetical protein
glpK_hypothetical protein
glgM_hypothetical protein
glgE1_hypothetical protein
glgC_hypothetical protein
glgB_hypothetical protein
glf_hypothetical protein
glcU_hypothetical protein
gatD_hypothetical protein
G181_gp30_hypothetical protein
G181_gp29_hypothetical protein
G181_gp24_hypothetical protein
G181_gp16_hypothetical protein
G181_gp14_hypothetical protein
G181_gp12_hypothetical protein
G181_gp11_hypothetical protein
G181_gp02_hypothetical protein
ftsX_hypothetical protein
ftsW_hypothetical protein
ftsH_hypothetical protein
fliP_hypothetical protein
fkbP_hypothetical protein
fixB_hypothetical protein
fhs_hypothetical protein
feoA_hypothetical protein
fatB_hypothetical protein
fabR_hypothetical protein
ecoRIR_hypothetical protein
ecfT_hypothetical protein
dnaG_hypothetical protein
dnaA_hypothetical protein
disA_hypothetical protein
dinG_hypothetical protein
dinB1_hypothetical protein
der_hypothetical protein
deoR_hypothetical protein
degA_hypothetical protein
deaD_hypothetical protein
dcsG_hypothetical protein
ctpE_hypothetical protein
cpoB_hypothetical protein
COQ5_hypothetical protein
cmoM_hypothetical protein
clsA_hypothetical protein
clcA_hypothetical protein
chuR_hypothetical protein
ccpA_hypothetical protein
carA_hypothetical protein
cah_hypothetical protein
caeB_hypothetical protein
C694_RS05525_hypothetical protein
C694_RS00080_hypothetical protein
btuD_hypothetical protein
btuB_hypothetical protein
betC_hypothetical protein
bcgIA_hypothetical protein
atpB_hypothetical protein
arcA_hypothetical protein

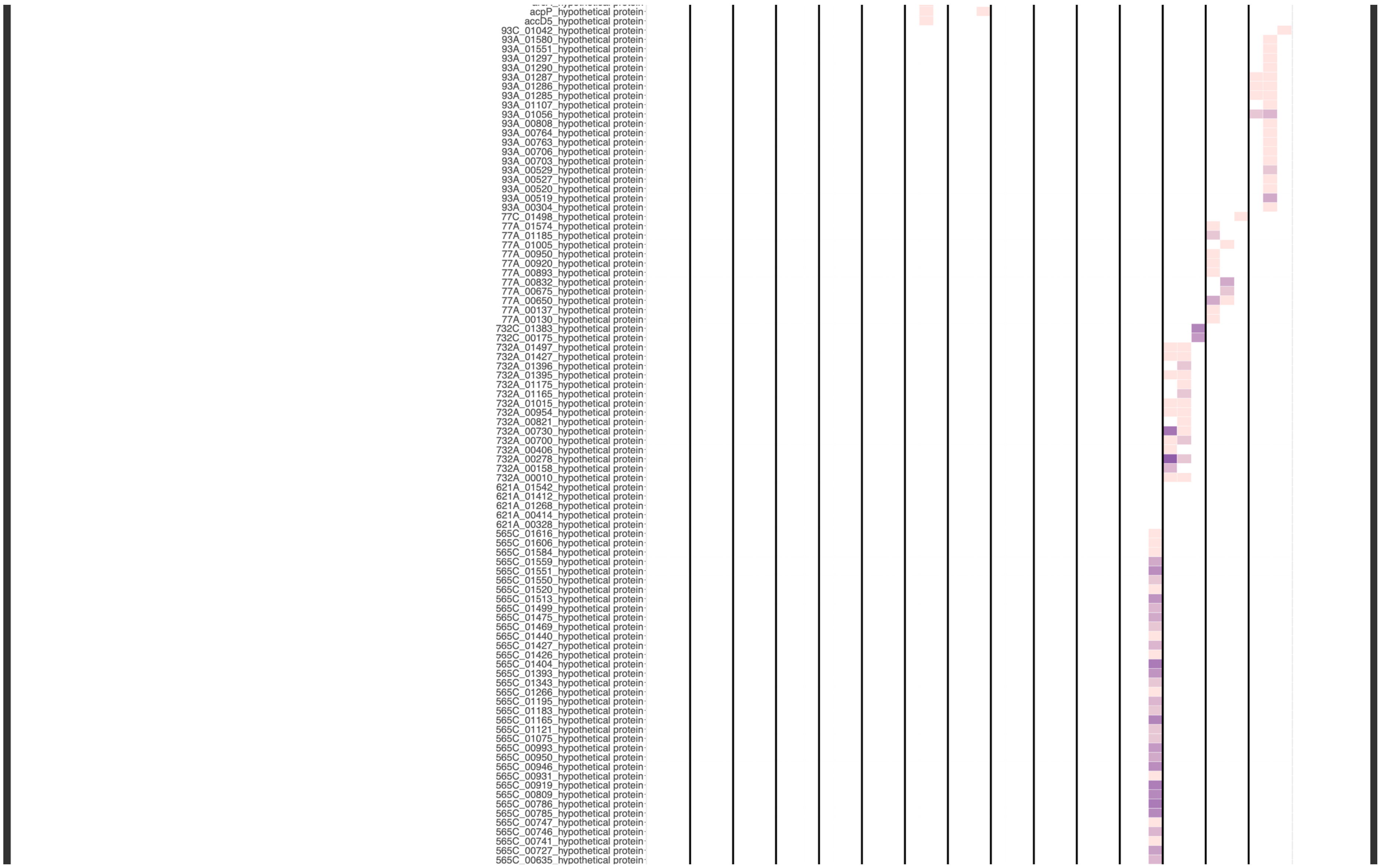

acpP_hypothetical protein
accD5_hypothetical protein
93C_01042_hypothetical protein
93A_01580_hypothetical protein
93A_01551_hypothetical protein
93A_01297_hypothetical protein
93A_01290_hypothetical protein
93A_01287_hypothetical protein
93A_01286_hypothetical protein
93A_01285_hypothetical protein
93A_01107_hypothetical protein
93A_01056_hypothetical protein
93A_00808_hypothetical protein
93A_00764_hypothetical protein
93A_00763_hypothetical protein
93A_00706_hypothetical protein
93A_00703_hypothetical protein
93A_00529_hypothetical protein
93A_00527_hypothetical protein
93A_00520_hypothetical protein
93A_00519_hypothetical protein
93A_00304_hypothetical protein
77C_01498_hypothetical protein
77A_01574_hypothetical protein
77A_01185_hypothetical protein
77A_01005_hypothetical protein
77A_00950_hypothetical protein
77A_00920_hypothetical protein
77A_00893_hypothetical protein
77A_00832_hypothetical protein
77A_00675_hypothetical protein
77A_00650_hypothetical protein
77A_00137_hypothetical protein
77A_00130_hypothetical protein
732C_01383_hypothetical protein
732C_00175_hypothetical protein
732A_01497_hypothetical protein
732A_01427_hypothetical protein
732A_01396_hypothetical protein
732A_01395_hypothetical protein
732A_01175_hypothetical protein
732A_01165_hypothetical protein
732A_01015_hypothetical protein
732A_00954_hypothetical protein
732A_00821_hypothetical protein
732A_00730_hypothetical protein
732A_00700_hypothetical protein
732A_00406_hypothetical protein
732A_00278_hypothetical protein
732A_00158_hypothetical protein
732A_00010_hypothetical protein
621A_01542_hypothetical protein
621A_01412_hypothetical protein
621A_01268_hypothetical protein
621A_00414_hypothetical protein
621A_00328_hypothetical protein
565C_01616_hypothetical protein
565C_01606_hypothetical protein
565C_01584_hypothetical protein
565C_01559_hypothetical protein
565C_01551_hypothetical protein
565C_01550_hypothetical protein
565C_01520_hypothetical protein
565C_01513_hypothetical protein
565C_01499_hypothetical protein
565C_01475_hypothetical protein
565C_01469_hypothetical protein
565C_01440_hypothetical protein
565C_01427_hypothetical protein
565C_01426_hypothetical protein
565C_01404_hypothetical protein
565C_01393_hypothetical protein
565C_01343_hypothetical protein
565C_01266_hypothetical protein
565C_01195_hypothetical protein
565C_01183_hypothetical protein
565C_01165_hypothetical protein
565C_01121_hypothetical protein
565C_01075_hypothetical protein
565C_00993_hypothetical protein
565C_00950_hypothetical protein
565C_00946_hypothetical protein
565C_00931_hypothetical protein
565C_00919_hypothetical protein
565C_00809_hypothetical protein
565C_00786_hypothetical protein
565C_00785_hypothetical protein
565C_00747_hypothetical protein
565C_00746_hypothetical protein
565C_00741_hypothetical protein
565C_00727_hypothetical protein
565C_00635_hypothetical protein

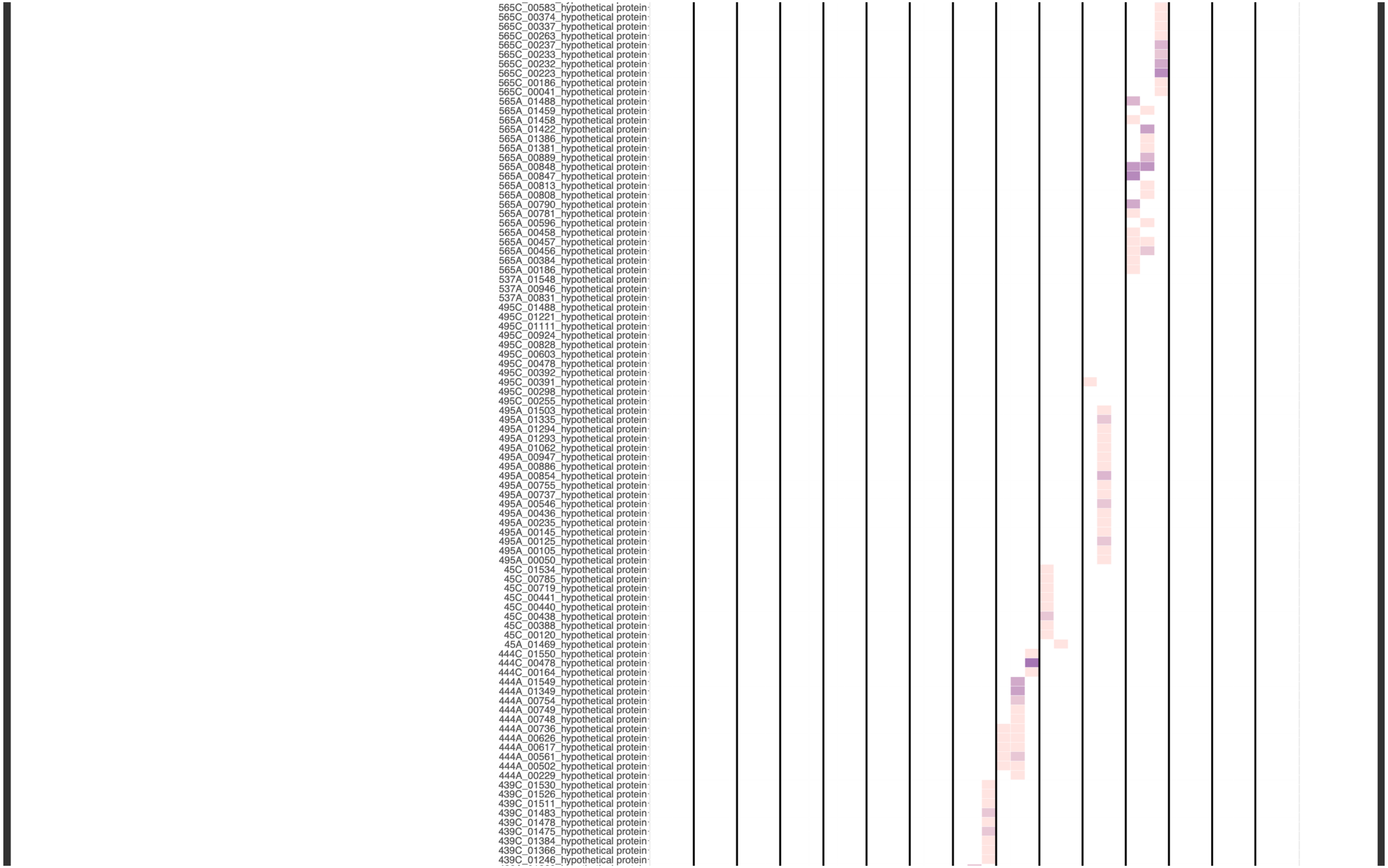
565C_00583_hypothetical protein
565C_00374_hypothetical protein
565C_00337_hypothetical protein
565C_00263_hypothetical protein
565C_00237_hypothetical protein
565C_00233_hypothetical protein
565C_00232_hypothetical protein
565C_00223_hypothetical protein
565C_00186_hypothetical protein
565C_00041_hypothetical protein
565A_01488_hypothetical protein
565A_01459_hypothetical protein
565A_01458_hypothetical protein
565A_01422_hypothetical protein
565A_01386_hypothetical protein
565A_01381_hypothetical protein
565A_00889_hypothetical protein
565A_00848_hypothetical protein
565A_00847_hypothetical protein
565A_00813_hypothetical protein
565A_00808_hypothetical protein
565A_00790_hypothetical protein
565A_00781_hypothetical protein
565A_00596_hypothetical protein
565A_00458_hypothetical protein
565A_00457_hypothetical protein
565A_00456_hypothetical protein
565A_00384_hypothetical protein
565A_00186_hypothetical protein
537A_01548_hypothetical protein
537A_00946_hypothetical protein
537A_00831_hypothetical protein
495C_01488_hypothetical protein
495C_01221_hypothetical protein
495C_01111_hypothetical protein
495C_00924_hypothetical protein
495C_00828_hypothetical protein
495C_00603_hypothetical protein
495C_00478_hypothetical protein
495C_00392_hypothetical protein
495C_00391_hypothetical protein
495C_00298_hypothetical protein
495C_00255_hypothetical protein
495A_01503_hypothetical protein
495A_01335_hypothetical protein
495A_01294_hypothetical protein
495A_01293_hypothetical protein
495A_01062_hypothetical protein
495A_00947_hypothetical protein
495A_00886_hypothetical protein
495A_00854_hypothetical protein
495A_00755_hypothetical protein
495A_00737_hypothetical protein
495A_00546_hypothetical protein
495A_00436_hypothetical protein
495A_00235_hypothetical protein
495A_00145_hypothetical protein
495A_00125_hypothetical protein
495A_00105_hypothetical protein
495A_00050_hypothetical protein
45C_01534_hypothetical protein
45C_00785_hypothetical protein
45C_00719_hypothetical protein
45C_00441_hypothetical protein
45C_00440_hypothetical protein
45C_00438_hypothetical protein
45C_00388_hypothetical protein
45C_00120_hypothetical protein
45A_01469_hypothetical protein
444C_01550_hypothetical protein
444C_00478_hypothetical protein
444C_00164_hypothetical protein
444A_01549_hypothetical protein
444A_01349_hypothetical protein
444A_00754_hypothetical protein
444A_00749_hypothetical protein
444A_00748_hypothetical protein
444A_00736_hypothetical protein
444A_00626_hypothetical protein
444A_00617_hypothetical protein
444A_00561_hypothetical protein
444A_00502_hypothetical protein
444A_00229_hypothetical protein
439C_01530_hypothetical protein
439C_01526_hypothetical protein
439C_01511_hypothetical protein
439C_01483_hypothetical protein
439C_01478_hypothetical protein
439C_01475_hypothetical protein
439C_01384_hypothetical protein
439C_01366_hypothetical protein
439C_01246_hypothetical protein

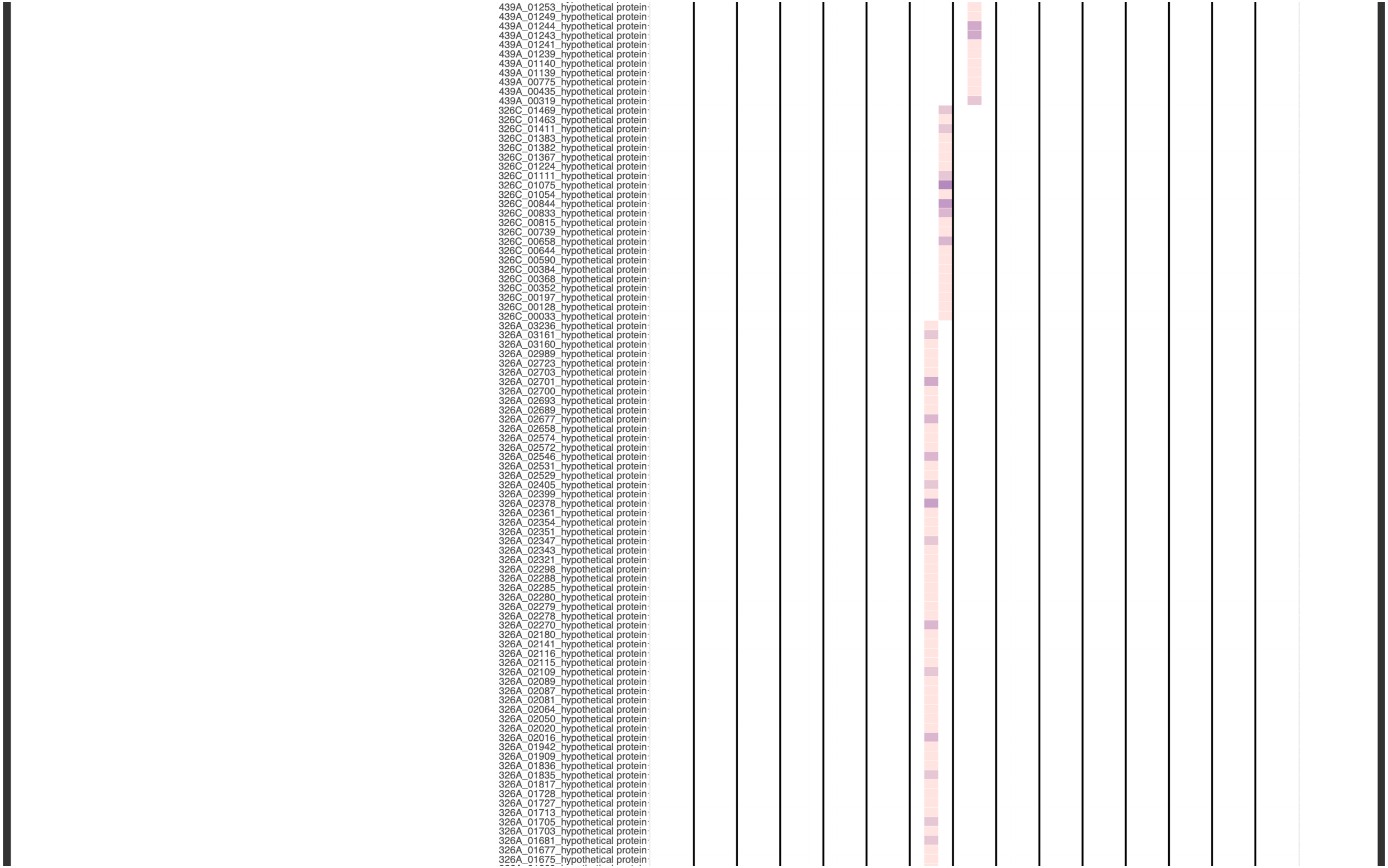

439A_01253_hypothetical protein
439A_01249_hypothetical protein
439A_01244_hypothetical protein
439A_01243_hypothetical protein
439A_01241_hypothetical protein
439A_01239_hypothetical protein
439A_01140_hypothetical protein
439A_01139_hypothetical protein
439A_00775_hypothetical protein
439A_00435_hypothetical protein
439A_00319_hypothetical protein
326C_01469_hypothetical protein
326C_01463_hypothetical protein
326C_01411_hypothetical protein
326C_01383_hypothetical protein
326C_01382_hypothetical protein
326C_01367_hypothetical protein
326C_01224_hypothetical protein
326C_01111_hypothetical protein
326C_01075_hypothetical protein
326C_01054_hypothetical protein
326C_00844_hypothetical protein
326C_00833_hypothetical protein
326C_00815_hypothetical protein
326C_00739_hypothetical protein
326C_00658_hypothetical protein
326C_00644_hypothetical protein
326C_00590_hypothetical protein
326C_00384_hypothetical protein
326C_00368_hypothetical protein
326C_00352_hypothetical protein
326C_00197_hypothetical protein
326C_00128_hypothetical protein
326C_00033_hypothetical protein
326A_03236_hypothetical protein
326A_03161_hypothetical protein
326A_03160_hypothetical protein
326A_02989_hypothetical protein
326A_02723_hypothetical protein
326A_02703_hypothetical protein
326A_02701_hypothetical protein
326A_02700_hypothetical protein
326A_02693_hypothetical protein
326A_02689_hypothetical protein
326A_02677_hypothetical protein
326A_02658_hypothetical protein
326A_02574_hypothetical protein
326A_02572_hypothetical protein
326A_02546_hypothetical protein
326A_02531_hypothetical protein
326A_02529_hypothetical protein
326A_02405_hypothetical protein
326A_02399_hypothetical protein
326A_02378_hypothetical protein
326A_02361_hypothetical protein
326A_02354_hypothetical protein
326A_02351_hypothetical protein
326A_02347_hypothetical protein
326A_02343_hypothetical protein
326A_02321_hypothetical protein
326A_02298_hypothetical protein
326A_02288_hypothetical protein
326A_02285_hypothetical protein
326A_02280_hypothetical protein
326A_02279_hypothetical protein
326A_02278_hypothetical protein
326A_02270_hypothetical protein
326A_02180_hypothetical protein
326A_02141_hypothetical protein
326A_02116_hypothetical protein
326A_02115_hypothetical protein
326A_02109_hypothetical protein
326A_02089_hypothetical protein
326A_02087_hypothetical protein
326A_02081_hypothetical protein
326A_02064_hypothetical protein
326A_02050_hypothetical protein
326A_02020_hypothetical protein
326A_02016_hypothetical protein
326A_01942_hypothetical protein
326A_01909_hypothetical protein
326A_01836_hypothetical protein
326A_01835_hypothetical protein
326A_01817_hypothetical protein
326A_01728_hypothetical protein
326A_01727_hypothetical protein
326A_01713_hypothetical protein
326A_01705_hypothetical protein
326A_01703_hypothetical protein
326A_01681_hypothetical protein
326A_01677_hypothetical protein
326A_01675_hypothetical protein

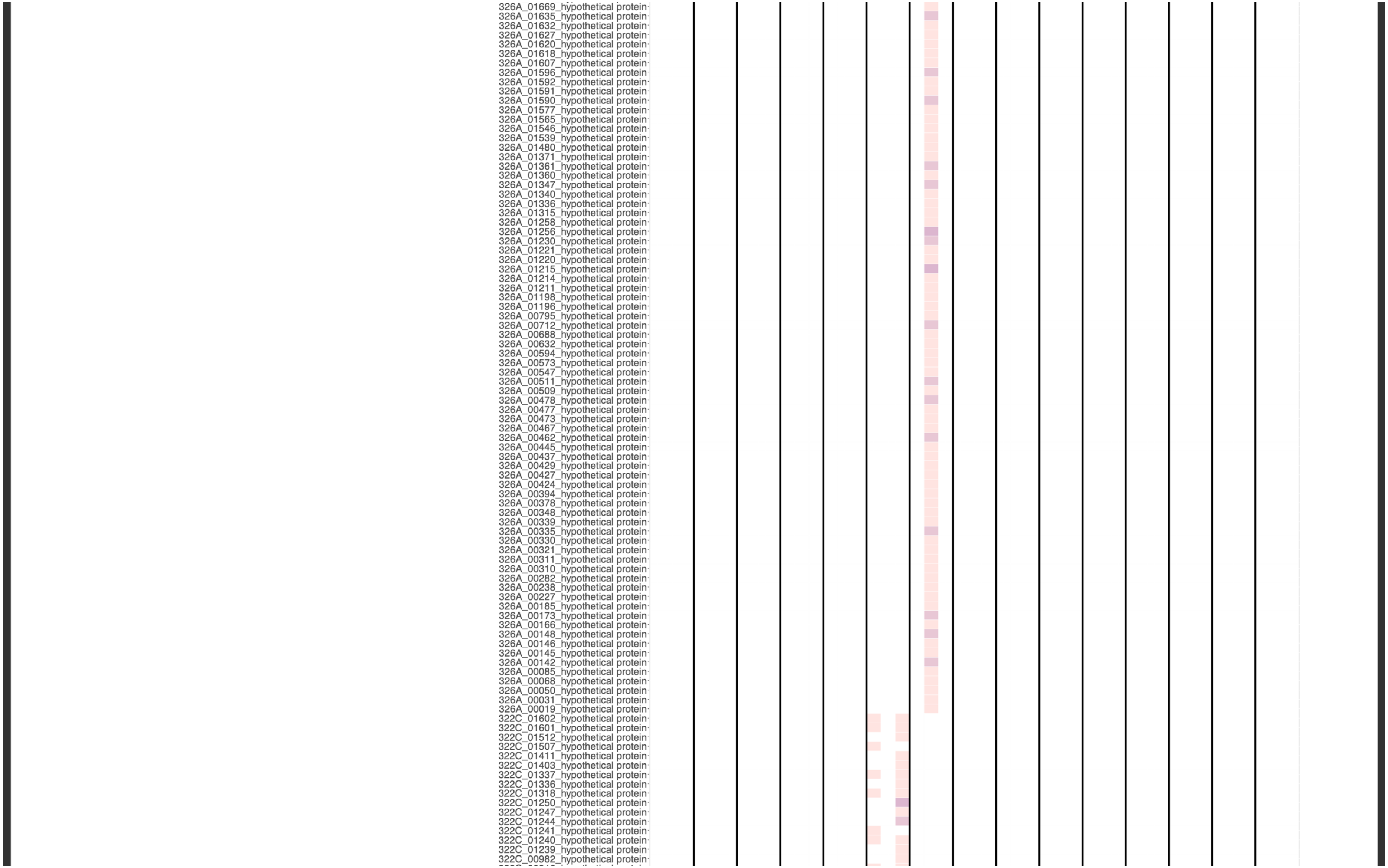
326A_01669_hypothetical protein
326A_01635_hypothetical protein
326A_01632_hypothetical protein
326A_01627_hypothetical protein
326A_01620_hypothetical protein
326A_01618_hypothetical protein
326A_01607_hypothetical protein
326A_01596_hypothetical protein
326A_01592_hypothetical protein
326A_01591_hypothetical protein
326A_01590_hypothetical protein
326A_01577_hypothetical protein
326A_01565_hypothetical protein
326A_01546_hypothetical protein
326A_01539_hypothetical protein
326A_01480_hypothetical protein
326A_01371_hypothetical protein
326A_01361_hypothetical protein
326A_01360_hypothetical protein
326A_01347_hypothetical protein
326A_01340_hypothetical protein
326A_01336_hypothetical protein
326A_01315_hypothetical protein
326A_01258_hypothetical protein
326A_01256_hypothetical protein
326A_01230_hypothetical protein
326A_01221_hypothetical protein
326A_01220_hypothetical protein
326A_01215_hypothetical protein
326A_01214_hypothetical protein
326A_01211_hypothetical protein
326A_01198_hypothetical protein
326A_01196_hypothetical protein
326A_00795_hypothetical protein
326A_00712_hypothetical protein
326A_00688_hypothetical protein
326A_00632_hypothetical protein
326A_00594_hypothetical protein
326A_00573_hypothetical protein
326A_00547_hypothetical protein
326A_00511_hypothetical protein
326A_00509_hypothetical protein
326A_00478_hypothetical protein
326A_00477_hypothetical protein
326A_00473_hypothetical protein
326A_00467_hypothetical protein
326A_00462_hypothetical protein
326A_00445_hypothetical protein
326A_00437_hypothetical protein
326A_00429_hypothetical protein
326A_00427_hypothetical protein
326A_00424_hypothetical protein
326A_00394_hypothetical protein
326A_00378_hypothetical protein
326A_00348_hypothetical protein
326A_00339_hypothetical protein
326A_00335_hypothetical protein
326A_00330_hypothetical protein
326A_00321_hypothetical protein
326A_00311_hypothetical protein
326A_00310_hypothetical protein
326A_00282_hypothetical protein
326A_00238_hypothetical protein
326A_00227_hypothetical protein
326A_00185_hypothetical protein
326A_00173_hypothetical protein
326A_00166_hypothetical protein
326A_00148_hypothetical protein
326A_00146_hypothetical protein
326A_00145_hypothetical protein
326A_00142_hypothetical protein
326A_00085_hypothetical protein
326A_00068_hypothetical protein
326A_00050_hypothetical protein
326A_00031_hypothetical protein
326A_00019_hypothetical protein
322C_01602_hypothetical protein
322C_01601_hypothetical protein
322C_01512_hypothetical protein
322C_01507_hypothetical protein
322C_01411_hypothetical protein
322C_01403_hypothetical protein
322C_01337_hypothetical protein
322C_01336_hypothetical protein
322C_01318_hypothetical protein
322C_01250_hypothetical protein
322C_01247_hypothetical protein
322C_01244_hypothetical protein
322C_01241_hypothetical protein
322C_01240_hypothetical protein
322C_01239_hypothetical protein
322C_00982_hypothetical protein

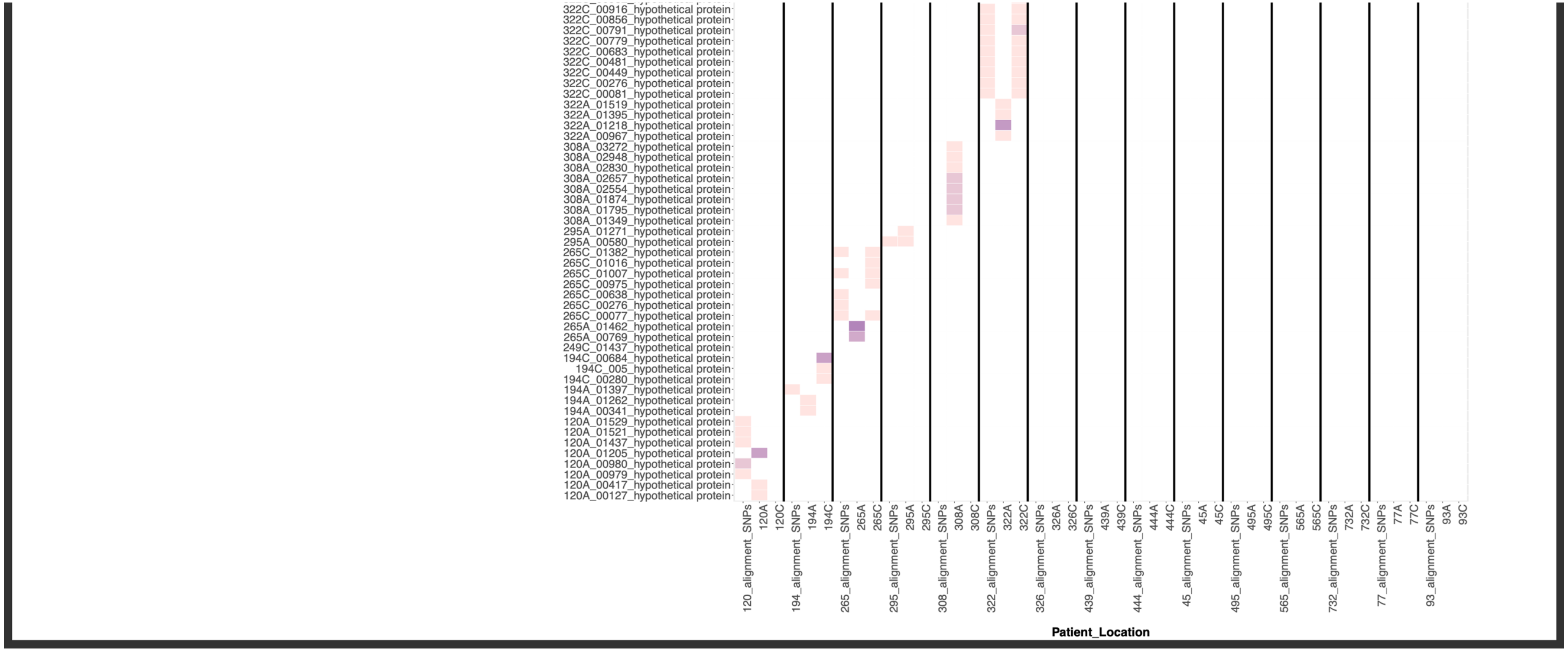


**Supp. Fig. 19**. Heat map showing the most variable genes and populations identified by a combination of analytical approaches (within and between bacterial populations from the antrum and corpus regions of patient stomachs). Each column contains data from one patient. Within each column, three datasets are shown. From left to right these are: between stomach region variation from whole genome alignment antrum versus corpus; within antrum minor allele variation; within corpus minor allele variation. Darker colour intensity indicates a larger number of variant bases within that gene/gene product. Only patients with paired antrum and corpus data were included. This figure combines the information presented in Fig. 3B plus Suppl. Fig. 17 (detection of minor allelic variants within each stomach region by read-mapping of deep sequencing reads back to the consensus genome) and Fig. 2A-B (detection of validated ‘red’ SNPs by whole genome alignment between stomach regions) plus Suppl. Fig. 2-15.

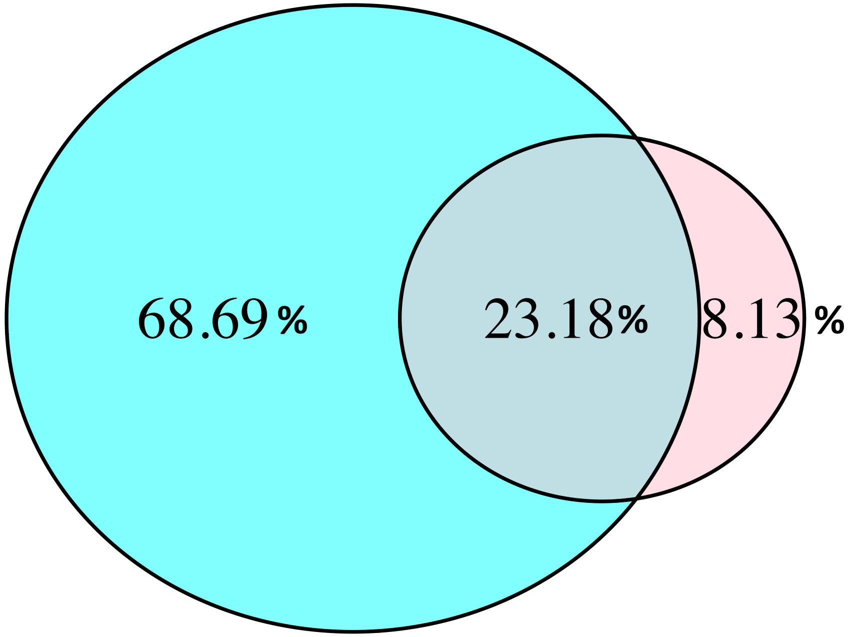


**Suppl. Fig. 20**. Venn diagram showing the percentage of allelic variant sites identified in this study by deep sequencing (blue) and single colony sequencing (pink) pipelines. Deep sequencing of the whole population detected the majority of the population diversity and was more powerful than the single colony method. However, some diversity (8.13%) was not identified by the deep sequencing dataset, presumably due to the stringent filtering and quality control methods we used to minimise identification of false positives.
